# Median based absolute quantification of proteins using Fully Unlabelled Generic Internal Standard (FUGIS)

**DOI:** 10.1101/2021.06.28.450203

**Authors:** Bharath Kumar Raghuraman, Aliona Bogdanova, HongKee Moon, Ignacy Rzagalinski, Eric R. Geertsma, Lena Hersemann, Andrej Shevchenko

## Abstract

By reporting molar abundances of proteins, absolute quantification determines their stoichiometry in complexes, pathways or networks. Typically, absolute quantification relies either on protein-specific isotopically labelled peptide standards or on a semi-empirical calibration against the average abundance of peptides chosen from arbitrary selected proteins. In contrast, a generic protein standard FUGIS (for <u>F</u>ully <u>U</u>nlabelled <u>G</u>eneric <u>I</u>nternal <u>S</u>tandard) requires no isotopic labelling, chemical synthesis or external calibration and is applicable to quantifying proteins of any organismal origin. The median intensity of peptide peaks produced by the digestion of FUGIS is used as a single point calibrant to determine the molar abundance of any co-digested protein. Powered by FUGIS, median based absolute quantification (MBAQ) outperformed other available methods of untargeted proteome-wide absolute quantification.

## Introduction

Proteomics envelopes multiple workflows for relative or absolute quantification of individual proteins. Relative quantification determines how the abundance of the same protein changes across multiple conditions on a proteome-wide scale. In contrast, absolute quantification determines the exact molar quantity of each protein in each condition. In this way, it is possible to relate the molar abundances of different proteins, estimate their expression level or determine their stoichiometry within a variety of molecular constellations from stable complexes to organelles or metabolic pathways and interaction networks^1–10^. Absolute quantification holds an important promise to deliver reference values of individual proteins in liquid and solid biopsies, which is a pre-requisite for robust molecular diagnostics.

A broad repertoire of absolute quantification techniques tailored towards analytical platforms, biological contexts and aims was developed^11–13^. It is usually presumed that the average abundance of a few representative peptides faithfully represents the abundance of corresponding source protein. In turn, peptides quantification relies either on isotopically labelled standards having exactly the same sequence or on a semi-empirical calibration against the abundances of selected (or, alternatively, of all detectable) peptides originating from arbitrary chosen standard proteins^12^,^14^. Targeted approaches are more accurate, yet they only cover a small selection of proteins that cannot be changed during the experiment. The latter methods work proteome-wide, however they rely on arbitrary assumptions and their accuracy is biased by experiment conditions and properties of individual proteins.

AQUA^15^ uses a set of isotopically labelled synthetic peptide standards identical to proteotypic peptides from endogenous proteins. Alternatively, QconCAT^16^, PSAQ^17^, PrEST^18^, PCS^19^, MEERCAT^20^, DOSCAT^21^ and GeLC-based MS Western^22^ employ metabolically labelled protein chimeras that, upon proteolytic cleavage, produce the desired peptide standards. MS Western relies on quantifying multiple proteotypic peptides per protein and validates the concordance of proteins determinations by monitoring the intensity ratios between XIC peaks of standards and corresponding endogenous peptides. Common discrepancies in these ratios point a unreliable quantification are typically due to miscleaved peptides or unexpected post-translational modifications.

To circumvent isotopic labelling, MIPA^23^ and SCAR^24^ standards use minimal sequence permutation or scrambling. It is assumed that scrambled and endogenous peptides share key physicochemical properties that results in equal instrument response^25^,^26^, which depends on analytical conditions and requires extensive validation.

Advances in robust and reproducible LC-MS/MS have led to a notion that generic measures of proteins molar abundance could be deduced either from raw intensities or spectral counts of peptide peaks, *e.g.* emPAI^27^, APEX^28^, SCAMPI^29^,^30^. Methods like Top3/Hi-3^6^, iBAQ^31^, Proteomic Ruler^32^,xTop^33^ and Pseudo-IS^34^ use averaged XIC intensities of selected or of all peptides matching the protein of interest. Because of limited inter-laboratory consistency, they are mostly used for supporting conventional proteomics workflows.

Hence, there is a need in developing a technology combining the accuracy and precision of the internal standards-based targeted quantification with broad (potentially, proteome-wide) coverage and ease of use of untargeted methods. To this end, we developed an untargeted proteome-wide quantification workflow termed **<u>m</u>**edian **<u>b</u>**ased **<u>a</u>**bsolute **<u>q</u>**uantification (MBAQ) that rely upon a **<u>f</u>**ully **<u>u</u>**nlabelled **<u>g</u>**eneric **<u>i</u>**nternal **<u>s</u>**tandard (FUGIS) based on common physicochemical properties of proteotypic peptides.

## Materials and Methods

### Protein extraction from HeLa cells

HeLa Kyoto cells were cultured in Dulbecco’s modified Eagle’s medium supplemented with 10% fetal calf serum and 1% penicillin-streptomycin (Gibco™ Life Technologies). HeLa cells were trypsinized, counted and washed 2x with PBS, before 1 ×10^6^ cells were lysed 30 mins on ice in either 1 mL or 0.5 mL RIPA buffer containing CLAAP protease inhibitors cocktail (10 μg/ml aprotinin, 10 μg/ml leupeptin, 10 μg/ml pepstatin, 10 μg/ml antipain and 0.4 mM phenylmethylsulfonyl fluoride (PMSF)). Subsequently, cells were further lysed by passing them 10 times through a 25g syringe. A post-nuclear supernatant was obtained from a 15 min centrifugation at 14.000 g. The supernatant was used for the further analysis by GeLC-MS/MS (**Supplementary information, GeLC-MS**) with both MS-Western standard and FUGIS standard in separate experiments.

### Absolute quantification of HeLa proteins using MS Western

Absolute protein quantification was performed using MS Western protocol^22^. The total protein content from HeLa cells from both dilutions were loaded on to a precast 4 to 20% gradient 1-mm thick polyacrylamide mini-gels purchased from Anamed Elektrophorese (Rodau, Germany) for 1D SDS PAGE. Separate gels were run for 1 pmol of BSA and isotopically labelled lysine (K) and arginine (R) incorporated chimeric standard containing 3-5 unique quantitypic peptides from target proteins The sample was cut into 3 gel fractions and each fraction was co-digested with known amount of BSA and the chimeric standard using Trypsin Gold, mass spectrometry grade, (Promega, Madison). The digest was analysed using GeLC-MS/MS workflow (**Supplementary information, GeLC-MS**). The peptide matching and chromatographic peak alignment from the raw files was carried out as described in (**Supplementary information, Database search and data processing**). The quantification was performed using the software developed in-house^9^.

### Absolute quantification of HeLa proteins using MBAQ and FUGIS

Similar to the MS Western experiments, the total HeLa cell lysate from both the dilutions were subjected to separation using 1D SDS PAGE. Separate gels were run for 1 pmol of BSA and the fully unlabelled generic internal standard (FUGIS). The sample was cut into three gel slices and each fraction was co-digested with the known amount of BSA and the FUGIS. The digests were again analysed using the GeLC-MS/MS workflow (**Supplementary information, GeLC-MS/MS**). The on-column amount of FUGIS was 200 fmol to 400 fmol; the loaded amount of chimeric proteins CP01 and CP02 (**Supplementary information, Expression and metabolic labelling of protein standards**) was 300 fmol. Peptide matching and chromatographic peak alignment were carried out as described in (**Supplementary information, Database search and data processing**). The output .csv files with sequences of matched peptides and areas of their XIC peaks were further processed by GlobeQuant software.

### GlobeQuant software for MBAQ quantification

GlobeQuant software was developed as a stand-alone Java script based application using in-memory SQL database (https://github.com/agershun/alasql) for fast access and search in the CSV file. GlobeQuant runs on a Windows 7 workstation with 16 GB RAM and 4-cores processor. The .csv output from the Progenesis LC-MS v.4.1 (Nonlinear Dynamics, UK) with peptide ID’s and their respective raw XIC peak areas were used by GlobeQuant software. List of FUGIS peptides was provided as an input. The software calculates the molar amount of the FUGIS standard by using the scrambled-native BSA peptide pairs and then calculated the median peak area of FUGIS peptides using areas of their XIC peaks. The calculated molar amount of the FUGIS standard is equated to the median of XIC peaks area and further used as a single point calibrant.

For BestN quantification peptides were chosen from a pool of Top3 peptides by calculating the coefficient of variation of all possible combination of Best2 and Best3 by default. If a protein did not contain Top3 peptides the Top2 peptides were taken as BestN peptides. Proteins identified with one peptide were excluded from the quantification. The BestN combination with the lowest coefficient of variation (<20%) was taken and averaged to provide the molar amounts of the protein. The software package is available at https://github.com/bharathkumar91/GlobeQuant.

## Results and Discussion

### MBAQ workflow for absolute quantification

MBAQ (for **<u>m</u>**ean **<u>b</u>**ased **<u>a</u>**bsolute **<u>q</u>**uantification) workflow relies on a recombinant protein standard consisting of concatenated peptides, whose sequences emulate physicochemical properties shared by typical proteotypic peptides. Its tryptic cleavage produces peptides in exactly equimolar concentration^16, 21,35^, as evidenced by the time course and relative abundances of rendered peptides^22^. Therefore, peptides concentration could be inferred from the known molar abundance of the chimera.

We therefore propose to determine the median value of areas of XIC peaks of peptides produced from chimeric protein and then use it a single point calibrant to calculate the molar abundance of other peptides from any co-digested protein. We note that proteotypic peptides included into the chimera protein standard are selected according to a few common rules, such as higher abundance of XIC peaks; no evidence of internal and also external miscleavages; no internal cysteine and methionine residues and no aspartic or glutamic acid residues at peptides N-termini^22^. We therefore hypothesized that peak areas corresponding to equimolar amount of proteotypic peptides released from the chimeric protein standard could cluster around some median value irrespective of their sequence. As compared to the targeted quantification by comparing the intensities of standard and analyte peaks, MBAQ can be less affected by biased yield of some peptide(s) because the abundance of all clustering peptides is used for calculating the median.

If so, we only have to: i) provide a sufficient number of such peptides to compute the robust median value under given experimental conditions; ii) select suitable peptides from those matched to proteins of interest and iii) check if the quantification of a target protein by its individual peptides is concordant. In our institute, we systematically produce large (40 to 270 kDa) protein chimeras comprising 40 to 250 proteotypic peptides from various proteins. To test the feasibility of MBAQ we further used CP01^9^ and CP02^4^ chimeras from our collection^22^ (**Supplementary information, Expression and metabolic labelling of protein standards**).

We first asked how areas of XIC peaks of proteotypic peptides chosen from different proteins and concatenated into a chimera are distributed around the median value and how many peptide would be required to estimate it with acceptable accuracy. To this end, we digested 267 kDa chimeric protein (CP01) comprising 250 proteotypic peptides selected from 53 *Caenorhabditis elegans* proteins^4^. Despite equimolar concentration of produced peptides, their peak areas differed by almost 10-fold (**Figure 1A**) (**Figure S1)**. However, the abundance of 48 % of peptides clustered near the median value (**Figure 1A; Figure S1)**. In order to ascertain that clustering does not depend on peptide sequence (again, these were all pre-selected proteotypic peptides from different proteins), we digested another 265 kDa chimera (CP02) harbouring proteotypic peptides from 48 proteins from *Drosophila melanogastet*^9^. We found that peak areas of 42 % of peptide were close to the median value (**Figure 1A; Figure S1**). We concluded that, independently of peptide sequences, approximately half of proteotypic peptides clustered around the same median, while others scattered around it. However, the commonality between peptide sequences within clustering and non-clustering groups was not immediately obvious.

**Figure 1.**
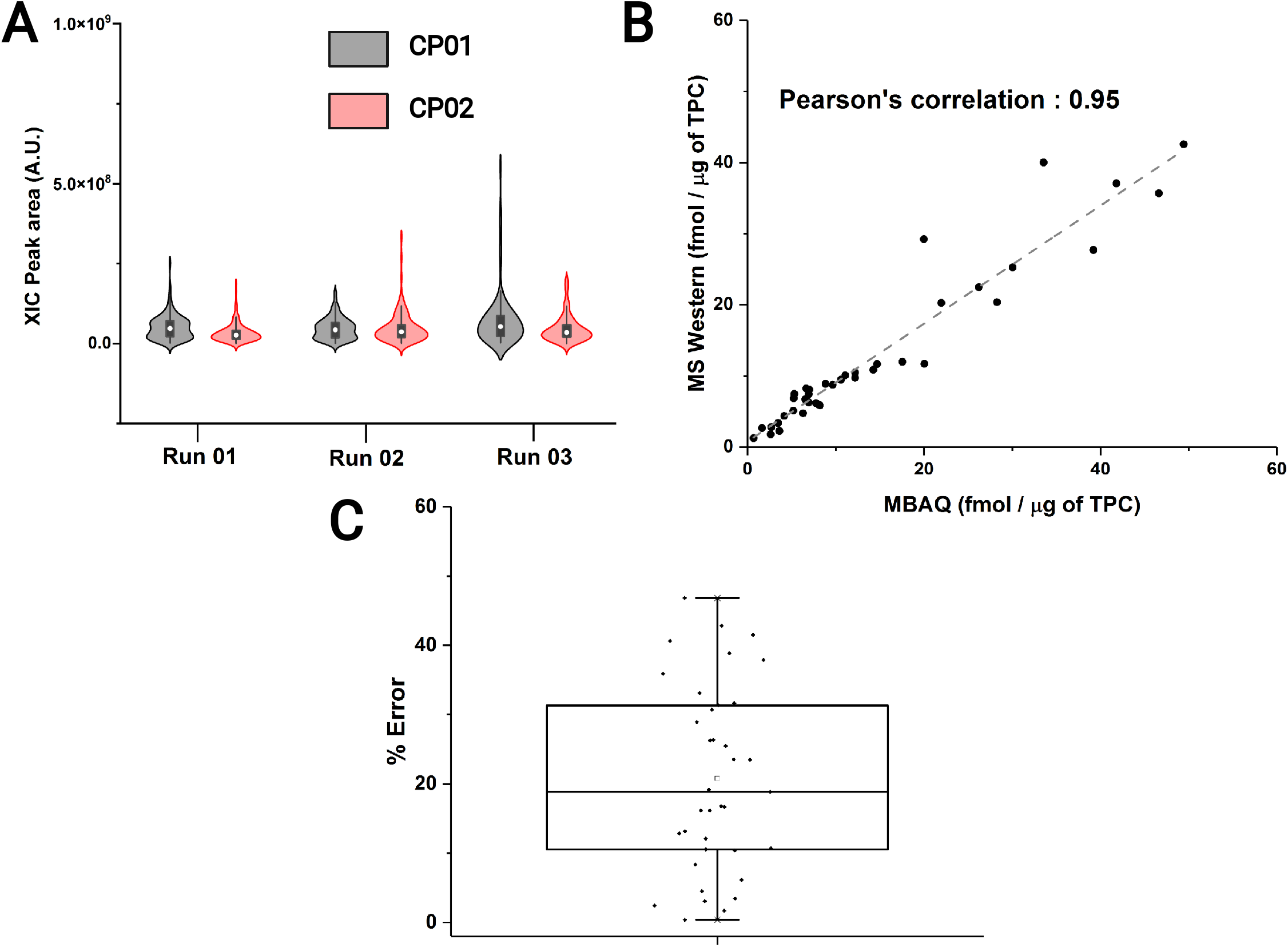
MBAQ Quantification. A: Distribution of XIC peak areas of peptides from chimeric proteins CP01 and CP02 in three independent chromatographic runs; B: Molar quantities of 48 metabolic enzymes from *Caenorhabditis elegans* quantified by MBAQ and MS Western; C: MBAQ quantification error (in %) relative to the values determined by MS Western with each data point signifying a protein

Since the “near-median” (NM) peptides were evenly distributed across the retention time range (**Figure S2**), we checked if the median value could faithfully represent the molar abundance of the chimera. We expect that, in this case, possible suppression of peptides ionization by a sample matrix would be likely randomized, compared to a hypothetical scenario if all peptides would be eluting together. For this purpose, we used the CP01 to quantify 48 metabolic enzymes from *Caenorhabditis elegans* by the MS Western protocol and, independently, by using a median value computed from the abundances of all CP01 peptides. We underscore that in MS Western workflow each enzyme was quantified using several isotopically labelled peptide standards that exactly matched sequences of corresponding native peptides^4^ with no recourse to other peptides. In contrast, in MBAQ workflow all peptides from the digested chimera were taken for calculating a single median value that was subsequently used for quantifying all proteins. The MBAQ was concordant with MS Western showing Pearson’s correlation of 95 % (**Figure 1B**) and median quantification error of 18 % (**Figure 1C**) within 3 orders of magnitude of molar abundance difference.

In a separate experiment, we quantified 30 proteins from the commercially available UPS2 protein standard (Sigma Aldrich, USA) using MBAQ and the median calculated from CP01 peptides. The Pearson’s correlation was 96 % and the median quantification error was less than 20 % (**Table S1**).

We therefore concluded that, if a sufficient number of equimolar prototypic peptides are detected by LC-MS/MS, their median abundance is invariant to their exact sequences and unaffected by other peptides included into the chimera. The use of median abundance as a single point calibrant delivers good quantification accuracy that is close to the accuracy of targeted quantification relying on identical peptide standards.

Though MBAQ workflow was accurate, the use of a large isotopically labelled CP deemed unnecessary. Effectively, we only used less than a half of its peptides and took no advantage of isotopic labelling, except for validating MBAQ by independent quantification of same proteins by MS Western. Therefore, we sought to design a generic (suitable for all proteins from all organisms) and fully unlabelled internal standard (FUGIS).

### Development of FUGIS

FUGIS was conceived as a relatively small protein chimera composed of concatenated proteotypic-like tryptic peptides that, however, share no sequence identity to any known protein. It also comprises a few reference peptides with close similarity to some common protein standard *e.g.* BSA. Upon co-digestion with quantified proteins, FUGIS should produce equimolar mix of peptide standards whose median abundance would support one-point MBAQ of all co-detected peptides from all proteins of interest. The exact amount of FUGIS is determined by comparison with the known amount of co-digested reference protein (here, BSA) in the same LC-MS/MS experiment.

We first asked, what is the minimum number of peptides required to arrive to a consistent median value? For this purpose, we performed a bootstrapping experiment over the abundances of tryptic peptides derived from CP01 and CP02. Median values were calculated by repetitive selection of defined (3 to 120) number of peptides (**Figure 2**). The data collected by 100 bootstrap iterations suggested that a consistent median value can be projected by considering peak areas of as little as 5 to 10 peptides. However, the medians spread (which depends on “internal” peptide properties and “external” conditions of ionization) decreased with the number of peptides and reached plateau at more than 30 peptides (**Figure 2A, B**). Bootstrapping also confirmed that, irrespective of peptides selection, same peptides tend to cluster around the median. The abundance of 32 % out of 230 peptides further termed as near-median (NM) peptides was within the range of 20 % of the median value. Therefore, for further work we selected 70 peptides, whose peak areas were also close to the median in several technical LC-MS/MS replicates.

**Figure 2.**
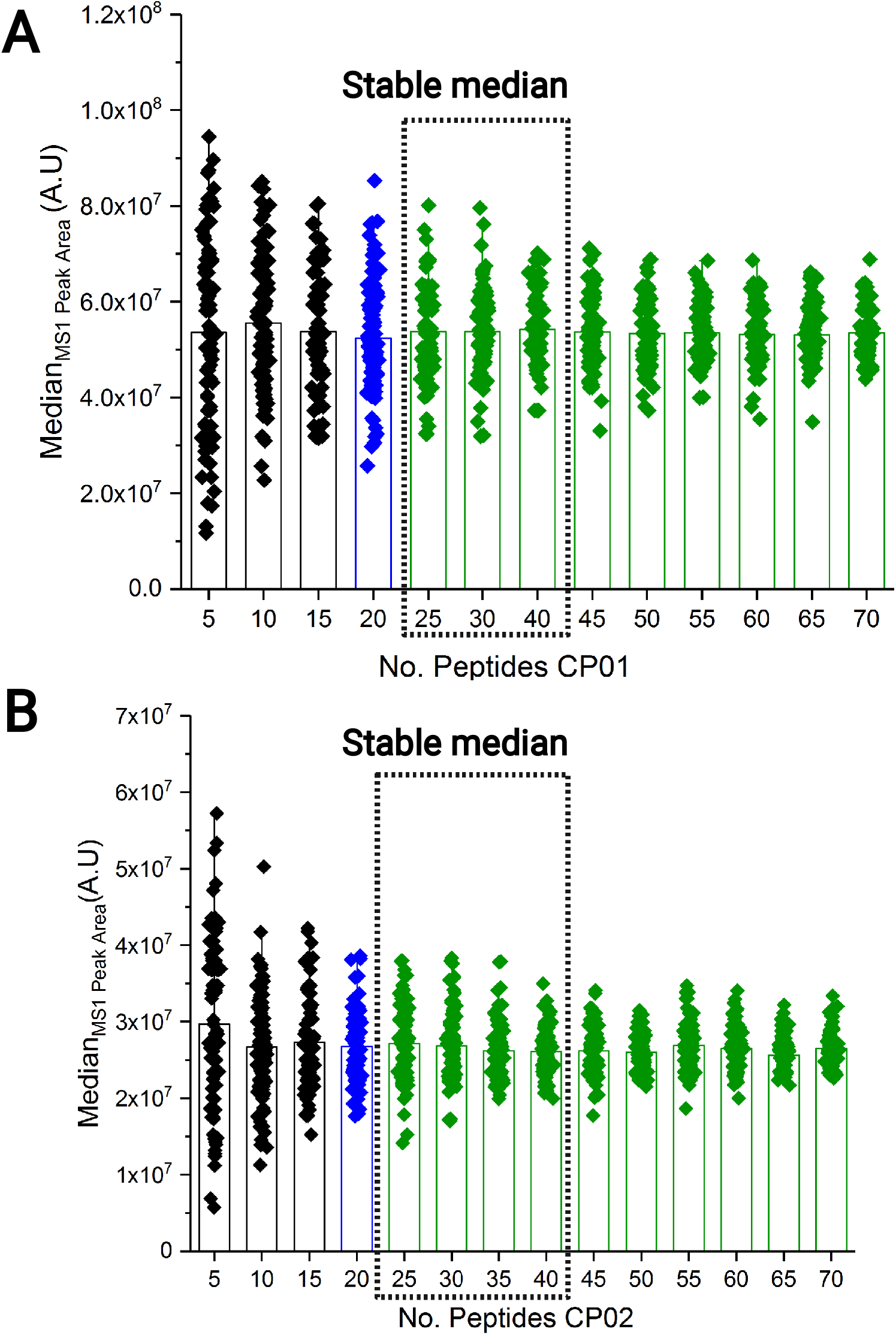
Minimum number of peptides for robust estimation of the median value. Bootstrapping of XIC peak areas of peptides from A: CP01 and B: CP02 over the range of 3 to 120 peptides in the total of 100 iterations. Filled diamonds represent median values determined by each bootstrapping iteration. Green bars represent the peptide number with stable median.

Next, we altered sequences of these near-median (NM) peptides such that they become different from any known sequence. Yet, we tried to preserve the similarity of their physicochemical properties, such as net charge, hydrophobicity index and location of polar (including C-terminal arginine or lysine) amino acid residues as compared to corresponding “source” peptides. We noted that the examination of sequences of NM and other peptides did not reveal an unequivocal rationale behind such grouping and therefore we tried several ways to alter their sequences.

We first selected a set of 40 out of total of 70 NM peptides and reversed their amino acid sequences (**Figure 3A**) except C-terminal lysine or arginine and assembled them into a chimeric protein GCP01 (**Table S2**) that was expressed and metabolically labelled with ^13^C^15^N-Arg and ^13^C-Lys in *Escherichia coli*^22^. Its band was excised from 1D SDS PAGE, co-digested with the band of 1 pmol BSA and analysed by LC-MS/MS^22^. Similar to previously published strategy^37^, the peptide abundances were normalised to the abundance of BSA peptides in the chimeric protein to check if the normalised median abundance (NMA) is close to unity (~1.0). A unit NMA means that the median abundance truly represents the amount of the FUGIS standard while any deviation contributes to the error in quantification.

**Figure 3.**
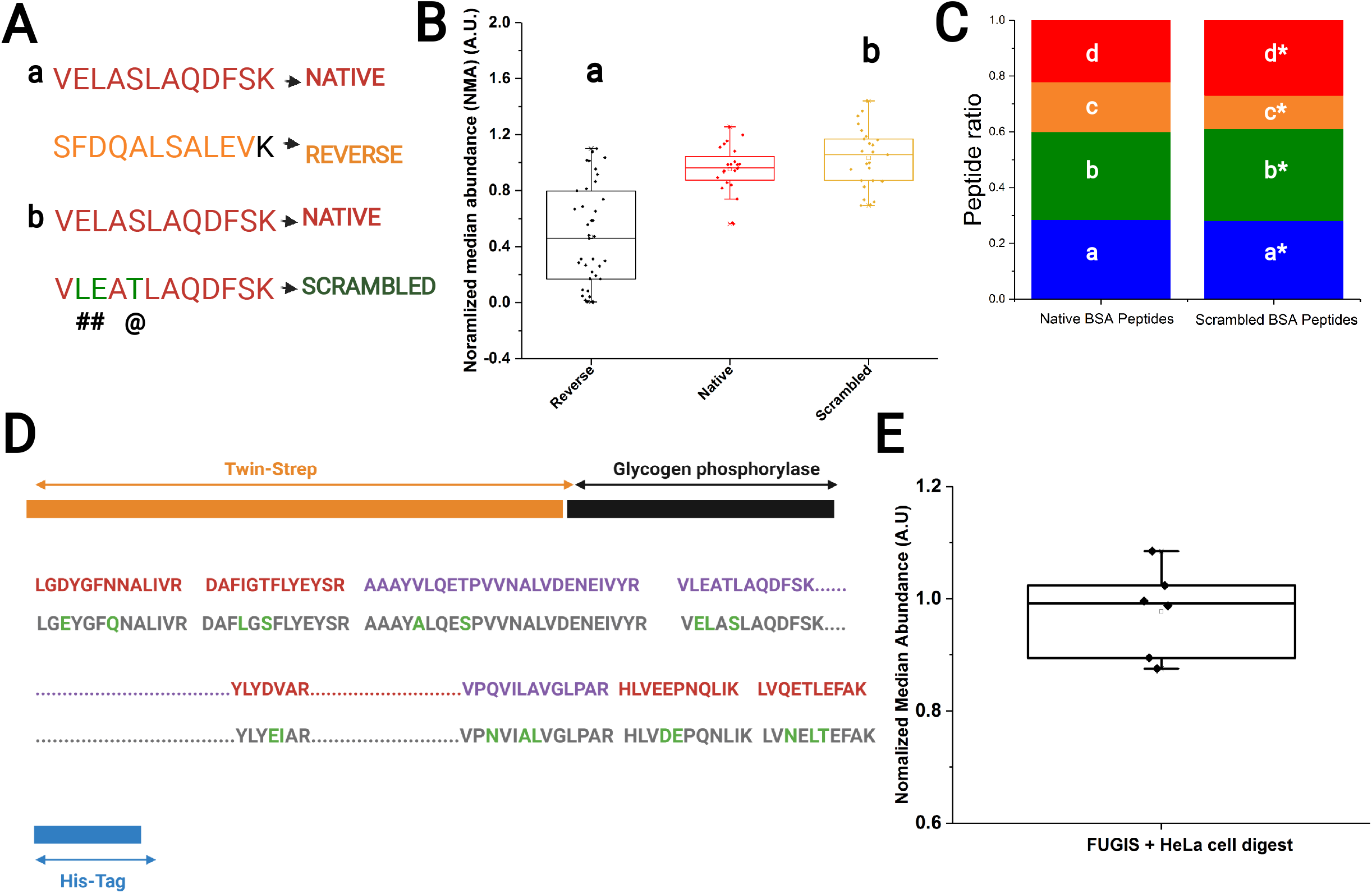
Design of FUGIS. A: Examples of reversing (a) and scrambling (b) of peptide sequences; # indicates a swap and @ indicates substitution of amino acid residues. B: Normalised median abundance (NMA) of reversed, native and scrambled peptide sequences. Each data point is a peptide. C: Distribution of relative abundances (peptide ratios) of native and scrambled peptides. Asterisks indicate scrambled sequences: a / a* – HLVDEPQNLIK / HLVEEPNQLIK; b / b* – LGEYFGQNALIVR / LGDYGFNNALIVR; c / c* – YLYEIAR / YLYDVAR; d / d* – DAFLGSFLYEYSR / DAFIGTFLYEYSR). D: Schematic diagram of th designed sequence of 80 kDa FUGIS protein. Sequence stretches in red are scrambled BSA peptides; in grey: native peptides; swap or substitution of amino acid residues id in green. Full-length sequence of FUGIS is in **Figure S2A**. E: NMA of FUGIS peptides in HeLa cell background. Each data point is technical replicate.

The NMA for the reversed sequences was 0.45 (**Figure 3B**), which was very far from the NMA of their native counterpart of 0.97. Thus, we concluded that reversing peptide sequences strongly biases the median, increases the spread and therefore it should not be used for designing a FUGIS chimera.

Next, we scrambled the peptide sequences by introducing point substitutions of amino acid residues. We allowed a maximum of two scrambling events per peptide that followed two intuitive rules. First, in each peptide only two amino acid residues were swapped (**Figure 3A**). Second, to create a mass shift, an amino acid residue preferably located in the middle of the peptide sequence was substituted with another amino acid having similar side chain *(e.g.,* Ser to Thr or vice versa) (**Figure 3A**). To minimize the retention time shift, aliphatic amino acids in the order of increasing hydrophobicity (G<A<V<L<I) were only substituted with an amino acid having similar hydrophobicity *(i.e.* substitutions V by L were allowed, but G by I were not). Altogether, 20 scrambled sequences together with corresponding 20 source “native” peptides were assembled into a chimera GCP02 (**Table S3**). Pairwise comparison of peak areas of native and scrambled sequences suggested that they differed by less than 5 %. Similar to GCP01, we calculated the NMA for peptides in GCP02. Scrambled peptides behaved similar to the native sequences with a NMA of 1.02 (**Figure 3B**). On average, the retention time difference between native and scrambled peptides was 3.21 (± 2.02) minutes. Therefore, these scrambled peptide sequences were selected for the FUGIS.

Isotopic labelling of GCP01 and GCP02 chimeras was unavoidable since their quantification was dependent on the reference BSA peptides. We found that reference BSA peptides scrambled in the same way behaved similar to the native peptides with retention time shift of 1.2 (± 0.5) minute. Also the relative abundances (peptide ratios) ^22^ of corresponding native and scrambled BSA peptide were very similar (**Figure 3C**). Therefore, metabolic labelling of a scrambled chimera was not any longer required.

Taken together, we designed and produced a FUGIS chimera having the molecular weight of 79.01 kDa (**Figure 3D**; **Figure S3**; **Table S4**), which harbours 43 scrambled near-median peptides and 5 sequences of scrambled reference peptides from BSA. All peptides showed no complete match to any protein sequence across organisms (**Table S4**).

### MBAQ quantification using FUGIS

To assess the feasibility and accuracy of MBAQ quantification using FUGIS we quantified 4 proteins from 1 million HeLa cells at 2 dilutions and compared it with the quantities previously determined using MS Western^22^. Since MBAQ quantification is based on the median abundance, we wanted to assess the accuracy of the median estimation in different matrix background. To this end, we pre-fractionated both dilutions of a HeLa cells lysate by 1D-SDS PAGE and excised 3 bands from each gel, which were co-digested with bands of 1 pm of BSA and FUGIS. Irrespective of protein background NMA calculated for FUGIS was 0.98 with less than 10 % error (**Figure 3E**).

We then proceeded to quantify the molar amounts of 4 proteins (PLK-1, TBA1A, CAT, G3P) from HeLa cells using MBAQ, MS Western^22^ and Top3/Hi-3 quantification^6^ (**Figure 4, Table S5 and Table S6**). We observed that molar abundances determined by MBAQ were much closer to MS Western than of Top-3/Hi-3.

**Figure 4.**
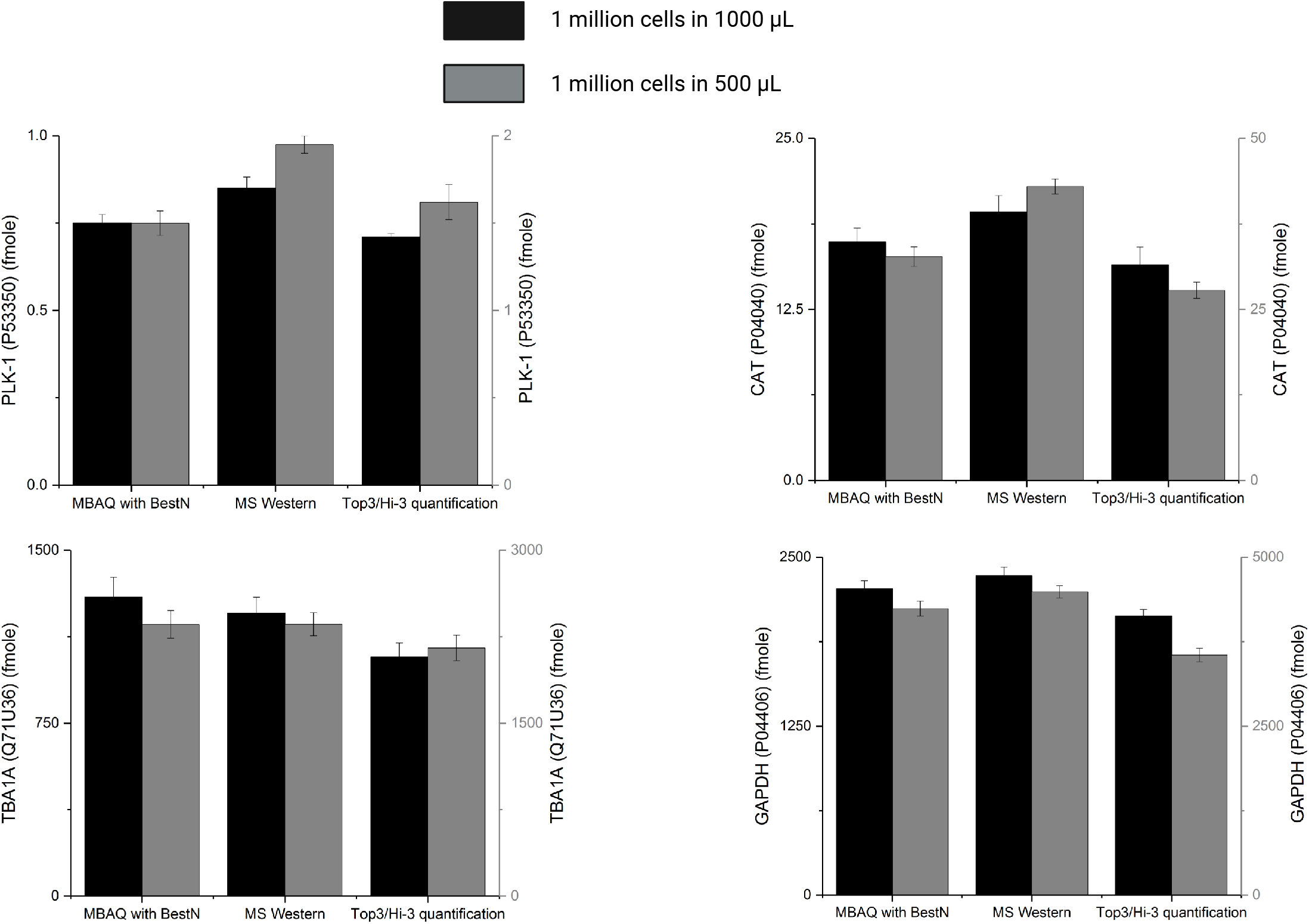
Molar quantities of proteins determined by MBAQ, MS Western and Hi3 quantification. MBAQ vs MS Western vs Top3/Hi3 quantification of PLK-1, CAT, G3P, TBA1A proteins from HeLa cell lysate and from its 2-fold dilution. Error bars represent +/− SEM of technical replicates.

For MBAQ in each target protein we selected peptides whose mean and median values differed by less than 15 %. We termed them as “BestN” peptides – in contrast to TopN peptides that corresponded to N most abundant peptides. To assess if BestN peptides delivered better accuracy, we looked into the quantification of one of the four proteins (glyceraldyhyde-3 phosphate dehydrogenase; G3P Human P04406) (**Figure 5A; Table S5; Table S6**). We estimated the concordance of its molar amount independently calculated from multiple peptides by the coefficient of variation (%CV)^9^. If calculated from BestN peptides it was 7%, which is significantly better than Hi-3 quantification (18 %) (**Figure 5B; Figure S4; Table S5; Table S6**).

**Figure 5.**
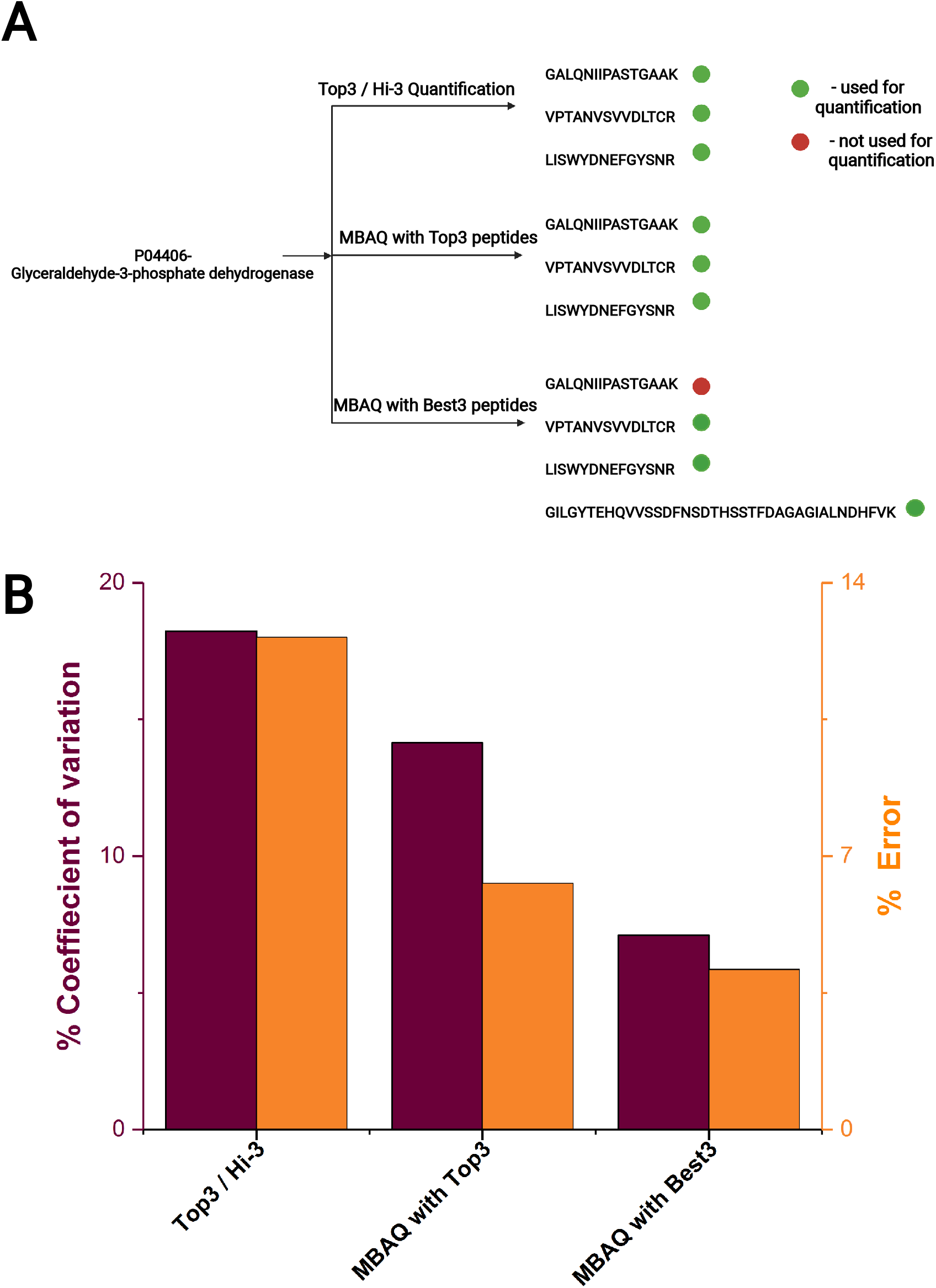
Molar quantification of human G3P protein by MBAQ and Top3 / Hi-3 methods. A: Selection of proteotypic peptides for each method. XIC peak areas of peptides are in **Figure S3**. B: Coefficient of variation (%) and % error (relative to the values determined by MS Western).

To understand why BestN peptides improved the quantification accuracy we considered the difference between BestN and TopN peptide sets. For human G3P (P04406) and tubulin-1 alpha (Q71U36) most abundant peptides were excluded from BestN set that reduced CV down to less than 10 % (**Table S7**). For human catalase (P04040) and serine/threonine protein kinase (P53350) Top2 and Best2 peptides were the same (**Table S7**). Therefore, BestN peptides is a subset of TopN peptides and a minimum of two peptides is required to provide reliable molar amounts.

Considering MS Western estimates as “true values”, we evaluated the accuracy of MBAQ quantification. MBAQ with BestN peptides provided the most accurate quantification with the accuracy of 96% (**Figure 5B**). We also observed that, when used together with TopN, MBAQ performed better than Top3 quantification with the accuracy of 94 % (**Figure 5B**).

GlobeQuant software supports MBAQ workflow (**Figure 6A**) by selecting BestN peptides from analysed proteins and using FUGIS to provide a robust single point calibrant. We employed GlobeQuant to quantify 1450 proteins identified with two or more matching peptides in HeLa lysate and, independently, in its 2-fold diluted aliquot. In each sample, proteins were quantified independently with no recourse to raw intensities of chromatographic peaks in another sample. Molar quantities of individual proteins (**Table S8)** are plotted as a ranked cumulative abundance in **Figure 6B**.

**Figure 6.**
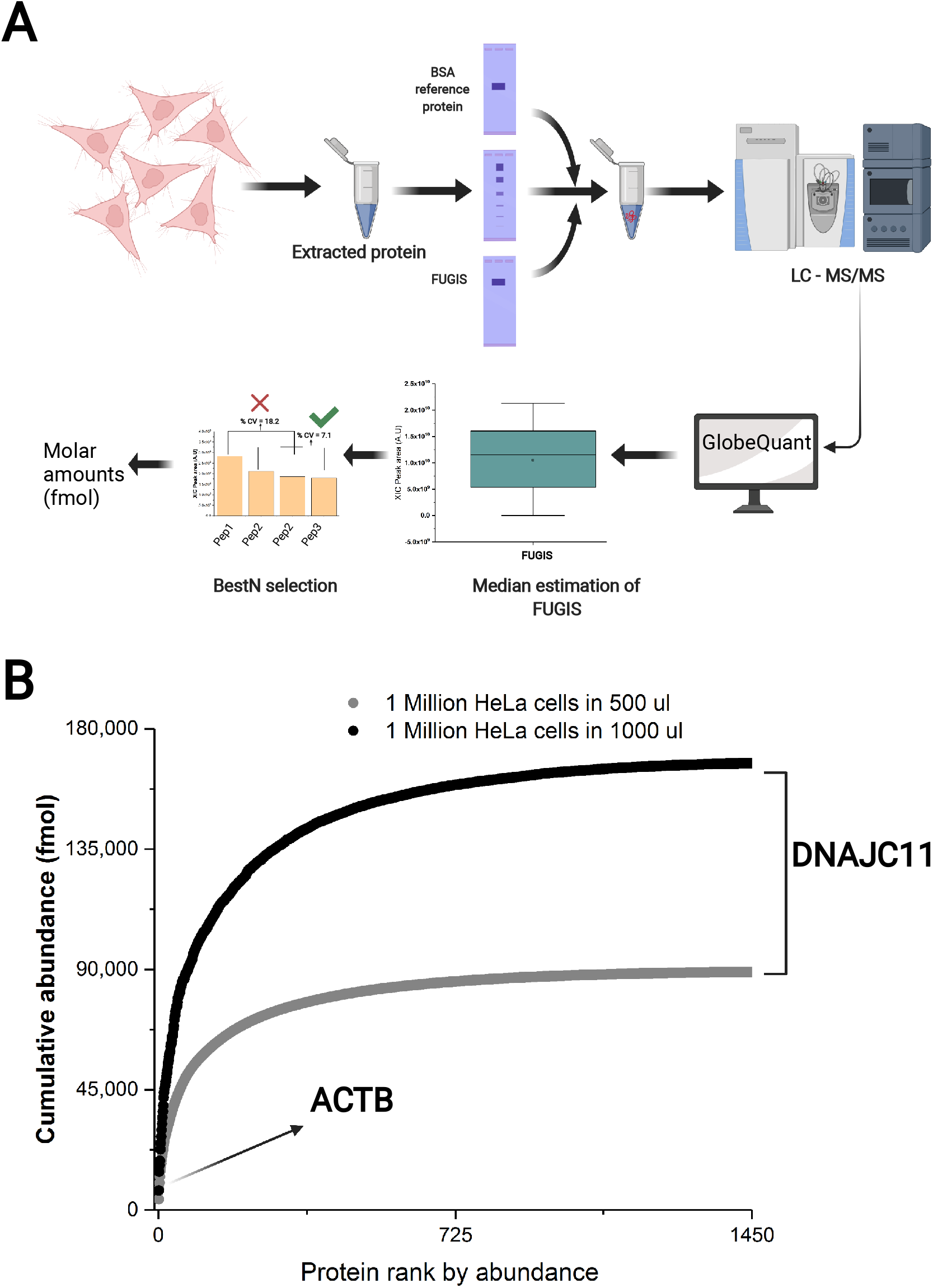
MBAQ quantification of HeLa proteome using GlobeQuant. A) Schematic representation MBAQ – GlobeQuant workflow. B) Ranked cumulative abundance of 1450 proteins from both dilutions of HeLa lysate with the least abundant protein at the right. ACTB was the most abundant protein.

MBAQ faithfully recapitulated the anticipated 2-fold difference with an average accuracy of 92%. Protein quantities (**Table S8**) provide useful resource for benchmarking of newly developed absolute quantification methods.

Finally, we checked if absolute quantification by MBAQ and by other proteome-wide techniques such as iBAQ and Proteomic Ruler is concordant. To this end, we converted molar abundance of the four HeLa proteins into the number of copies-per-cell and compared it with previous reports (**Table 1**). Copy numbers determined by two independent MS Western experiments were concordant and corroborated MBAQ. At the same time, MBAQ, iBAQ and Proteomic Ruler reported discordant quantities of the same four proteins, but also showed marginal concordance on the proteome-wide scale (**Table 1; Figure S5**). This is not surprising since both determinations by Proteomic Ruler do not correlate and are also discordant with iBAQ. Since MBAQ corroborated MS Western (**Table 1**), we argue that it provides more accurate estimate of molar abundance despite its apparent discordance to alternative methods.

**Table 1.**
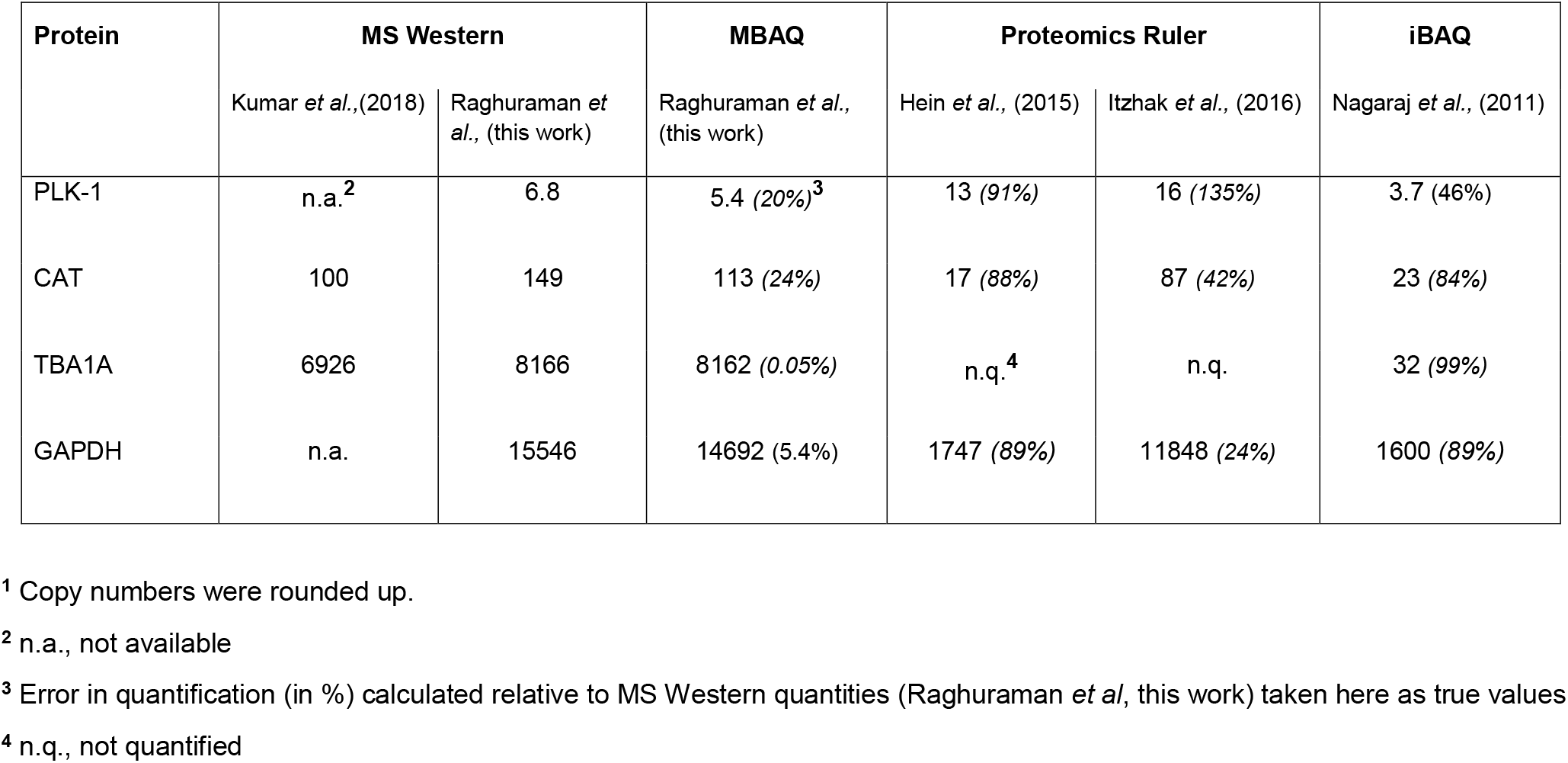
Number of copies-per-cell^1^ (×10^4^) in HeLa cells determined by MS Western^22^, MBAQ, Proteomics Ruler^32^ and iBAQ^31^.

### Conclusion and perspectives

We argue that, together with FUGIS standard, MBAQ workflow supported accurate absolute quantification of proteins at the proteome-wide scale. High level of expression in *E.coli,* good solubility and, last but not least, no interference with any known protein make FUGIS a preferred internal standard for label-free experiments aiming at the absolute but also relative quantification. Upon tryptic digestion, it produces 43 peptides in exactly known equimolar amount covering a common range of peptide retention times. Though the current workflow involves GeLC-MS/MS strategy, it can be easily adjusted for an in-solution protocols: since FUGIS is highly expressed in *E.coli* there is no need in its further purification.

It has long been noticed that the abundance of proteins could be inferred from the abundances of best detected (TopN) peptides, as in Hi-3 quantification^6^. However, relying on best ionized peptides biases its accuracy^33, 37, 38^. By selecting BestN (instead of TopN) peptides MBAQ improved the quantification consistency by disregarding peptides, whose ionization capacity is based on uniquely favourable sequence. It is also important that in MBAQ molar abundance of peptides is referred to a recognized commercial quantitative standard.

We speculate that employing MBAQ or similar quantification might be an important step towards establishing diagnostically relevant protein values in liquid and solid biopsies. MBAQ could quantify any protein detectable with multiple (three or more) peptides in any LC-MS/MS experiment, including data-independent acquisition (DIA). MBAQ does not rely on pre-conceived knowledge of the protein composition or availability of MS/MS spectra libraries.

Finally, charting the proteome and metabolome composition in molar quantities will facilitate our understanding of metabolic and signalling pathways that are controlled by molar ratios between multiple enzymes and substrates and help to uncover the molecular rationale of proteotype-phenotype relationships.

## Author Contributions

BKR and AS conceptualized and designed the FUGIS and MBAQ. BKR performed the experiment and interpreted the data. AB expressed the FUGIS standard. HKM and LH conceptualized and developed the software. BKR and AS wrote the manuscript. IR provided expert technical support and critical reading of the manuscript

## Acknowledgements

Work in AS group was supported by Max Planck Society. We acknowledge Dr. Sandra Segeletz for expert technical support. The figures were assembled using BioRender tool (created by BioRender.com)

## Data availability

Raw data have been deposited to the ProteomeXchange Consortium via the PRIDE partner repository with the dataset identifier PXD026886.

## SUPPORTING INFORMATION

The following supporting information is available free of charge at ACS website http://pubs.acs.org

## Supplementary methods

GeLC-MS

Expression and metabolic labelling of protein standards

Database search and data processing

Figure S 1. MS1 peak area of CP01 and CP02 in 3 independent chromatographic runs. Each data point represents a peptide.

Figure S 2. Retention times distribution of near-median peptides; the gradient elution profile is in grey.

Figure S 3. A) Full length sequence of FUGIS standard. B) Expressed FUGIS standard separated using 1D SDS PAGE. C) Retention time distribution of peptides in the FUGIS standard; the gradient elution profile is in grey.

Figure S 4. XIC peak area of Top4 peptides of GAPDH (P04406) with rational behind the selection of BestN peptides. %CV is the coefficient of variation of multiple selections.

Figure S 5. Comparison of copy numbers per HeLa cell for same proteins in: A) Nagaraj et al. (iBAQ)[1] vs. Raghuraman et al (this paper), B) Hein et al. (Proteomic Ruler)[2] vs. Raghuraman et al., C) Nagaraj et al. (iBAQ)[1] vs. Hein et al. (Proteomic Ruler)[2]; R is Spearmen correlation coefficient.

Table S 1. MBAQ quantification of UPS2 standard

Table S 2. Peptide sequence of GCP01

Table S 3. Peptide sequence of GCP02

Table S 4. Peptide sequence of FUGIS

Table S 5. MBAQ vs. MS Western vs. Hi-3 of 1 Million cells in 500 μl

Table S 6. MBAQ vs. MS Western vs. Hi-3 of 1 Million cells in 1000 μl

Table S 7. Best N peptides of 4 HeLa proteins

Table S 8. GlobeQuant quantification of 1450 HeLa proteins

Table S 9. MS data acquisition paratmeters

## For TOC onlyx

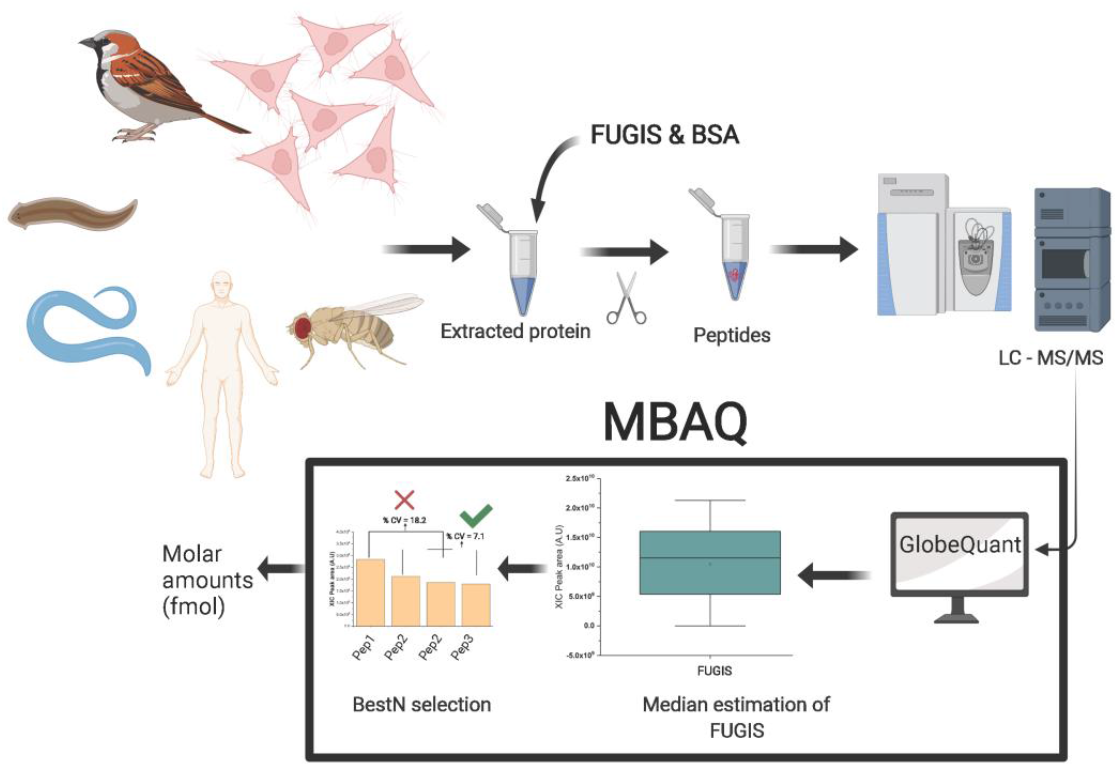

