## supplementary information for "Median based absolute quantification of proteins using Fully Unlabelled Generic Internal Standard (FUGIS)"

### Supplementary methods

#### GeLC-MS/MS

In-gel digestion workflow was adapted from the protocol by Shevchenko *et al* [1]. After electrophoresis the gel slab was stained with Coomassie Brilliant Blue R250 for 15 min at room temperature. After staining, gels were destained in 5:4:1 (v/v) of water: methanol: acetic acid. Slices corresponding to MW of interest were excised from the destained gel slab, cut into *ca.*1 mm size cubes and transferred to 1.5 ml LoBind Eppendorf tubes for further processing.

The gel pieces were destained completely by acetonitrile / water, and subsequently reduced by incubating the gels with 10 mM DTT at 56$^{o}$C for 45 minutes. After reduction proteins were alkylated using 55 mM of iodoacetamide for 30 minutes in dark at room temperature. The reduced and alkylated samples were digested overnight with trypsin (10 ng/µl) in 10 mM ammonium bicarbonate. After digestion tryptic peptides were recovered from gel matrix using water / acetonitrile / formic acid, dried under vacuum and stored at -20$^{o}$C for further analysis.

The dried extracts were reconstituted in 5% formic acid and 5 µl were injected into a Dionex Ultimate 3000 Nano-HPLC system equipped with two-column setup comprising of a trap column (5 mm × 300 µm i.d) and an analytical column (Acclaim PepMap100 C18 15 cm × 75 µm). Water with 0.1% (solvent A) and ACN with 0.1% (solvent B) were used as mobile phase. Samples were loaded into the trap column with a flow rate of 20 µl/min. Then the trap column was switched to the analytical column with a flow rate of 200 nL/min for peptides separation. The separation was carried out using Dionex Ultimate 3000 HPLC system (Thermo Scientific, Bremen,

Germany) running gradient elution program for 180 min and the output was hyphenated to Q-Exactive HF (Thermo Fisher Scientific, Bremen, Germany) and mass spectra were acquired in data-dependent acquisition mode. The acquisition parameters are provided in **Table S9**.

#### Expression and metabolic labelling of chimeric protein standards

We adapted the same expression workflow as described in [2]. Synthetic genes produced by GenScript (Piscataway NJ) were sub-cloned into pET expression vector and transformed into an *E.coli* strain that was dual auxotroph for arginine and lysine (ΔArgΔLysBL21 (DE3) T1 pRARE). Successful transformants were then diluted and sub cultured in MDAG-135 media [3]. They were induced by 0.2mM isopropyl β-d-1-thiogalactopyranoside (IPTG). After 4 to 6 hr post induction cells were pelleted, re-suspended in 2x phosphate-buffered saline (PBS). The suspended cells were then aliquoted, snap frozen in liquid nitrogen and stored at - 80°C until used for quantification.

#### Database searches and data processing

Peptide matching was carried out using Mascot v.2.2.04 software (Matrix Science, London, UK) against *Homo sapiens* (August 2020) proteome downloaded from Uniprot. A precursor mass tolerance of 5ppm and fragment mass tolerance of 0.03 Da was applied, fixed modification: carbamidomethyl (C); variable modifications: acetyl (protein N terminus), oxidation (M); labels: $^{13}C(6)$ (K) and $^{13}C(6)^{15}N(4)$ (R) (only for MS Western the labelling was used); cleavage specificity: trypsin, with up to 2 missed cleavages allowed. Peptides having the ions score above 15 were accepted (significance threshold $p < 0.05$). The chromatographic alignment and feature detection were carried using Progenesis LC-MS v.4.1 (Nonlinear Dynamics, UK). The absolute quantification was performed by calculating the abundances for the labelled and the unlabelled peptide using an in-house software as previously published [4]. The statistical analysis were carried out in OriginLab (2017) (OriginLab Corp., Northampton, Massachusetts, USA).

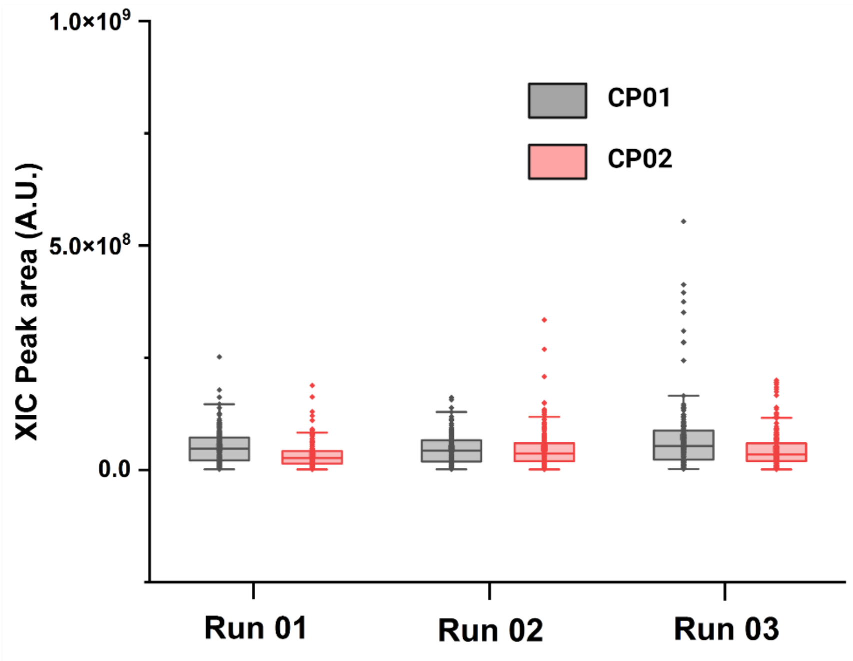

**FIGURE S 1. MS1 PEAK AREA OF CP01 AND CP02 IN 3 INDEPENDENT LC-MS/MS RUNS. EACH POINT REPRESENTS A PEPTIDE.**

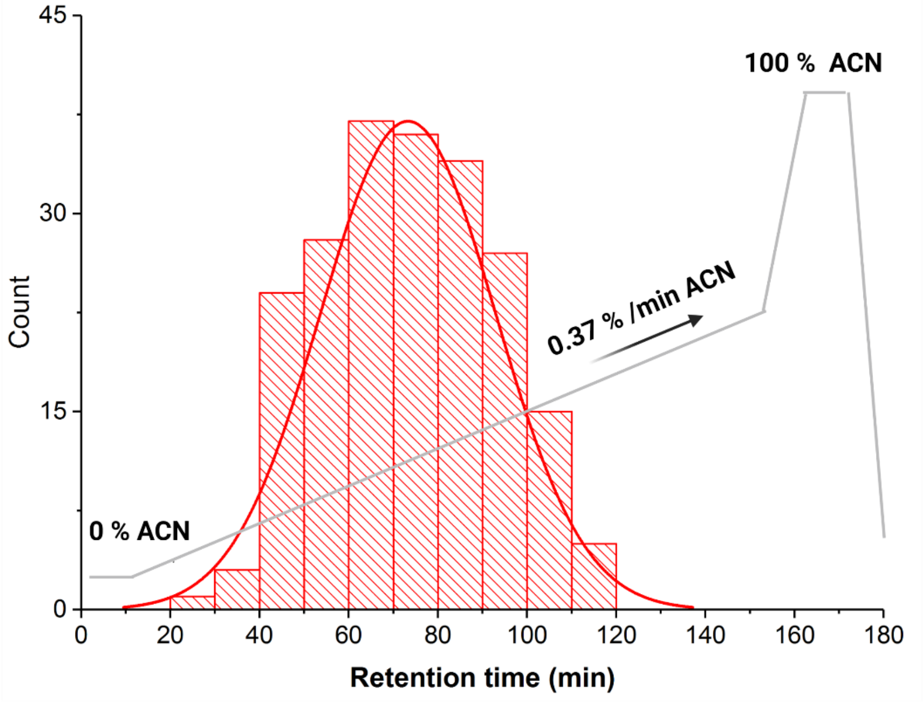

**FIGURE S 2 . RETENTION TIMES DISTRIBUTION OF NEAR MEDIAN PEPTIDES; GRADIENT ELUTION PROFILE IS IN GREY.**

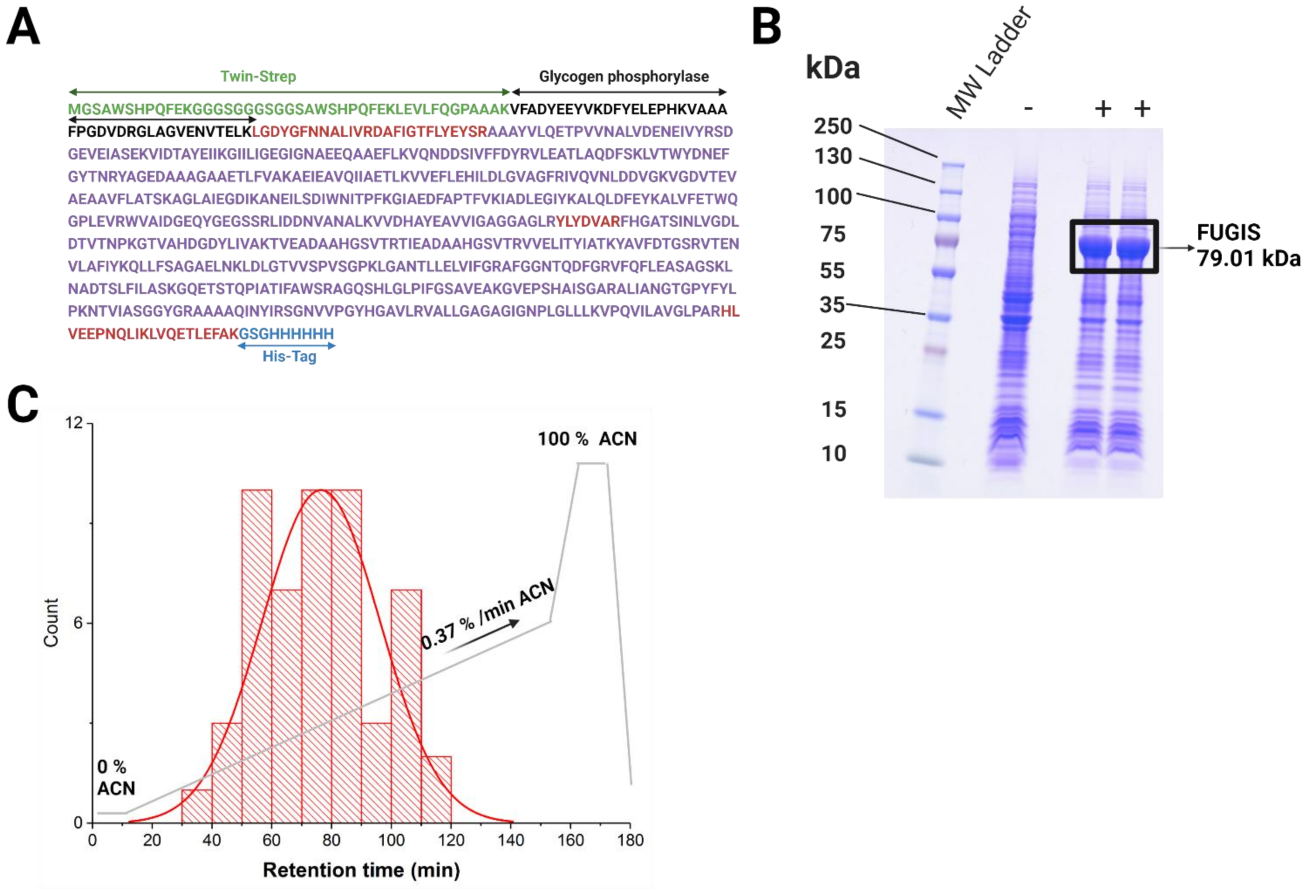

**Figure S 3. A) Full length sequence of FUGIS standard. B) Expressed FUGIS standard separated using 1D SDS PAGE. C) Retention time distribution of peptides in the FUGIS standard; gradient elution profile in grey.**

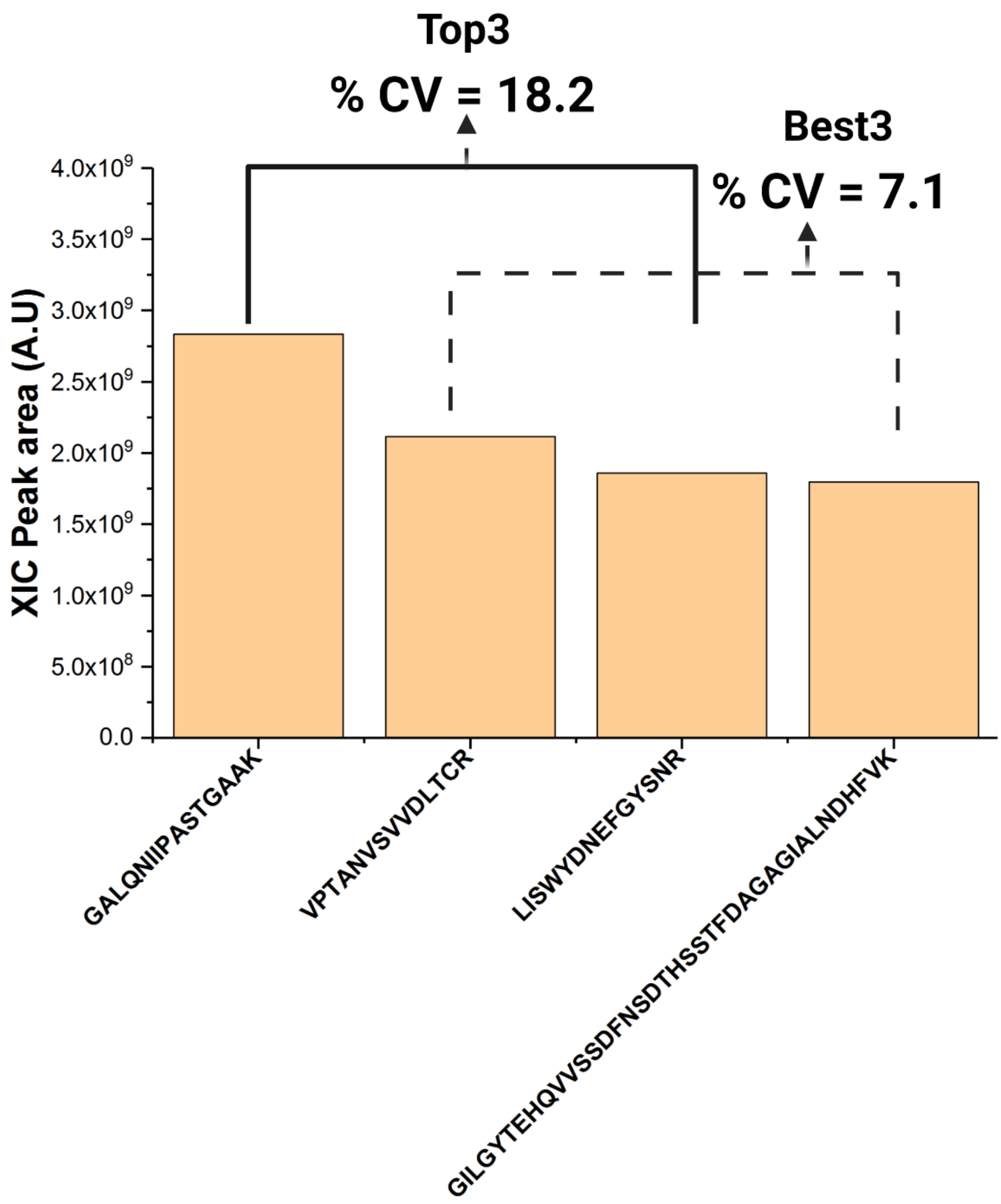

FIGURE S 4. XIC PEAK AREAS OF TOP4 PEPTIDES OF GAPDH (P04406) EXPLAINS THE RATIONAL BEHIND THE SELECTION OF BESTN PEPTIDES. %CV IS COEFFICIENT OF VARIATION OF MULTIPLE SELECTIONS.

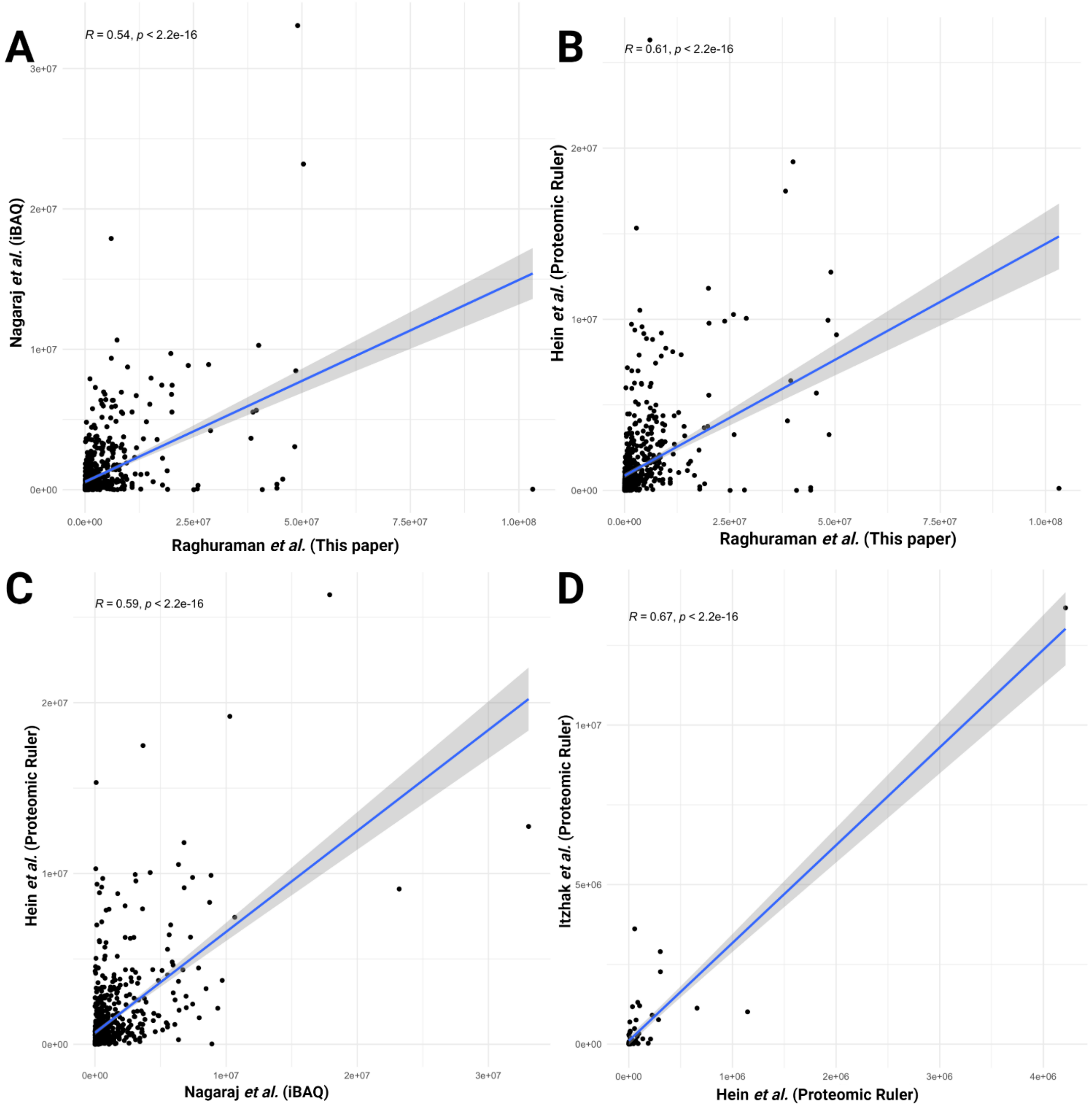

**FIGURE S 5. COMPARISON OF COPY NUMBER PER HELA CELL PROTEINS A) NAGARAJ *ET AL.* (IBAQ)[5] VS. RAGHURAMAN *ET AL* (THIS PAPER), (843 PROTEINS) B) HEIN *ET AL.* (PROTEOMIC RULER)[6] VS. RAGHURAMAN *ET AL*., (843 PROTEINS) , C) NAGARAJ *ET AL.* (IBAQ)[5] VS. HEIN *ET AL*. (PROTEOMIC RULER)[6], (843 PROTEINS) D) ITZHAK *ET AL.* (PROTEOMIC RULER)[7] VS. HEIN *ET AL* (PROTEOMIC RULER)[6] (82 PROTEINS). R IS SPEARMAN CORRELATION COEFFICIENT. ALL DATASETS WERE UNIFIED BY ENSEMBL GENE ID VIA GENE SYMBOL AND/OR MAJOR ID'S. ONLY PROTEINS UNEQUIVOCALLY MATCHED BETWEEN THE TWO COMPARED DATASETS WERE USED.**

TABLE S 1.MBAQ QUANTIFICATION OF UPS2 STANDARD

| Uniprot ID | Protein name | MBAQ (fmol on column) | UPS2 (fmol on column) | Quantification error (%) |
|---|---|---|---|---|
| P68871 | Haemoglobin beta chain | 1960.62 | 2173.91 | -9.81 |
| P02768 | Serum Albumin | 1883.46 | 2173.91 | -13.36 |
| P41159 | Leptin | 2159.04 | 2173.91 | -0.68 |
| P00915 | Carbonic anhydrase 1 | 1891.46 | 2173.91 | -12.99 |
| P00918 | Carbonic anhydrase 2 | 2255.1 | 2173.91 | 3.73 |
| P62988 | Ubiquitin | 1868.16 | 2173.91 | -14.06 |
| P62937 | Peptidicyl-prolyl cis-trans isomerase A | 238.38 | 217.39 | 9.65 |
| Q06830 | Peroxiredoxin 1 | 191.9 | 217.39 | -11.73 |
| P02144 | Myoglobulin C | 182.91 | 217.39 | -15.86 |
| P00167 | Cytochrome b5 | 201.19 | 217.39 | -7.45 |
| P04040 | Catalase | 138.13 | 217.39 | -36.46 |
| P15559 | NAD(p)H dehydrogenase (quinone) 1 C | 201.76 | 217.39 | -7.19 |
| P63165 | Small ubiquitin related modifier (SUMO-1) | 197.075 | 217.39 | -9.35 |
| P16083 | Ribosyldihydronicotinamide dehydrogenase | 15.65 | 21.74 | -28.01 |
| P06732 | Creatinine Kinase M-type | 12.64 | 21.74 | -41.86 |
| P61626 | Lysozyme C | 19.98 | 21.74 | -8.09 |
| P12081 | Histidyl tRNA synthetase | 19.24 | 21.74 | -11.50 |

TABLE S 2. PEPTIDE SEQUENCE OF GCP01

| |
|---|
| LTANAADITVQDDTADK |
| YVIENEDLVANVVPSEQLAYAAAR |
| ESAIDVGDSEK |
| YDFFIITDDNQVR |
| VETYAELGYEGLEK |
| VFDQNGLADWVDNNVR |
| ELFELTQELFGVGSNPSTDK |
| AVLDEYSEGSPYR |
| QANEYADELISGYK |
| VIDQVTYILEPGVGDGPIITVK |
| VLIEDDAPSPVPVQK |
| VYGIATEPIAIEEEGGAVK |
| AVFLSEAAAGAADEGAYK |

| |
|---|
| LFEAAQEEAQGGIEGILIIGK |
| SSGEGYNEDGIAVWR |
| EGQLIIPK |
| LFEDTEDAAYK |
| EAAPASGGSSAGDDKFDK |
| IQPFELAELVTEFVSK |
| GIIQEINEQEIGTSPLK |
| LAQAVQDDILK |
| AFDDFSLGQVIK |
| ELPTNVFHEDVSFAGDLLK |
| NSYGFENDYWSVLR |
| IIEYASDVVK |
| SFDQALSALEVK |
| VELPGNWTDFVLAR |
| DLGFETNVLAAAK |
| DQGAIDPATGHVSEFVAAGKG |
| ISEVANEISSSR |
| NTVTLDDPGVLQISTAGHFK |
| TVTGHAAEAEVTR |
| YVFALVQETVK |
| NLDAGATFLLQK |
| GFIVLDLLSNAGLR |
| SGATALFNFVEK |
| DLVLQSLVR |
| SWAFISAIPNTSTEQGR |
| LYFYPGPSGQAILAK |
| GYGGTALVTNR |
| IYNLNAAAAR |
| LVAHGYGPVVQGSR |
| AVALSVGLIANAGLNGK |
| APLGVLAIVNPVR |

TABLE S 3. PEPTIDE SEQUENCE OF GCP02

| |
|---|
| AAAYALQESPVVNAVLDENEIVYR |
| AAAYVLQETPVVNALVDENEIVYR |
| YVIENEDLVANVVPSEQLAYAAAR |
| ESDGVDIASEK |
| SDGEVEIASEK |
| SAIDVEEGDSK |
| VVDSAYEIIK |
| VIDTAYEIIK |
| IIEYASDVVK |
| GIILIGEIGGQAEEQAAEFLK |
| GIILIGEGIGNAEEQAAEFLK |

| |
|---|
| LFEAAQEEAQGGIEGILIIGK |
| VQNDDTIIFFDYR |
| VQNDDSIVFFDYR |
| YDFFIITDDNQVR |
| VELASLAQDFSK |
| VLEATLAQDFSK |
| SFDQALSALEVK |
| LVSWYDNEFGYSNR |
| LVTWYDNEFGYTNR |
| NSYGFENDYWSVLR |
| YAGEDAAGAAAESLFVAK |
| YAGEDAAAGAAETLFVAK |
| AVFLSEAAAGAADEGAYK |
| AEVEAVQIIAESLK |
| AEIEAVQIIAETLK |
| LSAIEIQVAEVEAK |
| VVEFLDHLIDLGVAGFR |
| VVEFLEHILDLGVAGFR |
| FGAVGLDILHDLFEVVR |
| IVQVQIDDVGK |
| IVQVNLDDVGK |
| GVDDIQVQVIK |
| VGDVTEVAEAVAFLASSK |
| VGDVTEVAEAAVFLATSK |
| SSALFAVAEAVETVDGVK |
| AGIAVEGDIK |
| AGLAIEGDIK |
| IDGEVAIGAK |
| ENAILTDIWNITPFK |
| ANEILSDIWNITPFK |
| FPTINEWIDTLIANK |
| GVAEDFAPSFVK |
| GIAEDFAPTFVK |
| VFSPAFDEAVGK |
| IDALEGVYK |
| IADLEGIYK |
| YVGELADIK |

*TABLE S 4. PEPTIDE SEQUENCE OF FUGIS*

| **Peptide sequence** | **Highest similarity found to a known sequence (%)** | **Sequence origin** |
|---|---|---|
| LGDYGFNNALIVR | 84.62 | *Bos taurus* |

| | | |
|---|---|---|
| DAFIGTFLYEYSR | 92.31 | *Bos taurus* |
| AAAYVLQETPVVNALVDENEIVYR | 83.33 | *Caenorhabditis elegans* |
| SDGEVEIASEK | 90 | *Drosophila melanogaster* |
| VIDTAYEIIK | 90 | *Caenorhabditis elegans* |
| GIILIGEGIGNAEEQAAEFLK | 86.36 | *Caenorhabditis elegans* |
| VQNDDSIVFFDYR | 83.41 | *Caenorhabditis elegans* |
| VLEATLAQDFSK | 90.91 | *Caenorhabditis elegans* |
| LVTWYDNEFGYTNR | 92.86 | *Caenorhabditis elegans* |
| YAGEDAAAGAAETLFVAK | 83.33 | *Caenorhabditis elegans* |
| AEIEAVQIIAETLK | 85.71 | *Drosophila melanogaster* |
| VVEFLEHILDLGVAGFR | 82.35 | *Drosophila melanogaster* |
| IVQVNLDDVGK | 90.91 | *Drosophila melanogaster* |
| VGDVTEVAEAAVFLATSK | 82.35 | *Drosophila melanogaster* |
| AGLAIEGDIK | 90 | *Drosophila melanogaster* |
| ANEILSDIWNITPFK | 85.71 | *Drosophila melanogaster* |
| GIAEDFAPTFVK | 83.33 | *Drosophila melanogaster* |
| IADLEGIYK | 88.89 | *Drosophila melanogaster* |
| ALQLDFEYK | 88.91 | *Drosophila melanogaster* |
| ALVFETWQGPLEVR | 85.71 | *Caenorhabditis elegans* |
| WVAIDGEQYGEGSSR | 80 | *Caenorhabditis elegans* |
| LIDDNVANALK | 83.33 | *Caenorhabditis elegans* |
| VVDHAYEAVVIGAGGAGLR | 89.47 | *Caenorhabditis elegans* |
| YLYDVAR | 95.23 | *Bos taurus* |
| FHGATSINLVGDLDTVTNPK | 85.71 | *Caenorhabditis elegans* |
| GTVAHDGDYLIVAK | 85.71 | *Caenorhabditis elegans* |
| TVEADAAHGSVTR | 92.31 | *Caenorhabditis elegans* |
| TIEADAAHGSVTR | 84.62 | *Caenorhabditis elegans* |
| VVELITYIATK | 81.82 | *Caenorhabditis elegans* |
| YAVFDTGSR | 88.89 | *Caenorhabditis elegans* |
| VTENVLAFIYK | 81.82 | *Caenorhabditis elegans* |
| QLLFSAGAELNK | 83.33 | *Caenorhabditis elegans* |
| LDLGTVVSPVSGPK | 92.86 | *Caenorhabditis elegans* |
| LGANTLLELVIFGR | 85.71 | *Caenorhabditis elegans* |
| AFGGNTQDFGR | 90 | *Caenorhabditis elegans* |
| VFQFLEASAGSK | 83.33 | *Caenorhabditis elegans* |
| LNADTSLFILASK | 84.62 | *Caenorhabditis elegans* |
| GQETSTQPIATIFAWSR | 88.24 | *Caenorhabditis elegans* |
| AGQSHLGLPIFGSAVEAK | 77.78 | *Caenorhabditis elegans* |
| GVEPSHAISGAR | 83.33 | *Caenorhabditis elegans* |
| ALIANGTGPYFYLPK | 93.33 | *Caenorhabditis elegans* |
| NTVIASGGYGR | 90.91 | *Caenorhabditis elegans* |
| AAAAQINYIR | 90.41 | *Caenorhabditis elegans* |
| SGNVVPGYHGAVLR | 86.67 | *Caenorhabditis elegans* |
| VALLGAGAGIGNPLGLLLK | 84.21 | *Caenorhabditis elegans* |
| VPQVILAVGLPAR | 69.23 | *Caenorhabditis elegans* |
| HLVEEPNQLIK | 90.91 | *Bos taurus* |

| | | |
|---|---|---|
| LVQETLEFAK | 90 | *Bos taurus* |

TABLE S 5. MBAQ VS. MS WESTERN VS. HI-3 OF 1 MILLION CELLS IN 500 µL

(SEM= STANDARD ERROR OF MEANS)

| Protein name | MBAQ (fmol) | SEM | MS Western (fmol) | SEM | Hi-3 quantification (fmol) | SEM | % Error (MBAQ vs MS Western) | % Error (Hi-3 vs MS Western) |
|---|---|---|---|---|---|---|---|---|
| PLK-1 | 1.6 | 0.04 | 1.95 | 0.03 | 1.62 | 0.06 | 17.95 | 16.92 |
| CAT | 32.75 | 1.45 | 42.97 | 1.04 | 27.81 | 1.08 | 23.78 | 35.28 |
| TBA1A | 2356.83 | 72.62 | 2357.92 | 60.11 | 2153.74 | 70.32 | 0.05 | 8.66 |
| GAPDH | 4242.35 | 110.21 | 4488.92 | 89.04 | 3552.61 | 74.32 | 5.49 | 20.86 |

TABLE S 6. MBAQ VS. MS WESTERN VS. HI-3 OF 1 MILLION CELLS IN 1000 µL

(SEM= STANDARD ERROR OF MEANS)

| Protein name | MBAQ (fmol) | SEM | MS Western (fmol) | SEM | Hi-3 quantification (fmol) | SEM | % Error (MBAQ vs MS Western) | % Error (Hi-3 vs MS Western) |
|---|---|---|---|---|---|---|---|---|
| PLK-1 | 0.751 | 0.002 | 0.852 | 0.005 | 0.715 | 0.009 | 11.85 | 16.08 |
| CAT | 17.65 | 1.82 | 19.61 | 1.21 | 15.75 | 1.32 | 9.99 | 19.68 |
| TBA1A | 1243.14 | 85.42 | 1226.09 | 70.21 | 1037.04 | 60.51 | 1.39 | 15.42 |
| GAPDH | 2266.94 | 63.25 | 2364.12 | 59.12 | 2066.41 | 48.32 | 4.11 | 12.59 |

TABLE S 7. BEST N PEPTIDES OF 4 HELA PROTEINS

| Protein name | Best N peptides used | %CV |
|---|---|---|
| PLK-1* | HINPVAASLIQK, FSIAPSSLDPSNR | 12.2 |
| CAT | FNTANDDNVTQVR, AFYVNVLNEEQR | 6.2 |
| TBA1A | AVFVDLEPTVIDEVR, TIGGGDDSFNTFFSETGAGK,NLDIERPTYTNLNR | 4.8 |
| GAPDH | VPTANVSVVDLTCR, LISWYDNEFGYSNR, GILGYTEHQVVSSDFNSDTHSSTFDAGAGIALNDHFVK | 7.1 |

*TABLE S 8. GLOBEQUANT QUANTIFICATION OF 1450 HELA PROTEINS*

| Proteins | 1 million HeLa cells in 1000 µl (fmol) | 1 million HeLa cells in 500 µl (fmol) | Fold Change | Log2FC |
|---|---|---|---|---|
| sp\|P60709\|ACTB_HUMAN Actin, cytoplasmic 1 OS=Homo sapiens OX=9606 GN=ACTB PE=1 SV=1 | 4088.96 | 7353.35 | 1.80 | 0.85 |
| sp\|P63261\|ACTG_HUMAN Actin, cytoplasmic 2 OS=Homo sapiens OX=9606 GN=ACTG1 PE=1 SV=1 | 3740.36 | 7013.21 | 1.88 | 0.91 |
| sp\|Q6S8J3\|POTEE_HUMAN POTE ankyrin domain family member E OS=Homo sapiens OX=9606 GN=POTEE PE=2 SV=3 | 2330.73 | 2402.03 | 1.03 | 0.04 |
| sp\|P0CG39\|POTEJ_HUMAN POTE ankyrin domain family member J OS=Homo sapiens OX=9606 GN=POTEJ PE=3 SV=1 | 2330.73 | 1622.79 | 0.70 | -0.52 |
| sp\|P04406\|G3P_HUMAN Glyceraldehyde-3-phosphate dehydrogenase OS=Homo sapiens OX=9606 GN=GAPDH PE=1 SV=3 | 2226.29 | 4307.67 | 1.93 | 0.95 |
| sp\|P68032\|ACTC_HUMAN Actin, alpha cardiac muscle 1 OS=Homo sapiens OX=9606 GN=ACTC1 PE=1 SV=1 | 1395.87 | 2340.83 | 1.68 | 0.75 |
| sp\|P68104\|EF1A1_HUMAN Elongation factor 1-alpha 1 OS=Homo sapiens OX=9606 GN=EEF1A1 PE=1 SV=1 | 1315.90 | 2412.97 | 1.83 | 0.87 |
| sp\|P68363\|TBA1B_HUMAN Tubulin alpha-1B chain OS=Homo sapiens OX=9606 GN=TUBA1B PE=1 SV=1 | 1253.39 | 2447.40 | 1.95 | 0.97 |
| sp\|Q71U36\|TBA1A_HUMAN Tubulin alpha-1A chain OS=Homo sapiens OX=9606 GN=TUBA1A PE=1 SV=1 | 1243.14 | 2356.83 | 1.90 | 0.92 |
| sp\|Q9BQE3\|TBA1C_HUMAN Tubulin alpha-1C chain OS=Homo sapiens OX=9606 GN=TUBA1C PE=1 SV=1 | 1175.81 | 2370.49 | 2.02 | 1.01 |
| sp\|P68366-2\|TBA4A_HUMAN Isoform 2 of Tubulin alpha-4A chain OS=Homo sapiens OX=9606 GN=TUBA4A | 1134.91 | 2257.92 | 1.99 | 0.99 |
| sp\|P08238\|HS90B_HUMAN Heat shock protein HSP 90-beta OS=Homo sapiens OX=9606 GN=HSP90AB1 PE=1 SV=4 | 1088.67 | 2202.28 | 2.02 | 1.02 |
| tr\|A0A2U3TZH3\|A0A2U3TZH3_HUMAN Elongation factor 1-alpha OS=Homo sapiens OX=9606 GN=EEF1A2 PE=1 SV=1 | 940.53 | 2979.98 | 3.17 | 1.66 |
| sp\|P07437\|TBB5_HUMAN Tubulin beta chain OS=Homo sapiens OX=9606 GN=TUBB PE=1 SV=2 | 926.28 | 1832.44 | 1.98 | 0.98 |

| | | | | |
|---|---|---|---|---|
| sp\|P06733\|ENOA_HUMAN Alpha-enolase OS=Homo sapiens OX=9606 GN=ENO1 PE=1 SV=2 | 904.37 | 1521.40 | 1.68 | 0.75 |
| tr\|K7EPT8\|K7EPT8_HUMAN Glial fibrillary acidic protein (Fragment) OS=Homo sapiens OX=9606 GN=GFAP PE=1 SV=2 | 718.96 | 482.84 | 0.67 | -0.57 |
| tr\|A0A0A0MRX1\|A0A0A0MRX1_HUMAN ELAV-like protein OS=Homo sapiens OX=9606 GN=ELAVL2 PE=1 SV=1 | 689.90 | 762.32 | 1.10 | 0.14 |
| sp\|P07900\|HS90A_HUMAN Heat shock protein HSP 90-alpha OS=Homo sapiens OX=9606 GN=HSP90AA1 PE=1 SV=5 | 645.76 | 1315.99 | 2.04 | 1.03 |
| sp\|P62805\|H4_HUMAN Histone H4 OS=Homo sapiens OX=9606 GN=HIST1H4A PE=1 SV=2 | 611.41 | 303.90 | 0.50 | -1.01 |
| sp\|P68371\|TBB4B_HUMAN Tubulin beta-4B chain OS=Homo sapiens OX=9606 GN=TUBB4B PE=1 SV=1 | 594.16 | 1403.17 | 2.36 | 1.24 |
| sp\|P11021\|BIP_HUMAN Endoplasmic reticulum chaperone BiP OS=Homo sapiens OX=9606 GN=HSPA5 PE=1 SV=2 | 581.80 | 1395.68 | 2.40 | 1.26 |
| sp\|P07195\|LDHB_HUMAN L-lactate dehydrogenase B chain OS=Homo sapiens OX=9606 GN=LDHB PE=1 SV=2 | 579.65 | 1117.72 | 1.93 | 0.95 |
| sp\|P04350\|TBB4A_HUMAN Tubulin beta-4A chain OS=Homo sapiens OX=9606 GN=TUBB4A PE=1 SV=2 | 575.87 | 1277.06 | 2.22 | 1.15 |
| sp\|Q9BVA1\|TBB2B_HUMAN Tubulin beta-2B chain OS=Homo sapiens OX=9606 GN=TUBB2B PE=1 SV=1 | 575.87 | 1277.06 | 2.22 | 1.15 |
| sp\|Q13509\|TBB3_HUMAN Tubulin beta-3 chain OS=Homo sapiens OX=9606 GN=TUBB3 PE=1 SV=2 | 575.87 | 1277.06 | 2.22 | 1.15 |
| sp\|Q9BUF5\|TBB6_HUMAN Tubulin beta-6 chain OS=Homo sapiens OX=9606 GN=TUBB6 PE=1 SV=1 | 575.87 | 1277.06 | 2.22 | 1.15 |
| sp\|P0C0S5\|H2AZ_HUMAN Histone H2A.Z OS=Homo sapiens OX=9606 GN=H2AFZ PE=1 SV=2 | 568.68 | 904.06 | 1.59 | 0.67 |
| sp\|Q7L7L0\|H2A3_HUMAN Histone H2A type 3 OS=Homo sapiens OX=9606 GN=HIST3H2A PE=1 SV=3 | 568.68 | 572.50 | 1.01 | 0.01 |
| sp\|Q6FI13\|H2A2A_HUMAN Histone H2A type 2-A OS=Homo sapiens OX=9606 GN=HIST2H2AA3 PE=1 SV=3 | 568.68 | 572.50 | 1.01 | 0.01 |
| sp\|P11142\|HSP7C_HUMAN Heat shock cognate 71 kDa protein OS=Homo sapiens OX=9606 GN=HSPA8 PE=1 SV=1 | 551.11 | 1768.44 | 3.21 | 1.68 |
| sp\|P00338\|LDHA_HUMAN L-lactate dehydrogenase A chain OS=Homo sapiens OX=9606 GN=LDHA PE=1 SV=2 | 549.56 | 980.09 | 1.78 | 0.83 |
| tr\|H3BTA2\|H3BTA2_HUMAN Serine/threonine-protein phosphatase (Fragment) OS=Homo sapiens OX=9606 GN=PPP4C PE=1 SV=1 | 538.08 | 36.09 | 0.07 | -3.90 |

| | | | | |
|---|---|---|---|---|
| sp\|P0DMV8\|HS71A_HUMAN Heat shock 70 kDa protein 1A OS=Homo sapiens OX=9606 GN=HSPA1A PE=1 SV=1 | 526.62 | 1182.21 | 2.24 | 1.17 |
| sp\|Q06830\|PRDX1_HUMAN Peroxiredoxin-1 OS=Homo sapiens OX=9606 GN=PRDX1 PE=1 SV=1 | 525.66 | 1453.85 | 2.77 | 1.47 |
| sp\|P60174-1\|TPIS_HUMAN Isoform 2 of Triosephosphate isomerase OS=Homo sapiens OX=9606 GN=TPI1 | 518.29 | 412.50 | 0.80 | -0.33 |
| sp\|P08670\|VIME_HUMAN Vimentin OS=Homo sapiens OX=9606 GN=VIM PE=1 SV=4 | 506.97 | 687.10 | 1.36 | 0.44 |
| sp\|P62937\|PPIA_HUMAN Peptidyl-prolyl cis-trans isomerase A OS=Homo sapiens OX=9606 GN=PPIA PE=1 SV=2 | 495.49 | 1836.04 | 3.71 | 1.89 |
| sp\|P54652\|HSP72_HUMAN Heat shock-related 70 kDa protein 2 OS=Homo sapiens OX=9606 GN=HSPA2 PE=1 SV=1 | 469.04 | 1179.88 | 2.52 | 1.33 |
| tr\|A0A2R8YFE2\|A0A2R8YFE2_HUMAN Actin, cytoplasmic 1 (Fragment) OS=Homo sapiens OX=9606 GN=ACTB PE=1 SV=1 | 469.00 | 2336.02 | 4.98 | 2.32 |
| sp\|P14618\|KPYM_HUMAN Pyruvate kinase PKM OS=Homo sapiens OX=9606 GN=PKM PE=1 SV=4 | 457.44 | 859.72 | 1.88 | 0.91 |
| sp\|P10809\|CH60_HUMAN 60 kDa heat shock protein, mitochondrial OS=Homo sapiens OX=9606 GN=HSPD1 PE=1 SV=2 | 417.31 | 747.53 | 1.79 | 0.84 |
| sp\|P23528\|COF1_HUMAN Cofilin-1 OS=Homo sapiens OX=9606 GN=CFL1 PE=1 SV=3 | 412.68 | 1415.25 | 3.43 | 1.78 |
| sp\|P04075\|ALDOA_HUMAN Fructose-bisphosphate aldolase A OS=Homo sapiens OX=9606 GN=ALDOA PE=1 SV=2 | 411.37 | 1026.12 | 2.49 | 1.32 |
| sp\|Q8WXI7\|MUC16_HUMAN Mucin-16 OS=Homo sapiens OX=9606 GN=MUC16 PE=1 SV=3 | 406.82 | 724.08 | 1.78 | 0.83 |
| tr\|A0A0J9YYC8\|A0A0J9YYC8_HUMAN Trypsin-2 OS=Homo sapiens OX=9606 GN=PRSS2 PE=1 SV=1 | 402.25 | 7.90 | 0.02 | -5.67 |
| sp\|Q58FF6\|H90B4_HUMAN Putative heat shock protein HSP 90-beta 4 OS=Homo sapiens OX=9606 GN=HSP90AB4P PE=5 SV=1 | 388.15 | 594.44 | 1.53 | 0.61 |
| sp\|P63104\|1433Z_HUMAN 14-3-3 protein zeta/delta OS=Homo sapiens OX=9606 GN=YWHAZ PE=1 SV=1 | 382.56 | 1140.22 | 2.98 | 1.58 |
| tr\|Q53FA3\|Q53FA3_HUMAN HSPA1L (Fragment) OS=Homo sapiens OX=9606 GN=HSPA1L PE=1 SV=1 | 380.88 | 1456.68 | 3.82 | 1.94 |
| sp\|P00558\|PGK1_HUMAN Phosphoglycerate kinase 1 OS=Homo sapiens OX=9606 GN=PGK1 PE=1 SV=3 | 375.13 | 751.94 | 2.00 | 1.00 |
| tr\|H0YA27\|H0YA27_HUMAN Cyclin-I (Fragment) OS=Homo sapiens OX=9606 GN=CCNI PE=1 SV=1 | 365.10 | 653.46 | 1.79 | 0.84 |

| | | | | |
|---|---|---|---|---|
| tr\|M0R0M7\|M0R0M7_HUMAN Syntaxin-binding protein 2 OS=Homo sapiens OX=9606 GN=STXBP2 PE=1 SV=1 | 365.10 | 19.65 | 0.05 | -4.22 |
| sp\|P07355\|ANXA2_HUMAN Annexin A2 OS=Homo sapiens OX=9606 GN=ANXA2 PE=1 SV=2 | 364.39 | 822.60 | 2.26 | 1.17 |
| sp\|P13639\|EF2_HUMAN Elongation factor 2 OS=Homo sapiens OX=9606 GN=EEF2 PE=1 SV=4 | 364.14 | 100.17 | 0.28 | -1.86 |
| sp\|P27797\|CALR_HUMAN Calreticulin OS=Homo sapiens OX=9606 GN=CALR PE=1 SV=1 | 362.86 | 1104.58 | 3.04 | 1.61 |
| tr\|A0A0C4DG17\|A0A0C4DG17_HUMAN 40S ribosomal protein SA OS=Homo sapiens OX=9606 GN=RPSA PE=1 SV=1 | 356.14 | 848.46 | 2.38 | 1.25 |
| sp\|P61981\|1433G_HUMAN 14-3-3 protein gamma OS=Homo sapiens OX=9606 GN=YWHAG PE=1 SV=2 | 351.94 | 226.27 | 0.64 | -0.64 |
| sp\|P60842\|IF4A1_HUMAN Eukaryotic initiation factor 4A-I OS=Homo sapiens OX=9606 GN=EIF4A1 PE=1 SV=1 | 345.54 | 592.08 | 1.71 | 0.78 |
| sp\|P05388\|RLA0_HUMAN 60S acidic ribosomal protein P0 OS=Homo sapiens OX=9606 GN=RPLP0 PE=1 SV=1 | 327.34 | 699.40 | 2.14 | 1.10 |
| sp\|P29401-2\|TKT_HUMAN Isoform 2 of Transketolase OS=Homo sapiens OX=9606 GN=TKT | 319.78 | 661.59 | 2.07 | 1.05 |
| sp\|P61978-2\|HNRPK_HUMAN Isoform 2 of Heterogeneous nuclear ribonucleoprotein K OS=Homo sapiens OX=9606 GN=HNRNPK | 319.39 | 382.18 | 1.20 | 0.26 |
| tr\|A0A0U1RRM8\|A0A0U1RRM8_HUMAN Fermitin family homolog 2 (Fragment) OS=Homo sapiens OX=9606 GN=FERMT2 PE=1 SV=1 | 313.63 | 6.24 | 0.02 | -5.65 |
| sp\|P68431\|H31_HUMAN Histone H3.1 OS=Homo sapiens OX=9606 GN=HIST1H3A PE=1 SV=2 | 308.81 | 99.58 | 0.32 | -1.63 |
| tr\|K7EK07\|K7EK07_HUMAN Histone H3 (Fragment) OS=Homo sapiens OX=9606 GN=H3F3B PE=1 SV=1 | 308.81 | 99.58 | 0.32 | -1.63 |
| sp\|Q71DI3\|H32_HUMAN Histone H3.2 OS=Homo sapiens OX=9606 GN=HIST2H3A PE=1 SV=3 | 308.81 | 99.58 | 0.32 | -1.63 |
| sp\|P05386\|RLA1_HUMAN 60S acidic ribosomal protein P1 OS=Homo sapiens OX=9606 GN=RPLP1 PE=1 SV=1 | 305.50 | 1155.85 | 3.78 | 1.92 |
| tr\|G8JLB6\|G8JLB6_HUMAN Heterogeneous nuclear ribonucleoprotein H OS=Homo sapiens OX=9606 GN=HNRNPH1 PE=1 SV=1 | 298.44 | 450.57 | 1.51 | 0.59 |
| tr\|B5MDF5\|B5MDF5_HUMAN GTP-binding nuclear protein Ran OS=Homo sapiens OX=9606 GN=RAN PE=1 SV=1 | 296.10 | 578.25 | 1.95 | 0.97 |

| | | | | |
|---|---|---|---|---|
| sp\|P27824-2\|CALX_HUMAN Isoform 2 of Calnexin OS=Homo sapiens OX=9606 GN=CANX | 275.16 | 14.68 | 0.05 | -4.23 |
| tr\|A0A0A0MTS2\|A0A0A0MTS2_HUMAN Glucose-6-phosphate isomerase (Fragment) OS=Homo sapiens OX=9606 GN=GPI PE=1 SV=1 | 267.44 | 495.07 | 1.85 | 0.89 |
| sp\|P07237\|PDIA1_HUMAN Protein disulfide-isomerase OS=Homo sapiens OX=9606 GN=P4HB PE=1 SV=3 | 265.91 | 263.99 | 0.99 | -0.01 |
| tr\|D6RB85\|D6RB85_HUMAN Calnexin (Fragment) OS=Homo sapiens OX=9606 GN=CANX PE=1 SV=1 | 265.71 | 56.42 | 0.21 | -2.24 |
| sp\|P16403\|H12_HUMAN Histone H1.2 OS=Homo sapiens OX=9606 GN=HIST1H1C PE=1 SV=2 | 244.74 | 408.21 | 1.67 | 0.74 |
| sp\|P22626\|ROA2_HUMAN Heterogeneous nuclear ribonucleoproteins A2/B1 OS=Homo sapiens OX=9606 GN=HNRNPA2B1 PE=1 SV=2 | 244.49 | 835.24 | 3.42 | 1.77 |
| tr\|A0A2R8Y811\|A0A2R8Y811_HUMAN 40S ribosomal protein S14 (Fragment) OS=Homo sapiens OX=9606 GN=RPS14 PE=1 SV=1 | 241.53 | 78.53 | 0.33 | -1.62 |
| sp\|O14556\|G3PT_HUMAN Glyceraldehyde-3-phosphate dehydrogenase, testis-specific OS=Homo sapiens OX=9606 GN=GAPDHS PE=1 SV=2 | 240.92 | 589.38 | 2.45 | 1.29 |
| sp\|P38646\|GRP75_HUMAN Stress-70 protein, mitochondrial OS=Homo sapiens OX=9606 GN=HSPA9 PE=1 SV=2 | 229.03 | 412.85 | 1.80 | 0.85 |
| sp\|Q96AY3\|FKB10_HUMAN Peptidyl-prolyl cis-trans isomerase FKBP10 OS=Homo sapiens OX=9606 GN=FKBP10 PE=1 SV=1 | 227.76 | 26.17 | 0.11 | -3.12 |
| tr\|E7EQG2\|E7EQG2_HUMAN Eukaryotic initiation factor 4A-II OS=Homo sapiens OX=9606 GN=EIF4A2 PE=1 SV=1 | 226.98 | 549.45 | 2.42 | 1.28 |
| sp\|P37802-2\|TAGL2_HUMAN Isoform 2 of Transgelin-2 OS=Homo sapiens OX=9606 GN=TAGLN2 | 222.46 | 318.62 | 1.43 | 0.52 |
| sp\|P30101\|PDIA3_HUMAN Protein disulfide-isomerase A3 OS=Homo sapiens OX=9606 GN=PDIA3 PE=1 SV=4 | 219.71 | 258.77 | 1.18 | 0.24 |
| sp\|P14625\|ENPL_HUMAN Endoplasmin OS=Homo sapiens OX=9606 GN=HSP90B1 PE=1 SV=1 | 218.68 | 215.11 | 0.98 | -0.02 |
| sp\|P62241\|RS8_HUMAN 40S ribosomal protein S8 OS=Homo sapiens OX=9606 GN=RPS8 PE=1 SV=2 | 214.13 | 579.57 | 2.71 | 1.44 |
| sp\|P62269\|RS18_HUMAN 40S ribosomal protein S18 OS=Homo sapiens OX=9606 GN=RPS18 PE=1 SV=3 | 208.55 | 523.90 | 2.51 | 1.33 |
| tr\|Q32Q12\|Q32Q12_HUMAN Nucleoside diphosphate kinase OS=Homo sapiens OX=9606 GN=NME1-NME2 PE=1 SV=1 | 204.47 | 724.04 | 3.54 | 1.82 |

| | | | | |
|---|---|---|---|---|
| sp\|P15531-2\|NDKA_HUMAN Isoform 2 of Nucleoside diphosphate kinase A OS=Homo sapiens OX=9606 GN=NME1 | 204.47 | 724.04 | 3.54 | 1.82 |
| tr\|H0YHX9\|H0YHX9_HUMAN Nascent polypeptide-associated complex subunit alpha (Fragment) OS=Homo sapiens OX=9606 GN=NACA PE=1 SV=1 | 200.98 | 453.20 | 2.25 | 1.17 |
| sp\|P15880\|RS2_HUMAN 40S ribosomal protein S2 OS=Homo sapiens OX=9606 GN=RPS2 PE=1 SV=2 | 187.89 | 342.19 | 1.82 | 0.86 |
| sp\|P62249\|RS16_HUMAN 40S ribosomal protein S16 OS=Homo sapiens OX=9606 GN=RPS16 PE=1 SV=2 | 186.77 | 479.64 | 2.57 | 1.36 |
| sp\|P06748-2\|NPM_HUMAN Isoform 2 of Nucleophosmin OS=Homo sapiens OX=9606 GN=NPM1 | 185.67 | 276.56 | 1.49 | 0.57 |
| sp\|P04792\|HSPB1_HUMAN Heat shock protein beta-1 OS=Homo sapiens OX=9606 GN=HSPB1 PE=1 SV=2 | 185.66 | 570.82 | 3.07 | 1.62 |
| sp\|P62081\|RS7_HUMAN 40S ribosomal protein S7 OS=Homo sapiens OX=9606 GN=RPS7 PE=1 SV=1 | 185.12 | 174.80 | 0.94 | -0.08 |
| sp\|P23284\|PPIB_HUMAN Peptidyl-prolyl cis-trans isomerase B OS=Homo sapiens OX=9606 GN=PPIB PE=1 SV=2 | 182.40 | 439.20 | 2.41 | 1.27 |
| sp\|P18669\|PGAM1_HUMAN Phosphoglycerate mutase 1 OS=Homo sapiens OX=9606 GN=PGAM1 PE=1 SV=2 | 181.28 | 170.43 | 0.94 | -0.09 |
| sp\|P63244\|RACK1_HUMAN Receptor of activated protein C kinase 1 OS=Homo sapiens OX=9606 GN=RACK1 PE=1 SV=3 | 180.24 | 444.13 | 2.46 | 1.30 |
| sp\|P26641-2\|EF1G_HUMAN Isoform 2 of Elongation factor 1-gamma OS=Homo sapiens OX=9606 GN=EEF1G | 175.09 | 335.79 | 1.92 | 0.94 |
| sp\|P50990\|TCPQ_HUMAN T-complex protein 1 subunit theta OS=Homo sapiens OX=9606 GN=CCT8 PE=1 SV=4 | 173.24 | 334.00 | 1.93 | 0.95 |
| tr\|E5RI99\|E5RI99_HUMAN 60S ribosomal protein L30 (Fragment) OS=Homo sapiens OX=9606 GN=RPL30 PE=1 SV=1 | 172.03 | 370.97 | 2.16 | 1.11 |
| sp\|P63241\|IF5A1_HUMAN Eukaryotic translation initiation factor 5A-1 OS=Homo sapiens OX=9606 GN=EIF5A PE=1 SV=2 | 171.93 | 387.53 | 2.25 | 1.17 |
| sp\|Q58FG1\|HS904_HUMAN Putative heat shock protein HSP 90-alpha A4 OS=Homo sapiens OX=9606 GN=HSP90AA4P PE=5 SV=1 | 167.90 | 385.20 | 2.29 | 1.20 |
| tr\|A0A2R8Y4W8\|A0A2R8Y4W8_HUMAN Girdin OS=Homo sapiens OX=9606 GN=CCDC88A PE=1 SV=1 | 166.46 | 98.08 | 0.59 | -0.76 |
| sp\|P62851\|RS25_HUMAN 40S ribosomal protein S25 OS=Homo sapiens OX=9606 GN=RPS25 PE=1 SV=1 | 163.34 | 357.86 | 2.19 | 1.13 |

| | | | | |
|---|---|---|---|---|
| sp\|P48643\|TCPE_HUMAN T-complex protein 1 subunit epsilon OS=Homo sapiens OX=9606 GN=CCT5 PE=1 SV=1 | 162.72 | 122.27 | 0.75 | -0.41 |
| sp\|P30050\|RL12_HUMAN 60S ribosomal protein L12 OS=Homo sapiens OX=9606 GN=RPL12 PE=1 SV=1 | 162.42 | 312.89 | 1.93 | 0.95 |
| sp\|P04083\|ANXA1_HUMAN Annexin A1 OS=Homo sapiens OX=9606 GN=ANXA1 PE=1 SV=2 | 162.08 | 330.58 | 2.04 | 1.03 |
| tr\|G3V203\|G3V203_HUMAN 60S ribosomal protein L18 OS=Homo sapiens OX=9606 GN=RPL18 PE=1 SV=1 | 158.78 | 510.99 | 3.22 | 1.69 |
| sp\|Q9BRZ2\|TRI56_HUMAN E3 ubiquitin-protein ligase TRIM56 OS=Homo sapiens OX=9606 GN=TRIM56 PE=1 SV=3 | 154.97 | 11.88 | 0.08 | -3.71 |
| tr\|F5H157\|F5H157_HUMAN Ras-related protein Rab-35 (Fragment) OS=Homo sapiens OX=9606 GN=RAB35 PE=1 SV=1 | 154.46 | 313.69 | 2.03 | 1.02 |
| sp\|P61026\|RAB10_HUMAN Ras-related protein Rab-10 OS=Homo sapiens OX=9606 GN=RAB10 PE=1 SV=1 | 154.46 | 313.69 | 2.03 | 1.02 |
| sp\|P61006\|RAB8A_HUMAN Ras-related protein Rab-8A OS=Homo sapiens OX=9606 GN=RAB8A PE=1 SV=1 | 154.46 | 313.69 | 2.03 | 1.02 |
| sp\|P14174\|MIF_HUMAN Macrophage migration inhibitory factor OS=Homo sapiens OX=9606 GN=MIF PE=1 SV=4 | 154.30 | 575.93 | 3.73 | 1.90 |
| tr\|A0A1B0GTU8\|A0A1B0GTU8_HUMAN Renin receptor OS=Homo sapiens OX=9606 GN=ATP6AP2 PE=1 SV=1 | 152.73 | 13.44 | 0.09 | -3.51 |
| sp\|P54727\|RD23B_HUMAN UV excision repair protein RAD23 homolog B OS=Homo sapiens OX=9606 GN=RAD23B PE=1 SV=1 | 152.13 | 28.62 | 0.19 | -2.41 |
| tr\|H7BZJ3\|H7BZJ3_HUMAN Protein disulfide-isomerase A3 (Fragment) OS=Homo sapiens OX=9606 GN=PDIA3 PE=1 SV=1 | 149.29 | 253.79 | 1.70 | 0.77 |
| sp\|P62854\|RS26_HUMAN 40S ribosomal protein S26 OS=Homo sapiens OX=9606 GN=RPS26 PE=1 SV=3 | 149.15 | 53.14 | 0.36 | -1.49 |
| sp\|P61247\|RS3A_HUMAN 40S ribosomal protein S3a OS=Homo sapiens OX=9606 GN=RPS3A PE=1 SV=2 | 149.15 | 205.93 | 1.38 | 0.47 |
| sp\|P06576\|ATPB_HUMAN ATP synthase subunit beta, mitochondrial OS=Homo sapiens OX=9606 GN=ATP5F1B PE=1 SV=3 | 148.57 | 271.61 | 1.83 | 0.87 |
| sp\|P50454\|SERPH_HUMAN Serpin H1 OS=Homo sapiens OX=9606 GN=SERPINH1 PE=1 SV=2 | 148.28 | 277.94 | 1.87 | 0.91 |
| sp\|P81605\|DCD_HUMAN Dermcidin OS=Homo sapiens OX=9606 GN=DCD PE=1 SV=2 | 147.65 | 67.80 | 0.46 | -1.12 |
| tr\|M0R0F0\|M0R0F0_HUMAN 40S ribosomal protein S5 (Fragment) OS=Homo sapiens OX=9606 GN=RPS5 PE=1 SV=1 | 147.58 | 575.63 | 3.90 | 1.96 |

| | | | | |
|---|---|---|---|---|
| tr\|A0A0U1RRM4\|A0A0U1RRM4_HUMAN Polypyrimidine tract-binding protein 1 OS=Homo sapiens OX=9606 GN=PTBP1 PE=1 SV=1 | 146.10 | 42.68 | 0.29 | -1.78 |
| sp\|P46783\|RS10_HUMAN 40S ribosomal protein S10 OS=Homo sapiens OX=9606 GN=RPS10 PE=1 SV=1 | 145.43 | 408.47 | 2.81 | 1.49 |
| sp\|P30041\|PRDX6_HUMAN Peroxiredoxin-6 OS=Homo sapiens OX=9606 GN=PRDX6 PE=1 SV=3 | 144.63 | 283.14 | 1.96 | 0.97 |
| tr\|V9GZ17\|V9GZ17_HUMAN Tubulin alpha-8 chain (Fragment) OS=Homo sapiens OX=9606 GN=TUBA8 PE=1 SV=1 | 144.62 | 328.83 | 2.27 | 1.19 |
| sp\|P62277\|RS13_HUMAN 40S ribosomal protein S13 OS=Homo sapiens OX=9606 GN=RPS13 PE=1 SV=2 | 142.19 | 340.74 | 2.40 | 1.26 |
| sp\|P36578\|RL4_HUMAN 60S ribosomal protein L4 OS=Homo sapiens OX=9606 GN=RPL4 PE=1 SV=5 | 141.39 | 252.19 | 1.78 | 0.83 |
| sp\|Q01650\|LAT1_HUMAN Large neutral amino acids transporter small subunit 1 OS=Homo sapiens OX=9606 GN=SLC7A5 PE=1 SV=2 | 139.63 | 81.21 | 0.58 | -0.78 |
| sp\|Q02878\|RL6_HUMAN 60S ribosomal protein L6 OS=Homo sapiens OX=9606 GN=RPL6 PE=1 SV=3 | 138.94 | 388.74 | 2.80 | 1.48 |
| sp\|P49411\|EFTU_HUMAN Elongation factor Tu, mitochondrial OS=Homo sapiens OX=9606 GN=TUFM PE=1 SV=2 | 138.46 | 254.76 | 1.84 | 0.88 |
| sp\|Q15084-2\|PDIA6_HUMAN Isoform 2 of Protein disulfide-isomerase A6 OS=Homo sapiens OX=9606 GN=PDIA6 | 138.19 | 147.72 | 1.07 | 0.10 |
| sp\|P62244\|RS15A_HUMAN 40S ribosomal protein S15a OS=Homo sapiens OX=9606 GN=RPS15A PE=1 SV=2 | 136.16 | 371.64 | 2.73 | 1.45 |
| sp\|Q15365\|PCBP1_HUMAN Poly(rC)-binding protein 1 OS=Homo sapiens OX=9606 GN=PCBP1 PE=1 SV=2 | 134.37 | 206.86 | 1.54 | 0.62 |
| sp\|P25705\|ATPA_HUMAN ATP synthase subunit alpha, mitochondrial OS=Homo sapiens OX=9606 GN=ATP5F1A PE=1 SV=1 | 134.09 | 301.54 | 2.25 | 1.17 |
| sp\|P00966\|ASSY_HUMAN Argininosuccinate synthase OS=Homo sapiens OX=9606 GN=ASS1 PE=1 SV=2 | 132.36 | 259.68 | 1.96 | 0.97 |
| sp\|P62701\|RS4X_HUMAN 40S ribosomal protein S4, X isoform OS=Homo sapiens OX=9606 GN=RPS4X PE=1 SV=2 | 129.69 | 256.55 | 1.98 | 0.98 |
| sp\|O00299\|CLIC1_HUMAN Chloride intracellular channel protein 1 OS=Homo sapiens OX=9606 GN=CLIC1 PE=1 SV=4 | 129.48 | 115.21 | 0.89 | -0.17 |
| tr\|J3KPF3\|J3KPF3_HUMAN 4F2 cell-surface antigen heavy chain OS=Homo sapiens OX=9606 GN=SLC3A2 PE=1 SV=1 | 129.04 | 264.47 | 2.05 | 1.04 |

| | | | | |
|---|---|---|---|---|
| sp\|P18124\|RL7_HUMAN 60S ribosomal protein L7 OS=Homo sapiens OX=9606 GN=RPL7 PE=1 SV=1 | 127.52 | 206.78 | 1.62 | 0.70 |
| sp\|P23396\|RS3_HUMAN 40S ribosomal protein S3 OS=Homo sapiens OX=9606 GN=RPS3 PE=1 SV=2 | 123.54 | 431.74 | 3.49 | 1.81 |
| sp\|P47914\|RL29_HUMAN 60S ribosomal protein L29 OS=Homo sapiens OX=9606 GN=RPL29 PE=1 SV=2 | 123.54 | 181.95 | 1.47 | 0.56 |
| sp\|Q5VT79\|AXA81_HUMAN Annexin A8-like protein 1 OS=Homo sapiens OX=9606 GN=ANXA8L1 PE=2 SV=2 | 123.37 | 369.55 | 3.00 | 1.58 |
| sp\|P38919\|IF4A3_HUMAN Eukaryotic initiation factor 4A-III OS=Homo sapiens OX=9606 GN=EIF4A3 PE=1 SV=4 | 123.19 | 228.49 | 1.85 | 0.89 |
| sp\|P62424\|RL7A_HUMAN 60S ribosomal protein L7a OS=Homo sapiens OX=9606 GN=RPL7A PE=1 SV=2 | 123.11 | 335.43 | 2.72 | 1.45 |
| sp\|Q9UQ80\|PA2G4_HUMAN Proliferation-associated protein 2G4 OS=Homo sapiens OX=9606 GN=PA2G4 PE=1 SV=3 | 121.35 | 216.66 | 1.79 | 0.84 |
| sp\|P05141\|ADT2_HUMAN ADP/ATP translocase 2 OS=Homo sapiens OX=9606 GN=SLC25A5 PE=1 SV=7 | 121.15 | 205.42 | 1.70 | 0.76 |
| tr\|A0A024R4M0\|A0A024R4M0_HUMAN 40S ribosomal protein S9 OS=Homo sapiens OX=9606 GN=RPS9 PE=1 SV=1 | 120.94 | 375.10 | 3.10 | 1.63 |
| sp\|P78371\|TCPB_HUMAN T-complex protein 1 subunit beta OS=Homo sapiens OX=9606 GN=CCT2 PE=1 SV=4 | 120.75 | 546.76 | 4.53 | 2.18 |
| tr\|J3KTE4\|J3KTE4_HUMAN Ribosomal protein L19 OS=Homo sapiens OX=9606 GN=RPL19 PE=1 SV=1 | 120.50 | 59.85 | 0.50 | -1.01 |
| tr\|E7EQR4\|E7EQR4_HUMAN Ezrin OS=Homo sapiens OX=9606 GN=EZR PE=1 SV=3 | 120.21 | 179.43 | 1.49 | 0.58 |
| tr\|A0A0A0MR02\|A0A0A0MR02_HUMAN Voltage-dependent anion-selective channel protein 2 (Fragment) OS=Homo sapiens OX=9606 GN=VDAC2 PE=1 SV=1 | 118.53 | 239.80 | 2.02 | 1.02 |
| sp\|P35232\|PHB_HUMAN Prohibitin OS=Homo sapiens OX=9606 GN=PHB PE=1 SV=1 | 116.63 | 260.65 | 2.23 | 1.16 |
| sp\|P26038\|MOES_HUMAN Moesin OS=Homo sapiens OX=9606 GN=MSN PE=1 SV=3 | 116.61 | 262.88 | 2.25 | 1.17 |
| tr\|F8W6I7\|F8W6I7_HUMAN Heterogeneous nuclear ribonucleoprotein A1 OS=Homo sapiens OX=9606 GN=HNRNPA1 PE=1 SV=2 | 115.87 | 252.96 | 2.18 | 1.13 |
| tr\|H0YIV4\|H0YIV4_HUMAN Nucleosome assembly protein 1-like 1 (Fragment) OS=Homo sapiens OX=9606 GN=NAP1L1 PE=1 SV=1 | 115.41 | 145.08 | 1.26 | 0.33 |
| sp\|P52272-2\|HNRPM_HUMAN Isoform 2 of Heterogeneous nuclear ribonucleoprotein M OS=Homo sapiens OX=9606 GN=HNRNPM | 115.18 | 221.93 | 1.93 | 0.95 |

| | | | | |
|---|---|---|---|---|
| sp\|Q16658\|FSCN1_HUMAN Fascin OS=Homo sapiens OX=9606 GN=FSCN1 PE=1 SV=3 | 114.88 | 152.33 | 1.33 | 0.41 |
| tr\|A0A2R8Y5S7\|A0A2R8Y5S7_HUMAN Radixin OS=Homo sapiens OX=9606 GN=RDX PE=1 SV=1 | 114.62 | 262.88 | 2.29 | 1.20 |
| sp\|P02545\|LMNA_HUMAN Prelamin-A/C OS=Homo sapiens OX=9606 GN=LMNA PE=1 SV=1 | 114.47 | 113.18 | 0.99 | -0.02 |
| sp\|P62906\|RL10A_HUMAN 60S ribosomal protein L10a OS=Homo sapiens OX=9606 GN=RPL10A PE=1 SV=2 | 113.80 | 121.60 | 1.07 | 0.10 |
| sp\|Q8WZ42-11\|TITIN_HUMAN Isoform 11 of Titin OS=Homo sapiens OX=9606 GN=TTN | 113.18 | 5.19 | 0.05 | -4.45 |
| tr\|B2R5W2\|B2R5W2_HUMAN Heterogeneous nuclear ribonucleoproteins C1/C2 OS=Homo sapiens OX=9606 GN=HNRNPC PE=1 SV=1 | 113.14 | 106.94 | 0.95 | -0.08 |
| sp\|P12277\|KCRB_HUMAN Creatine kinase B-type OS=Homo sapiens OX=9606 GN=CKB PE=1 SV=1 | 112.82 | 219.66 | 1.95 | 0.96 |
| sp\|P50502\|F10A1_HUMAN Hsc70-interacting protein OS=Homo sapiens OX=9606 GN=ST13 PE=1 SV=2 | 111.98 | 148.62 | 1.33 | 0.41 |
| sp\|P61353\|RL27_HUMAN 60S ribosomal protein L27 OS=Homo sapiens OX=9606 GN=RPL27 PE=1 SV=2 | 110.28 | 241.83 | 2.19 | 1.13 |
| sp\|P21796\|VDAC1_HUMAN Voltage-dependent anion-selective channel protein 1 OS=Homo sapiens OX=9606 GN=VDAC1 PE=1 SV=2 | 110.19 | 276.11 | 2.51 | 1.33 |
| sp\|P40227\|TCPZ_HUMAN T-complex protein 1 subunit zeta OS=Homo sapiens OX=9606 GN=CCT6A PE=1 SV=3 | 109.07 | 218.48 | 2.00 | 1.00 |
| tr\|A0A286YF22\|A0A286YF22_HUMAN D-3-phosphoglycerate dehydrogenase OS=Homo sapiens OX=9606 GN=PHGDH PE=1 SV=1 | 107.97 | 198.44 | 1.84 | 0.88 |
| sp\|P12236\|ADT3_HUMAN ADP/ATP translocase 3 OS=Homo sapiens OX=9606 GN=SLC25A6 PE=1 SV=4 | 107.65 | 130.86 | 1.22 | 0.28 |
| tr\|A0A087WXM6\|A0A087WXM6_HUMAN 60S ribosomal protein L17 (Fragment) OS=Homo sapiens OX=9606 GN=RPL17 PE=3 SV=1 | 106.51 | 155.76 | 1.46 | 0.55 |
| tr\|E7EPB3\|E7EPB3_HUMAN 60S ribosomal protein L14 OS=Homo sapiens OX=9606 GN=RPL14 PE=1 SV=1 | 105.49 | 113.12 | 1.07 | 0.10 |
| sp\|P61604\|CH10_HUMAN 10 kDa heat shock protein, mitochondrial OS=Homo sapiens OX=9606 GN=HSPE1 PE=1 SV=2 | 105.06 | 159.60 | 1.52 | 0.60 |
| sp\|P31946-2\|1433B_HUMAN Isoform Short of 14-3-3 protein beta/alpha OS=Homo sapiens OX=9606 GN=YWHAB | 104.27 | 246.52 | 2.36 | 1.24 |
| tr\|A0A0B4J1R4\|A0A0B4J1R4_HUMAN 4-hydroxyphenylpyruvate dioxygenase OS=Homo sapiens OX=9606 GN=HPD PE=1 SV=1 | 101.52 | 135.67 | 1.34 | 0.42 |

| | | | | |
|---|---|---|---|---|
| sp\|P09382\|LEG1_HUMAN Galectin-1 OS=Homo sapiens OX=9606 GN=LGALS1 PE=1 SV=2 | 101.49 | 64.93 | 0.64 | -0.64 |
| sp\|P30048-2\|PRDX3_HUMAN Isoform 2 of Thioredoxin-dependent peroxide reductase, mitochondrial OS=Homo sapiens OX=9606 GN=PRDX3 | 100.77 | 258.29 | 2.56 | 1.36 |
| sp\|Q99832\|TCPH_HUMAN T-complex protein 1 subunit eta OS=Homo sapiens OX=9606 GN=CCT7 PE=1 SV=2 | 100.40 | 151.21 | 1.51 | 0.59 |
| sp\|P62258\|1433E_HUMAN 14-3-3 protein epsilon OS=Homo sapiens OX=9606 GN=YWHAE PE=1 SV=1 | 99.86 | 249.66 | 2.50 | 1.32 |
| sp\|P23526\|SAHH_HUMAN Adenosylhomocysteinase OS=Homo sapiens OX=9606 GN=AHCY PE=1 SV=4 | 99.51 | 214.37 | 2.15 | 1.11 |
| sp\|P39019\|RS19_HUMAN 40S ribosomal protein S19 OS=Homo sapiens OX=9606 GN=RPS19 PE=1 SV=2 | 99.20 | 238.51 | 2.40 | 1.27 |
| sp\|P67809\|YBOX1_HUMAN Nuclease-sensitive element-binding protein 1 OS=Homo sapiens OX=9606 GN=YBX1 PE=1 SV=3 | 97.19 | 233.91 | 2.41 | 1.27 |
| tr\|C9J4Z3\|C9J4Z3_HUMAN 60S ribosomal protein L37a OS=Homo sapiens OX=9606 GN=RPL37A PE=1 SV=1 | 97.02 | 266.24 | 2.74 | 1.46 |
| sp\|P84077\|ARF1_HUMAN ADP-ribosylation factor 1 OS=Homo sapiens OX=9606 GN=ARF1 PE=1 SV=2 | 96.96 | 250.48 | 2.58 | 1.37 |
| sp\|P52597\|HNRPF_HUMAN Heterogeneous nuclear ribonucleoprotein F OS=Homo sapiens OX=9606 GN=HNRNPF PE=1 SV=3 | 94.16 | 178.24 | 1.89 | 0.92 |
| sp\|P63173\|RL38_HUMAN 60S ribosomal protein L38 OS=Homo sapiens OX=9606 GN=RPL38 PE=1 SV=2 | 92.97 | 226.96 | 2.44 | 1.29 |
| tr\|J3KPX7\|J3KPX7_HUMAN Prohibitin-2 OS=Homo sapiens OX=9606 GN=PHB2 PE=1 SV=2 | 92.80 | 269.60 | 2.91 | 1.54 |
| sp\|Q13162\|PRDX4_HUMAN Peroxiredoxin-4 OS=Homo sapiens OX=9606 GN=PRDX4 PE=1 SV=1 | 92.41 | 246.06 | 2.66 | 1.41 |
| tr\|A0A087WUS0\|A0A087WUS0_HUMAN 40S ribosomal protein S24 OS=Homo sapiens OX=9606 GN=RPS24 PE=1 SV=1 | 91.04 | 256.86 | 2.82 | 1.50 |
| sp\|O43760-2\|SNG2_HUMAN Isoform 2 of Synaptogyrin-2 OS=Homo sapiens OX=9606 GN=SYNGR2 | 90.71 | 89.11 | 0.98 | -0.03 |
| sp\|P32119\|PRDX2_HUMAN Peroxiredoxin-2 OS=Homo sapiens OX=9606 GN=PRDX2 PE=1 SV=5 | 90.70 | 212.99 | 2.35 | 1.23 |
| sp\|Q14019\|COTL1_HUMAN Coactosin-like protein OS=Homo sapiens OX=9606 GN=COTL1 PE=1 SV=3 | 90.15 | 76.74 | 0.85 | -0.23 |

| | | | | |
|---|---|---|---|---|
| sp\|P00441\|SODC_HUMAN Superoxide dismutase [Cu-Zn] OS=Homo sapiens OX=9606 GN=SOD1 PE=1 SV=2 | 90.08 | 43.68 | 0.48 | -1.04 |
| sp\|P55795\|HNRH2_HUMAN Heterogeneous nuclear ribonucleoprotein H2 OS=Homo sapiens OX=9606 GN=HNRNPH2 PE=1 SV=1 | 89.37 | 140.15 | 1.57 | 0.65 |
| tr\|D6R9P3\|D6R9P3_HUMAN Heterogeneous nuclear ribonucleoprotein A/B OS=Homo sapiens OX=9606 GN=HNRNPAB PE=1 SV=1 | 89.15 | 204.21 | 2.29 | 1.20 |
| sp\|P84103-2\|SRSF3_HUMAN Isoform 2 of Serine/arginine-rich splicing factor 3 OS=Homo sapiens OX=9606 GN=SRSF3 | 89.08 | 236.64 | 2.66 | 1.41 |
| tr\|M0R117\|M0R117_HUMAN 60S ribosomal protein L18a OS=Homo sapiens OX=9606 GN=RPL18A PE=1 SV=1 | 88.95 | 187.23 | 2.10 | 1.07 |
| sp\|P62857\|RS28_HUMAN 40S ribosomal protein S28 OS=Homo sapiens OX=9606 GN=RPS28 PE=1 SV=1 | 88.83 | 199.70 | 2.25 | 1.17 |
| sp\|P35613-2\|BASI_HUMAN Isoform 2 of Basigin OS=Homo sapiens OX=9606 GN=BSG | 88.56 | 113.44 | 1.28 | 0.36 |
| sp\|P06703\|S10A6_HUMAN Protein S100-A6 OS=Homo sapiens OX=9606 GN=S100A6 PE=1 SV=1 | 87.98 | 165.76 | 1.88 | 0.91 |
| sp\|P32969\|RL9_HUMAN 60S ribosomal protein L9 OS=Homo sapiens OX=9606 GN=RPL9 PE=1 SV=1 | 87.89 | 187.85 | 2.14 | 1.10 |
| tr\|A0A1B0GV23\|A0A1B0GV23_HUMAN Cathepsin D OS=Homo sapiens OX=9606 GN=CTSD PE=1 SV=1 | 86.35 | 16.46 | 0.19 | -2.39 |
| sp\|P07954-2\|FUMH_HUMAN Isoform Cytoplasmic of Fumarate hydratase, mitochondrial OS=Homo sapiens OX=9606 GN=FH | 85.61 | 178.32 | 2.08 | 1.06 |
| tr\|F8W1A4\|F8W1A4_HUMAN Adenylate kinase 2, mitochondrial OS=Homo sapiens OX=9606 GN=AK2 PE=1 SV=1 | 85.14 | 164.73 | 1.93 | 0.95 |
| sp\|P13667\|PDIA4_HUMAN Protein disulfide-isomerase A4 OS=Homo sapiens OX=9606 GN=PDIA4 PE=1 SV=2 | 84.18 | 248.95 | 2.96 | 1.56 |
| sp\|P50991\|TCPD_HUMAN T-complex protein 1 subunit delta OS=Homo sapiens OX=9606 GN=CCT4 PE=1 SV=4 | 83.52 | 147.57 | 1.77 | 0.82 |
| sp\|P62280\|RS11_HUMAN 40S ribosomal protein S11 OS=Homo sapiens OX=9606 GN=RPS11 PE=1 SV=3 | 83.34 | 177.08 | 2.12 | 1.09 |
| sp\|P00505\|AATM_HUMAN Aspartate aminotransferase, mitochondrial OS=Homo sapiens OX=9606 GN=GOT2 PE=1 SV=3 | 81.99 | 146.42 | 1.79 | 0.84 |
| sp\|P31689\|DNJA1_HUMAN DnaJ homolog subfamily A member 1 OS=Homo sapiens OX=9606 GN=DNAJA1 PE=1 SV=2 | 81.43 | 150.89 | 1.85 | 0.89 |
| sp\|P55072\|TERA_HUMAN Transitional endoplasmic reticulum ATPase OS=Homo sapiens OX=9606 GN=VCP PE=1 SV=4 | 80.27 | 13.13 | 0.16 | -2.61 |

| | | | | |
|---|---|---|---|---|
| sp\|P19338\|NUCL_HUMAN Nucleolin OS=Homo sapiens OX=9606 GN=NCL PE=1 SV=3 | 80.05 | 196.72 | 2.46 | 1.30 |
| tr\|A0A0B4J1Z1\|A0A0B4J1Z1_HUMAN Serine/arginine-rich-splicing factor 7 OS=Homo sapiens OX=9606 GN=SRSF7 PE=1 SV=1 | 80.03 | 158.50 | 1.98 | 0.99 |
| sp\|P26639\|SYTC_HUMAN Threonine--tRNA ligase, cytoplasmic OS=Homo sapiens OX=9606 GN=TARS PE=1 SV=3 | 79.88 | 202.52 | 2.54 | 1.34 |
| sp\|P62273\|RS29_HUMAN 40S ribosomal protein S29 OS=Homo sapiens OX=9606 GN=RPS29 PE=1 SV=2 | 79.00 | 60.82 | 0.77 | -0.38 |
| sp\|P40926\|MDHM_HUMAN Malate dehydrogenase, mitochondrial OS=Homo sapiens OX=9606 GN=MDH2 PE=1 SV=3 | 77.97 | 192.45 | 2.47 | 1.30 |
| sp\|P40925-3\|MDHC_HUMAN Isoform 3 of Malate dehydrogenase, cytoplasmic OS=Homo sapiens OX=9606 GN=MDH1 | 77.23 | 193.27 | 2.50 | 1.32 |
| sp\|P08758\|ANXA5_HUMAN Annexin A5 OS=Homo sapiens OX=9606 GN=ANXA5 PE=1 SV=2 | 77.20 | 170.90 | 2.21 | 1.15 |
| sp\|P84085\|ARF5_HUMAN ADP-ribosylation factor 5 OS=Homo sapiens OX=9606 GN=ARF5 PE=1 SV=2 | 77.08 | 515.69 | 6.69 | 2.74 |
| sp\|P18085\|ARF4_HUMAN ADP-ribosylation factor 4 OS=Homo sapiens OX=9606 GN=ARF4 PE=1 SV=3 | 77.08 | 515.69 | 6.69 | 2.74 |
| sp\|P15121\|ALDR_HUMAN Aldose reductase OS=Homo sapiens OX=9606 GN=AKR1B1 PE=1 SV=3 | 76.07 | 176.46 | 2.32 | 1.21 |
| sp\|P17987\|TCPA_HUMAN T-complex protein 1 subunit alpha OS=Homo sapiens OX=9606 GN=TCP1 PE=1 SV=1 | 75.99 | 128.33 | 1.69 | 0.76 |
| sp\|P26373\|RL13_HUMAN 60S ribosomal protein L13 OS=Homo sapiens OX=9606 GN=RPL13 PE=1 SV=4 | 75.94 | 431.20 | 5.68 | 2.51 |
| sp\|P38159\|RBMX_HUMAN RNA-binding motif protein, X chromosome OS=Homo sapiens OX=9606 GN=RBMX PE=1 SV=3 | 75.91 | 73.89 | 0.97 | -0.04 |
| sp\|P51149\|RAB7A_HUMAN Ras-related protein Rab-7a OS=Homo sapiens OX=9606 GN=RAB7A PE=1 SV=1 | 74.67 | 99.95 | 1.34 | 0.42 |
| sp\|P25398\|RS12_HUMAN 40S ribosomal protein S12 OS=Homo sapiens OX=9606 GN=RPS12 PE=1 SV=3 | 74.21 | 175.39 | 2.36 | 1.24 |
| sp\|P27348\|1433T_HUMAN 14-3-3 protein theta OS=Homo sapiens OX=9606 GN=YWHAQ PE=1 SV=1 | 74.11 | 182.37 | 2.46 | 1.30 |
| sp\|O75396\|SC22B_HUMAN Vesicle-trafficking protein SEC22b OS=Homo sapiens OX=9606 GN=SEC22B PE=1 SV=4 | 74.04 | 24.74 | 0.33 | -1.58 |
| sp\|P28066\|PSA5_HUMAN Proteasome subunit alpha type-5 OS=Homo sapiens OX=9606 GN=PSMA5 PE=1 SV=3 | 73.63 | 75.11 | 1.02 | 0.03 |

| | | | | |
|---|---|---|---|---|
| sp\|P16401\|H15_HUMAN Histone H1.5 OS=Homo sapiens OX=9606 GN=HIST1H1B PE=1 SV=3 | 73.31 | 66.74 | 0.91 | -0.14 |
| sp\|P49368\|TCPG_HUMAN T-complex protein 1 subunit gamma OS=Homo sapiens OX=9606 GN=CCT3 PE=1 SV=4 | 73.08 | 184.43 | 2.52 | 1.34 |
| sp\|P30086\|PEBP1_HUMAN Phosphatidylethanolamine-binding protein 1 OS=Homo sapiens OX=9606 GN=PEBP1 PE=1 SV=3 | 72.98 | 202.02 | 2.77 | 1.47 |
| sp\|Q13151\|ROA0_HUMAN Heterogeneous nuclear ribonucleoprotein A0 OS=Homo sapiens OX=9606 GN=HNRNPA0 PE=1 SV=1 | 72.90 | 151.75 | 2.08 | 1.06 |
| sp\|P51991\|ROA3_HUMAN Heterogeneous nuclear ribonucleoprotein A3 OS=Homo sapiens OX=9606 GN=HNRNPA3 PE=1 SV=2 | 72.49 | 170.79 | 2.36 | 1.24 |
| sp\|Q04917\|1433F_HUMAN 14-3-3 protein eta OS=Homo sapiens OX=9606 GN=YWHAH PE=1 SV=4 | 71.82 | 211.89 | 2.95 | 1.56 |
| sp\|Q01105\|SET_HUMAN Protein SET OS=Homo sapiens OX=9606 GN=SET PE=1 SV=3 | 71.75 | 102.79 | 1.43 | 0.52 |
| sp\|Q96AG4\|LRC59_HUMAN Leucine-rich repeat-containing protein 59 OS=Homo sapiens OX=9606 GN=LRRC59 PE=1 SV=1 | 69.53 | 188.98 | 2.72 | 1.44 |
| sp\|P11940\|PABP1_HUMAN Polyadenylate-binding protein 1 OS=Homo sapiens OX=9606 GN=PABPC1 PE=1 SV=2 | 69.34 | 132.64 | 1.91 | 0.94 |
| tr\|E9PLL6\|E9PLL6_HUMAN 60S ribosomal protein L27a OS=Homo sapiens OX=9606 GN=RPL27A PE=1 SV=1 | 69.00 | 145.43 | 2.11 | 1.08 |
| sp\|P49643\|PRI2_HUMAN DNA primase large subunit OS=Homo sapiens OX=9606 GN=PRIM2 PE=1 SV=2 | 68.80 | 16.08 | 0.23 | -2.10 |
| sp\|Q6EEV6\|SUMO4_HUMAN Small ubiquitin-related modifier 4 OS=Homo sapiens OX=9606 GN=SUMO4 PE=1 SV=2 | 68.75 | 96.14 | 1.40 | 0.48 |
| tr\|C9JNW5\|C9JNW5_HUMAN 60S ribosomal protein L24 OS=Homo sapiens OX=9606 GN=RPL24 PE=1 SV=1 | 68.61 | 242.59 | 3.54 | 1.82 |
| sp\|P62917\|RL8_HUMAN 60S ribosomal protein L8 OS=Homo sapiens OX=9606 GN=RPL8 PE=1 SV=2 | 68.57 | 178.01 | 2.60 | 1.38 |
| sp\|Q9H0U4\|RAB1B_HUMAN Ras-related protein Rab-1B OS=Homo sapiens OX=9606 GN=RAB1B PE=1 SV=1 | 68.30 | 182.91 | 2.68 | 1.42 |
| sp\|P62820\|RAB1A_HUMAN Ras-related protein Rab-1A OS=Homo sapiens OX=9606 GN=RAB1A PE=1 SV=3 | 68.30 | 182.91 | 2.68 | 1.42 |
| sp\|Q14974\|IMB1_HUMAN Importin subunit beta-1 OS=Homo sapiens OX=9606 GN=KPNB1 PE=1 SV=2 | 67.41 | 32.99 | 0.49 | -1.03 |

| | | | | |
|---|---|---|---|---|
| sp\|Q02790\|FKBP4_HUMAN Peptidyl-prolyl cis-trans isomerase FKBP4 OS=Homo sapiens OX=9606 GN=FKBP4 PE=1 SV=3 | 67.32 | 121.01 | 1.80 | 0.85 |
| tr\|A0A1C7CYX9\|A0A1C7CYX9_HUMAN Dihydropyrimidinase-related protein 2 OS=Homo sapiens OX=9606 GN=DPYSL2 PE=1 SV=1 | 64.96 | 30.79 | 0.47 | -1.08 |
| sp\|P39748\|FEN1_HUMAN Flap endonuclease 1 OS=Homo sapiens OX=9606 GN=FEN1 PE=1 SV=1 | 64.89 | 104.61 | 1.61 | 0.69 |
| sp\|P35637-2\|FUS_HUMAN Isoform Short of RNA-binding protein FUS OS=Homo sapiens OX=9606 GN=FUS | 64.17 | 98.28 | 1.53 | 0.62 |
| sp\|O14818\|PSA7_HUMAN Proteasome subunit alpha type-7 OS=Homo sapiens OX=9606 GN=PSMA7 PE=1 SV=1 | 63.74 | 178.98 | 2.81 | 1.49 |
| sp\|P60866-2\|RS20_HUMAN Isoform 2 of 40S ribosomal protein S20 OS=Homo sapiens OX=9606 GN=RPS20 | 63.56 | 185.66 | 2.92 | 1.55 |
| sp\|P62879\|GBB2_HUMAN Guanine nucleotide-binding protein G(I)/G(S)/G(T) subunit beta-2 OS=Homo sapiens OX=9606 GN=GNB2 PE=1 SV=3 | 63.43 | 66.91 | 1.05 | 0.08 |
| sp\|P62873-2\|GBB1_HUMAN Isoform 2 of Guanine nucleotide-binding protein G(I)/G(S)/G(T) subunit beta-1 OS=Homo sapiens OX=9606 GN=GNB1 | 63.43 | 23.19 | 0.37 | -1.45 |
| sp\|Q9HAV0\|GBB4_HUMAN Guanine nucleotide-binding protein subunit beta-4 OS=Homo sapiens OX=9606 GN=GNB4 PE=1 SV=3 | 63.43 | 14.62 | 0.23 | -2.12 |
| sp\|P12956\|XRCC6_HUMAN X-ray repair cross-complementing protein 6 OS=Homo sapiens OX=9606 GN=XRCC6 PE=1 SV=2 | 63.28 | 148.84 | 2.35 | 1.23 |
| tr\|A0A0C4DG40\|A0A0C4DG40_HUMAN Nesprin-1 OS=Homo sapiens OX=9606 GN=SYNE1 PE=1 SV=1 | 62.74 | 195.44 | 3.11 | 1.64 |
| sp\|P29692\|EF1D_HUMAN Elongation factor 1-delta OS=Homo sapiens OX=9606 GN=EEF1D PE=1 SV=5 | 62.66 | 216.01 | 3.45 | 1.79 |
| sp\|P10599\|THIO_HUMAN Thioredoxin OS=Homo sapiens OX=9606 GN=TXN PE=1 SV=3 | 62.27 | 150.47 | 2.42 | 1.27 |
| tr\|H0YEY5\|H0YEY5_HUMAN Plakophilin-3 (Fragment) OS=Homo sapiens OX=9606 GN=PKP3 PE=1 SV=1 | 61.89 | 193.91 | 3.13 | 1.65 |
| sp\|Q12931-2\|TRAP1_HUMAN Isoform 2 of Heat shock protein 75 kDa, mitochondrial OS=Homo sapiens OX=9606 GN=TRAP1 | 61.39 | 150.52 | 2.45 | 1.29 |
| sp\|P18077\|RL35A_HUMAN 60S ribosomal protein L35a OS=Homo sapiens OX=9606 GN=RPL35A PE=1 SV=2 | 61.28 | 94.96 | 1.55 | 0.63 |

| | | | | |
|---|---|---|---|---|
| sp\|P24534\|EF1B_HUMAN Elongation factor 1-beta OS=Homo sapiens OX=9606 GN=EEF1B2 PE=1 SV=3 | 60.56 | 147.00 | 2.43 | 1.28 |
| tr\|A0A087WSW9\|A0A087WSW9_HUMAN Thioredoxin reductase 1, cytoplasmic OS=Homo sapiens OX=9606 GN=TXNRD1 PE=1 SV=1 | 60.26 | 121.91 | 2.02 | 1.02 |
| tr\|M0QYS1\|M0QYS1_HUMAN 60S ribosomal protein L13a (Fragment) OS=Homo sapiens OX=9606 GN=RPL13A PE=1 SV=2 | 60.24 | 156.34 | 2.60 | 1.38 |
| sp\|P23381\|SYWC_HUMAN Tryptophan--tRNA ligase, cytoplasmic OS=Homo sapiens OX=9606 GN=WARS PE=1 SV=2 | 60.24 | 107.39 | 1.78 | 0.83 |
| sp\|P50395\|GDIB_HUMAN Rab GDP dissociation inhibitor beta OS=Homo sapiens OX=9606 GN=GDI2 PE=1 SV=2 | 60.16 | 103.29 | 1.72 | 0.78 |
| sp\|Q99497\|PARK7_HUMAN Protein/nucleic acid deglycase DJ-1 OS=Homo sapiens OX=9606 GN=PARK7 PE=1 SV=2 | 60.15 | 123.86 | 2.06 | 1.04 |
| tr\|D3YTB1\|D3YTB1_HUMAN 60S ribosomal protein L32 (Fragment) OS=Homo sapiens OX=9606 GN=RPL32 PE=1 SV=1 | 59.54 | 59.52 | 1.00 | 0.00 |
| tr\|E9PBS1\|E9PBS1_HUMAN Multifunctional protein ADE2 (Fragment) OS=Homo sapiens OX=9606 GN=PAICS PE=1 SV=1 | 58.71 | 143.50 | 2.44 | 1.29 |
| sp\|P09972\|ALDOC_HUMAN Fructose-bisphosphate aldolase C OS=Homo sapiens OX=9606 GN=ALDOC PE=1 SV=2 | 57.49 | 239.47 | 4.17 | 2.06 |
| tr\|B4DJV2\|B4DJV2_HUMAN Citrate synthase OS=Homo sapiens OX=9606 GN=CS PE=1 SV=1 | 57.39 | 105.29 | 1.83 | 0.88 |
| tr\|H0YIC4\|H0YIC4_HUMAN Citrate synthase (Fragment) OS=Homo sapiens OX=9606 GN=CS PE=1 SV=1 | 57.39 | 90.92 | 1.58 | 0.66 |
| tr\|Q5VV89\|Q5VV89_HUMAN Microsomal glutathione S-transferase 3 OS=Homo sapiens OX=9606 GN=MGST3 PE=1 SV=1 | 57.21 | 103.66 | 1.81 | 0.86 |
| sp\|P05387\|RLA2_HUMAN 60S acidic ribosomal protein P2 OS=Homo sapiens OX=9606 GN=RPLP2 PE=1 SV=1 | 57.12 | 173.97 | 3.05 | 1.61 |
| sp\|P42766\|RL35_HUMAN 60S ribosomal protein L35 OS=Homo sapiens OX=9606 GN=RPL35 PE=1 SV=2 | 56.20 | 112.84 | 2.01 | 1.01 |
| tr\|H0YKD8\|H0YKD8_HUMAN 60S ribosomal protein L28 OS=Homo sapiens OX=9606 GN=RPL28 PE=1 SV=1 | 56.01 | 132.95 | 2.37 | 1.25 |
| sp\|P52292\|IMA1_HUMAN Importin subunit alpha-1 OS=Homo sapiens OX=9606 GN=KPNA2 PE=1 SV=1 | 55.40 | 155.16 | 2.80 | 1.49 |
| tr\|A0A0C4DFV9\|A0A0C4DFV9_HUMAN Protein SET OS=Homo sapiens OX=9606 GN=SET PE=1 SV=1 | 55.16 | 116.57 | 2.11 | 1.08 |

| | | | | |
|---|---|---|---|---|
| sp\|O75947\|ATP5H_HUMAN ATP synthase subunit d, mitochondrial OS=Homo sapiens OX=9606 GN=ATP5PD PE=1 SV=3 | 54.85 | 22.32 | 0.41 | -1.30 |
| sp\|P80723\|BASP1_HUMAN Brain acid soluble protein 1 OS=Homo sapiens OX=9606 GN=BASP1 PE=1 SV=2 | 54.55 | 43.88 | 0.80 | -0.31 |
| sp\|Q09028-3\|RBBP4_HUMAN Isoform 3 of Histone-binding protein RBBP4 OS=Homo sapiens OX=9606 GN=RBBP4 | 53.85 | 47.32 | 0.88 | -0.19 |
| sp\|P49721\|PSB2_HUMAN Proteasome subunit beta type-2 OS=Homo sapiens OX=9606 GN=PSMB2 PE=1 SV=1 | 53.12 | 47.18 | 0.89 | -0.17 |
| sp\|P62829\|RL23_HUMAN 60S ribosomal protein L23 OS=Homo sapiens OX=9606 GN=RPL23 PE=1 SV=1 | 52.89 | 122.28 | 2.31 | 1.21 |
| sp\|P13797\|PLST_HUMAN Plastin-3 OS=Homo sapiens OX=9606 GN=PLS3 PE=1 SV=4 | 52.87 | 118.31 | 2.24 | 1.16 |
| tr\|B1ANR0\|B1ANR0_HUMAN Polyadenylate-binding protein OS=Homo sapiens OX=9606 GN=PABPC4 PE=1 SV=1 | 52.78 | 167.06 | 3.17 | 1.66 |
| tr\|A0A087WX29\|A0A087WX29_HUMAN TAR DNA-binding protein 43 (Fragment) OS=Homo sapiens OX=9606 GN=TARDBP PE=1 SV=1 | 52.69 | 77.72 | 1.48 | 0.56 |
| sp\|P31153\|METK2_HUMAN S-adenosylmethionine synthase isoform type-2 OS=Homo sapiens OX=9606 GN=MAT2A PE=1 SV=1 | 52.42 | 59.29 | 1.13 | 0.18 |
| sp\|P62753\|RS6_HUMAN 40S ribosomal protein S6 OS=Homo sapiens OX=9606 GN=RPS6 PE=1 SV=1 | 52.31 | 104.91 | 2.01 | 1.00 |
| sp\|P61020\|RAB5B_HUMAN Ras-related protein Rab-5B OS=Homo sapiens OX=9606 GN=RAB5B PE=1 SV=1 | 52.23 | 152.75 | 2.92 | 1.55 |
| tr\|H3BNC9\|H3BNC9_HUMAN Uncharacterized protein OS=Homo sapiens OX=9606 PE=3 SV=2 | 52.18 | 117.48 | 2.25 | 1.17 |
| sp\|P04080\|CYTB_HUMAN Cystatin-B OS=Homo sapiens OX=9606 GN=CSTB PE=1 SV=2 | 51.77 | 162.84 | 3.15 | 1.65 |
| sp\|P25786-2\|PSA1_HUMAN Isoform Long of Proteasome subunit alpha type-1 OS=Homo sapiens OX=9606 GN=PSMA1 | 51.70 | 142.24 | 2.75 | 1.46 |
| sp\|P52209-2\|6PGD_HUMAN Isoform 2 of 6-phosphogluconate dehydrogenase, decarboxylating OS=Homo sapiens OX=9606 GN=PGD | 51.25 | 73.54 | 1.43 | 0.52 |
| tr\|H0YN26\|H0YN26_HUMAN Acidic leucine-rich nuclear phosphoprotein 32 family member A OS=Homo sapiens OX=9606 GN=ANP32A PE=1 SV=1 | 50.97 | 92.38 | 1.81 | 0.86 |
| sp\|Q01518\|CAP1_HUMAN Adenylyl cyclase-associated protein 1 OS=Homo sapiens OX=9606 GN=CAP1 PE=1 SV=5 | 50.83 | 84.07 | 1.65 | 0.73 |

| | | | | |
|---|---|---|---|---|
| sp\|P20290-2\|BTF3_HUMAN Isoform 2 of Transcription factor BTF3 OS=Homo sapiens OX=9606 GN=BTF3 | 50.61 | 167.34 | 3.31 | 1.73 |
| sp\|Q15233\|NONO_HUMAN Non-POU domain-containing octamer-binding protein OS=Homo sapiens OX=9606 GN=NONO PE=1 SV=4 | 50.50 | 58.67 | 1.16 | 0.22 |
| sp\|Q15181\|IPYR_HUMAN Inorganic pyrophosphatase OS=Homo sapiens OX=9606 GN=PPA1 PE=1 SV=2 | 50.48 | 147.88 | 2.93 | 1.55 |
| sp\|O60506\|HNRPQ_HUMAN Heterogeneous nuclear ribonucleoprotein Q OS=Homo sapiens OX=9606 GN=SYNCRIP PE=1 SV=2 | 50.48 | 99.16 | 1.96 | 0.97 |
| sp\|P39023\|RL3_HUMAN 60S ribosomal protein L3 OS=Homo sapiens OX=9606 GN=RPL3 PE=1 SV=2 | 50.40 | 50.64 | 1.00 | 0.01 |
| sp\|P31040\|SDHA_HUMAN Succinate dehydrogenase [ubiquinone] flavoprotein subunit, mitochondrial OS=Homo sapiens OX=9606 GN=SDHA PE=1 SV=2 | 50.36 | 48.28 | 0.96 | -0.06 |
| sp\|P55809\|SCOT1_HUMAN Succinyl-CoA:3-ketoacid coenzyme A transferase 1, mitochondrial OS=Homo sapiens OX=9606 GN=OXCT1 PE=1 SV=1 | 49.87 | 87.97 | 1.76 | 0.82 |
| tr\|A0A1W2PQB2\|A0A1W2PQB2_HUMAN BTB/POZ domain-containing protein KCTD7 (Fragment) OS=Homo sapiens OX=9606 GN=KCTD7 PE=1 SV=1 | 49.39 | 60.69 | 1.23 | 0.30 |
| sp\|Q04941-2\|PLP2_HUMAN Isoform 2 of Proteolipid protein 2 OS=Homo sapiens OX=9606 GN=PLP2 | 49.15 | 136.27 | 2.77 | 1.47 |
| tr\|A0A0D9SF53\|A0A0D9SF53_HUMAN ATP-dependent RNA helicase DDX3X OS=Homo sapiens OX=9606 GN=DDX3X PE=1 SV=1 | 48.86 | 103.77 | 2.12 | 1.09 |
| sp\|P51858\|HDGF_HUMAN Hepatoma-derived growth factor OS=Homo sapiens OX=9606 GN=HDGF PE=1 SV=1 | 48.47 | 188.96 | 3.90 | 1.96 |
| sp\|P48047\|ATPO_HUMAN ATP synthase subunit O, mitochondrial OS=Homo sapiens OX=9606 GN=ATP5PO PE=1 SV=1 | 48.08 | 142.34 | 2.96 | 1.57 |
| sp\|P11413-2\|G6PD_HUMAN Isoform Long of Glucose-6-phosphate 1-dehydrogenase OS=Homo sapiens OX=9606 GN=G6PD | 48.07 | 75.26 | 1.57 | 0.65 |
| sp\|Q9HB71\|CYBP_HUMAN Calcyclin-binding protein OS=Homo sapiens OX=9606 GN=CACYBP PE=1 SV=2 | 47.53 | 132.20 | 2.78 | 1.48 |
| sp\|Q07021\|C1QBP_HUMAN Complement component 1 Q subcomponent-binding protein, mitochondrial OS=Homo sapiens OX=9606 GN=C1QBP PE=1 SV=1 | 47.42 | 119.95 | 2.53 | 1.34 |
| sp\|P51148-2\|RAB5C_HUMAN Isoform 2 of Ras-related protein Rab-5C OS=Homo sapiens OX=9606 GN=RAB5C | 46.95 | 126.06 | 2.69 | 1.43 |

| | | | | |
|---|---|---|---|---|
| sp\|P28070\|PSB4_HUMAN Proteasome subunit beta type-4 OS=Homo sapiens OX=9606 GN=PSMB4 PE=1 SV=4 | 46.63 | 23.44 | 0.50 | -0.99 |
| tr\|A0A087WYT3\|A0A087WYT3_HUMAN Prostaglandin E synthase 3 OS=Homo sapiens OX=9606 GN=PTGES3 PE=1 SV=1 | 46.62 | 265.28 | 5.69 | 2.51 |
| sp\|Q13247-3\|SRSF6_HUMAN Isoform SRP55-3 of Serine/arginine-rich splicing factor 6 OS=Homo sapiens OX=9606 GN=SRSF6 | 46.61 | 87.18 | 1.87 | 0.90 |
| sp\|Q15907-2\|RB11B_HUMAN Isoform 2 of Ras-related protein Rab-11B OS=Homo sapiens OX=9606 GN=RAB11B | 46.57 | 71.54 | 1.54 | 0.62 |
| sp\|P36871\|PGM1_HUMAN Phosphoglucomutase-1 OS=Homo sapiens OX=9606 GN=PGM1 PE=1 SV=3 | 46.33 | 208.28 | 4.50 | 2.17 |
| tr\|G3V5Z7\|G3V5Z7_HUMAN Proteasome subunit alpha type OS=Homo sapiens OX=9606 GN=PSMA6 PE=1 SV=1 | 46.31 | 194.77 | 4.21 | 2.07 |
| sp\|P41091\|IF2G_HUMAN Eukaryotic translation initiation factor 2 subunit 3 OS=Homo sapiens OX=9606 GN=EIF2S3 PE=1 SV=3 | 46.31 | 57.02 | 1.23 | 0.30 |
| sp\|P06753-2\|TPM3_HUMAN Isoform 2 of Tropomyosin alpha-3 chain OS=Homo sapiens OX=9606 GN=TPM3 | 45.97 | 42.47 | 0.92 | -0.11 |
| tr\|E5RIW3\|E5RIW3_HUMAN Tubulin-specific chaperone A OS=Homo sapiens OX=9606 GN=TBCA PE=1 SV=1 | 45.48 | 19.27 | 0.42 | -1.24 |
| tr\|J3KQN4\|J3KQN4_HUMAN 60S ribosomal protein L36a OS=Homo sapiens OX=9606 GN=RPL36A PE=1 SV=1 | 45.24 | 79.42 | 1.76 | 0.81 |
| sp\|P34897-2\|GLYM_HUMAN Isoform 2 of Serine hydroxymethyltransferase, mitochondrial OS=Homo sapiens OX=9606 GN=SHMT2 | 45.14 | 93.99 | 2.08 | 1.06 |
| tr\|H0YIZ0\|H0YIZ0_HUMAN Serine hydroxymethyltransferase, mitochondrial (Fragment) OS=Homo sapiens OX=9606 GN=SHMT2 PE=1 SV=1 | 45.14 | 93.99 | 2.08 | 1.06 |
| sp\|O00303\|EIF3F_HUMAN Eukaryotic translation initiation factor 3 subunit F OS=Homo sapiens OX=9606 GN=EIF3F PE=1 SV=1 | 44.78 | 87.19 | 1.95 | 0.96 |
| sp\|O00148\|DX39A_HUMAN ATP-dependent RNA helicase DDX39A OS=Homo sapiens OX=9606 GN=DDX39A PE=1 SV=2 | 44.38 | 62.36 | 1.40 | 0.49 |
| sp\|P21266\|GSTM3_HUMAN Glutathione S-transferase Mu 3 OS=Homo sapiens OX=9606 GN=GSTM3 PE=1 SV=3 | 44.19 | 95.41 | 2.16 | 1.11 |
| tr\|K7EK33\|K7EK33_HUMAN DAZ-associated protein 1 OS=Homo sapiens OX=9606 GN=DAZAP1 PE=1 SV=2 | 44.11 | 68.23 | 1.55 | 0.63 |

| | | | | |
|---|---|---|---|---|
| sp\|P61586\|RHOA_HUMAN Transforming protein RhoA OS=Homo sapiens OX=9606 GN=RHOA PE=1 SV=1 | 44.08 | 111.82 | 2.54 | 1.34 |
| sp\|P05455\|LA_HUMAN Lupus La protein OS=Homo sapiens OX=9606 GN=SSB PE=1 SV=2 | 43.48 | 36.03 | 0.83 | -0.27 |
| sp\|P25789\|PSA4_HUMAN Proteasome subunit alpha type-4 OS=Homo sapiens OX=9606 GN=PSMA4 PE=1 SV=1 | 43.14 | 95.46 | 2.21 | 1.15 |
| sp\|P56537\|IF6_HUMAN Eukaryotic translation initiation factor 6 OS=Homo sapiens OX=9606 GN=EIF6 PE=1 SV=1 | 42.81 | 122.79 | 2.87 | 1.52 |
| sp\|P02786\|TFR1_HUMAN Transferrin receptor protein 1 OS=Homo sapiens OX=9606 GN=TFRC PE=1 SV=2 | 42.71 | 120.22 | 2.82 | 1.49 |
| sp\|P12004\|PCNA_HUMAN Proliferating cell nuclear antigen OS=Homo sapiens OX=9606 GN=PCNA PE=1 SV=1 | 42.40 | 113.60 | 2.68 | 1.42 |
| tr\|C9JFR7\|C9JFR7_HUMAN Cytochrome c (Fragment) OS=Homo sapiens OX=9606 GN=CYCS PE=1 SV=1 | 42.39 | 133.96 | 3.16 | 1.66 |
| tr\|R4GNH3\|R4GNH3_HUMAN 26S proteasome regulatory subunit 6A OS=Homo sapiens OX=9606 GN=PSMC3 PE=1 SV=1 | 42.24 | 50.40 | 1.19 | 0.25 |
| tr\|A0A0B4J239\|A0A0B4J239_HUMAN Dolichyl-diphosphooligosaccharide--protein glycosyltransferase subunit DAD1 OS=Homo sapiens OX=9606 GN=DAD1 PE=1 SV=1 | 41.96 | 111.35 | 2.65 | 1.41 |
| tr\|J3KTL2\|J3KTL2_HUMAN Serine/arginine-rich-splicing factor 1 OS=Homo sapiens OX=9606 GN=SRSF1 PE=1 SV=1 | 41.91 | 67.74 | 1.62 | 0.69 |
| sp\|P37837\|TALDO_HUMAN Transaldolase OS=Homo sapiens OX=9606 GN=TALDO1 PE=1 SV=2 | 41.44 | 142.26 | 3.43 | 1.78 |
| sp\|P08754\|GNAI3_HUMAN Guanine nucleotide-binding protein G(k) subunit alpha OS=Homo sapiens OX=9606 GN=GNAI3 PE=1 SV=3 | 41.31 | 111.31 | 2.69 | 1.43 |
| sp\|O60888-2\|CUTA_HUMAN Isoform A of Protein CutA OS=Homo sapiens OX=9606 GN=CUTA | 41.28 | 139.34 | 3.38 | 1.76 |
| tr\|Q5JR08\|Q5JR08_HUMAN Rho-related GTP-binding protein RhoC (Fragment) OS=Homo sapiens OX=9606 GN=RHOC PE=1 SV=8 | 41.23 | 104.13 | 2.53 | 1.34 |
| sp\|P68036-3\|UB2L3_HUMAN Isoform 3 of Ubiquitin-conjugating enzyme E2 L3 OS=Homo sapiens OX=9606 GN=UBE2L3 | 40.98 | 227.60 | 5.55 | 2.47 |
| sp\|Q9Y5M8\|SRPRB_HUMAN Signal recognition particle receptor subunit beta OS=Homo sapiens OX=9606 GN=SRPRB PE=1 SV=3 | 40.90 | 38.49 | 0.94 | -0.09 |
| sp\|P60468\|SC61B_HUMAN Protein transport protein Sec61 subunit beta OS=Homo sapiens OX=9606 GN=SEC61B PE=1 SV=2 | 40.69 | 23.24 | 0.57 | -0.81 |

| | | | | |
|---|---|---|---|---|
| tr\|C9IZQ1\|C9IZQ1_HUMAN Translocon-associated protein subunit alpha OS=Homo sapiens OX=9606 GN=SSR1 PE=1 SV=1 | 40.33 | 87.89 | 2.18 | 1.12 |
| sp\|P46778\|RL21_HUMAN 60S ribosomal protein L21 OS=Homo sapiens OX=9606 GN=RPL21 PE=1 SV=2 | 39.88 | 138.01 | 3.46 | 1.79 |
| tr\|A0A075B7D9\|A0A075B7D9_HUMAN TATA-binding protein-associated factor 2N OS=Homo sapiens OX=9606 GN=TAF15 PE=1 SV=1 | 39.88 | 6.39 | 0.16 | -2.64 |
| sp\|P61163\|ACTZ_HUMAN Alpha-centractin OS=Homo sapiens OX=9606 GN=ACTR1A PE=1 SV=1 | 39.77 | 67.22 | 1.69 | 0.76 |
| sp\|P31948\|STIP1_HUMAN Stress-induced-phosphoprotein 1 OS=Homo sapiens OX=9606 GN=STIP1 PE=1 SV=1 | 39.61 | 88.35 | 2.23 | 1.16 |
| sp\|Q99536-3\|VAT1_HUMAN Isoform 3 of Synaptic vesicle membrane protein VAT-1 homolog OS=Homo sapiens OX=9606 GN=VAT1 | 39.58 | 63.00 | 1.59 | 0.67 |
| sp\|P24539\|AT5F1_HUMAN ATP synthase F(0) complex subunit B1, mitochondrial OS=Homo sapiens OX=9606 GN=ATP5PB PE=1 SV=2 | 39.20 | 125.24 | 3.19 | 1.68 |
| sp\|Q96QD8-2\|S38A2_HUMAN Isoform 2 of Sodium-coupled neutral amino acid transporter 2 OS=Homo sapiens OX=9606 GN=SLC38A2 | 38.92 | 67.96 | 1.75 | 0.80 |
| sp\|P11908-2\|PRPS2_HUMAN Isoform 2 of Ribose-phosphate pyrophosphokinase 2 OS=Homo sapiens OX=9606 GN=PRPS2 | 38.91 | 133.83 | 3.44 | 1.78 |
| tr\|B1ALA9\|B1ALA9_HUMAN Ribose-phosphate pyrophosphokinase 1 OS=Homo sapiens OX=9606 GN=PRPS1 PE=1 SV=2 | 38.91 | 133.83 | 3.44 | 1.78 |
| sp\|P28072\|PSB6_HUMAN Proteasome subunit beta type-6 OS=Homo sapiens OX=9606 GN=PSMB6 PE=1 SV=4 | 38.90 | 46.64 | 1.20 | 0.26 |
| sp\|P06493\|CDK1_HUMAN Cyclin-dependent kinase 1 OS=Homo sapiens OX=9606 GN=CDK1 PE=1 SV=3 | 38.85 | 62.35 | 1.61 | 0.68 |
| tr\|K7EQA9\|K7EQA9_HUMAN Hsp90 co-chaperone Cdc37 (Fragment) OS=Homo sapiens OX=9606 GN=CDC37 PE=1 SV=1 | 38.78 | 35.41 | 0.91 | -0.13 |
| sp\|P13010\|XRCC5_HUMAN X-ray repair cross-complementing protein 5 OS=Homo sapiens OX=9606 GN=XRCC5 PE=1 SV=3 | 38.78 | 73.05 | 1.88 | 0.91 |
| sp\|P61289-2\|PSME3_HUMAN Isoform 2 of Proteasome activator complex subunit 3 OS=Homo sapiens OX=9606 GN=PSME3 | 38.60 | 83.00 | 2.15 | 1.10 |
| tr\|Q5T4U5\|Q5T4U5_HUMAN Acyl-Coenzyme A dehydrogenase, C-4 to C-12 straight chain, isoform CRA_a OS=Homo sapiens OX=9606 GN=ACADM PE=1 SV=1 | 38.37 | 76.28 | 1.99 | 0.99 |
| tr\|E9PEB5\|E9PEB5_HUMAN Far upstream element-binding protein 1 OS=Homo sapiens OX=9606 GN=FUBP1 PE=1 SV=1 | 38.21 | 114.64 | 3.00 | 1.58 |

| | | | | |
|---|---|---|---|---|
| sp\|P23921\|RIR1_HUMAN Ribonucleoside-diphosphate reductase large subunit OS=Homo sapiens OX=9606 GN=RRM1 PE=1 SV=1 | 37.78 | 47.14 | 1.25 | 0.32 |
| sp\|O60664-4\|PLIN3_HUMAN Isoform 4 of Perilipin-3 OS=Homo sapiens OX=9606 GN=PLIN3 | 37.63 | 89.21 | 2.37 | 1.25 |
| sp\|Q13283\|G3BP1_HUMAN Ras GTPase-activating protein-binding protein 1 OS=Homo sapiens OX=9606 GN=G3BP1 PE=1 SV=1 | 37.47 | 60.57 | 1.62 | 0.69 |
| sp\|Q06323\|PSME1_HUMAN Proteasome activator complex subunit 1 OS=Homo sapiens OX=9606 GN=PSME1 PE=1 SV=1 | 37.21 | 37.81 | 1.02 | 0.02 |
| sp\|P60953\|CDC42_HUMAN Cell division control protein 42 homolog OS=Homo sapiens OX=9606 GN=CDC42 PE=1 SV=2 | 37.20 | 70.18 | 1.89 | 0.92 |
| tr\|A0A1W2PQ51\|A0A1W2PQ51_HUMAN Probable ATP-dependent RNA helicase DDX17 OS=Homo sapiens OX=9606 GN=DDX17 PE=1 SV=1 | 37.18 | 92.61 | 2.49 | 1.32 |
| sp\|P00403\|COX2_HUMAN Cytochrome c oxidase subunit 2 OS=Homo sapiens OX=9606 GN=MT-CO2 PE=1 SV=1 | 37.02 | 37.38 | 1.01 | 0.01 |
| sp\|Q13185\|CBX3_HUMAN Chromobox protein homolog 3 OS=Homo sapiens OX=9606 GN=CBX3 PE=1 SV=4 | 36.99 | 105.06 | 2.84 | 1.51 |
| sp\|P05161\|ISG15_HUMAN Ubiquitin-like protein ISG15 OS=Homo sapiens OX=9606 GN=ISG15 PE=1 SV=5 | 36.84 | 110.75 | 3.01 | 1.59 |
| sp\|P62136\|PP1A_HUMAN Serine/threonine-protein phosphatase PP1-alpha catalytic subunit OS=Homo sapiens OX=9606 GN=PPP1CA PE=1 SV=1 | 36.77 | 79.62 | 2.17 | 1.11 |
| tr\|F8VYE8\|F8VYE8_HUMAN Serine/threonine-protein phosphatase OS=Homo sapiens OX=9606 GN=PPP1CC PE=1 SV=1 | 36.77 | 79.62 | 2.17 | 1.11 |
| sp\|P62140\|PP1B_HUMAN Serine/threonine-protein phosphatase PP1-beta catalytic subunit OS=Homo sapiens OX=9606 GN=PPP1CB PE=1 SV=3 | 36.77 | 79.62 | 2.17 | 1.11 |
| sp\|O75340\|PDCD6_HUMAN Programmed cell death protein 6 OS=Homo sapiens OX=9606 GN=PDCD6 PE=1 SV=1 | 36.70 | 115.93 | 3.16 | 1.66 |
| sp\|P13804-2\|ETFA_HUMAN Isoform 2 of Electron transfer flavoprotein subunit alpha, mitochondrial OS=Homo sapiens OX=9606 GN=ETFA | 36.68 | 67.97 | 1.85 | 0.89 |
| tr\|G3V325\|G3V325_HUMAN ATP5MF-PTCD1 readthrough OS=Homo sapiens OX=9606 GN=ATP5MF-PTCD1 PE=4 SV=1 | 36.51 | 85.83 | 2.35 | 1.23 |
| sp\|Q99714\|HCD2_HUMAN 3-hydroxyacyl-CoA dehydrogenase type-2 OS=Homo sapiens OX=9606 GN=HSD17B10 PE=1 SV=3 | 36.48 | 115.50 | 3.17 | 1.66 |

| | | | | |
|---|---|---|---|---|
| sp\|Q9UBR2\|CATZ_HUMAN Cathepsin Z OS=Homo sapiens OX=9606 GN=CTSZ PE=1 SV=1 | 36.10 | 27.66 | 0.77 | -0.38 |
| sp\|Q9Y3F4-2\|STRAP_HUMAN Isoform 2 of Serine-threonine kinase receptor-associated protein OS=Homo sapiens OX=9606 GN=STRAP | 36.06 | 94.78 | 2.63 | 1.39 |
| sp\|Q9BS40\|LXN_HUMAN Latexin OS=Homo sapiens OX=9606 GN=LXN PE=1 SV=2 | 35.56 | 99.28 | 2.79 | 1.48 |
| sp\|P28074\|PSB5_HUMAN Proteasome subunit beta type-5 OS=Homo sapiens OX=9606 GN=PSMB5 PE=1 SV=3 | 35.48 | 93.25 | 2.63 | 1.39 |
| sp\|P53999\|TCP4_HUMAN Activated RNA polymerase II transcriptional coactivator p15 OS=Homo sapiens OX=9606 GN=SUB1 PE=1 SV=3 | 35.37 | 117.21 | 3.31 | 1.73 |
| tr\|A0A024RA52\|A0A024RA52_HUMAN Proteasome subunit alpha type OS=Homo sapiens OX=9606 GN=PSMA2 PE=1 SV=1 | 35.30 | 107.51 | 3.05 | 1.61 |
| sp\|P49720\|PSB3_HUMAN Proteasome subunit beta type-3 OS=Homo sapiens OX=9606 GN=PSMB3 PE=1 SV=2 | 35.19 | 112.85 | 3.21 | 1.68 |
| sp\|P52907\|CAZA1_HUMAN F-actin-capping protein subunit alpha-1 OS=Homo sapiens OX=9606 GN=CAPZA1 PE=1 SV=3 | 34.90 | 72.85 | 2.09 | 1.06 |
| sp\|P21291\|CSRP1_HUMAN Cysteine and glycine-rich protein 1 OS=Homo sapiens OX=9606 GN=CSRP1 PE=1 SV=3 | 34.89 | 112.66 | 3.23 | 1.69 |
| sp\|P62834\|RAP1A_HUMAN Ras-related protein Rap-1A OS=Homo sapiens OX=9606 GN=RAP1A PE=1 SV=1 | 34.68 | 81.75 | 2.36 | 1.24 |
| sp\|P17812\|PYRG1_HUMAN CTP synthase 1 OS=Homo sapiens OX=9606 GN=CTPS1 PE=1 SV=2 | 34.67 | 45.27 | 1.31 | 0.39 |
| sp\|A6NHL2-2\|TBAL3_HUMAN Isoform 2 of Tubulin alpha chain-like 3 OS=Homo sapiens OX=9606 GN=TUBAL3 | 34.59 | 105.86 | 3.06 | 1.61 |
| sp\|Q15645\|PCH2_HUMAN Pachytene checkpoint protein 2 homolog OS=Homo sapiens OX=9606 GN=TRIP13 PE=1 SV=2 | 34.57 | 57.51 | 1.66 | 0.73 |
| sp\|O14929\|HAT1_HUMAN Histone acetyltransferase type B catalytic subunit OS=Homo sapiens OX=9606 GN=HAT1 PE=1 SV=1 | 34.47 | 48.98 | 1.42 | 0.51 |
| sp\|P41250\|GARS_HUMAN Glycine--tRNA ligase OS=Homo sapiens OX=9606 GN=GARS PE=1 SV=3 | 34.37 | 75.21 | 2.19 | 1.13 |
| sp\|P30044-2\|PRDX5_HUMAN Isoform Cytoplasmic+peroxisomal of Peroxiredoxin-5, mitochondrial OS=Homo sapiens OX=9606 GN=PRDX5 | 34.32 | 110.58 | 3.22 | 1.69 |
| sp\|P00387-2\|NB5R3_HUMAN Isoform 2 of NADH-cytochrome b5 reductase 3 OS=Homo sapiens OX=9606 GN=CYB5R3 | 34.29 | 47.12 | 1.37 | 0.46 |

| | | | | |
|---|---|---|---|---|
| sp\|P22695\|QCR2_HUMAN Cytochrome b-c1 complex subunit 2, mitochondrial OS=Homo sapiens OX=9606 GN=UQCRC2 PE=1 SV=3 | 34.28 | 63.70 | 1.86 | 0.89 |
| sp\|Q04837\|SSBP_HUMAN Single-stranded DNA-binding protein, mitochondrial OS=Homo sapiens OX=9606 GN=SSBP1 PE=1 SV=1 | 33.97 | 101.92 | 3.00 | 1.59 |
| sp\|Q9UNL2-2\|SSRG_HUMAN Isoform 2 of Translocon-associated protein subunit gamma OS=Homo sapiens OX=9606 GN=SSR3 | 33.94 | 56.14 | 1.65 | 0.73 |
| sp\|Q10471\|GALT2_HUMAN Polypeptide N-acetylgalactosaminyltransferase 2 OS=Homo sapiens OX=9606 GN=GALNT2 PE=1 SV=1 | 33.92 | 49.22 | 1.45 | 0.54 |
| sp\|Q92769-3\|HDAC2_HUMAN Isoform 2 of Histone deacetylase 2 OS=Homo sapiens OX=9606 GN=HDAC2 | 33.83 | 76.82 | 2.27 | 1.18 |
| sp\|Q13547\|HDAC1_HUMAN Histone deacetylase 1 OS=Homo sapiens OX=9606 GN=HDAC1 PE=1 SV=1 | 33.83 | 32.50 | 0.96 | -0.06 |
| sp\|O75534-2\|CSDE1_HUMAN Isoform 2 of Cold shock domain-containing protein E1 OS=Homo sapiens OX=9606 GN=CSDE1 | 33.70 | 47.93 | 1.42 | 0.51 |
| sp\|Q7L1Q6-3\|BZW1_HUMAN Isoform 3 of Basic leucine zipper and W2 domain-containing protein 1 OS=Homo sapiens OX=9606 GN=BZW1 | 33.62 | 51.80 | 1.54 | 0.62 |
| sp\|P60903\|S10AA_HUMAN Protein S100-A10 OS=Homo sapiens OX=9606 GN=S100A10 PE=1 SV=2 | 33.57 | 16.12 | 0.48 | -1.06 |
| sp\|Q10589-2\|BST2_HUMAN Isoform 2 of Bone marrow stromal antigen 2 OS=Homo sapiens OX=9606 GN=BST2 | 33.48 | 70.20 | 2.10 | 1.07 |
| tr\|A0A087WYS1\|A0A087WYS1_HUMAN UTP--glucose-1-phosphate uridylyltransferase OS=Homo sapiens OX=9606 GN=UGP2 PE=1 SV=1 | 33.45 | 47.52 | 1.42 | 0.51 |
| sp\|O96008\|TOM40_HUMAN Mitochondrial import receptor subunit TOM40 homolog OS=Homo sapiens OX=9606 GN=TOMM40 PE=1 SV=1 | 32.90 | 73.45 | 2.23 | 1.16 |
| sp\|Q15631\|TSN_HUMAN Translin OS=Homo sapiens OX=9606 GN=TSN PE=1 SV=1 | 32.23 | 36.25 | 1.12 | 0.17 |
| sp\|Q15293\|RCN1_HUMAN Reticulocalbin-1 OS=Homo sapiens OX=9606 GN=RCN1 PE=1 SV=1 | 32.05 | 47.76 | 1.49 | 0.58 |
| sp\|O43390\|HNRPR_HUMAN Heterogeneous nuclear ribonucleoprotein R OS=Homo sapiens OX=9606 GN=HNRNPR PE=1 SV=1 | 31.94 | 68.46 | 2.14 | 1.10 |
| sp\|Q15056-2\|IF4H_HUMAN Isoform Short of Eukaryotic translation initiation factor 4H OS=Homo sapiens OX=9606 GN=EIF4H | 31.82 | 103.55 | 3.25 | 1.70 |
| sp\|P07741\|APT_HUMAN Adenine phosphoribosyltransferase OS=Homo sapiens OX=9606 GN=APRT PE=1 SV=2 | 31.81 | 76.40 | 2.40 | 1.26 |

| | | | | |
|---|---|---|---|---|
| tr\|F6WQW2\|F6WQW2_HUMAN Ran-specific GTPase-activating protein OS=Homo sapiens OX=9606 GN=RANBP1 PE=1 SV=1 | 31.81 | 49.77 | 1.56 | 0.65 |
| sp\|Q9Y230\|RUVB2_HUMAN RuvB-like 2 OS=Homo sapiens OX=9606 GN=RUVBL2 PE=1 SV=3 | 31.78 | 55.10 | 1.73 | 0.79 |
| sp\|P61088\|UBE2N_HUMAN Ubiquitin-conjugating enzyme E2 N OS=Homo sapiens OX=9606 GN=UBE2N PE=1 SV=1 | 31.74 | 126.21 | 3.98 | 1.99 |
| sp\|P26447\|S10A4_HUMAN Protein S100-A4 OS=Homo sapiens OX=9606 GN=S100A4 PE=1 SV=1 | 31.69 | 28.12 | 0.89 | -0.17 |
| sp\|Q96HV5\|TM41A_HUMAN Transmembrane protein 41A OS=Homo sapiens OX=9606 GN=TMEM41A PE=1 SV=1 | 31.61 | 24.28 | 0.77 | -0.38 |
| sp\|Q7Z7H5-2\|TMED4_HUMAN Isoform 2 of Transmembrane emp24 domain-containing protein 4 OS=Homo sapiens OX=9606 GN=TMED4 | 31.38 | 19.69 | 0.63 | -0.67 |
| sp\|Q13155\|AIMP2_HUMAN Aminoacyl tRNA synthase complex-interacting multifunctional protein 2 OS=Homo sapiens OX=9606 GN=AIMP2 PE=1 SV=2 | 31.38 | 90.44 | 2.88 | 1.53 |
| tr\|A6NFX8\|A6NFX8_HUMAN ADP-sugar pyrophosphatase OS=Homo sapiens OX=9606 GN=NUDT5 PE=1 SV=1 | 31.35 | 76.31 | 2.43 | 1.28 |
| tr\|A0A087WTS4\|A0A087WTS4_HUMAN Peptidyl-prolyl cis-trans isomerase FKBP1A OS=Homo sapiens OX=9606 GN=FKBP1A PE=4 SV=1 | 31.11 | 20.30 | 0.65 | -0.62 |
| sp\|Q9Y617\|SERC_HUMAN Phosphoserine aminotransferase OS=Homo sapiens OX=9606 GN=PSAT1 PE=1 SV=2 | 31.10 | 71.45 | 2.30 | 1.20 |
| tr\|E9PN17\|E9PN17_HUMAN ATP synthase subunit g, mitochondrial OS=Homo sapiens OX=9606 GN=ATP5MG PE=1 SV=1 | 31.04 | 92.66 | 2.99 | 1.58 |
| sp\|P58546\|MTPN_HUMAN Myotrophin OS=Homo sapiens OX=9606 GN=MTPN PE=1 SV=2 | 31.02 | 89.92 | 2.90 | 1.54 |
| sp\|P37108\|SRP14_HUMAN Signal recognition particle 14 kDa protein OS=Homo sapiens OX=9606 GN=SRP14 PE=1 SV=2 | 30.95 | 75.39 | 2.44 | 1.28 |
| sp\|P60228\|EIF3E_HUMAN Eukaryotic translation initiation factor 3 subunit E OS=Homo sapiens OX=9606 GN=EIF3E PE=1 SV=1 | 30.69 | 55.14 | 1.80 | 0.85 |
| sp\|Q9Y265\|RUVB1_HUMAN RuvB-like 1 OS=Homo sapiens OX=9606 GN=RUVBL1 PE=1 SV=1 | 30.63 | 65.04 | 2.12 | 1.09 |
| sp\|P63167\|DYL1_HUMAN Dynein light chain 1, cytoplasmic OS=Homo sapiens OX=9606 GN=DYNLL1 PE=1 SV=1 | 30.47 | 25.73 | 0.84 | -0.24 |

| | | | | |
|---|---|---|---|---|
| sp\|P63000-2\|RAC1_HUMAN Isoform B of Ras-related C3 botulinum toxin substrate 1 OS=Homo sapiens OX=9606 GN=RAC1 | 30.35 | 49.87 | 1.64 | 0.72 |
| sp\|P12268\|IMDH2_HUMAN Inosine-5'-monophosphate dehydrogenase 2 OS=Homo sapiens OX=9606 GN=IMPDH2 PE=1 SV=2 | 30.19 | 54.42 | 1.80 | 0.85 |
| sp\|O43809\|CPSF5_HUMAN Cleavage and polyadenylation specificity factor subunit 5 OS=Homo sapiens OX=9606 GN=NUDT21 PE=1 SV=1 | 30.15 | 83.35 | 2.76 | 1.47 |
| tr\|A0A0J9YVP6\|A0A0J9YVP6_HUMAN Poly(U)-binding-splicing factor PUF60 (Fragment) OS=Homo sapiens OX=9606 GN=PUF60 PE=1 SV=1 | 29.99 | 72.18 | 2.41 | 1.27 |
| tr\|B7Z7P8\|B7Z7P8_HUMAN Eukaryotic peptide chain release factor subunit 1 OS=Homo sapiens OX=9606 GN=ETF1 PE=1 SV=1 | 29.92 | 45.64 | 1.53 | 0.61 |
| sp\|P52565\|GDIR1_HUMAN Rho GDP-dissociation inhibitor 1 OS=Homo sapiens OX=9606 GN=ARHGDIA PE=1 SV=3 | 29.65 | 68.44 | 2.31 | 1.21 |
| sp\|Q9NR31\|SAR1A_HUMAN GTP-binding protein SAR1a OS=Homo sapiens OX=9606 GN=SAR1A PE=1 SV=1 | 29.61 | 90.97 | 3.07 | 1.62 |
| sp\|P14866\|HNRPL_HUMAN Heterogeneous nuclear ribonucleoprotein L OS=Homo sapiens OX=9606 GN=HNRNPL PE=1 SV=2 | 29.52 | 66.25 | 2.24 | 1.17 |
| sp\|P20618\|PSB1_HUMAN Proteasome subunit beta type-1 OS=Homo sapiens OX=9606 GN=PSMB1 PE=1 SV=2 | 29.42 | 41.49 | 1.41 | 0.50 |
| sp\|P24752\|THIL_HUMAN Acetyl-CoA acetyltransferase, mitochondrial OS=Homo sapiens OX=9606 GN=ACAT1 PE=1 SV=1 | 29.36 | 70.43 | 2.40 | 1.26 |
| sp\|P00367\|DHE3_HUMAN Glutamate dehydrogenase 1, mitochondrial OS=Homo sapiens OX=9606 GN=GLUD1 PE=1 SV=2 | 29.30 | 43.80 | 1.50 | 0.58 |
| tr\|B0QY89\|B0QY89_HUMAN Eukaryotic translation initiation factor 3 subunit L OS=Homo sapiens OX=9606 GN=EIF3L PE=1 SV=1 | 29.29 | 67.18 | 2.29 | 1.20 |
| sp\|P08574\|CY1_HUMAN Cytochrome c1, heme protein, mitochondrial OS=Homo sapiens OX=9606 GN=CYC1 PE=1 SV=3 | 29.26 | 43.45 | 1.48 | 0.57 |
| tr\|C9J381\|C9J381_HUMAN Inosine-5'-monophosphate dehydrogenase OS=Homo sapiens OX=9606 GN=IMPDH1 PE=1 SV=1 | 29.22 | 7.95 | 0.27 | -1.88 |
| sp\|P32322\|P5CR1_HUMAN Pyrroline-5-carboxylate reductase 1, mitochondrial OS=Homo sapiens OX=9606 GN=PYCR1 PE=1 SV=2 | 29.20 | 83.78 | 2.87 | 1.52 |
| sp\|P30040\|ERP29_HUMAN Endoplasmic reticulum resident protein 29 OS=Homo sapiens OX=9606 GN=ERP29 PE=1 SV=4 | 29.13 | 63.67 | 2.19 | 1.13 |

| | | | | |
|---|---|---|---|---|
| sp\|Q07065\|CKAP4_HUMAN Cytoskeleton-associated protein 4 OS=Homo sapiens OX=9606 GN=CKAP4 PE=1 SV=2 | 28.87 | 67.47 | 2.34 | 1.22 |
| sp\|P31939-2\|PUR9_HUMAN Isoform 2 of Bifunctional purine biosynthesis protein PURH OS=Homo sapiens OX=9606 GN=ATIC | 28.81 | 57.24 | 1.99 | 0.99 |
| sp\|P04843\|RPN1_HUMAN Dolichyl-diphosphooligosaccharide--protein glycosyltransferase subunit 1 OS=Homo sapiens OX=9606 GN=RPN1 PE=1 SV=1 | 28.67 | 61.94 | 2.16 | 1.11 |
| sp\|P38117-2\|ETFB_HUMAN Isoform 2 of Electron transfer flavoprotein subunit beta OS=Homo sapiens OX=9606 GN=ETFB | 28.63 | 32.91 | 1.15 | 0.20 |
| sp\|P31942-2\|HNRH3_HUMAN Isoform 2 of Heterogeneous nuclear ribonucleoprotein H3 OS=Homo sapiens OX=9606 GN=HNRNPH3 | 28.53 | 80.76 | 2.83 | 1.50 |
| sp\|Q9Y5S9-2\|RBM8A_HUMAN Isoform 2 of RNA-binding protein 8A OS=Homo sapiens OX=9606 GN=RBM8A | 28.43 | 53.30 | 1.88 | 0.91 |
| tr\|A8K878\|A8K878_HUMAN Mesencephalic astrocyte-derived neurotrophic factor OS=Homo sapiens OX=9606 GN=MANF PE=1 SV=1 | 28.40 | 52.14 | 1.84 | 0.88 |
| tr\|H0Y2P0\|H0Y2P0_HUMAN CD44 antigen (Fragment) OS=Homo sapiens OX=9606 GN=CD44 PE=1 SV=1 | 28.29 | 38.23 | 1.35 | 0.43 |
| sp\|Q13347\|EIF3I_HUMAN Eukaryotic translation initiation factor 3 subunit I OS=Homo sapiens OX=9606 GN=EIF3I PE=1 SV=1 | 28.28 | 38.06 | 1.35 | 0.43 |
| tr\|X1WI28\|X1WI28_HUMAN 60S ribosomal protein L10 (Fragment) OS=Homo sapiens OX=9606 GN=RPL10 PE=1 SV=7 | 28.17 | 57.70 | 2.05 | 1.03 |
| sp\|O60701\|UGDH_HUMAN UDP-glucose 6-dehydrogenase OS=Homo sapiens OX=9606 GN=UGDH PE=1 SV=1 | 28.13 | 32.23 | 1.15 | 0.20 |
| sp\|P17931\|LEG3_HUMAN Galectin-3 OS=Homo sapiens OX=9606 GN=LGALS3 PE=1 SV=5 | 28.05 | 65.94 | 2.35 | 1.23 |
| sp\|Q9Y2T3-3\|GUAD_HUMAN Isoform 3 of Guanine deaminase OS=Homo sapiens OX=9606 GN=GDA | 27.94 | 80.34 | 2.88 | 1.52 |
| sp\|P78417\|GSTO1_HUMAN Glutathione S-transferase omega-1 OS=Homo sapiens OX=9606 GN=GSTO1 PE=1 SV=2 | 27.90 | 86.27 | 3.09 | 1.63 |
| sp\|P16083\|NQO2_HUMAN Ribosyldihydronicotinamide dehydrogenase [quinone] OS=Homo sapiens OX=9606 GN=NQO2 PE=1 SV=5 | 27.70 | 14.90 | 0.54 | -0.89 |
| sp\|O60869-2\|EDF1_HUMAN Isoform 2 of Endothelial differentiation-related factor 1 OS=Homo sapiens OX=9606 GN=EDF1 | 27.42 | 31.88 | 1.16 | 0.22 |

| | | | | |
|---|---|---|---|---|
| sp\|P62266\|RS23_HUMAN 40S ribosomal protein S23 OS=Homo sapiens OX=9606 GN=RPS23 PE=1 SV=3 | 27.33 | 29.83 | 1.09 | 0.13 |
| sp\|Q99733-2\|NP1L4_HUMAN Isoform 2 of Nucleosome assembly protein 1-like 4 OS=Homo sapiens OX=9606 GN=NAP1L4 | 27.13 | 144.07 | 5.31 | 2.41 |
| sp\|Q96QK1\|VPS35_HUMAN Vacuolar protein sorting-associated protein 35 OS=Homo sapiens OX=9606 GN=VPS35 PE=1 SV=2 | 26.81 | 66.47 | 2.48 | 1.31 |
| sp\|P62314\|SMD1_HUMAN Small nuclear ribonucleoprotein Sm D1 OS=Homo sapiens OX=9606 GN=SNRPD1 PE=1 SV=1 | 26.80 | 48.18 | 1.80 | 0.85 |
| sp\|O00483\|NDUA4_HUMAN Cytochrome c oxidase subunit NDUFA4 OS=Homo sapiens OX=9606 GN=NDUFA4 PE=1 SV=1 | 26.62 | 44.07 | 1.66 | 0.73 |
| tr\|K7ELL7\|K7ELL7_HUMAN Glucosidase 2 subunit beta OS=Homo sapiens OX=9606 GN=PRKCSH PE=1 SV=1 | 26.51 | 99.29 | 3.75 | 1.91 |
| sp\|P54577\|SYYC_HUMAN Tyrosine--tRNA ligase, cytoplasmic OS=Homo sapiens OX=9606 GN=YARS PE=1 SV=4 | 26.48 | 68.82 | 2.60 | 1.38 |
| sp\|P63208\|SKP1_HUMAN S-phase kinase-associated protein 1 OS=Homo sapiens OX=9606 GN=SKP1 PE=1 SV=2 | 26.27 | 15.85 | 0.60 | -0.73 |
| sp\|P42677\|RS27_HUMAN 40S ribosomal protein S27 OS=Homo sapiens OX=9606 GN=RPS27 PE=1 SV=3 | 26.12 | 39.53 | 1.51 | 0.60 |
| sp\|O00487\|PSDE_HUMAN 26S proteasome non-ATPase regulatory subunit 14 OS=Homo sapiens OX=9606 GN=PSMD14 PE=1 SV=1 | 26.01 | 66.25 | 2.55 | 1.35 |
| sp\|P30046\|DOPD_HUMAN D-dopachrome decarboxylase OS=Homo sapiens OX=9606 GN=DDT PE=1 SV=3 | 25.99 | 132.95 | 5.11 | 2.35 |
| sp\|P30084\|ECHM_HUMAN Enoyl-CoA hydratase, mitochondrial OS=Homo sapiens OX=9606 GN=ECHS1 PE=1 SV=4 | 25.83 | 78.03 | 3.02 | 1.60 |
| sp\|Q15019-2\|SEPT2_HUMAN Isoform 2 of Septin-2 OS=Homo sapiens OX=9606 GN=SEPT2 | 25.65 | 45.03 | 1.76 | 0.81 |
| sp\|P13489\|RINI_HUMAN Ribonuclease inhibitor OS=Homo sapiens OX=9606 GN=RNH1 PE=1 SV=2 | 25.52 | 50.18 | 1.97 | 0.98 |
| sp\|P23246\|SFPQ_HUMAN Splicing factor, proline- and glutamine-rich OS=Homo sapiens OX=9606 GN=SFPQ PE=1 SV=2 | 25.45 | 19.57 | 0.77 | -0.38 |
| tr\|Q8WVC2\|Q8WVC2_HUMAN 40S ribosomal protein S21 OS=Homo sapiens OX=9606 GN=RPS21 PE=1 SV=1 | 25.44 | 74.02 | 2.91 | 1.54 |
| sp\|Q8NBS9-2\|TXND5_HUMAN Isoform 2 of Thioredoxin domain-containing protein 5 OS=Homo sapiens OX=9606 GN=TXNDC5 | 25.37 | 36.79 | 1.45 | 0.54 |
| sp\|P30153\|2AAA_HUMAN Serine/threonine-protein phosphatase 2A 65 kDa regulatory | 25.20 | 50.64 | 2.01 | 1.01 |

| | | | | |
|---|---|---|---|---|
| subunit A alpha isoform OS=Homo sapiens OX=9606 GN=PPP2R1A PE=1 SV=4 | | | | |
| sp\|P82979\|SARNP_HUMAN SAP domain-containing ribonucleoprotein OS=Homo sapiens OX=9606 GN=SARNP PE=1 SV=3 | 25.08 | 55.84 | 2.23 | 1.15 |
| sp\|O76021\|RL1D1_HUMAN Ribosomal L1 domain-containing protein 1 OS=Homo sapiens OX=9606 GN=RSL1D1 PE=1 SV=3 | 24.71 | 39.18 | 1.59 | 0.66 |
| tr\|Q6P452\|Q6P452_HUMAN Annexin OS=Homo sapiens OX=9606 GN=ANXA4 PE=1 SV=1 | 24.60 | 24.91 | 1.01 | 0.02 |
| tr\|F8VV56\|F8VV56_HUMAN CD63 antigen OS=Homo sapiens OX=9606 GN=CD63 PE=1 SV=1 | 24.49 | 34.28 | 1.40 | 0.48 |
| sp\|Q92688-2\|AN32B_HUMAN Isoform 2 of Acidic leucine-rich nuclear phosphoprotein 32 family member B OS=Homo sapiens OX=9606 GN=ANP32B | 24.40 | 113.69 | 4.66 | 2.22 |
| sp\|P0DP23\|CALM1_HUMAN Calmodulin-1 OS=Homo sapiens OX=9606 GN=CALM1 PE=1 SV=1 | 24.31 | 155.12 | 6.38 | 2.67 |
| sp\|P49755\|TMEDA_HUMAN Transmembrane emp24 domain-containing protein 10 OS=Homo sapiens OX=9606 GN=TMED10 PE=1 SV=2 | 24.30 | 80.61 | 3.32 | 1.73 |
| sp\|Q8NC51\|PAIRB_HUMAN Plasminogen activator inhibitor 1 RNA-binding protein OS=Homo sapiens OX=9606 GN=SERBP1 PE=1 SV=2 | 24.29 | 25.52 | 1.05 | 0.07 |
| sp\|Q9P0S9\|TM14C_HUMAN Transmembrane protein 14C OS=Homo sapiens OX=9606 GN=TMEM14C PE=1 SV=1 | 24.27 | 41.06 | 1.69 | 0.76 |
| sp\|P16435\|NCPR_HUMAN NADPH--cytochrome P450 reductase OS=Homo sapiens OX=9606 GN=POR PE=1 SV=2 | 24.20 | 60.99 | 2.52 | 1.33 |
| sp\|P16152\|CBR1_HUMAN Carbonyl reductase [NADPH] 1 OS=Homo sapiens OX=9606 GN=CBR1 PE=1 SV=3 | 24.09 | 47.40 | 1.97 | 0.98 |
| tr\|J3QT28\|J3QT28_HUMAN Mitotic checkpoint protein BUB3 (Fragment) OS=Homo sapiens OX=9606 GN=BUB3 PE=1 SV=1 | 23.97 | 28.52 | 1.19 | 0.25 |
| sp\|Q7L2H7\|EIF3M_HUMAN Eukaryotic translation initiation factor 3 subunit M OS=Homo sapiens OX=9606 GN=EIF3M PE=1 SV=1 | 23.97 | 64.71 | 2.70 | 1.43 |
| sp\|P27695\|APEX1_HUMAN DNA-(apurinic or apyrimidinic site) lyase OS=Homo sapiens OX=9606 GN=APEX1 PE=1 SV=2 | 23.97 | 50.08 | 2.09 | 1.06 |
| sp\|P00492\|HPRT_HUMAN Hypoxanthine-guanine phosphoribosyltransferase OS=Homo sapiens OX=9606 GN=HPRT1 PE=1 SV=2 | 23.90 | 15.56 | 0.65 | -0.62 |
| sp\|Q15717-2\|ELAV1_HUMAN Isoform 2 of ELAV-like protein 1 OS=Homo sapiens OX=9606 GN=ELAVL1 | 23.74 | 42.50 | 1.79 | 0.84 |

| | | | | |
|---|---|---|---|---|
| tr\|A0A0C4DGQ5\|A0A0C4DGQ5_HUMAN Calpain small subunit 1 OS=Homo sapiens OX=9606 GN=CAPNS1 PE=1 SV=1 | 23.74 | 64.50 | 2.72 | 1.44 |
| sp\|P61158\|ARP3_HUMAN Actin-related protein 3 OS=Homo sapiens OX=9606 GN=ACTR3 PE=1 SV=3 | 23.65 | 57.68 | 2.44 | 1.29 |
| sp\|P12429\|ANXA3_HUMAN Annexin A3 OS=Homo sapiens OX=9606 GN=ANXA3 PE=1 SV=3 | 23.58 | 23.52 | 1.00 | 0.00 |
| sp\|P33993\|MCM7_HUMAN DNA replication licensing factor MCM7 OS=Homo sapiens OX=9606 GN=MCM7 PE=1 SV=4 | 23.37 | 67.02 | 2.87 | 1.52 |
| sp\|P25788-2\|PSA3_HUMAN Isoform 2 of Proteasome subunit alpha type-3 OS=Homo sapiens OX=9606 GN=PSMA3 | 23.33 | 20.23 | 0.87 | -0.21 |
| tr\|A0A087X1Z3\|A0A087X1Z3_HUMAN Proteasome activator complex subunit 2 OS=Homo sapiens OX=9606 GN=PSME2 PE=1 SV=1 | 23.23 | 63.80 | 2.75 | 1.46 |
| tr\|A0A0C4DGS1\|A0A0C4DGS1_HUMAN Dolichyl-diphosphooligosaccharide--protein glycosyltransferase 48 kDa subunit OS=Homo sapiens OX=9606 GN=DDOST PE=1 SV=1 | 23.17 | 36.94 | 1.59 | 0.67 |
| sp\|P30154-2\|2AAB_HUMAN Isoform 2 of Serine/threonine-protein phosphatase 2A 65 kDa regulatory subunit A beta isoform OS=Homo sapiens OX=9606 GN=PPP2R1B | 23.16 | 3.47 | 0.15 | -2.74 |
| sp\|P20042\|IF2B_HUMAN Eukaryotic translation initiation factor 2 subunit 2 OS=Homo sapiens OX=9606 GN=EIF2S2 PE=1 SV=2 | 23.06 | 52.48 | 2.28 | 1.19 |
| sp\|P15170-2\|ERF3A_HUMAN Isoform 2 of Eukaryotic peptide chain release factor GTP-binding subunit ERF3A OS=Homo sapiens OX=9606 GN=GSPT1 | 23.06 | 33.17 | 1.44 | 0.52 |
| sp\|Q9HBL8\|NMRL1_HUMAN NmrA-like family domain-containing protein 1 OS=Homo sapiens OX=9606 GN=NMRAL1 PE=1 SV=1 | 23.02 | 30.18 | 1.31 | 0.39 |
| sp\|P20674\|COX5A_HUMAN Cytochrome c oxidase subunit 5A, mitochondrial OS=Homo sapiens OX=9606 GN=COX5A PE=1 SV=2 | 22.97 | 28.21 | 1.23 | 0.30 |
| sp\|P14324-2\|FPPS_HUMAN Isoform 2 of Farnesyl pyrophosphate synthase OS=Homo sapiens OX=9606 GN=FDPS | 22.96 | 73.69 | 3.21 | 1.68 |
| sp\|O60488-2\|ACSL4_HUMAN Isoform Short of Long-chain-fatty-acid--CoA ligase 4 OS=Homo sapiens OX=9606 GN=ACSL4 | 22.94 | 12.14 | 0.53 | -0.92 |
| tr\|G3V1V0\|G3V1V0_HUMAN Myosin light polypeptide 6 OS=Homo sapiens OX=9606 GN=MYL6 PE=1 SV=1 | 22.92 | 151.77 | 6.62 | 2.73 |
| sp\|Q9NQR4\|NIT2_HUMAN Omega-amidase NIT2 OS=Homo sapiens OX=9606 GN=NIT2 PE=1 SV=1 | 22.82 | 12.88 | 0.56 | -0.83 |

| | | | | |
|---|---|---|---|---|
| sp\|O95831\|AIFM1_HUMAN Apoptosis-inducing factor 1, mitochondrial OS=Homo sapiens OX=9606 GN=AIFM1 PE=1 SV=1 | 22.79 | 39.35 | 1.73 | 0.79 |
| sp\|P08133\|ANXA6_HUMAN Annexin A6 OS=Homo sapiens OX=9606 GN=ANXA6 PE=1 SV=3 | 22.58 | 42.16 | 1.87 | 0.90 |
| sp\|P62191\|PRS4_HUMAN 26S proteasome regulatory subunit 4 OS=Homo sapiens OX=9606 GN=PSMC1 PE=1 SV=1 | 22.42 | 40.81 | 1.82 | 0.86 |
| sp\|Q9Y266\|NUDC_HUMAN Nuclear migration protein nudC OS=Homo sapiens OX=9606 GN=NUDC PE=1 SV=1 | 22.23 | 51.89 | 2.33 | 1.22 |
| sp\|Q13011\|ECH1_HUMAN Delta(3,5)-Delta(2,4)-dienoyl-CoA isomerase, mitochondrial OS=Homo sapiens OX=9606 GN=ECH1 PE=1 SV=2 | 22.13 | 44.54 | 2.01 | 1.01 |
| sp\|P62195\|PRS8_HUMAN 26S proteasome regulatory subunit 8 OS=Homo sapiens OX=9606 GN=PSMC5 PE=1 SV=1 | 21.94 | 43.76 | 1.99 | 1.00 |
| sp\|Q15758\|AAAT_HUMAN Neutral amino acid transporter B(0) OS=Homo sapiens OX=9606 GN=SLC1A5 PE=1 SV=2 | 21.94 | 45.90 | 2.09 | 1.07 |
| sp\|Q16891-2\|MIC60_HUMAN Isoform 2 of MICOS complex subunit MIC60 OS=Homo sapiens OX=9606 GN=IMMT | 21.92 | 48.89 | 2.23 | 1.16 |
| sp\|P26583\|HMGB2_HUMAN High mobility group protein B2 OS=Homo sapiens OX=9606 GN=HMGB2 PE=1 SV=2 | 21.84 | 66.09 | 3.03 | 1.60 |
| sp\|P62316\|SMD2_HUMAN Small nuclear ribonucleoprotein Sm D2 OS=Homo sapiens OX=9606 GN=SNRPD2 PE=1 SV=1 | 21.81 | 36.90 | 1.69 | 0.76 |
| sp\|Q96PK6\|RBM14_HUMAN RNA-binding protein 14 OS=Homo sapiens OX=9606 GN=RBM14 PE=1 SV=2 | 21.74 | 114.71 | 5.28 | 2.40 |
| sp\|Q96IX5\|USMG5_HUMAN Up-regulated during skeletal muscle growth protein 5 OS=Homo sapiens OX=9606 GN=ATP5MD PE=1 SV=1 | 21.73 | 55.27 | 2.54 | 1.35 |
| tr\|F5GX77\|F5GX77_HUMAN Multifunctional methyltransferase subunit TRM112-like protein OS=Homo sapiens OX=9606 GN=TRMT112 PE=1 SV=1 | 21.46 | 47.26 | 2.20 | 1.14 |
| sp\|Q15691\|MARE1_HUMAN Microtubule-associated protein RP/EB family member 1 OS=Homo sapiens OX=9606 GN=MAPRE1 PE=1 SV=3 | 21.16 | 12.10 | 0.57 | -0.81 |
| sp\|Q06210-2\|GFPT1_HUMAN Isoform 2 of Glutamine--fructose-6-phosphate aminotransferase [isomerizing] 1 OS=Homo sapiens OX=9606 GN=GFPT1 | 21.13 | 33.40 | 1.58 | 0.66 |
| sp\|P18754-2\|RCC1_HUMAN Isoform 2 of Regulator of chromosome condensation OS=Homo sapiens OX=9606 GN=RCC1 | 21.11 | 40.60 | 1.92 | 0.94 |

| | | | | |
|---|---|---|---|---|
| tr\|H3BN98\|H3BN98_HUMAN Uncharacterized protein (Fragment) OS=Homo sapiens OX=9606 PE=4 SV=2 | 21.01 | 69.36 | 3.30 | 1.72 |
| tr\|B4E0Y9\|B4E0Y9_HUMAN Serine/threonine-protein kinase 26 OS=Homo sapiens OX=9606 GN=STK26 PE=1 SV=1 | 20.92 | 27.44 | 1.31 | 0.39 |
| sp\|P51452\|DUS3_HUMAN Dual specificity protein phosphatase 3 OS=Homo sapiens OX=9606 GN=DUSP3 PE=1 SV=1 | 20.91 | 37.91 | 1.81 | 0.86 |
| sp\|P43897\|EFTS_HUMAN Elongation factor Ts, mitochondrial OS=Homo sapiens OX=9606 GN=TSFM PE=1 SV=2 | 20.82 | 31.21 | 1.50 | 0.58 |
| sp\|Q07666-2\|KHDR1_HUMAN Isoform 2 of KH domain-containing, RNA-binding, signal transduction-associated protein 1 OS=Homo sapiens OX=9606 GN=KHDRBS1 | 20.82 | 20.55 | 0.99 | -0.02 |
| sp\|P54136\|SYRC_HUMAN Arginine--tRNA ligase, cytoplasmic OS=Homo sapiens OX=9606 GN=RARS PE=1 SV=2 | 20.80 | 58.09 | 2.79 | 1.48 |
| sp\|Q9BY32-2\|ITPA_HUMAN Isoform 2 of Inosine triphosphate pyrophosphatase OS=Homo sapiens OX=9606 GN=ITPA | 20.71 | 4.74 | 0.23 | -2.13 |
| sp\|Q01813\|PFKAP_HUMAN ATP-dependent 6-phosphofructokinase, platelet type OS=Homo sapiens OX=9606 GN=PFKP PE=1 SV=2 | 20.69 | 44.96 | 2.17 | 1.12 |
| tr\|J3QRS3\|J3QRS3_HUMAN Myosin regulatory light chain 12A OS=Homo sapiens OX=9606 GN=MYL12A PE=1 SV=1 | 20.61 | 84.18 | 4.08 | 2.03 |
| sp\|Q9NR45\|SIAS_HUMAN Sialic acid synthase OS=Homo sapiens OX=9606 GN=NANS PE=1 SV=2 | 20.59 | 44.09 | 2.14 | 1.10 |
| sp\|Q9Y696\|CLIC4_HUMAN Chloride intracellular channel protein 4 OS=Homo sapiens OX=9606 GN=CLIC4 PE=1 SV=4 | 20.54 | 15.02 | 0.73 | -0.45 |
| tr\|H0YKC5\|H0YKC5_HUMAN Deoxyuridine 5'-triphosphate nucleotidohydrolase, mitochondrial (Fragment) OS=Homo sapiens OX=9606 GN=DUT PE=1 SV=1 | 20.49 | 53.05 | 2.59 | 1.37 |
| sp\|P01891\|1A68_HUMAN HLA class I histocompatibility antigen, A-68 alpha chain OS=Homo sapiens OX=9606 GN=HLA-A PE=1 SV=4 | 20.45 | 34.51 | 1.69 | 0.75 |
| sp\|Q8WUM4\|PDC6I_HUMAN Programmed cell death 6-interacting protein OS=Homo sapiens OX=9606 GN=PDCD6IP PE=1 SV=1 | 20.22 | 47.00 | 2.32 | 1.22 |
| sp\|O75607\|NPM3_HUMAN Nucleoplasmin-3 OS=Homo sapiens OX=9606 GN=NPM3 PE=1 SV=3 | 20.19 | 74.16 | 3.67 | 1.88 |
| sp\|O00264\|PGRC1_HUMAN Membrane-associated progesterone receptor component 1 OS=Homo sapiens OX=9606 GN=PGRMC1 PE=1 SV=3 | 20.02 | 39.80 | 1.99 | 0.99 |

| | | | | |
|---|---|---|---|---|
| sp\|P40121-2\|CAPG_HUMAN Isoform 2 of Macrophage-capping protein OS=Homo sapiens OX=9606 GN=CAPG | 19.99 | 13.54 | 0.68 | -0.56 |
| tr\|G3V153\|G3V153_HUMAN Caprin-1 OS=Homo sapiens OX=9606 GN=CAPRIN1 PE=1 SV=1 | 19.99 | 32.49 | 1.63 | 0.70 |
| sp\|P49773\|HINT1_HUMAN Histidine triad nucleotide-binding protein 1 OS=Homo sapiens OX=9606 GN=HINT1 PE=1 SV=2 | 19.89 | 20.90 | 1.05 | 0.07 |
| sp\|P35998\|PRS7_HUMAN 26S proteasome regulatory subunit 7 OS=Homo sapiens OX=9606 GN=PSMC2 PE=1 SV=3 | 19.85 | 40.92 | 2.06 | 1.04 |
| sp\|Q9Y333\|LSM2_HUMAN U6 snRNA-associated Sm-like protein LSm2 OS=Homo sapiens OX=9606 GN=LSM2 PE=1 SV=1 | 19.84 | 38.46 | 1.94 | 0.95 |
| sp\|P13073\|COX41_HUMAN Cytochrome c oxidase subunit 4 isoform 1, mitochondrial OS=Homo sapiens OX=9606 GN=COX4I1 PE=1 SV=1 | 19.77 | 18.80 | 0.95 | -0.07 |
| sp\|Q9UMS4\|PRP19_HUMAN Pre-mRNA-processing factor 19 OS=Homo sapiens OX=9606 GN=PRPF19 PE=1 SV=1 | 19.48 | 45.66 | 2.34 | 1.23 |
| tr\|A0A2R8YD50\|A0A2R8YD50_HUMAN Peroxisomal multifunctional enzyme type 2 OS=Homo sapiens OX=9606 GN=HSD17B4 PE=1 SV=1 | 19.48 | 36.94 | 1.90 | 0.92 |
| tr\|Q5JRI1\|Q5JRI1_HUMAN Serine/arginine-rich-splicing factor 10 OS=Homo sapiens OX=9606 GN=SRSF10 PE=1 SV=1 | 19.33 | 40.92 | 2.12 | 1.08 |
| sp\|Q15046-2\|SYK_HUMAN Isoform Mitochondrial of Lysine--tRNA ligase OS=Homo sapiens OX=9606 GN=KARS | 19.27 | 62.58 | 3.25 | 1.70 |
| tr\|F8VVA7\|F8VVA7_HUMAN Coatomer subunit zeta-1 OS=Homo sapiens OX=9606 GN=COPZ1 PE=1 SV=1 | 19.22 | 27.87 | 1.45 | 0.54 |
| sp\|P00390-3\|GSHR_HUMAN Isoform 2 of Glutathione reductase, mitochondrial OS=Homo sapiens OX=9606 GN=GSR | 19.20 | 29.81 | 1.55 | 0.63 |
| sp\|O43776\|SYNC_HUMAN Asparagine--tRNA ligase, cytoplasmic OS=Homo sapiens OX=9606 GN=NARS PE=1 SV=1 | 19.09 | 70.30 | 3.68 | 1.88 |
| sp\|P17174\|AATC_HUMAN Aspartate aminotransferase, cytoplasmic OS=Homo sapiens OX=9606 GN=GOT1 PE=1 SV=3 | 18.94 | 43.12 | 2.28 | 1.19 |
| tr\|F8VVM2\|F8VVM2_HUMAN Phosphate carrier protein, mitochondrial OS=Homo sapiens OX=9606 GN=SLC25A3 PE=1 SV=1 | 18.89 | 27.44 | 1.45 | 0.54 |
| sp\|P22102-2\|PUR2_HUMAN Isoform Short of Trifunctional purine biosynthetic protein adenosine-3 OS=Homo sapiens OX=9606 GN=GART | 18.85 | 27.11 | 1.44 | 0.52 |
| tr\|E9PK05\|E9PK05_HUMAN Phospholipid transfer protein C2CD2L OS=Homo sapiens OX=9606 GN=C2CD2L PE=1 SV=1 | 18.71 | 10.83 | 0.58 | -0.79 |

| | | | | |
|---|---|---|---|---|
| sp\|O43852-3\|CALU_HUMAN Isoform 3 of Calumenin OS=Homo sapiens OX=9606 GN=CALU | 18.70 | 19.28 | 1.03 | 0.04 |
| sp\|Q8NCW5-2\|NNRE_HUMAN Isoform 2 of NAD(P)H-hydrate epimerase OS=Homo sapiens OX=9606 GN=NAXE | 18.54 | 48.25 | 2.60 | 1.38 |
| sp\|O75874\|IDHC_HUMAN Isocitrate dehydrogenase [NADP] cytoplasmic OS=Homo sapiens OX=9606 GN=IDH1 PE=1 SV=2 | 18.51 | 45.87 | 2.48 | 1.31 |
| sp\|P35659\|DEK_HUMAN Protein DEK OS=Homo sapiens OX=9606 GN=DEK PE=1 SV=1 | 18.48 | 42.06 | 2.28 | 1.19 |
| sp\|O75083\|WDR1_HUMAN WD repeat-containing protein 1 OS=Homo sapiens OX=9606 GN=WDR1 PE=1 SV=4 | 18.47 | 20.34 | 1.10 | 0.14 |
| sp\|P61019\|RAB2A_HUMAN Ras-related protein Rab-2A OS=Homo sapiens OX=9606 GN=RAB2A PE=1 SV=1 | 18.26 | 45.62 | 2.50 | 1.32 |
| sp\|Q15738\|NSDHL_HUMAN Sterol-4-alpha-carboxylate 3-dehydrogenase, decarboxylating OS=Homo sapiens OX=9606 GN=NSDHL PE=1 SV=2 | 18.19 | 15.55 | 0.85 | -0.23 |
| sp\|P00491\|PNPH_HUMAN Purine nucleoside phosphorylase OS=Homo sapiens OX=9606 GN=PNP PE=1 SV=2 | 18.05 | 48.09 | 2.66 | 1.41 |
| sp\|P09661\|RU2A_HUMAN U2 small nuclear ribonucleoprotein A' OS=Homo sapiens OX=9606 GN=SNRPA1 PE=1 SV=2 | 18.05 | 43.34 | 2.40 | 1.26 |
| tr\|B8ZZQ6\|B8ZZQ6_HUMAN Prothymosin alpha OS=Homo sapiens OX=9606 GN=PTMA PE=1 SV=1 | 17.99 | 33.57 | 1.87 | 0.90 |
| sp\|Q13057-2\|COASY_HUMAN Isoform 2 of Bifunctional coenzyme A synthase OS=Homo sapiens OX=9606 GN=COASY | 17.86 | 26.24 | 1.47 | 0.55 |
| sp\|Q9Y678\|COPG1_HUMAN Coatomer subunit gamma-1 OS=Homo sapiens OX=9606 GN=COPG1 PE=1 SV=1 | 17.81 | 3.34 | 0.19 | -2.42 |
| sp\|Q12904-2\|AIMP1_HUMAN Isoform 2 of Aminoacyl tRNA synthase complex-interacting multifunctional protein 1 OS=Homo sapiens OX=9606 GN=AIMP1 | 17.77 | 37.31 | 2.10 | 1.07 |
| sp\|P68402\|PA1B2_HUMAN Platelet-activating factor acetylhydrolase IB subunit beta OS=Homo sapiens OX=9606 GN=PAFAH1B2 PE=1 SV=1 | 17.73 | 37.79 | 2.13 | 1.09 |
| tr\|B1AK87\|B1AK87_HUMAN Capping protein (Actin filament) muscle Z-line, beta, isoform CRA_a OS=Homo sapiens OX=9606 GN=CAPZB PE=1 SV=1 | 17.70 | 25.18 | 1.42 | 0.51 |
| sp\|Q9BVK6\|TMED9_HUMAN Transmembrane emp24 domain-containing protein 9 OS=Homo sapiens OX=9606 GN=TMED9 PE=1 SV=2 | 17.70 | 18.38 | 1.04 | 0.05 |

| | | | | |
|---|---|---|---|---|
| sp\|P09012\|SNRPA_HUMAN U1 small nuclear ribonucleoprotein A OS=Homo sapiens OX=9606 GN=SNRPA PE=1 SV=3 | 17.69 | 50.20 | 2.84 | 1.51 |
| sp\|P49915\|GUAA_HUMAN GMP synthase [glutamine-hydrolyzing] OS=Homo sapiens OX=9606 GN=GMPS PE=1 SV=1 | 17.62 | 41.27 | 2.34 | 1.23 |
| sp\|P04040\|CATA_HUMAN Catalase OS=Homo sapiens OX=9606 GN=CAT PE=1 SV=3 | 17.60 | 32.76 | 1.86 | 0.90 |
| sp\|Q6NZI2\|CAVN1_HUMAN Caveolae-associated protein 1 OS=Homo sapiens OX=9606 GN=CAVIN1 PE=1 SV=1 | 17.59 | 30.15 | 1.71 | 0.78 |
| sp\|Q9BQ67\|GRWD1_HUMAN Glutamate-rich WD repeat-containing protein 1 OS=Homo sapiens OX=9606 GN=GRWD1 PE=1 SV=1 | 17.38 | 24.04 | 1.38 | 0.47 |
| sp\|P62714\|PP2AB_HUMAN Serine/threonine-protein phosphatase 2A catalytic subunit beta isoform OS=Homo sapiens OX=9606 GN=PPP2CB PE=1 SV=1 | 17.36 | 108.40 | 6.24 | 2.64 |
| sp\|Q9UBF2-2\|COPG2_HUMAN Isoform 2 of Coatomer subunit gamma-2 OS=Homo sapiens OX=9606 GN=COPG2 | 17.31 | 11.36 | 0.66 | -0.61 |
| tr\|F8WCF6\|F8WCF6_HUMAN Actin-related protein 2/3 complex subunit 4 OS=Homo sapiens OX=9606 GN=ARPC4-TTLL3 PE=3 SV=1 | 17.24 | 40.66 | 2.36 | 1.24 |
| tr\|A0A087WUK2\|A0A087WUK2_HUMAN Heterogeneous nuclear ribonucleoprotein D-like, isoform CRA_b OS=Homo sapiens OX=9606 GN=HNRNPDL PE=1 SV=1 | 17.23 | 102.90 | 5.97 | 2.58 |
| sp\|Q92499\|DDX1_HUMAN ATP-dependent RNA helicase DDX1 OS=Homo sapiens OX=9606 GN=DDX1 PE=1 SV=2 | 17.21 | 29.32 | 1.70 | 0.77 |
| sp\|O15427\|MOT4_HUMAN Monocarboxylate transporter 4 OS=Homo sapiens OX=9606 GN=SLC16A3 PE=1 SV=1 | 17.17 | 10.46 | 0.61 | -0.71 |
| tr\|E7ETZ4\|E7ETZ4_HUMAN Basic leucine zipper and W2 domain-containing protein 2 (Fragment) OS=Homo sapiens OX=9606 GN=BZW2 PE=1 SV=1 | 17.11 | 58.12 | 3.40 | 1.76 |
| tr\|A0A087WZT3\|A0A087WZT3_HUMAN BolA-like protein 2 OS=Homo sapiens OX=9606 GN=BOLA2 PE=1 SV=2 | 17.04 | 46.56 | 2.73 | 1.45 |
| sp\|P05198\|IF2A_HUMAN Eukaryotic translation initiation factor 2 subunit 1 OS=Homo sapiens OX=9606 GN=EIF2S1 PE=1 SV=3 | 17.00 | 37.25 | 2.19 | 1.13 |
| sp\|P30085\|KCY_HUMAN UMP-CMP kinase OS=Homo sapiens OX=9606 GN=CMPK1 PE=1 SV=3 | 16.98 | 47.34 | 2.79 | 1.48 |
| sp\|Q96DG6\|CMBL_HUMAN Carboxymethylenebutenolidase homolog OS=Homo sapiens OX=9606 GN=CMBL PE=1 SV=1 | 16.93 | 24.35 | 1.44 | 0.52 |

| | | | | |
|---|---|---|---|---|
| sp\|P10606\|COX5B_HUMAN Cytochrome c oxidase subunit 5B, mitochondrial OS=Homo sapiens OX=9606 GN=COX5B PE=1 SV=2 | 16.91 | 18.96 | 1.12 | 0.17 |
| sp\|P47985\|UCRI_HUMAN Cytochrome b-c1 complex subunit Rieske, mitochondrial OS=Homo sapiens OX=9606 GN=UQCRFS1 PE=1 SV=2 | 16.90 | 15.20 | 0.90 | -0.15 |
| tr\|E7EMS6\|E7EMS6_HUMAN Catechol O-methyltransferase (Fragment) OS=Homo sapiens OX=9606 GN=COMT PE=1 SV=1 | 16.89 | 34.15 | 2.02 | 1.02 |
| sp\|Q12906-2\|ILF3_HUMAN Isoform 2 of Interleukin enhancer-binding factor 3 OS=Homo sapiens OX=9606 GN=ILF3 | 16.75 | 63.78 | 3.81 | 1.93 |
| sp\|P30043\|BLVRB_HUMAN Flavin reductase (NADPH) OS=Homo sapiens OX=9606 GN=BLVRB PE=1 SV=3 | 16.74 | 39.53 | 2.36 | 1.24 |
| sp\|Q14498-2\|RBM39_HUMAN Isoform 2 of RNA-binding protein 39 OS=Homo sapiens OX=9606 GN=RBM39 | 16.71 | 16.95 | 1.01 | 0.02 |
| sp\|P26196\|DDX6_HUMAN Probable ATP-dependent RNA helicase DDX6 OS=Homo sapiens OX=9606 GN=DDX6 PE=1 SV=2 | 16.70 | 30.20 | 1.81 | 0.85 |
| sp\|P10620\|MGST1_HUMAN Microsomal glutathione S-transferase 1 OS=Homo sapiens OX=9606 GN=MGST1 PE=1 SV=1 | 16.69 | 68.55 | 4.11 | 2.04 |
| tr\|A0A1B0GVA9\|A0A1B0GVA9_HUMAN Ryanodine receptor 3 (Fragment) OS=Homo sapiens OX=9606 GN=RYR3 PE=4 SV=1 | 16.53 | 6.45 | 0.39 | -1.36 |
| sp\|P61970\|NTF2_HUMAN Nuclear transport factor 2 OS=Homo sapiens OX=9606 GN=NUTF2 PE=1 SV=1 | 16.50 | 53.54 | 3.24 | 1.70 |
| tr\|B4E3S0\|B4E3S0_HUMAN Coronin OS=Homo sapiens OX=9606 GN=CORO1C PE=1 SV=1 | 16.47 | 5.15 | 0.31 | -1.68 |
| sp\|P08579\|RU2B_HUMAN U2 small nuclear ribonucleoprotein B'' OS=Homo sapiens OX=9606 GN=SNRPB2 PE=1 SV=1 | 16.44 | 13.23 | 0.80 | -0.31 |
| sp\|P51572-2\|BAP31_HUMAN Isoform 2 of B-cell receptor-associated protein 31 OS=Homo sapiens OX=9606 GN=BCAP31 | 16.41 | 43.87 | 2.67 | 1.42 |
| sp\|P54709\|AT1B3_HUMAN Sodium/potassium-transporting ATPase subunit beta-3 OS=Homo sapiens OX=9606 GN=ATP1B3 PE=1 SV=1 | 16.37 | 36.82 | 2.25 | 1.17 |
| sp\|P36957\|ODO2_HUMAN Dihydrolipoyllysine-residue succinyltransferase component of 2-oxoglutarate dehydrogenase complex, mitochondrial OS=Homo sapiens OX=9606 GN=DLST PE=1 SV=4 | 16.36 | 32.19 | 1.97 | 0.98 |
| sp\|O15371-2\|EIF3D_HUMAN Isoform 2 of Eukaryotic translation initiation factor 3 subunit D OS=Homo sapiens OX=9606 GN=EIF3D | 16.34 | 47.97 | 2.94 | 1.55 |

| | | | | |
|---|---|---|---|---|
| sp\|Q9UBQ5-2\|EIF3K_HUMAN Isoform 2 of Eukaryotic translation initiation factor 3 subunit K OS=Homo sapiens OX=9606 GN=EIF3K | 16.32 | 82.48 | 5.05 | 2.34 |
| tr\|A0A0B4J2C3\|A0A0B4J2C3_HUMAN Translationally-controlled tumor protein OS=Homo sapiens OX=9606 GN=TPT1 PE=1 SV=1 | 16.28 | 32.74 | 2.01 | 1.01 |
| sp\|P84090\|ERH_HUMAN Enhancer of rudimentary homolog OS=Homo sapiens OX=9606 GN=ERH PE=1 SV=1 | 16.25 | 63.07 | 3.88 | 1.96 |
| sp\|P51571\|SSRD_HUMAN Translocon-associated protein subunit delta OS=Homo sapiens OX=9606 GN=SSR4 PE=1 SV=1 | 16.24 | 70.17 | 4.32 | 2.11 |
| tr\|A0A0C4DFL7\|A0A0C4DFL7_HUMAN Lanosterol 14-alpha demethylase OS=Homo sapiens OX=9606 GN=CYP51A1 PE=1 SV=1 | 16.21 | 4.76 | 0.29 | -1.77 |
| sp\|P14923\|PLAK_HUMAN Junction plakoglobin OS=Homo sapiens OX=9606 GN=JUP PE=1 SV=3 | 16.07 | 10.90 | 0.68 | -0.56 |
| sp\|Q14847-2\|LASP1_HUMAN Isoform 2 of LIM and SH3 domain protein 1 OS=Homo sapiens OX=9606 GN=LASP1 | 16.01 | 17.89 | 1.12 | 0.16 |
| sp\|O60684\|IMA7_HUMAN Importin subunit alpha-7 OS=Homo sapiens OX=9606 GN=KPNA6 PE=1 SV=1 | 16.00 | 31.25 | 1.95 | 0.97 |
| sp\|O95816-2\|BAG2_HUMAN Isoform 2 of BAG family molecular chaperone regulator 2 OS=Homo sapiens OX=9606 GN=BAG2 | 15.93 | 10.85 | 0.68 | -0.55 |
| sp\|Q92979\|NEP1_HUMAN Ribosomal RNA small subunit methyltransferase NEP1 OS=Homo sapiens OX=9606 GN=EMG1 PE=1 SV=4 | 15.83 | 43.63 | 2.76 | 1.46 |
| sp\|Q9BS26\|ERP44_HUMAN Endoplasmic reticulum resident protein 44 OS=Homo sapiens OX=9606 GN=ERP44 PE=1 SV=1 | 15.72 | 27.28 | 1.73 | 0.79 |
| tr\|G5EA30\|G5EA30_HUMAN CUG triplet repeat, RNA binding protein 1, isoform CRA_c OS=Homo sapiens OX=9606 GN=CELF1 PE=1 SV=1 | 15.70 | 24.67 | 1.57 | 0.65 |
| sp\|O76003\|GLRX3_HUMAN Glutaredoxin-3 OS=Homo sapiens OX=9606 GN=GLRX3 PE=1 SV=2 | 15.63 | 89.33 | 5.72 | 2.51 |
| sp\|O76070\|SYUG_HUMAN Gamma-synuclein OS=Homo sapiens OX=9606 GN=SNCG PE=1 SV=2 | 15.57 | 61.41 | 3.94 | 1.98 |
| tr\|A0A0D9SFS3\|A0A0D9SFS3_HUMAN 2-oxoglutarate dehydrogenase, mitochondrial OS=Homo sapiens OX=9606 GN=OGDH PE=1 SV=1 | 15.47 | 69.67 | 4.50 | 2.17 |
| sp\|P20700\|LMNB1_HUMAN Lamin-B1 OS=Homo sapiens OX=9606 GN=LMNB1 PE=1 SV=2 | 15.36 | 27.43 | 1.79 | 0.84 |

| | | | | |
|---|---|---|---|---|
| sp\|Q9BPW8\|NIPS1_HUMAN Protein NipSnap homolog 1 OS=Homo sapiens OX=9606 GN=NIPSNAP1 PE=1 SV=1 | 15.33 | 2.34 | 0.15 | -2.72 |
| sp\|O00232\|PSD12_HUMAN 26S proteasome non-ATPase regulatory subunit 12 OS=Homo sapiens OX=9606 GN=PSMD12 PE=1 SV=3 | 15.26 | 53.34 | 3.49 | 1.81 |
| sp\|A6NDG6\|PGP_HUMAN Glycerol-3-phosphate phosphatase OS=Homo sapiens OX=9606 GN=PGP PE=1 SV=1 | 15.19 | 49.75 | 3.28 | 1.71 |
| sp\|Q7RTV0\|PHF5A_HUMAN PHD finger-like domain-containing protein 5A OS=Homo sapiens OX=9606 GN=PHF5A PE=1 SV=1 | 15.18 | 10.15 | 0.67 | -0.58 |
| sp\|P30520\|PURA2_HUMAN Adenylosuccinate synthetase isozyme 2 OS=Homo sapiens OX=9606 GN=ADSS PE=1 SV=3 | 15.15 | 30.23 | 2.00 | 1.00 |
| tr\|I3L0A0\|I3L0A0_HUMAN HCG2044781 OS=Homo sapiens OX=9606 GN=TMEM189-UBE2V1 PE=4 SV=1 | 14.99 | 44.44 | 2.97 | 1.57 |
| tr\|E9PKG1\|E9PKG1_HUMAN Protein arginine N-methyltransferase 1 OS=Homo sapiens OX=9606 GN=PRMT1 PE=1 SV=1 | 14.92 | 30.43 | 2.04 | 1.03 |
| sp\|Q86YZ3\|HORN_HUMAN Hornerin OS=Homo sapiens OX=9606 GN=HRNR PE=1 SV=2 | 14.89 | 10.78 | 0.72 | -0.47 |
| sp\|O75531\|BAF_HUMAN Barrier-to-autointegration factor OS=Homo sapiens OX=9606 GN=BANF1 PE=1 SV=1 | 14.88 | 36.19 | 2.43 | 1.28 |
| sp\|P55735\|SEC13_HUMAN Protein SEC13 homolog OS=Homo sapiens OX=9606 GN=SEC13 PE=1 SV=3 | 14.81 | 28.47 | 1.92 | 0.94 |
| sp\|Q13595-2\|TRA2A_HUMAN Isoform Short of Transformer-2 protein homolog alpha OS=Homo sapiens OX=9606 GN=TRA2A | 14.79 | 10.03 | 0.68 | -0.56 |
| sp\|P31949\|S10AB_HUMAN Protein S100-A11 OS=Homo sapiens OX=9606 GN=S100A11 PE=1 SV=2 | 14.70 | 81.69 | 5.56 | 2.47 |
| sp\|P47897\|SYQ_HUMAN Glutamine--tRNA ligase OS=Homo sapiens OX=9606 GN=QARS PE=1 SV=1 | 14.70 | 54.41 | 3.70 | 1.89 |
| tr\|F8VXJ7\|F8VXJ7_HUMAN Protein canopy homolog 2 (Fragment) OS=Homo sapiens OX=9606 GN=CNPY2 PE=1 SV=1 | 14.66 | 15.86 | 1.08 | 0.11 |
| sp\|Q9H2P9-4\|DPH5_HUMAN Isoform 4 of Diphthine methyl ester synthase OS=Homo sapiens OX=9606 GN=DPH5 | 14.64 | 30.44 | 2.08 | 1.06 |
| sp\|O00764\|PDXK_HUMAN Pyridoxal kinase OS=Homo sapiens OX=9606 GN=PDXK PE=1 SV=1 | 14.63 | 3.94 | 0.27 | -1.89 |
| sp\|Q7Z333-3\|SETX_HUMAN Isoform 3 of Probable helicase senataxin OS=Homo sapiens OX=9606 GN=SETX | 14.55 | 30.78 | 2.12 | 1.08 |
| sp\|P17655\|CAN2_HUMAN Calpain-2 catalytic subunit OS=Homo sapiens OX=9606 GN=CAPN2 PE=1 SV=6 | 14.51 | 35.79 | 2.47 | 1.30 |

| | | | | |
|---|---|---|---|---|
| sp\|Q9BUL8\|PDC10_HUMAN Programmed cell death protein 10 OS=Homo sapiens OX=9606 GN=PDCD10 PE=1 SV=1 | 14.43 | 43.68 | 3.03 | 1.60 |
| sp\|O75489\|NDUS3_HUMAN NADH dehydrogenase [ubiquinone] iron-sulfur protein 3, mitochondrial OS=Homo sapiens OX=9606 GN=NDUFS3 PE=1 SV=1 | 14.37 | 26.09 | 1.82 | 0.86 |
| tr\|A0A1B0GTJ7\|A0A1B0GTJ7_HUMAN Adenylosuccinate lyase OS=Homo sapiens OX=9606 GN=ADSL PE=1 SV=1 | 14.37 | 32.14 | 2.24 | 1.16 |
| sp\|Q16401\|PSMD5_HUMAN 26S proteasome non-ATPase regulatory subunit 5 OS=Homo sapiens OX=9606 GN=PSMD5 PE=1 SV=3 | 14.35 | 26.30 | 1.83 | 0.87 |
| sp\|P42765\|THIM_HUMAN 3-ketoacyl-CoA thiolase, mitochondrial OS=Homo sapiens OX=9606 GN=ACAA2 PE=1 SV=2 | 14.35 | 42.26 | 2.94 | 1.56 |
| sp\|Q00688\|FKBP3_HUMAN Peptidyl-prolyl cis-trans isomerase FKBP3 OS=Homo sapiens OX=9606 GN=FKBP3 PE=1 SV=1 | 14.32 | 7.55 | 0.53 | -0.92 |
| sp\|Q96C19\|EFHD2_HUMAN EF-hand domain-containing protein D2 OS=Homo sapiens OX=9606 GN=EFHD2 PE=1 SV=1 | 14.31 | 15.98 | 1.12 | 0.16 |
| sp\|P54578-2\|UBP14_HUMAN Isoform 2 of Ubiquitin carboxyl-terminal hydrolase 14 OS=Homo sapiens OX=9606 GN=USP14 | 14.28 | 45.96 | 3.22 | 1.69 |
| tr\|K7ER16\|K7ER16_HUMAN Phenylalanine--tRNA ligase alpha subunit OS=Homo sapiens OX=9606 GN=FARSA PE=1 SV=1 | 14.24 | 9.62 | 0.68 | -0.57 |
| sp\|P50402\|EMD_HUMAN Emerin OS=Homo sapiens OX=9606 GN=EMD PE=1 SV=1 | 14.24 | 12.61 | 0.89 | -0.18 |
| tr\|A2A274\|A2A274_HUMAN Aconitate hydratase, mitochondrial OS=Homo sapiens OX=9606 GN=ACO2 PE=1 SV=1 | 14.23 | 34.83 | 2.45 | 1.29 |
| sp\|Q3MHD2-2\|LSM12_HUMAN Isoform 2 of Protein LSM12 homolog OS=Homo sapiens OX=9606 GN=LSM12 | 14.21 | 28.34 | 1.99 | 1.00 |
| sp\|Q00796\|DHSO_HUMAN Sorbitol dehydrogenase OS=Homo sapiens OX=9606 GN=SORD PE=1 SV=4 | 14.20 | 41.92 | 2.95 | 1.56 |
| sp\|Q96EY1-2\|DNJA3_HUMAN Isoform 2 of DnaJ homolog subfamily A member 3, mitochondrial OS=Homo sapiens OX=9606 GN=DNAJA3 | 14.09 | 21.75 | 1.54 | 0.63 |
| tr\|J3KNQ4\|J3KNQ4_HUMAN Alpha-parvin OS=Homo sapiens OX=9606 GN=PARVA PE=1 SV=1 | 14.08 | 13.58 | 0.96 | -0.05 |
| sp\|P30405\|PPIF_HUMAN Peptidyl-prolyl cis-trans isomerase F, mitochondrial OS=Homo sapiens OX=9606 GN=PPIF PE=1 SV=1 | 14.07 | 47.62 | 3.38 | 1.76 |
| sp\|Q9Y3I0\|RTCB_HUMAN tRNA-splicing ligase RtcB homolog OS=Homo sapiens OX=9606 GN=RTCB PE=1 SV=1 | 14.06 | 47.73 | 3.39 | 1.76 |
| sp\|Q16795\|NDUA9_HUMAN NADH dehydrogenase [ubiquinone] 1 alpha | 13.94 | 18.26 | 1.31 | 0.39 |

| | | | | |
|---|---|---|---|---|
| subcomplex subunit 9, mitochondrial OS=Homo sapiens OX=9606 GN=NDUFA9 PE=1 SV=2 | | | | |
| sp\|P61081\|UBC12_HUMAN NEDD8-conjugating enzyme Ubc12 OS=Homo sapiens OX=9606 GN=UBE2M PE=1 SV=1 | 13.93 | 35.83 | 2.57 | 1.36 |
| sp\|P12081-4\|SYHC_HUMAN Isoform 4 of Histidine--tRNA ligase, cytoplasmic OS=Homo sapiens OX=9606 GN=HARS | 13.92 | 28.71 | 2.06 | 1.04 |
| tr\|G3V5T9\|G3V5T9_HUMAN Cyclin-dependent kinase 2 OS=Homo sapiens OX=9606 GN=CDK2 PE=1 SV=1 | 13.91 | 22.33 | 1.60 | 0.68 |
| sp\|Q9UHV9\|PFD2_HUMAN Prefoldin subunit 2 OS=Homo sapiens OX=9606 GN=PFDN2 PE=1 SV=1 | 13.79 | 61.40 | 4.45 | 2.15 |
| tr\|H0YFD6\|H0YFD6_HUMAN Trifunctional enzyme subunit alpha, mitochondrial OS=Homo sapiens OX=9606 GN=HADHA PE=1 SV=2 | 13.73 | 34.70 | 2.53 | 1.34 |
| sp\|Q08945\|SSRP1_HUMAN FACT complex subunit SSRP1 OS=Homo sapiens OX=9606 GN=SSRP1 PE=1 SV=1 | 13.69 | 8.53 | 0.62 | -0.68 |
| tr\|A0A0A0MRF6\|A0A0A0MRF6_HUMAN A-kinase anchor protein 9 OS=Homo sapiens OX=9606 GN=AKAP9 PE=1 SV=1 | 13.67 | 11.30 | 0.83 | -0.27 |
| tr\|A0A0A0MRJ6\|A0A0A0MRJ6_HUMAN Protein-L-isoaspartate O-methyltransferase OS=Homo sapiens OX=9606 GN=PCMT1 PE=1 SV=1 | 13.58 | 69.77 | 5.14 | 2.36 |
| sp\|Q9UBI6\|GBG12_HUMAN Guanine nucleotide-binding protein G(I)/G(S)/G(O) subunit gamma-12 OS=Homo sapiens OX=9606 GN=GNG12 PE=1 SV=3 | 13.56 | 10.77 | 0.79 | -0.33 |
| tr\|J3KQ32\|J3KQ32_HUMAN Obg-like ATPase 1 OS=Homo sapiens OX=9606 GN=OLA1 PE=1 SV=1 | 13.55 | 15.84 | 1.17 | 0.22 |
| tr\|F8W7Q4\|F8W7Q4_HUMAN Protein FAM162A OS=Homo sapiens OX=9606 GN=FAM162A PE=1 SV=1 | 13.48 | 87.69 | 6.50 | 2.70 |
| sp\|P20073-2\|ANXA7_HUMAN Isoform 2 of Annexin A7 OS=Homo sapiens OX=9606 GN=ANXA7 | 13.39 | 49.26 | 3.68 | 1.88 |
| sp\|P54886-2\|P5CS_HUMAN Isoform Short of Delta-1-pyrroline-5-carboxylate synthase OS=Homo sapiens OX=9606 GN=ALDH18A1 | 13.37 | 20.75 | 1.55 | 0.63 |
| sp\|Q92973-2\|TNPO1_HUMAN Isoform 2 of Transportin-1 OS=Homo sapiens OX=9606 GN=TNPO1 | 13.36 | 27.03 | 2.02 | 1.02 |
| tr\|F5H4X0\|F5H4X0_HUMAN Scavenger receptor class B member 1 OS=Homo sapiens OX=9606 GN=SCARB1 PE=1 SV=1 | 13.32 | 13.51 | 1.01 | 0.02 |
| tr\|B4DR61\|B4DR61_HUMAN cDNA FLJ59739, highly similar to Protein transport protein Sec61 | 13.23 | 6.23 | 0.47 | -1.09 |

| | | | | |
|---|---|---|---|---|
| subunit alpha isoform 1 OS=Homo sapiens OX=9606 GN=SEC61A1 PE=1 SV=1 | | | | |
| tr\|B1AJY5\|B1AJY5_HUMAN 26S proteasome non-ATPase regulatory subunit 10 OS=Homo sapiens OX=9606 GN=PSMD10 PE=1 SV=1 | 13.23 | 12.63 | 0.95 | -0.07 |
| sp\|Q9UHD8-2\|SEPT9_HUMAN Isoform 2 of Septin-9 OS=Homo sapiens OX=9606 GN=SEPT9 | 13.21 | 14.16 | 1.07 | 0.10 |
| sp\|P29966\|MARCS_HUMAN Myristoylated alanine-rich C-kinase substrate OS=Homo sapiens OX=9606 GN=MARCKS PE=1 SV=4 | 13.16 | 9.90 | 0.75 | -0.41 |
| tr\|H7BYV1\|H7BYV1_HUMAN Interferon-induced transmembrane protein 2 (Fragment) OS=Homo sapiens OX=9606 GN=IFITM2 PE=4 SV=1 | 13.07 | 49.00 | 3.75 | 1.91 |
| sp\|Q15637-2\|SF01_HUMAN Isoform 2 of Splicing factor 1 OS=Homo sapiens OX=9606 GN=SF1 | 13.04 | 20.32 | 1.56 | 0.64 |
| tr\|A0A2R8Y3N3\|A0A2R8Y3N3_HUMAN Probable histidine--tRNA ligase, mitochondrial OS=Homo sapiens OX=9606 GN=HARS2 PE=1 SV=1 | 13.01 | 25.08 | 1.93 | 0.95 |
| tr\|B4DUC8\|B4DUC8_HUMAN S-methyl-5'-thioadenosine phosphorylase OS=Homo sapiens OX=9606 GN=MTAP PE=1 SV=1 | 12.97 | 68.25 | 5.26 | 2.40 |
| sp\|Q13501-2\|SQSTM_HUMAN Isoform 2 of Sequestosome-1 OS=Homo sapiens OX=9606 GN=SQSTM1 | 12.92 | 33.10 | 2.56 | 1.36 |
| sp\|P61106\|RAB14_HUMAN Ras-related protein Rab-14 OS=Homo sapiens OX=9606 GN=RAB14 PE=1 SV=4 | 12.87 | 14.93 | 1.16 | 0.21 |
| sp\|O00592-2\|PODXL_HUMAN Isoform 2 of Podocalyxin OS=Homo sapiens OX=9606 GN=PODXL | 12.85 | 30.42 | 2.37 | 1.24 |
| sp\|P43490\|NAMPT_HUMAN Nicotinamide phosphoribosyltransferase OS=Homo sapiens OX=9606 GN=NAMPT PE=1 SV=1 | 12.79 | 31.84 | 2.49 | 1.32 |
| sp\|Q14914-2\|PTGR1_HUMAN Isoform 2 of Prostaglandin reductase 1 OS=Homo sapiens OX=9606 GN=PTGR1 | 12.77 | 15.02 | 1.18 | 0.23 |
| sp\|P45973\|CBX5_HUMAN Chromobox protein homolog 5 OS=Homo sapiens OX=9606 GN=CBX5 PE=1 SV=1 | 12.73 | 18.05 | 1.42 | 0.50 |
| sp\|P43121\|MUC18_HUMAN Cell surface glycoprotein MUC18 OS=Homo sapiens OX=9606 GN=MCAM PE=1 SV=2 | 12.72 | 29.01 | 2.28 | 1.19 |
| tr\|M0QXB4\|M0QXB4_HUMAN Coatomer protein complex, subunit epsilon, isoform CRA_g OS=Homo sapiens OX=9606 GN=COPE PE=1 SV=1 | 12.65 | 49.03 | 3.88 | 1.95 |
| sp\|Q8N4V1-2\|MMGT1_HUMAN Isoform 2 of Membrane magnesium transporter 1 OS=Homo sapiens OX=9606 GN=MMGT1 | 12.57 | 45.78 | 3.64 | 1.86 |

| | | | | |
|---|---|---|---|---|
| sp\|Q9HCN8\|SDF2L_HUMAN Stromal cell-derived factor 2-like protein 1 OS=Homo sapiens OX=9606 GN=SDF2L1 PE=1 SV=2 | 12.57 | 41.01 | 3.26 | 1.71 |
| sp\|Q9BY43-2\|CHM4A_HUMAN Isoform 2 of Charged multivesicular body protein 4a OS=Homo sapiens OX=9606 GN=CHMP4A | 12.56 | 44.92 | 3.58 | 1.84 |
| sp\|O75629\|CREG1_HUMAN Protein CREG1 OS=Homo sapiens OX=9606 GN=CREG1 PE=1 SV=1 | 12.54 | 27.20 | 2.17 | 1.12 |
| sp\|P53007\|TXTP_HUMAN Tricarboxylate transport protein, mitochondrial OS=Homo sapiens OX=9606 GN=SLC25A1 PE=1 SV=2 | 12.51 | 28.81 | 2.30 | 1.20 |
| tr\|H0YIB4\|H0YIB4_HUMAN Serine/arginine-rich-splicing factor 9 (Fragment) OS=Homo sapiens OX=9606 GN=SRSF9 PE=1 SV=8 | 12.46 | 0.95 | 0.08 | -3.72 |
| tr\|H7C2Q8\|H7C2Q8_HUMAN EBNA1 binding protein 2, isoform CRA_d OS=Homo sapiens OX=9606 GN=EBNA1BP2 PE=1 SV=1 | 12.44 | 15.94 | 1.28 | 0.36 |
| tr\|A0A087WZN1\|A0A087WZN1_HUMAN Isocitrate dehydrogenase [NAD] subunit, mitochondrial OS=Homo sapiens OX=9606 GN=IDH3B PE=1 SV=1 | 12.35 | 8.09 | 0.65 | -0.61 |
| sp\|P0DN79\|CBSL_HUMAN Cystathionine beta-synthase-like protein OS=Homo sapiens OX=9606 GN=CBSL PE=1 SV=1 | 12.35 | 20.48 | 1.66 | 0.73 |
| tr\|A0A1B0GTG2\|A0A1B0GTG2_HUMAN Alpha-aminoadipic semialdehyde dehydrogenase OS=Homo sapiens OX=9606 GN=ALDH7A1 PE=1 SV=1 | 12.34 | 27.87 | 2.26 | 1.18 |
| sp\|Q14166\|TTL12_HUMAN Tubulin--tyrosine ligase-like protein 12 OS=Homo sapiens OX=9606 GN=TTLL12 PE=1 SV=2 | 12.31 | 27.51 | 2.24 | 1.16 |
| tr\|G5E9R5\|G5E9R5_HUMAN Acid phosphatase 1, soluble, isoform CRA_d OS=Homo sapiens OX=9606 GN=ACP1 PE=1 SV=1 | 12.25 | 34.72 | 2.84 | 1.50 |
| sp\|Q9BRA2\|TXD17_HUMAN Thioredoxin domain-containing protein 17 OS=Homo sapiens OX=9606 GN=TXNDC17 PE=1 SV=1 | 12.23 | 44.80 | 3.66 | 1.87 |
| tr\|F5GZQ3\|F5GZQ3_HUMAN Trifunctional enzyme subunit beta, mitochondrial OS=Homo sapiens OX=9606 GN=HADHB PE=1 SV=1 | 12.23 | 25.98 | 2.12 | 1.09 |
| sp\|Q8TDN6\|BRX1_HUMAN Ribosome biogenesis protein BRX1 homolog OS=Homo sapiens OX=9606 GN=BRIX1 PE=1 SV=2 | 12.15 | 19.08 | 1.57 | 0.65 |
| tr\|B1AHB1\|B1AHB1_HUMAN DNA helicase OS=Homo sapiens OX=9606 GN=MCM5 PE=1 SV=1 | 12.02 | 11.20 | 0.93 | -0.10 |
| sp\|Q96EK6\|GNA1_HUMAN Glucosamine 6-phosphate N-acetyltransferase OS=Homo sapiens OX=9606 GN=GNPNAT1 PE=1 SV=1 | 12.02 | 35.08 | 2.92 | 1.55 |
| tr\|E7EX17\|E7EX17_HUMAN Eukaryotic translation initiation factor 4B OS=Homo sapiens OX=9606 GN=EIF4B PE=1 SV=1 | 11.98 | 23.39 | 1.95 | 0.97 |

| | | | | |
|---|---|---|---|---|
| sp\|Q9NQ88\|TIGAR_HUMAN Fructose-2,6-bisphosphatase TIGAR OS=Homo sapiens OX=9606 GN=TIGAR PE=1 SV=1 | 11.95 | 27.14 | 2.27 | 1.18 |
| sp\|O95292\|VAPB_HUMAN Vesicle-associated membrane protein-associated protein B/C OS=Homo sapiens OX=9606 GN=VAPB PE=1 SV=3 | 11.92 | 27.13 | 2.28 | 1.19 |
| tr\|Q5T5C7\|Q5T5C7_HUMAN Serine--tRNA ligase, cytoplasmic OS=Homo sapiens OX=9606 GN=SARS PE=1 SV=1 | 11.90 | 35.93 | 3.02 | 1.59 |
| tr\|F8VQE1\|F8VQE1_HUMAN LIM domain and actin-binding protein 1 OS=Homo sapiens OX=9606 GN=LIMA1 PE=1 SV=1 | 11.87 | 57.87 | 4.88 | 2.29 |
| sp\|P63096\|GNAI1_HUMAN Guanine nucleotide-binding protein G(i) subunit alpha-1 OS=Homo sapiens OX=9606 GN=GNAI1 PE=1 SV=2 | 11.86 | 8.22 | 0.69 | -0.53 |
| tr\|D6RHJ3\|D6RHJ3_HUMAN Calnexin (Fragment) OS=Homo sapiens OX=9606 GN=CANX PE=1 SV=8 | 11.86 | 56.42 | 4.76 | 2.25 |
| sp\|P31930\|QCR1_HUMAN Cytochrome b-c1 complex subunit 1, mitochondrial OS=Homo sapiens OX=9606 GN=UQCRC1 PE=1 SV=3 | 11.85 | 32.81 | 2.77 | 1.47 |
| sp\|Q14165\|MLEC_HUMAN Malectin OS=Homo sapiens OX=9606 GN=MLEC PE=1 SV=1 | 11.84 | 30.62 | 2.59 | 1.37 |
| sp\|Q9BTV4\|TMM43_HUMAN Transmembrane protein 43 OS=Homo sapiens OX=9606 GN=TMEM43 PE=1 SV=1 | 11.83 | 10.81 | 0.91 | -0.13 |
| sp\|P40261\|NNMT_HUMAN Nicotinamide N-methyltransferase OS=Homo sapiens OX=9606 GN=NNMT PE=1 SV=1 | 11.76 | 8.70 | 0.74 | -0.44 |
| sp\|P10301\|RRAS_HUMAN Ras-related protein R-Ras OS=Homo sapiens OX=9606 GN=RRAS PE=1 SV=1 | 11.76 | 0.84 | 0.07 | -3.80 |
| sp\|P33991\|MCM4_HUMAN DNA replication licensing factor MCM4 OS=Homo sapiens OX=9606 GN=MCM4 PE=1 SV=5 | 11.64 | 1.05 | 0.09 | -3.47 |
| sp\|O95336\|6PGL_HUMAN 6-phosphogluconolactonase OS=Homo sapiens OX=9606 GN=PGLS PE=1 SV=2 | 11.63 | 42.40 | 3.65 | 1.87 |
| tr\|M0R0I3\|M0R0I3_HUMAN Endophilin-A2 (Fragment) OS=Homo sapiens OX=9606 GN=SH3GL1 PE=1 SV=1 | 11.62 | 37.70 | 3.24 | 1.70 |
| sp\|P0DN76\|U2AF5_HUMAN Splicing factor U2AF 35 kDa subunit-like protein OS=Homo sapiens OX=9606 GN=U2AF1L5 PE=1 SV=1 | 11.58 | 43.64 | 3.77 | 1.91 |
| sp\|P11172\|UMPS_HUMAN Uridine 5'-monophosphate synthase OS=Homo sapiens OX=9606 GN=UMPS PE=1 SV=1 | 11.58 | 18.85 | 1.63 | 0.70 |
| sp\|O95340-2\|PAPS2_HUMAN Isoform B of Bifunctional 3'-phosphoadenosine 5'-phosphosulfate synthase 2 OS=Homo sapiens OX=9606 GN=PAPSS2 | 11.57 | 39.71 | 3.43 | 1.78 |

| | | | | |
|---|---|---|---|---|
| sp\|O96019\|ACL6A_HUMAN Actin-like protein 6A OS=Homo sapiens OX=9606 GN=ACTL6A PE=1 SV=1 | 11.55 | 18.35 | 1.59 | 0.67 |
| tr\|J3KN29\|J3KN29_HUMAN 26S proteasome non-ATPase regulatory subunit 9 OS=Homo sapiens OX=9606 GN=PSMD9 PE=1 SV=1 | 11.55 | 3.30 | 0.29 | -1.81 |
| tr\|H0Y368\|H0Y368_HUMAN Dolichol-phosphate mannosyltransferase subunit 1 (Fragment) OS=Homo sapiens OX=9606 GN=DPM1 PE=1 SV=1 | 11.53 | 14.32 | 1.24 | 0.31 |
| sp\|Q05682\|CALD1_HUMAN Caldesmon OS=Homo sapiens OX=9606 GN=CALD1 PE=1 SV=3 | 11.53 | 10.53 | 0.91 | -0.13 |
| tr\|I3L1Y9\|I3L1Y9_HUMAN FLYWCH family member 2 OS=Homo sapiens OX=9606 GN=FLYWCH2 PE=1 SV=1 | 11.37 | 8.49 | 0.75 | -0.42 |
| sp\|Q96RS6-2\|NUDC1_HUMAN Isoform 2 of NudC domain-containing protein 1 OS=Homo sapiens OX=9606 GN=NUDCD1 | 11.28 | 12.89 | 1.14 | 0.19 |
| sp\|Q15388\|TOM20_HUMAN Mitochondrial import receptor subunit TOM20 homolog OS=Homo sapiens OX=9606 GN=TOMM20 PE=1 SV=1 | 11.26 | 32.83 | 2.92 | 1.54 |
| sp\|Q9H4A4\|AMPB_HUMAN Aminopeptidase B OS=Homo sapiens OX=9606 GN=RNPEP PE=1 SV=2 | 11.21 | 20.00 | 1.78 | 0.84 |
| sp\|Q9UNE7\|CHIP_HUMAN E3 ubiquitin-protein ligase CHIP OS=Homo sapiens OX=9606 GN=STUB1 PE=1 SV=2 | 11.19 | 28.09 | 2.51 | 1.33 |
| sp\|P49189\|AL9A1_HUMAN 4-trimethylaminobutyraldehyde dehydrogenase OS=Homo sapiens OX=9606 GN=ALDH9A1 PE=1 SV=3 | 11.19 | 17.72 | 1.58 | 0.66 |
| sp\|P42025\|ACTY_HUMAN Beta-centractin OS=Homo sapiens OX=9606 GN=ACTR1B PE=1 SV=1 | 11.15 | 2.25 | 0.20 | -2.31 |
| sp\|P20645\|MPRD_HUMAN Cation-dependent mannose-6-phosphate receptor OS=Homo sapiens OX=9606 GN=M6PR PE=1 SV=1 | 11.07 | 23.30 | 2.10 | 1.07 |
| sp\|O15144\|ARPC2_HUMAN Actin-related protein 2/3 complex subunit 2 OS=Homo sapiens OX=9606 GN=ARPC2 PE=1 SV=1 | 10.96 | 28.95 | 2.64 | 1.40 |
| tr\|K7ERJ1\|K7ERJ1_HUMAN Thymidine kinase (Fragment) OS=Homo sapiens OX=9606 GN=TK1 PE=1 SV=1 | 10.96 | 34.67 | 3.16 | 1.66 |
| sp\|Q9Y2V2\|CHSP1_HUMAN Calcium-regulated heat-stable protein 1 OS=Homo sapiens OX=9606 GN=CARHSP1 PE=1 SV=2 | 10.95 | 43.25 | 3.95 | 1.98 |
| sp\|P11233\|RALA_HUMAN Ras-related protein Ral-A OS=Homo sapiens OX=9606 GN=RALA PE=1 SV=1 | 10.92 | 3.52 | 0.32 | -1.63 |
| tr\|A0A286YEX5\|A0A286YEX5_HUMAN MICOS complex subunit (Fragment) OS=Homo sapiens OX=9606 GN=CHCHD3 PE=1 SV=1 | 10.87 | 2.05 | 0.19 | -2.41 |

| | | | | |
|---|---|---|---|---|
| sp\|Q07960\|RHG01_HUMAN Rho GTPase-activating protein 1 OS=Homo sapiens OX=9606 GN=ARHGAP1 PE=1 SV=1 | 10.87 | 19.66 | 1.81 | 0.85 |
| sp\|P36507\|MP2K2_HUMAN Dual specificity mitogen-activated protein kinase kinase 2 OS=Homo sapiens OX=9606 GN=MAP2K2 PE=1 SV=1 | 10.86 | 14.52 | 1.34 | 0.42 |
| sp\|Q02750\|MP2K1_HUMAN Dual specificity mitogen-activated protein kinase kinase 1 OS=Homo sapiens OX=9606 GN=MAP2K1 PE=1 SV=2 | 10.86 | 14.13 | 1.30 | 0.38 |
| tr\|H0Y2V1\|H0Y2V1_HUMAN Microtubule-associated protein (Fragment) OS=Homo sapiens OX=9606 GN=MAP4 PE=1 SV=1 | 10.77 | 14.46 | 1.34 | 0.42 |
| tr\|G3V1C3\|G3V1C3_HUMAN Apoptosis inhibitor 5 OS=Homo sapiens OX=9606 GN=API5 PE=1 SV=1 | 10.76 | 43.20 | 4.02 | 2.01 |
| tr\|R4GMR5\|R4GMR5_HUMAN 26S proteasome non-ATPase regulatory subunit 8 OS=Homo sapiens OX=9606 GN=PSMD8 PE=1 SV=1 | 10.74 | 31.90 | 2.97 | 1.57 |
| sp\|Q53EL6-2\|PDCD4_HUMAN Isoform 2 of Programmed cell death protein 4 OS=Homo sapiens OX=9606 GN=PDCD4 | 10.70 | 16.50 | 1.54 | 0.63 |
| tr\|H0Y8X4\|H0Y8X4_HUMAN 2'-deoxynucleoside 5'-phosphate N-hydrolase 1 (Fragment) OS=Homo sapiens OX=9606 GN=DNPH1 PE=1 SV=1 | 10.69 | 37.47 | 3.51 | 1.81 |
| tr\|B1AMS2\|B1AMS2_HUMAN Septin 6, isoform CRA_b OS=Homo sapiens OX=9606 GN=SEPT6 PE=1 SV=1 | 10.66 | 25.47 | 2.39 | 1.26 |
| sp\|Q9H074-2\|PAIP1_HUMAN Isoform 2 of Polyadenylate-binding protein-interacting protein 1 OS=Homo sapiens OX=9606 GN=PAIP1 | 10.66 | 14.96 | 1.40 | 0.49 |
| sp\|P38606-2\|VATA_HUMAN Isoform 2 of V-type proton ATPase catalytic subunit A OS=Homo sapiens OX=9606 GN=ATP6V1A | 10.66 | 14.67 | 1.38 | 0.46 |
| sp\|Q96IZ0\|PAWR_HUMAN PRKC apoptosis WT1 regulator protein OS=Homo sapiens OX=9606 GN=PAWR PE=1 SV=1 | 10.62 | 1.48 | 0.14 | -2.84 |
| sp\|Q15646\|OASL_HUMAN 2'-5'-oligoadenylate synthase-like protein OS=Homo sapiens OX=9606 GN=OASL PE=1 SV=2 | 10.56 | 9.82 | 0.93 | -0.10 |
| sp\|P52943\|CRIP2_HUMAN Cysteine-rich protein 2 OS=Homo sapiens OX=9606 GN=CRIP2 PE=1 SV=1 | 10.54 | 36.74 | 3.48 | 1.80 |
| sp\|P27694\|RFA1_HUMAN Replication protein A 70 kDa DNA-binding subunit OS=Homo sapiens OX=9606 GN=RPA1 PE=1 SV=2 | 10.53 | 11.77 | 1.12 | 0.16 |
| tr\|K7ENG2\|K7ENG2_HUMAN U2 snRNP auxiliary factor large subunit OS=Homo sapiens OX=9606 GN=U2AF2 PE=1 SV=1 | 10.51 | 35.32 | 3.36 | 1.75 |

| | | | | |
|---|---|---|---|---|
| sp\|Q9NZ45\|CISD1_HUMAN CDGSH iron-sulfur domain-containing protein 1 OS=Homo sapiens OX=9606 GN=CISD1 PE=1 SV=1 | 10.51 | 33.47 | 3.19 | 1.67 |
| sp\|P61086\|UBE2K_HUMAN Ubiquitin-conjugating enzyme E2 K OS=Homo sapiens OX=9606 GN=UBE2K PE=1 SV=3 | 10.45 | 18.14 | 1.74 | 0.80 |
| sp\|Q969X5\|ERGI1_HUMAN Endoplasmic reticulum-Golgi intermediate compartment protein 1 OS=Homo sapiens OX=9606 GN=ERGIC1 PE=1 SV=1 | 10.44 | 30.69 | 2.94 | 1.56 |
| sp\|P51665\|PSMD7_HUMAN 26S proteasome non-ATPase regulatory subunit 7 OS=Homo sapiens OX=9606 GN=PSMD7 PE=1 SV=2 | 10.44 | 29.73 | 2.85 | 1.51 |
| sp\|Q9Y224\|RTRAF_HUMAN RNA transcription, translation and transport factor protein OS=Homo sapiens OX=9606 GN=RTRAF PE=1 SV=1 | 10.40 | 30.65 | 2.95 | 1.56 |
| sp\|O43143\|DHX15_HUMAN Pre-mRNA-splicing factor ATP-dependent RNA helicase DHX15 OS=Homo sapiens OX=9606 GN=DHX15 PE=1 SV=2 | 10.30 | 15.47 | 1.50 | 0.59 |
| sp\|P13861-2\|KAP2_HUMAN Isoform 2 of cAMP-dependent protein kinase type II-alpha regulatory subunit OS=Homo sapiens OX=9606 GN=PRKAR2A | 10.30 | 14.46 | 1.40 | 0.49 |
| tr\|E9PEX6\|E9PEX6_HUMAN Dihydrolipoyl dehydrogenase OS=Homo sapiens OX=9606 GN=DLD PE=1 SV=1 | 10.25 | 17.22 | 1.68 | 0.75 |
| sp\|O95433\|AHSA1_HUMAN Activator of 90 kDa heat shock protein ATPase homolog 1 OS=Homo sapiens OX=9606 GN=AHSA1 PE=1 SV=1 | 10.24 | 46.85 | 4.58 | 2.19 |
| sp\|P16949-2\|STMN1_HUMAN Isoform 2 of Stathmin OS=Homo sapiens OX=9606 GN=STMN1 | 10.23 | 29.94 | 2.93 | 1.55 |
| sp\|P51970\|NDUA8_HUMAN NADH dehydrogenase [ubiquinone] 1 alpha subcomplex subunit 8 OS=Homo sapiens OX=9606 GN=NDUFA8 PE=1 SV=3 | 10.22 | 20.62 | 2.02 | 1.01 |
| sp\|P21912\|SDHB_HUMAN Succinate dehydrogenase [ubiquinone] iron-sulfur subunit, mitochondrial OS=Homo sapiens OX=9606 GN=SDHB PE=1 SV=3 | 10.20 | 23.98 | 2.35 | 1.23 |
| sp\|P46060\|RAGP1_HUMAN Ran GTPase-activating protein 1 OS=Homo sapiens OX=9606 GN=RANGAP1 PE=1 SV=1 | 10.15 | 22.34 | 2.20 | 1.14 |
| tr\|D6RBV2\|D6RBV2_HUMAN Vesicular integral-membrane protein VIP36 OS=Homo sapiens OX=9606 GN=LMAN2 PE=1 SV=1 | 10.11 | 21.69 | 2.15 | 1.10 |
| sp\|Q96HE7\|ERO1A_HUMAN ERO1-like protein alpha OS=Homo sapiens OX=9606 GN=ERO1A PE=1 SV=2 | 10.10 | 10.60 | 1.05 | 0.07 |

| | | | | |
|---|---|---|---|---|
| sp\|P48507\|GSH0_HUMAN Glutamate--cysteine ligase regulatory subunit OS=Homo sapiens OX=9606 GN=GCLM PE=1 SV=1 | 10.06 | 26.96 | 2.68 | 1.42 |
| sp\|P42166\|LAP2A_HUMAN Lamina-associated polypeptide 2, isoform alpha OS=Homo sapiens OX=9606 GN=TMPO PE=1 SV=2 | 10.04 | 15.27 | 1.52 | 0.61 |
| sp\|Q9BWF3\|RBM4_HUMAN RNA-binding protein 4 OS=Homo sapiens OX=9606 GN=RBM4 PE=1 SV=1 | 9.98 | 31.57 | 3.16 | 1.66 |
| sp\|P52294\|IMA5_HUMAN Importin subunit alpha-5 OS=Homo sapiens OX=9606 GN=KPNA1 PE=1 SV=3 | 9.97 | 22.33 | 2.24 | 1.16 |
| tr\|J3QLE5\|J3QLE5_HUMAN Small nuclear ribonucleoprotein-associated protein N (Fragment) OS=Homo sapiens OX=9606 GN=SNRPN PE=1 SV=1 | 9.96 | 69.50 | 6.98 | 2.80 |
| sp\|Q9NUQ9\|FA49B_HUMAN Protein FAM49B OS=Homo sapiens OX=9606 GN=FAM49B PE=1 SV=1 | 9.92 | 27.86 | 2.81 | 1.49 |
| tr\|E5RFX4\|E5RFX4_HUMAN Lymphokine-activated killer T-cell-originated protein kinase (Fragment) OS=Homo sapiens OX=9606 GN=PBK PE=1 SV=2 | 9.92 | 36.72 | 3.70 | 1.89 |
| sp\|Q9Y2Z4\|SYYM_HUMAN Tyrosine--tRNA ligase, mitochondrial OS=Homo sapiens OX=9606 GN=YARS2 PE=1 SV=2 | 9.88 | 2.84 | 0.29 | -1.80 |
| tr\|F5GX39\|F5GX39_HUMAN Transmembrane emp24 domain-containing protein 2 OS=Homo sapiens OX=9606 GN=TMED2 PE=1 SV=1 | 9.88 | 7.22 | 0.73 | -0.45 |
| sp\|Q15370-2\|ELOB_HUMAN Isoform 2 of Elongin-B OS=Homo sapiens OX=9606 GN=ELOB | 9.87 | 14.18 | 1.44 | 0.52 |
| sp\|O94925-3\|GLSK_HUMAN Isoform 3 of Glutaminase kidney isoform, mitochondrial OS=Homo sapiens OX=9606 GN=GLS | 9.83 | 14.76 | 1.50 | 0.59 |
| sp\|P82933\|RT09_HUMAN 28S ribosomal protein S9, mitochondrial OS=Homo sapiens OX=9606 GN=MRPS9 PE=1 SV=2 | 9.79 | 20.24 | 2.07 | 1.05 |
| sp\|Q9H7Z7\|PGES2_HUMAN Prostaglandin E synthase 2 OS=Homo sapiens OX=9606 GN=PTGES2 PE=1 SV=1 | 9.77 | 22.84 | 2.34 | 1.23 |
| sp\|P62304\|RUXE_HUMAN Small nuclear ribonucleoprotein E OS=Homo sapiens OX=9606 GN=SNRPE PE=1 SV=1 | 9.74 | 27.02 | 2.78 | 1.47 |
| tr\|B4DLR8\|B4DLR8_HUMAN NAD(P)H dehydrogenase [quinone] 1 OS=Homo sapiens OX=9606 GN=NQO1 PE=1 SV=1 | 9.73 | 7.93 | 0.81 | -0.30 |
| tr\|H0YKU1\|H0YKU1_HUMAN Tropomodulin-3 (Fragment) OS=Homo sapiens OX=9606 GN=TMOD3 PE=1 SV=1 | 9.68 | 25.52 | 2.64 | 1.40 |
| tr\|C9JKI3\|C9JKI3_HUMAN Caveolin (Fragment) OS=Homo sapiens OX=9606 GN=CAV1 PE=1 SV=1 | 9.68 | 21.04 | 2.17 | 1.12 |

| | | | | |
|---|---|---|---|---|
| sp\|P25815\|S100P_HUMAN Protein S100-P OS=Homo sapiens OX=9606 GN=S100P PE=1 SV=2 | 9.64 | 30.57 | 3.17 | 1.66 |
| sp\|Q6NUQ4-2\|TM214_HUMAN Isoform 2 of Transmembrane protein 214 OS=Homo sapiens OX=9606 GN=TMEM214 | 9.59 | 0.44 | 0.05 | -4.46 |
| sp\|Q9UBT2\|SAE2_HUMAN SUMO-activating enzyme subunit 2 OS=Homo sapiens OX=9606 GN=UBA2 PE=1 SV=2 | 9.56 | 1.92 | 0.20 | -2.32 |
| sp\|Q15008-4\|PSMD6_HUMAN Isoform 4 of 26S proteasome non-ATPase regulatory subunit 6 OS=Homo sapiens OX=9606 GN=PSMD6 | 9.53 | 13.49 | 1.42 | 0.50 |
| sp\|Q7L5N1\|CSN6_HUMAN COP9 signalosome complex subunit 6 OS=Homo sapiens OX=9606 GN=COPS6 PE=1 SV=1 | 9.51 | 15.41 | 1.62 | 0.70 |
| tr\|A0A2U3TZL5\|A0A2U3TZL5_HUMAN CD59 glycoprotein (Fragment) OS=Homo sapiens OX=9606 GN=CD59 PE=4 SV=1 | 9.51 | 16.02 | 1.69 | 0.75 |
| sp\|Q9UJS0-2\|CMC2_HUMAN Isoform 2 of Calcium-binding mitochondrial carrier protein Aralar2 OS=Homo sapiens OX=9606 GN=SLC25A13 | 9.46 | 15.01 | 1.59 | 0.67 |
| sp\|O14745\|NHRF1_HUMAN Na(+)/H(+) exchange regulatory cofactor NHE-RF1 OS=Homo sapiens OX=9606 GN=SLC9A3R1 PE=1 SV=4 | 9.41 | 6.40 | 0.68 | -0.56 |
| sp\|P51151\|RAB9A_HUMAN Ras-related protein Rab-9A OS=Homo sapiens OX=9606 GN=RAB9A PE=1 SV=1 | 9.41 | 26.99 | 2.87 | 1.52 |
| tr\|A0A1W2PNR9\|A0A1W2PNR9_HUMAN Immediate early response 3-interacting protein 1 OS=Homo sapiens OX=9606 GN=IER3IP1 PE=1 SV=1 | 9.36 | 24.35 | 2.60 | 1.38 |
| tr\|J3QRG6\|J3QRG6_HUMAN Cyclin-dependent kinase inhibitor 2A OS=Homo sapiens OX=9606 GN=CDKN2A PE=1 SV=1 | 9.35 | 34.60 | 3.70 | 1.89 |
| sp\|Q96B26\|EXOS8_HUMAN Exosome complex component RRP43 OS=Homo sapiens OX=9606 GN=EXOSC8 PE=1 SV=1 | 9.35 | 20.79 | 2.22 | 1.15 |
| sp\|Q9Y2Z0-2\|SGT1_HUMAN Isoform 2 of Protein SGT1 homolog OS=Homo sapiens OX=9606 GN=SUGT1 | 9.33 | 15.87 | 1.70 | 0.77 |
| sp\|Q6NUK1-2\|SCMC1_HUMAN Isoform 2 of Calcium-binding mitochondrial carrier protein SCaMC-1 OS=Homo sapiens OX=9606 GN=SLC25A24 | 9.29 | 12.73 | 1.37 | 0.45 |
| sp\|P50995-2\|ANX11_HUMAN Isoform 2 of Annexin A11 OS=Homo sapiens OX=9606 GN=ANXA11 | 9.24 | 18.41 | 1.99 | 0.99 |
| sp\|Q13868\|EXOS2_HUMAN Exosome complex component RRP4 OS=Homo sapiens OX=9606 GN=EXOSC2 PE=1 SV=2 | 9.23 | 22.76 | 2.47 | 1.30 |

| | | | | |
|---|---|---|---|---|
| sp\|Q9UDY8\|MALT1_HUMAN Mucosa-associated lymphoid tissue lymphoma translocation protein 1 OS=Homo sapiens OX=9606 GN=MALT1 PE=1 SV=1 | 9.21 | 20.54 | 2.23 | 1.16 |
| sp\|Q9Y383\|LC7L2_HUMAN Putative RNA-binding protein Luc7-like 2 OS=Homo sapiens OX=9606 GN=LUC7L2 PE=1 SV=2 | 9.13 | 8.37 | 0.92 | -0.13 |
| tr\|A0A0A6YYA0\|A0A0A6YYA0_HUMAN Protein TMED7-TICAM2 OS=Homo sapiens OX=9606 GN=TMED7-TICAM2 PE=3 SV=1 | 9.10 | 13.72 | 1.51 | 0.59 |
| sp\|Q9UJZ1-2\|STML2_HUMAN Isoform 2 of Stomatin-like protein 2, mitochondrial OS=Homo sapiens OX=9606 GN=STOML2 | 9.07 | 20.22 | 2.23 | 1.16 |
| tr\|D6W5Y5\|D6W5Y5_HUMAN Cold inducible RNA binding protein, isoform CRA_c OS=Homo sapiens OX=9606 GN=CIRBP PE=1 SV=1 | 9.07 | 15.84 | 1.75 | 0.80 |
| sp\|Q7L9L4-2\|MOB1B_HUMAN Isoform 2 of MOB kinase activator 1B OS=Homo sapiens OX=9606 GN=MOB1B | 9.05 | 14.07 | 1.55 | 0.64 |
| sp\|P53004\|BIEA_HUMAN Biliverdin reductase A OS=Homo sapiens OX=9606 GN=BLVRA PE=1 SV=2 | 9.03 | 34.15 | 3.78 | 1.92 |
| sp\|Q5JWF2-2\|GNAS1_HUMAN Isoform XLas-2 of Guanine nucleotide-binding protein G(s) subunit alpha isoforms XLas OS=Homo sapiens OX=9606 GN=GNAS | 9.03 | 26.05 | 2.89 | 1.53 |
| sp\|Q09161\|NCBP1_HUMAN Nuclear cap-binding protein subunit 1 OS=Homo sapiens OX=9606 GN=NCBP1 PE=1 SV=1 | 9.01 | 12.92 | 1.43 | 0.52 |
| sp\|Q9NPD8\|UBE2T_HUMAN Ubiquitin-conjugating enzyme E2 T OS=Homo sapiens OX=9606 GN=UBE2T PE=1 SV=1 | 9.01 | 24.85 | 2.76 | 1.46 |
| sp\|P49257\|LMAN1_HUMAN Protein ERGIC-53 OS=Homo sapiens OX=9606 GN=LMAN1 PE=1 SV=2 | 8.97 | 23.28 | 2.59 | 1.38 |
| sp\|O60493\|SNX3_HUMAN Sorting nexin-3 OS=Homo sapiens OX=9606 GN=SNX3 PE=1 SV=3 | 8.96 | 33.86 | 3.78 | 1.92 |
| sp\|Q9H444\|CHM4B_HUMAN Charged multivesicular body protein 4b OS=Homo sapiens OX=9606 GN=CHMP4B PE=1 SV=1 | 8.95 | 6.21 | 0.69 | -0.53 |
| sp\|P55010\|IF5_HUMAN Eukaryotic translation initiation factor 5 OS=Homo sapiens OX=9606 GN=EIF5 PE=1 SV=2 | 8.91 | 10.35 | 1.16 | 0.22 |
| tr\|M0QYZ2\|M0QYZ2_HUMAN AP complex subunit sigma OS=Homo sapiens OX=9606 GN=AP2S1 PE=1 SV=1 | 8.84 | 8.57 | 0.97 | -0.05 |
| sp\|Q15785\|TOM34_HUMAN Mitochondrial import receptor subunit TOM34 OS=Homo sapiens OX=9606 GN=TOMM34 PE=1 SV=2 | 8.79 | 10.49 | 1.19 | 0.26 |
| sp\|Q9P258\|RCC2_HUMAN Protein RCC2 OS=Homo sapiens OX=9606 GN=RCC2 PE=1 SV=2 | 8.73 | 4.21 | 0.48 | -1.05 |

| | | | | |
|---|---|---|---|---|
| tr\|A0A087WVZ9\|A0A087WVZ9_HUMAN DNA-directed RNA polymerases I, II, and III subunit RPABC1 OS=Homo sapiens OX=9606 GN=POLR2E PE=1 SV=1 | 8.72 | 11.34 | 1.30 | 0.38 |
| sp\|P08621\|RU17_HUMAN U1 small nuclear ribonucleoprotein 70 kDa OS=Homo sapiens OX=9606 GN=SNRNP70 PE=1 SV=2 | 8.72 | 10.04 | 1.15 | 0.20 |
| tr\|A0A2R8Y653\|A0A2R8Y653_HUMAN Uridine-cytidine kinase 2 OS=Homo sapiens OX=9606 GN=UCK2 PE=1 SV=1 | 8.71 | 27.66 | 3.18 | 1.67 |
| tr\|Q5TEJ7\|Q5TEJ7_HUMAN Replication protein A 32 kDa subunit (Fragment) OS=Homo sapiens OX=9606 GN=RPA2 PE=1 SV=1 | 8.70 | 17.16 | 1.97 | 0.98 |
| sp\|Q92734-2\|TFG_HUMAN Isoform 2 of Protein TFG OS=Homo sapiens OX=9606 GN=TFG | 8.67 | 18.36 | 2.12 | 1.08 |
| sp\|P61758\|PFD3_HUMAN Prefoldin subunit 3 OS=Homo sapiens OX=9606 GN=VBP1 PE=1 SV=3 | 8.65 | 13.99 | 1.62 | 0.69 |
| tr\|G3V1U5\|G3V1U5_HUMAN Golgi transport 1 homolog B (S. cerevisiae), isoform CRA_c OS=Homo sapiens OX=9606 GN=GOLT1B PE=1 SV=1 | 8.63 | 15.01 | 1.74 | 0.80 |
| sp\|Q9NVP1\|DDX18_HUMAN ATP-dependent RNA helicase DDX18 OS=Homo sapiens OX=9606 GN=DDX18 PE=1 SV=2 | 8.62 | 12.35 | 1.43 | 0.52 |
| sp\|O14949\|QCR8_HUMAN Cytochrome b-c1 complex subunit 8 OS=Homo sapiens OX=9606 GN=UQCRQ PE=1 SV=4 | 8.60 | 23.40 | 2.72 | 1.44 |
| sp\|P84095\|RHOG_HUMAN Rho-related GTP-binding protein RhoG OS=Homo sapiens OX=9606 GN=RHOG PE=1 SV=1 | 8.59 | 46.63 | 5.43 | 2.44 |
| tr\|H7C1U8\|H7C1U8_HUMAN MICOS complex subunit (Fragment) OS=Homo sapiens OX=9606 GN=APOO PE=1 SV=1 | 8.52 | 7.92 | 0.93 | -0.11 |
| tr\|I3L3Q4\|I3L3Q4_HUMAN Glyoxalase domain-containing protein 4 (Fragment) OS=Homo sapiens OX=9606 GN=GLOD4 PE=1 SV=1 | 8.49 | 13.81 | 1.63 | 0.70 |
| sp\|Q9BRJ6\|CG050_HUMAN Uncharacterized protein C7orf50 OS=Homo sapiens OX=9606 GN=C7orf50 PE=1 SV=1 | 8.49 | 30.99 | 3.65 | 1.87 |
| sp\|O75746-2\|CMC1_HUMAN Isoform 2 of Calcium-binding mitochondrial carrier protein Aralar1 OS=Homo sapiens OX=9606 GN=SLC25A12 | 8.48 | 14.16 | 1.67 | 0.74 |
| sp\|P55327-2\|TPD52_HUMAN Isoform 2 of Tumor protein D52 OS=Homo sapiens OX=9606 GN=TPD52 | 8.40 | 30.66 | 3.65 | 1.87 |
| sp\|Q13492-2\|PICAL_HUMAN Isoform 2 of Phosphatidylinositol-binding clathrin assembly protein OS=Homo sapiens OX=9606 GN=PICALM | 8.39 | 19.20 | 2.29 | 1.19 |

| | | | | |
|---|---|---|---|---|
| tr\|G3V0E4\|G3V0E4_HUMAN Mitochondrial-processing peptidase subunit beta OS=Homo sapiens OX=9606 GN=PMPCB PE=1 SV=1 | 8.38 | 14.91 | 1.78 | 0.83 |
| sp\|O95456-2\|PSMG1_HUMAN Isoform 2 of Proteasome assembly chaperone 1 OS=Homo sapiens OX=9606 GN=PSMG1 | 8.37 | 29.59 | 3.54 | 1.82 |
| sp\|Q9BX68\|HINT2_HUMAN Histidine triad nucleotide-binding protein 2, mitochondrial OS=Homo sapiens OX=9606 GN=HINT2 PE=1 SV=1 | 8.36 | 1.40 | 0.17 | -2.58 |
| sp\|P43034\|LIS1_HUMAN Platelet-activating factor acetylhydrolase IB subunit alpha OS=Homo sapiens OX=9606 GN=PAFAH1B1 PE=1 SV=2 | 8.36 | 17.37 | 2.08 | 1.06 |
| tr\|B1AH49\|B1AH49_HUMAN 3-mercaptopyruvate sulfurtransferase OS=Homo sapiens OX=9606 GN=MPST PE=1 SV=1 | 8.33 | 28.68 | 3.44 | 1.78 |
| sp\|O00625\|PIR_HUMAN Pirin OS=Homo sapiens OX=9606 GN=PIR PE=1 SV=1 | 8.32 | 21.54 | 2.59 | 1.37 |
| tr\|Q5T948\|Q5T948_HUMAN Serine/threonine-protein phosphatase 2A activator (Fragment) OS=Homo sapiens OX=9606 GN=PTPA PE=1 SV=8 | 8.32 | 12.04 | 1.45 | 0.53 |
| tr\|D6RAX7\|D6RAX7_HUMAN COP9 constitutive photomorphogenic-like protein subunit 4 isoform 2 OS=Homo sapiens OX=9606 GN=COPS4 PE=1 SV=1 | 8.29 | 23.88 | 2.88 | 1.53 |
| sp\|Q9UHG3\|PCYOX_HUMAN Prenylcysteine oxidase 1 OS=Homo sapiens OX=9606 GN=PCYOX1 PE=1 SV=3 | 8.27 | 16.26 | 1.97 | 0.98 |
| sp\|Q8NBU5-2\|ATAD1_HUMAN Isoform 2 of ATPase family AAA domain-containing protein 1 OS=Homo sapiens OX=9606 GN=ATAD1 | 8.26 | 19.82 | 2.40 | 1.26 |
| sp\|Q8NC51-2\|PAIRB_HUMAN Isoform 2 of Plasminogen activator inhibitor 1 RNA-binding protein OS=Homo sapiens OX=9606 GN=SERBP1 | 8.25 | 5.05 | 0.61 | -0.71 |
| sp\|Q9NSD9\|SYFB_HUMAN Phenylalanine--tRNA ligase beta subunit OS=Homo sapiens OX=9606 GN=FARSB PE=1 SV=3 | 8.24 | 11.40 | 1.38 | 0.47 |
| sp\|P19623\|SPEE_HUMAN Spermidine synthase OS=Homo sapiens OX=9606 GN=SRM PE=1 SV=1 | 8.20 | 26.43 | 3.22 | 1.69 |
| tr\|A0A087WYR3\|A0A087WYR3_HUMAN Tumor protein D54 OS=Homo sapiens OX=9606 GN=TPD52L2 PE=1 SV=1 | 8.19 | 24.98 | 3.05 | 1.61 |
| sp\|P55036\|PSMD4_HUMAN 26S proteasome non-ATPase regulatory subunit 4 OS=Homo sapiens OX=9606 GN=PSMD4 PE=1 SV=1 | 8.18 | 18.91 | 2.31 | 1.21 |
| sp\|P49207\|RL34_HUMAN 60S ribosomal protein L34 OS=Homo sapiens OX=9606 GN=RPL34 PE=1 SV=3 | 8.13 | 17.45 | 2.15 | 1.10 |

| | | | | |
|---|---|---|---|---|
| sp\|O00425\|IF2B3_HUMAN Insulin-like growth factor 2 mRNA-binding protein 3 OS=Homo sapiens OX=9606 GN=IGF2BP3 PE=1 SV=2 | 8.12 | 5.52 | 0.68 | -0.56 |
| tr\|H0YL72\|H0YL72_HUMAN Isocitrate dehydrogenase [NAD] subunit alpha, mitochondrial OS=Homo sapiens OX=9606 GN=IDH3A PE=1 SV=1 | 8.10 | 49.12 | 6.06 | 2.60 |
| sp\|Q9Y6C9\|MTCH2_HUMAN Mitochondrial carrier homolog 2 OS=Homo sapiens OX=9606 GN=MTCH2 PE=1 SV=1 | 8.08 | 22.49 | 2.78 | 1.48 |
| sp\|Q9BRX2\|PELO_HUMAN Protein pelota homolog OS=Homo sapiens OX=9606 GN=PELO PE=1 SV=2 | 8.06 | 14.64 | 1.82 | 0.86 |
| sp\|Q8WUX2\|CHAC2_HUMAN Glutathione-specific gamma-glutamylcyclotransferase 2 OS=Homo sapiens OX=9606 GN=CHAC2 PE=1 SV=1 | 8.05 | 17.09 | 2.12 | 1.09 |
| sp\|Q99720-2\|SGMR1_HUMAN Isoform 2 of Sigma non-opioid intracellular receptor 1 OS=Homo sapiens OX=9606 GN=SIGMAR1 | 8.05 | 18.48 | 2.30 | 1.20 |
| sp\|Q9BV57\|MTND_HUMAN 1,2-dihydroxy-3-keto-5-methylthiopentene dioxygenase OS=Homo sapiens OX=9606 GN=ADI1 PE=1 SV=1 | 8.04 | 11.55 | 1.44 | 0.52 |
| sp\|P50336\|PPOX_HUMAN Protoporphyrinogen oxidase OS=Homo sapiens OX=9606 GN=PPOX PE=1 SV=1 | 8.03 | 1.82 | 0.23 | -2.14 |
| sp\|Q6IBS0\|TWF2_HUMAN Twinfilin-2 OS=Homo sapiens OX=9606 GN=TWF2 PE=1 SV=2 | 8.03 | 13.28 | 1.65 | 0.73 |
| tr\|F5H442\|F5H442_HUMAN Tumor susceptibility gene 101 protein OS=Homo sapiens OX=9606 GN=TSG101 PE=1 SV=1 | 7.99 | 11.39 | 1.43 | 0.51 |
| tr\|A0A096LPJ3\|A0A096LPJ3_HUMAN COP9 signalosome complex subunit 1 OS=Homo sapiens OX=9606 GN=GPS1 PE=1 SV=1 | 7.91 | 15.99 | 2.02 | 1.02 |
| sp\|Q9Y3B4\|SF3B6_HUMAN Splicing factor 3B subunit 6 OS=Homo sapiens OX=9606 GN=SF3B6 PE=1 SV=1 | 7.87 | 15.64 | 1.99 | 0.99 |
| sp\|Q15050\|RRS1_HUMAN Ribosome biogenesis regulatory protein homolog OS=Homo sapiens OX=9606 GN=RRS1 PE=1 SV=2 | 7.87 | 7.24 | 0.92 | -0.12 |
| tr\|B4DKY1\|B4DKY1_HUMAN Cysteine--tRNA ligase, cytoplasmic OS=Homo sapiens OX=9606 GN=CARS PE=1 SV=1 | 7.83 | 23.93 | 3.06 | 1.61 |
| tr\|F5H013\|F5H013_HUMAN Small nuclear ribonucleoprotein G OS=Homo sapiens OX=9606 GN=SNRPG PE=1 SV=1 | 7.82 | 11.32 | 1.45 | 0.53 |
| sp\|Q9H3P7\|GCP60_HUMAN Golgi resident protein GCP60 OS=Homo sapiens OX=9606 GN=ACBD3 PE=1 SV=4 | 7.77 | 3.58 | 0.46 | -1.12 |

| | | | | |
|---|---|---|---|---|
| tr\|K4DI93\|K4DI93_HUMAN Cullin 4B, isoform CRA_e OS=Homo sapiens OX=9606 GN=CUL4B PE=1 SV=1 | 7.75 | 21.02 | 2.71 | 1.44 |
| sp\|Q9H9A6\|LRC40_HUMAN Leucine-rich repeat-containing protein 40 OS=Homo sapiens OX=9606 GN=LRRC40 PE=1 SV=1 | 7.74 | 15.58 | 2.01 | 1.01 |
| sp\|Q9NQP4\|PFD4_HUMAN Prefoldin subunit 4 OS=Homo sapiens OX=9606 GN=PFDN4 PE=1 SV=1 | 7.72 | 6.80 | 0.88 | -0.18 |
| sp\|Q9UBB4\|ATX10_HUMAN Ataxin-10 OS=Homo sapiens OX=9606 GN=ATXN10 PE=1 SV=1 | 7.72 | 49.42 | 6.40 | 2.68 |
| sp\|Q16719\|KYNU_HUMAN Kynureninase OS=Homo sapiens OX=9606 GN=KYNU PE=1 SV=1 | 7.71 | 9.35 | 1.21 | 0.28 |
| sp\|Q9Y241-2\|HIG1A_HUMAN Isoform 2 of HIG1 domain family member 1A, mitochondrial OS=Homo sapiens OX=9606 GN=HIGD1A | 7.64 | 13.04 | 1.71 | 0.77 |
| tr\|H0YDD4\|H0YDD4_HUMAN Acetyltransferase component of pyruvate dehydrogenase complex (Fragment) OS=Homo sapiens OX=9606 GN=DLAT PE=1 SV=1 | 7.62 | 14.94 | 1.96 | 0.97 |
| sp\|Q9UHD1-2\|CHRD1_HUMAN Isoform 2 of Cysteine and histidine-rich domain-containing protein 1 OS=Homo sapiens OX=9606 GN=CHORDC1 | 7.62 | 21.11 | 2.77 | 1.47 |
| sp\|P40222\|TXLNA_HUMAN Alpha-taxilin OS=Homo sapiens OX=9606 GN=TXLNA PE=1 SV=3 | 7.62 | 2.31 | 0.30 | -1.72 |
| sp\|Q15006\|EMC2_HUMAN ER membrane protein complex subunit 2 OS=Homo sapiens OX=9606 GN=EMC2 PE=1 SV=1 | 7.54 | 20.74 | 2.75 | 1.46 |
| sp\|Q96PZ0-2\|PUS7_HUMAN Isoform 2 of Pseudouridylate synthase 7 homolog OS=Homo sapiens OX=9606 GN=PUS7 | 7.54 | 6.53 | 0.87 | -0.21 |
| tr\|A0A0A0MSE2\|A0A0A0MSE2_HUMAN Hydroxyacyl-coenzyme A dehydrogenase, mitochondrial OS=Homo sapiens OX=9606 GN=HADH PE=1 SV=2 | 7.53 | 16.24 | 2.16 | 1.11 |
| sp\|P42167\|LAP2B_HUMAN Lamina-associated polypeptide 2, isoforms beta/gamma OS=Homo sapiens OX=9606 GN=TMPO PE=1 SV=2 | 7.51 | 43.75 | 5.83 | 2.54 |
| tr\|H0YG54\|H0YG54_HUMAN Oligoribonuclease, mitochondrial OS=Homo sapiens OX=9606 GN=REXO2 PE=1 SV=2 | 7.49 | 14.93 | 1.99 | 1.00 |
| tr\|A0A2R8Y5A0\|A0A2R8Y5A0_HUMAN Casein kinase II subunit alpha OS=Homo sapiens OX=9606 GN=CSNK2A1 PE=1 SV=1 | 7.46 | 40.49 | 5.43 | 2.44 |
| sp\|Q9NXG2\|THUM1_HUMAN THUMP domain-containing protein 1 OS=Homo sapiens OX=9606 GN=THUMPD1 PE=1 SV=2 | 7.41 | 11.97 | 1.61 | 0.69 |
| sp\|P18031\|PTN1_HUMAN Tyrosine-protein phosphatase non-receptor type 1 OS=Homo sapiens OX=9606 GN=PTPN1 PE=1 SV=1 | 7.37 | 13.11 | 1.78 | 0.83 |

| | | | | |
|---|---|---|---|---|
| tr\|A0A087X020\|A0A087X020_HUMAN Ribosome maturation protein SBDS OS=Homo sapiens OX=9606 GN=SBDS PE=1 SV=1 | 7.36 | 36.01 | 4.89 | 2.29 |
| sp\|Q8IUR0\|TPPC5_HUMAN Trafficking protein particle complex subunit 5 OS=Homo sapiens OX=9606 GN=TRAPPC5 PE=1 SV=1 | 7.35 | 3.94 | 0.54 | -0.90 |
| sp\|O43765\|SGTA_HUMAN Small glutamine-rich tetratricopeptide repeat-containing protein alpha OS=Homo sapiens OX=9606 GN=SGTA PE=1 SV=1 | 7.34 | 22.67 | 3.09 | 1.63 |
| sp\|Q9Y3D6\|FIS1_HUMAN Mitochondrial fission 1 protein OS=Homo sapiens OX=9606 GN=FIS1 PE=1 SV=2 | 7.34 | 14.60 | 1.99 | 0.99 |
| sp\|P55060-3\|XPO2_HUMAN Isoform 3 of Exportin-2 OS=Homo sapiens OX=9606 GN=CSE1L | 7.31 | 0.86 | 0.12 | -3.09 |
| sp\|Q9NS69\|TOM22_HUMAN Mitochondrial import receptor subunit TOM22 homolog OS=Homo sapiens OX=9606 GN=TOMM22 PE=1 SV=3 | 7.31 | 46.43 | 6.35 | 2.67 |
| tr\|A0A087X2I1\|A0A087X2I1_HUMAN 26S proteasome regulatory subunit 10B OS=Homo sapiens OX=9606 GN=PSMC6 PE=1 SV=1 | 7.31 | 46.60 | 6.38 | 2.67 |
| tr\|B2WTI3\|B2WTI3_HUMAN Bifunctional arginine demethylase and lysyl-hydroxylase JMJD6 OS=Homo sapiens OX=9606 GN=JMJD6 PE=1 SV=1 | 7.29 | 11.87 | 1.63 | 0.70 |
| sp\|P47755\|CAZA2_HUMAN F-actin-capping protein subunit alpha-2 OS=Homo sapiens OX=9606 GN=CAPZA2 PE=1 SV=3 | 7.28 | 26.58 | 3.65 | 1.87 |
| sp\|O60884\|DNJA2_HUMAN DnaJ homolog subfamily A member 2 OS=Homo sapiens OX=9606 GN=DNAJA2 PE=1 SV=1 | 7.25 | 33.99 | 4.69 | 2.23 |
| sp\|P36952\|SPB5_HUMAN Serpin B5 OS=Homo sapiens OX=9606 GN=SERPINB5 PE=1 SV=2 | 7.25 | 54.78 | 7.56 | 2.92 |
| sp\|Q8NBJ5\|GT251_HUMAN Procollagen galactosyltransferase 1 OS=Homo sapiens OX=9606 GN=COLGALT1 PE=1 SV=1 | 7.25 | 8.00 | 1.10 | 0.14 |
| tr\|A0A1W2PR36\|A0A1W2PR36_HUMAN Guanidinoacetate N-methyltransferase OS=Homo sapiens OX=9606 GN=GAMT PE=1 SV=1 | 7.24 | 19.54 | 2.70 | 1.43 |
| sp\|Q14318-2\|FKBP8_HUMAN Isoform 2 of Peptidyl-prolyl cis-trans isomerase FKBP8 OS=Homo sapiens OX=9606 GN=FKBP8 | 7.22 | 2.58 | 0.36 | -1.49 |
| sp\|Q9Y570-4\|PPME1_HUMAN Isoform 4 of Protein phosphatase methylesterase 1 OS=Homo sapiens OX=9606 GN=PPME1 | 7.18 | 13.74 | 1.91 | 0.94 |
| sp\|Q9UDW1\|QCR9_HUMAN Cytochrome b-c1 complex subunit 9 OS=Homo sapiens OX=9606 GN=UQCR10 PE=1 SV=3 | 7.18 | 24.18 | 3.37 | 1.75 |

| | | | | |
|---|---|---|---|---|
| sp\|Q8NE86-3\|MCU_HUMAN Isoform 3 of Calcium uniporter protein, mitochondrial OS=Homo sapiens OX=9606 GN=MCU | 7.16 | 18.48 | 2.58 | 1.37 |
| sp\|P49321-2\|NASP_HUMAN Isoform 2 of Nuclear autoantigenic sperm protein OS=Homo sapiens OX=9606 GN=NASP | 7.16 | 19.71 | 2.75 | 1.46 |
| sp\|O75436\|VP26A_HUMAN Vacuolar protein sorting-associated protein 26A OS=Homo sapiens OX=9606 GN=VPS26A PE=1 SV=2 | 7.13 | 17.05 | 2.39 | 1.26 |
| sp\|Q9BZD4\|NUF2_HUMAN Kinetochore protein Nuf2 OS=Homo sapiens OX=9606 GN=NUF2 PE=1 SV=2 | 7.13 | 6.75 | 0.95 | -0.08 |
| sp\|P50552\|VASP_HUMAN Vasodilator-stimulated phosphoprotein OS=Homo sapiens OX=9606 GN=VASP PE=1 SV=3 | 7.10 | 7.46 | 1.05 | 0.07 |
| sp\|P28838-2\|AMPL_HUMAN Isoform 2 of Cytosol aminopeptidase OS=Homo sapiens OX=9606 GN=LAP3 | 7.10 | 13.05 | 1.84 | 0.88 |
| sp\|O43252\|PAPS1_HUMAN Bifunctional 3'-phosphoadenosine 5'-phosphosulfate synthase 1 OS=Homo sapiens OX=9606 GN=PAPSS1 PE=1 SV=2 | 7.08 | 24.64 | 3.48 | 1.80 |
| sp\|Q9H9B4\|SFXN1_HUMAN Sideroflexin-1 OS=Homo sapiens OX=9606 GN=SFXN1 PE=1 SV=4 | 7.07 | 5.37 | 0.76 | -0.40 |
| sp\|Q14116\|IL18_HUMAN Interleukin-18 OS=Homo sapiens OX=9606 GN=IL18 PE=1 SV=1 | 7.03 | 29.57 | 4.20 | 2.07 |
| tr\|F8WJN3\|F8WJN3_HUMAN Cleavage and polyadenylation-specificity factor subunit 6 OS=Homo sapiens OX=9606 GN=CPSF6 PE=1 SV=1 | 7.00 | 27.40 | 3.91 | 1.97 |
| sp\|Q5TIE3-2\|VW5B1_HUMAN Isoform 2 of von Willebrand factor A domain-containing protein 5B1 OS=Homo sapiens OX=9606 GN=VWA5B1 | 7.00 | 12.65 | 1.81 | 0.85 |
| sp\|Q9Y3C1\|NOP16_HUMAN Nucleolar protein 16 OS=Homo sapiens OX=9606 GN=NOP16 PE=1 SV=2 | 6.98 | 14.56 | 2.08 | 1.06 |
| tr\|A0A0A0MSG2\|A0A0A0MSG2_HUMAN Four and a half LIM domains protein 2 OS=Homo sapiens OX=9606 GN=FHL2 PE=1 SV=1 | 6.98 | 4.75 | 0.68 | -0.56 |
| tr\|J3KT68\|J3KT68_HUMAN Sigma intracellular receptor 2 OS=Homo sapiens OX=9606 GN=TMEM97 PE=1 SV=1 | 6.98 | 16.36 | 2.34 | 1.23 |
| sp\|P04818-2\|TYSY_HUMAN Isoform 2 of Thymidylate synthase OS=Homo sapiens OX=9606 GN=TYMS | 6.97 | 13.36 | 1.92 | 0.94 |
| sp\|Q8WY22\|BRI3B_HUMAN BRI3-binding protein OS=Homo sapiens OX=9606 GN=BRI3BP PE=1 SV=1 | 6.93 | 20.99 | 3.03 | 1.60 |
| sp\|Q5RI15-2\|COX20_HUMAN Isoform 2 of Cytochrome c oxidase assembly protein | 6.93 | 15.86 | 2.29 | 1.19 |

| | | | | |
|---|---|---|---|---|
| COX20, mitochondrial OS=Homo sapiens OX=9606 GN=COX20 | | | | |
| tr\|H0UI06\|H0UI06_HUMAN Cytochrome c oxidase subunit 7A2, mitochondrial OS=Homo sapiens OX=9606 GN=COX7A2 PE=1 SV=1 | 6.93 | 26.38 | 3.81 | 1.93 |
| sp\|O43172\|PRP4_HUMAN U4/U6 small nuclear ribonucleoprotein Prp4 OS=Homo sapiens OX=9606 GN=PRPF4 PE=1 SV=2 | 6.90 | 18.25 | 2.64 | 1.40 |
| sp\|Q9BTT0\|AN32E_HUMAN Acidic leucine-rich nuclear phosphoprotein 32 family member E OS=Homo sapiens OX=9606 GN=ANP32E PE=1 SV=1 | 6.89 | 47.29 | 6.86 | 2.78 |
| tr\|A6NP24\|A6NP24_HUMAN Quinone oxidoreductase (Fragment) OS=Homo sapiens OX=9606 GN=CRYZ PE=1 SV=1 | 6.86 | 27.97 | 4.08 | 2.03 |
| sp\|Q9BVP2-2\|GNL3_HUMAN Isoform 2 of Guanine nucleotide-binding protein-like 3 OS=Homo sapiens OX=9606 GN=GNL3 | 6.84 | 11.02 | 1.61 | 0.69 |
| sp\|Q9NZI8\|IF2B1_HUMAN Insulin-like growth factor 2 mRNA-binding protein 1 OS=Homo sapiens OX=9606 GN=IGF2BP1 PE=1 SV=2 | 6.84 | 27.86 | 4.07 | 2.03 |
| tr\|A0A2R8Y430\|A0A2R8Y430_HUMAN Glutathione synthetase OS=Homo sapiens OX=9606 GN=GSS PE=1 SV=1 | 6.82 | 12.29 | 1.80 | 0.85 |
| sp\|Q969Z0\|FAKD4_HUMAN FAST kinase domain-containing protein 4 OS=Homo sapiens OX=9606 GN=TBRG4 PE=1 SV=1 | 6.81 | 13.94 | 2.04 | 1.03 |
| tr\|A0A0A0MSV9\|A0A0A0MSV9_HUMAN Tapasin OS=Homo sapiens OX=9606 GN=TAPBP PE=1 SV=1 | 6.81 | 13.76 | 2.02 | 1.02 |
| tr\|K7EM18\|K7EM18_HUMAN Eukaryotic translation initiation factor 1 OS=Homo sapiens OX=9606 GN=EIF1 PE=1 SV=1 | 6.81 | 24.14 | 3.55 | 1.83 |
| tr\|F5H669\|F5H669_HUMAN Cleavage and polyadenylation-specificity factor subunit 7 (Fragment) OS=Homo sapiens OX=9606 GN=CPSF7 PE=1 SV=1 | 6.77 | 12.22 | 1.80 | 0.85 |
| sp\|P53597\|SUCA_HUMAN Succinate--CoA ligase [ADP/GDP-forming] subunit alpha, mitochondrial OS=Homo sapiens OX=9606 GN=SUCLG1 PE=1 SV=4 | 6.70 | 10.85 | 1.62 | 0.70 |
| tr\|A0A087X1B2\|A0A087X1B2_HUMAN U4/U6.U5 tri-snRNP-associated protein 2 OS=Homo sapiens OX=9606 GN=USP39 PE=1 SV=1 | 6.69 | 16.95 | 2.53 | 1.34 |
| sp\|Q96M27-2\|PRRC1_HUMAN Isoform 2 of Protein PRRC1 OS=Homo sapiens OX=9606 GN=PRRC1 | 6.68 | 12.74 | 1.91 | 0.93 |
| tr\|H7C174\|H7C174_HUMAN Hepatocyte growth factor receptor (Fragment) OS=Homo sapiens OX=9606 GN=MET PE=1 SV=1 | 6.64 | 46.01 | 6.93 | 2.79 |
| sp\|Q96CS3\|FAF2_HUMAN FAS-associated factor 2 OS=Homo sapiens OX=9606 GN=FAF2 PE=1 SV=2 | 6.64 | 12.27 | 1.85 | 0.89 |

| | | | | |
|---|---|---|---|---|
| sp\|P55769\|NH2L1_HUMAN NHP2-like protein 1 OS=Homo sapiens OX=9606 GN=SNU13 PE=1 SV=3 | 6.64 | 18.37 | 2.77 | 1.47 |
| sp\|Q9BYT8\|NEUL_HUMAN Neurolysin, mitochondrial OS=Homo sapiens OX=9606 GN=NLN PE=1 SV=1 | 6.62 | 14.66 | 2.21 | 1.15 |
| tr\|A0A0C4DGA2\|A0A0C4DGA2_HUMAN Enoyl-CoA delta isomerase 2, mitochondrial OS=Homo sapiens OX=9606 GN=ECI2 PE=1 SV=1 | 6.61 | 49.53 | 7.49 | 2.90 |
| sp\|P61201\|CSN2_HUMAN COP9 signalosome complex subunit 2 OS=Homo sapiens OX=9606 GN=COPS2 PE=1 SV=1 | 6.61 | 15.25 | 2.31 | 1.21 |
| sp\|Q08209-2\|PP2BA_HUMAN Isoform 2 of Serine/threonine-protein phosphatase 2B catalytic subunit alpha isoform OS=Homo sapiens OX=9606 GN=PPP3CA | 6.60 | 14.30 | 2.17 | 1.12 |
| sp\|O00560-2\|SDCB1_HUMAN Isoform 2 of Syntenin-1 OS=Homo sapiens OX=9606 GN=SDCBP | 6.58 | 36.16 | 5.50 | 2.46 |
| tr\|R4GMN1\|R4GMN1_HUMAN Motile sperm domain-containing protein 2 OS=Homo sapiens OX=9606 GN=MOSPD2 PE=1 SV=1 | 6.55 | 8.33 | 1.27 | 0.35 |
| sp\|O75638-2\|CTAG2_HUMAN Isoform LAGE-1A of Cancer/testis antigen 2 OS=Homo sapiens OX=9606 GN=CTAG2 | 6.53 | 16.45 | 2.52 | 1.33 |
| sp\|Q6P587-2\|FAHD1_HUMAN Isoform 2 of Acylpyruvase FAHD1, mitochondrial OS=Homo sapiens OX=9606 GN=FAHD1 | 6.53 | 16.18 | 2.48 | 1.31 |
| tr\|C9JIZ6\|C9JIZ6_HUMAN Prosaposin OS=Homo sapiens OX=9606 GN=PSAP PE=1 SV=2 | 6.52 | 18.23 | 2.79 | 1.48 |
| sp\|O43681\|ASNA_HUMAN ATPase ASNA1 OS=Homo sapiens OX=9606 GN=ASNA1 PE=1 SV=2 | 6.50 | 26.62 | 4.09 | 2.03 |
| tr\|A0A0R4J2E8\|A0A0R4J2E8_HUMAN Matrin-3 OS=Homo sapiens OX=9606 GN=MATR3 PE=1 SV=1 | 6.46 | 16.00 | 2.48 | 1.31 |
| sp\|P04844-2\|RPN2_HUMAN Isoform 2 of Dolichyl-diphosphooligosaccharide--protein glycosyltransferase subunit 2 OS=Homo sapiens OX=9606 GN=RPN2 | 6.42 | 10.46 | 1.63 | 0.71 |
| sp\|P52888\|THOP1_HUMAN Thimet oligopeptidase OS=Homo sapiens OX=9606 GN=THOP1 PE=1 SV=2 | 6.41 | 22.36 | 3.49 | 1.80 |
| sp\|P11166\|GTR1_HUMAN Solute carrier family 2, facilitated glucose transporter member 1 OS=Homo sapiens OX=9606 GN=SLC2A1 PE=1 SV=2 | 6.40 | 0.59 | 0.09 | -3.44 |
| sp\|O75937\|DNJC8_HUMAN DnaJ homolog subfamily C member 8 OS=Homo sapiens OX=9606 GN=DNAJC8 PE=1 SV=2 | 6.40 | 10.84 | 1.69 | 0.76 |
| tr\|H0Y5K5\|H0Y5K5_HUMAN Endoplasmic reticulum-Golgi intermediate compartment | 6.36 | 11.30 | 1.78 | 0.83 |

| | | | | |
|---|---|---|---|---|
| protein 3 (Fragment) OS=Homo sapiens OX=9606 GN=ERGIC3 PE=1 SV=1 | | | | |
| sp\|Q5M9N0\|CD158_HUMAN Coiled-coil domain-containing protein 158 OS=Homo sapiens OX=9606 GN=CCDC158 PE=2 SV=2 | 6.35 | 35.42 | 5.58 | 2.48 |
| tr\|A0A286YFF7\|A0A286YFF7_HUMAN Palmitoyl-protein thioesterase 1 OS=Homo sapiens OX=9606 GN=PPT1 PE=1 SV=1 | 6.33 | 19.63 | 3.10 | 1.63 |
| sp\|Q9UNS2\|CSN3_HUMAN COP9 signalosome complex subunit 3 OS=Homo sapiens OX=9606 GN=COPS3 PE=1 SV=3 | 6.27 | 2.27 | 0.36 | -1.47 |
| sp\|Q9P287-2\|BCCIP_HUMAN Isoform 2 of BRCA2 and CDKN1A-interacting protein OS=Homo sapiens OX=9606 GN=BCCIP | 6.26 | 16.92 | 2.71 | 1.44 |
| sp\|Q02809-2\|PLOD1_HUMAN Isoform 2 of Procollagen-lysine,2-oxoglutarate 5-dioxygenase 1 OS=Homo sapiens OX=9606 GN=PLOD1 | 6.24 | 16.06 | 2.57 | 1.36 |
| sp\|P46108-2\|CRK_HUMAN Isoform Crk-I of Adapter molecule crk OS=Homo sapiens OX=9606 GN=CRK | 6.24 | 11.96 | 1.92 | 0.94 |
| tr\|C9JGI3\|C9JGI3_HUMAN Thymidine phosphorylase (Fragment) OS=Homo sapiens OX=9606 GN=TYMP PE=1 SV=1 | 6.21 | 26.91 | 4.33 | 2.12 |
| tr\|B4DLN1\|B4DLN1_HUMAN cDNA FLJ60124, highly similar to Mitochondrial dicarboxylate carrier OS=Homo sapiens OX=9606 PE=2 SV=1 | 6.19 | 7.25 | 1.17 | 0.23 |
| sp\|O43395\|PRPF3_HUMAN U4/U6 small nuclear ribonucleoprotein Prp3 OS=Homo sapiens OX=9606 GN=PRPF3 PE=1 SV=2 | 6.16 | 14.40 | 2.34 | 1.23 |
| sp\|P13473-2\|LAMP2_HUMAN Isoform LAMP-2B of Lysosome-associated membrane glycoprotein 2 OS=Homo sapiens OX=9606 GN=LAMP2 | 6.15 | 9.86 | 1.60 | 0.68 |
| sp\|O60568\|PLOD3_HUMAN Multifunctional procollagen lysine hydroxylase and glycosyltransferase LH3 OS=Homo sapiens OX=9606 GN=PLOD3 PE=1 SV=1 | 6.15 | 11.77 | 1.92 | 0.94 |
| sp\|Q9UBQ0\|VPS29_HUMAN Vacuolar protein sorting-associated protein 29 OS=Homo sapiens OX=9606 GN=VPS29 PE=1 SV=1 | 6.11 | 9.67 | 1.58 | 0.66 |
| sp\|P62993\|GRB2_HUMAN Growth factor receptor-bound protein 2 OS=Homo sapiens OX=9606 GN=GRB2 PE=1 SV=1 | 6.07 | 12.49 | 2.06 | 1.04 |
| tr\|A0A2R8Y891\|A0A2R8Y891_HUMAN ATP-dependent 6-phosphofructokinase OS=Homo sapiens OX=9606 GN=PFKM PE=1 SV=1 | 6.06 | 10.75 | 1.77 | 0.83 |
| sp\|Q14258\|TRI25_HUMAN E3 ubiquitin/ISG15 ligase TRIM25 OS=Homo sapiens OX=9606 GN=TRIM25 PE=1 SV=2 | 6.06 | 2.13 | 0.35 | -1.50 |
| sp\|Q96RP9\|EFGM_HUMAN Elongation factor G, mitochondrial OS=Homo sapiens OX=9606 GN=GFM1 PE=1 SV=2 | 6.03 | 10.98 | 1.82 | 0.86 |

| | | | | |
|---|---|---|---|---|
| tr\|Q5T6H7\|Q5T6H7_HUMAN Xaa-Pro aminopeptidase 1 OS=Homo sapiens OX=9606 GN=XPNPEP1 PE=1 SV=1 | 6.00 | 9.41 | 1.57 | 0.65 |
| tr\|K7EII7\|K7EII7_HUMAN Galactokinase (Fragment) OS=Homo sapiens OX=9606 GN=GALK1 PE=1 SV=1 | 5.99 | 7.40 | 1.24 | 0.31 |
| sp\|Q9NQZ2\|SAS10_HUMAN Something about silencing protein 10 OS=Homo sapiens OX=9606 GN=UTP3 PE=1 SV=1 | 5.97 | 10.09 | 1.69 | 0.76 |
| sp\|Q99436\|PSB7_HUMAN Proteasome subunit beta type-7 OS=Homo sapiens OX=9606 GN=PSMB7 PE=1 SV=1 | 5.94 | 18.54 | 3.12 | 1.64 |
| sp\|Q9Y2R4\|DDX52_HUMAN Probable ATP-dependent RNA helicase DDX52 OS=Homo sapiens OX=9606 GN=DDX52 PE=1 SV=3 | 5.94 | 0.56 | 0.09 | -3.41 |
| sp\|O43242\|PSMD3_HUMAN 26S proteasome non-ATPase regulatory subunit 3 OS=Homo sapiens OX=9606 GN=PSMD3 PE=1 SV=2 | 5.94 | 8.77 | 1.48 | 0.56 |
| tr\|I3L505\|I3L505_HUMAN Acyl carrier protein (Fragment) OS=Homo sapiens OX=9606 GN=NDUFAB1 PE=1 SV=1 | 5.93 | 16.50 | 2.78 | 1.48 |
| tr\|A0A0J9YXF2\|A0A0J9YXF2_HUMAN Paraoxonase 2, isoform CRA_a OS=Homo sapiens OX=9606 GN=PON2 PE=1 SV=1 | 5.91 | 13.24 | 2.24 | 1.16 |
| sp\|Q8WVM8-2\|SCFD1_HUMAN Isoform 2 of Sec1 family domain-containing protein 1 OS=Homo sapiens OX=9606 GN=SCFD1 | 5.91 | 4.46 | 0.75 | -0.41 |
| sp\|Q9Y4P3\|TBL2_HUMAN Transducin beta-like protein 2 OS=Homo sapiens OX=9606 GN=TBL2 PE=1 SV=1 | 5.89 | 15.47 | 2.63 | 1.39 |
| sp\|P14550\|AK1A1_HUMAN Alcohol dehydrogenase [NADP(+)] OS=Homo sapiens OX=9606 GN=AKR1A1 PE=1 SV=3 | 5.86 | 29.73 | 5.07 | 2.34 |
| sp\|Q96FQ6\|S10AG_HUMAN Protein S100-A16 OS=Homo sapiens OX=9606 GN=S100A16 PE=1 SV=1 | 5.85 | 7.98 | 1.36 | 0.45 |
| tr\|A0A2R8Y478\|A0A2R8Y478_HUMAN Tetraspanin OS=Homo sapiens OX=9606 GN=CD9 PE=1 SV=1 | 5.85 | 8.44 | 1.44 | 0.53 |
| sp\|P07099\|HYEP_HUMAN Epoxide hydrolase 1 OS=Homo sapiens OX=9606 GN=EPHX1 PE=1 SV=1 | 5.83 | 3.44 | 0.59 | -0.76 |
| sp\|O94776\|MTA2_HUMAN Metastasis-associated protein MTA2 OS=Homo sapiens OX=9606 GN=MTA2 PE=1 SV=1 | 5.82 | 14.86 | 2.55 | 1.35 |
| sp\|O15173-2\|PGRC2_HUMAN Isoform 2 of Membrane-associated progesterone receptor component 2 OS=Homo sapiens OX=9606 GN=PGRMC2 | 5.80 | 20.34 | 3.50 | 1.81 |
| tr\|J3QLR8\|J3QLR8_HUMAN 28S ribosomal protein S23, mitochondrial OS=Homo sapiens OX=9606 GN=MRPS23 PE=1 SV=1 | 5.79 | 16.71 | 2.88 | 1.53 |

| | | | | |
|---|---|---|---|---|
| sp\|Q6DKJ4\|NXN_HUMAN Nucleoredoxin OS=Homo sapiens OX=9606 GN=NXN PE=1 SV=2 | 5.77 | 8.18 | 1.42 | 0.50 |
| sp\|Q08380\|LG3BP_HUMAN Galectin-3-binding protein OS=Homo sapiens OX=9606 GN=LGALS3BP PE=1 SV=1 | 5.77 | 15.08 | 2.61 | 1.39 |
| tr\|B5MCF9\|B5MCF9_HUMAN Pescadillo homolog OS=Homo sapiens OX=9606 GN=PES1 PE=1 SV=1 | 5.74 | 6.35 | 1.11 | 0.15 |
| sp\|Q9Y2S7\|PDIP2_HUMAN Polymerase delta-interacting protein 2 OS=Homo sapiens OX=9606 GN=POLDIP2 PE=1 SV=1 | 5.73 | 7.45 | 1.30 | 0.38 |
| sp\|P30740\|ILEU_HUMAN Leukocyte elastase inhibitor OS=Homo sapiens OX=9606 GN=SERPINB1 PE=1 SV=1 | 5.72 | 16.38 | 2.86 | 1.52 |
| sp\|Q15067-2\|ACOX1_HUMAN Isoform 2 of Peroxisomal acyl-coenzyme A oxidase 1 OS=Homo sapiens OX=9606 GN=ACOX1 | 5.69 | 7.72 | 1.35 | 0.44 |
| sp\|O00743-2\|PPP6_HUMAN Isoform 2 of Serine/threonine-protein phosphatase 6 catalytic subunit OS=Homo sapiens OX=9606 GN=PPP6C | 5.69 | 15.70 | 2.76 | 1.46 |
| tr\|M0R389\|M0R389_HUMAN Platelet-activating factor acetylhydrolase IB subunit gamma (Fragment) OS=Homo sapiens OX=9606 GN=PAFAH1B3 PE=1 SV=8 | 5.68 | 16.72 | 2.94 | 1.56 |
| tr\|H7BYN3\|H7BYN3_HUMAN Transcription factor A, mitochondrial (Fragment) OS=Homo sapiens OX=9606 GN=TFAM PE=1 SV=1 | 5.67 | 20.73 | 3.66 | 1.87 |
| tr\|F6VRR5\|F6VRR5_HUMAN Polymerase delta-interacting protein 3 OS=Homo sapiens OX=9606 GN=POLDIP3 PE=1 SV=1 | 5.63 | 11.09 | 1.97 | 0.98 |
| tr\|Q5VZU3\|Q5VZU3_HUMAN RNA 3'-terminal phosphate cyclase-like protein OS=Homo sapiens OX=9606 GN=RCL1 PE=1 SV=2 | 5.60 | 10.04 | 1.79 | 0.84 |
| sp\|Q9UBK8-2\|MTRR_HUMAN Isoform B of Methionine synthase reductase OS=Homo sapiens OX=9606 GN=MTRR | 5.59 | 9.74 | 1.74 | 0.80 |
| sp\|Q9BT73\|PSMG3_HUMAN Proteasome assembly chaperone 3 OS=Homo sapiens OX=9606 GN=PSMG3 PE=1 SV=1 | 5.58 | 18.91 | 3.39 | 1.76 |
| sp\|P61011\|SRP54_HUMAN Signal recognition particle 54 kDa protein OS=Homo sapiens OX=9606 GN=SRP54 PE=1 SV=1 | 5.53 | 1.92 | 0.35 | -1.52 |
| sp\|Q9NTX5-2\|ECHD1_HUMAN Isoform 2 of Ethylmalonyl-CoA decarboxylase OS=Homo sapiens OX=9606 GN=ECHDC1 | 5.53 | 9.87 | 1.79 | 0.84 |
| sp\|Q8TCS8\|PNPT1_HUMAN Polyribonucleotide nucleotidyltransferase 1, mitochondrial OS=Homo sapiens OX=9606 GN=PNPT1 PE=1 SV=2 | 5.50 | 12.29 | 2.23 | 1.16 |
| sp\|Q96A33-2\|CCD47_HUMAN Isoform 2 of Coiled-coil domain-containing protein 47 OS=Homo sapiens OX=9606 GN=CCDC47 | 5.50 | 1.80 | 0.33 | -1.61 |

| | | | | |
|---|---|---|---|---|
| sp\|Q13409-2\|DC1I2_HUMAN Isoform 2B of Cytoplasmic dynein 1 intermediate chain 2 OS=Homo sapiens OX=9606 GN=DYNC1I2 | 5.49 | 13.01 | 2.37 | 1.24 |
| tr\|K7EQG1\|K7EQG1_HUMAN Glutaminyl-peptide cyclotransferase-like protein OS=Homo sapiens OX=9606 GN=QPCTL PE=1 SV=1 | 5.49 | 3.65 | 0.67 | -0.59 |
| tr\|C9JP16\|C9JP16_HUMAN Cartilage-associated protein OS=Homo sapiens OX=9606 GN=CRTAP PE=1 SV=1 | 5.49 | 10.13 | 1.85 | 0.88 |
| sp\|P11177-2\|ODPB_HUMAN Isoform 2 of Pyruvate dehydrogenase E1 component subunit beta, mitochondrial OS=Homo sapiens OX=9606 GN=PDHB | 5.48 | 14.66 | 2.67 | 1.42 |
| tr\|B1AP13\|B1AP13_HUMAN Complement decay-accelerating factor OS=Homo sapiens OX=9606 GN=CD55 PE=1 SV=1 | 5.46 | 13.73 | 2.51 | 1.33 |
| sp\|O76071\|CIAO1_HUMAN Probable cytosolic iron-sulfur protein assembly protein CIAO1 OS=Homo sapiens OX=9606 GN=CIAO1 PE=1 SV=1 | 5.44 | 9.83 | 1.81 | 0.85 |
| tr\|C9J1E7\|C9J1E7_HUMAN AP-1 complex subunit beta-1 (Fragment) OS=Homo sapiens OX=9606 GN=AP1B1 PE=1 SV=1 | 5.42 | 13.84 | 2.55 | 1.35 |
| tr\|H0YDU8\|H0YDU8_HUMAN Serine/threonine-protein phosphatase (Fragment) OS=Homo sapiens OX=9606 GN=PPP5C PE=1 SV=1 | 5.39 | 15.47 | 2.87 | 1.52 |
| sp\|O15269\|SPTC1_HUMAN Serine palmitoyltransferase 1 OS=Homo sapiens OX=9606 GN=SPTLC1 PE=1 SV=1 | 5.37 | 9.68 | 1.80 | 0.85 |
| sp\|Q9UL25\|RAB21_HUMAN Ras-related protein Rab-21 OS=Homo sapiens OX=9606 GN=RAB21 PE=1 SV=3 | 5.33 | 14.78 | 2.77 | 1.47 |
| tr\|A0A087X0R6\|A0A087X0R6_HUMAN Sorting nexin-12 OS=Homo sapiens OX=9606 GN=SNX12 PE=1 SV=1 | 5.32 | 3.35 | 0.63 | -0.67 |
| sp\|Q9P0L0-2\|VAPA_HUMAN Isoform 2 of Vesicle-associated membrane protein-associated protein A OS=Homo sapiens OX=9606 GN=VAPA | 5.32 | 11.28 | 2.12 | 1.09 |
| sp\|Q9H2U2-2\|IPYR2_HUMAN Isoform 2 of Inorganic pyrophosphatase 2, mitochondrial OS=Homo sapiens OX=9606 GN=PPA2 | 5.31 | 15.19 | 2.86 | 1.52 |
| sp\|Q12792-3\|TWF1_HUMAN Isoform 3 of Twinfilin-1 OS=Homo sapiens OX=9606 GN=TWF1 | 5.31 | 11.52 | 2.17 | 1.12 |
| sp\|Q4G0N4-2\|NAKD2_HUMAN Isoform 2 of NAD kinase 2, mitochondrial OS=Homo sapiens OX=9606 GN=NADK2 | 5.28 | 11.86 | 2.25 | 1.17 |
| sp\|Q15024\|EXOS7_HUMAN Exosome complex component RRP42 OS=Homo sapiens OX=9606 GN=EXOSC7 PE=1 SV=3 | 5.28 | 4.55 | 0.86 | -0.21 |

| | | | | |
|---|---|---|---|---|
| tr\|J3QRD1\|J3QRD1_HUMAN Fatty aldehyde dehydrogenase OS=Homo sapiens OX=9606 GN=ALDH3A2 PE=1 SV=1 | 5.26 | 4.14 | 0.79 | -0.35 |
| tr\|B0QZ18\|B0QZ18_HUMAN Copine-1 OS=Homo sapiens OX=9606 GN=CPNE1 PE=1 SV=1 | 5.26 | 25.45 | 4.84 | 2.27 |
| tr\|G5EA06\|G5EA06_HUMAN 28S ribosomal protein S27, mitochondrial OS=Homo sapiens OX=9606 GN=MRPS27 PE=1 SV=1 | 5.21 | 9.06 | 1.74 | 0.80 |
| sp\|Q96HY6\|DDRGK_HUMAN DDRGK domain-containing protein 1 OS=Homo sapiens OX=9606 GN=DDRGK1 PE=1 SV=2 | 5.19 | 6.60 | 1.27 | 0.35 |
| tr\|H7C2S1\|H7C2S1_HUMAN Guanine nucleotide exchange factor DBS (Fragment) OS=Homo sapiens OX=9606 GN=MCF2L PE=1 SV=1 | 5.19 | 16.53 | 3.19 | 1.67 |
| tr\|Q5T9B7\|Q5T9B7_HUMAN Adenylate kinase isoenzyme 1 OS=Homo sapiens OX=9606 GN=AK1 PE=1 SV=1 | 5.18 | 21.07 | 4.06 | 2.02 |
| tr\|G5E977\|G5E977_HUMAN Nicotinate phosphoribosyltransferase domain containing 1, isoform CRA_d OS=Homo sapiens OX=9606 GN=NAPRT PE=1 SV=1 | 5.17 | 12.38 | 2.39 | 1.26 |
| tr\|A0A087WWF6\|A0A087WWF6_HUMAN DNA polymerase delta subunit 2 OS=Homo sapiens OX=9606 GN=POLD2 PE=1 SV=1 | 5.16 | 10.49 | 2.03 | 1.02 |
| sp\|O95573\|ACSL3_HUMAN Long-chain-fatty-acid--CoA ligase 3 OS=Homo sapiens OX=9606 GN=ACSL3 PE=1 SV=3 | 5.15 | 1.79 | 0.35 | -1.52 |
| tr\|H0Y512\|H0Y512_HUMAN Adipocyte plasma membrane-associated protein (Fragment) OS=Homo sapiens OX=9606 GN=APMAP PE=1 SV=1 | 5.15 | 9.10 | 1.77 | 0.82 |
| sp\|P63151-2\|2ABA_HUMAN Isoform 2 of Serine/threonine-protein phosphatase 2A 55 kDa regulatory subunit B alpha isoform OS=Homo sapiens OX=9606 GN=PPP2R2A | 5.14 | 7.49 | 1.46 | 0.54 |
| sp\|Q9UGI8-2\|TES_HUMAN Isoform 2 of Testin OS=Homo sapiens OX=9606 GN=TES | 5.14 | 5.94 | 1.16 | 0.21 |
| sp\|P35270\|SPRE_HUMAN Sepiapterin reductase OS=Homo sapiens OX=9606 GN=SPR PE=1 SV=1 | 5.13 | 19.03 | 3.71 | 1.89 |
| tr\|A0A0A0MRG8\|A0A0A0MRG8_HUMAN Bcl-2 homologous antagonist/killer OS=Homo sapiens OX=9606 GN=BAK1 PE=1 SV=1 | 5.12 | 4.98 | 0.97 | -0.04 |
| tr\|K7EMW4\|K7EMW4_HUMAN Nicalin homolog (Zebrafish), isoform CRA_e OS=Homo sapiens OX=9606 GN=NCLN PE=1 SV=1 | 5.06 | 11.87 | 2.35 | 1.23 |
| tr\|B4DJV5\|B4DJV5_HUMAN cDNA FLJ51513, highly similar to Periodic tryptophan protein 1 homolog OS=Homo sapiens OX=9606 GN=PWP1 PE=1 SV=1 | 5.05 | 9.42 | 1.87 | 0.90 |

| | | | | |
|---|---|---|---|---|
| tr\|A0A087WX59\|A0A087WX59_HUMAN Proteasomal ubiquitin receptor ADRM1 OS=Homo sapiens OX=9606 GN=ADRM1 PE=1 SV=1 | 5.04 | 9.46 | 1.88 | 0.91 |
| tr\|G3V192\|G3V192_HUMAN Ferritin OS=Homo sapiens OX=9606 GN=FTH1 PE=1 SV=1 | 5.04 | 25.50 | 5.06 | 2.34 |
| sp\|Q6IAA8\|LTOR1_HUMAN Ragulator complex protein LAMTOR1 OS=Homo sapiens OX=9606 GN=LAMTOR1 PE=1 SV=2 | 5.03 | 21.61 | 4.29 | 2.10 |
| tr\|A0A087WUD3\|A0A087WUD3_HUMAN Oligosaccharyltransferase complex subunit OSTC OS=Homo sapiens OX=9606 GN=OSTC PE=1 SV=1 | 5.03 | 8.41 | 1.67 | 0.74 |
| tr\|F8W733\|F8W733_HUMAN BRISC and BRCA1-A complex member 2 (Fragment) OS=Homo sapiens OX=9606 GN=BABAM2 PE=1 SV=1 | 5.03 | 8.59 | 1.71 | 0.77 |
| sp\|Q9Y3B7-3\|RM11_HUMAN Isoform 3 of 39S ribosomal protein L11, mitochondrial OS=Homo sapiens OX=9606 GN=MRPL11 | 5.02 | 15.63 | 3.11 | 1.64 |
| sp\|Q96TA1-2\|NIBL1_HUMAN Isoform 2 of Niban-like protein 1 OS=Homo sapiens OX=9606 GN=FAM129B | 5.00 | 7.51 | 1.50 | 0.59 |
| sp\|Q96G03\|PGM2_HUMAN Phosphoglucomutase-2 OS=Homo sapiens OX=9606 GN=PGM2 PE=1 SV=4 | 5.00 | 1.42 | 0.28 | -1.81 |
| tr\|A0A0A6YYL4\|A0A0A6YYL4_HUMAN Coronin OS=Homo sapiens OX=9606 GN=CORO7-PAM16 PE=3 SV=1 | 4.97 | 26.75 | 5.38 | 2.43 |
| sp\|Q15397\|PUM3_HUMAN Pumilio homolog 3 OS=Homo sapiens OX=9606 GN=PUM3 PE=1 SV=3 | 4.95 | 3.97 | 0.80 | -0.32 |
| tr\|H0YDR3\|H0YDR3_HUMAN Tetratricopeptide repeat protein 9C (Fragment) OS=Homo sapiens OX=9606 GN=TTC9C PE=1 SV=1 | 4.94 | 19.95 | 4.04 | 2.01 |
| tr\|A0A087WSV8\|A0A087WSV8_HUMAN Nucleobindin 2, isoform CRA_b OS=Homo sapiens OX=9606 GN=NUCB2 PE=1 SV=1 | 4.94 | 5.03 | 1.02 | 0.03 |
| sp\|P48147\|PPCE_HUMAN Prolyl endopeptidase OS=Homo sapiens OX=9606 GN=PREP PE=1 SV=2 | 4.90 | 6.66 | 1.36 | 0.44 |
| sp\|O14744\|ANM5_HUMAN Protein arginine N-methyltransferase 5 OS=Homo sapiens OX=9606 GN=PRMT5 PE=1 SV=4 | 4.87 | 13.64 | 2.80 | 1.49 |
| sp\|O75306-2\|NDUS2_HUMAN Isoform 2 of NADH dehydrogenase [ubiquinone] iron-sulfur protein 2, mitochondrial OS=Homo sapiens OX=9606 GN=NDUFS2 | 4.86 | 13.29 | 2.74 | 1.45 |
| sp\|Q13217\|DNJC3_HUMAN DnaJ homolog subfamily C member 3 OS=Homo sapiens OX=9606 GN=DNAJC3 PE=1 SV=1 | 4.82 | 3.03 | 0.63 | -0.67 |
| sp\|Q9Y512\|SAM50_HUMAN Sorting and assembly machinery component 50 homolog | 4.81 | 10.75 | 2.23 | 1.16 |

| | | | | |
|---|---|---|---|---|
| OS=Homo sapiens OX=9606 GN=SAMM50 PE=1 SV=3 | | | | |
| sp\|P05362\|ICAM1_HUMAN Intercellular adhesion molecule 1 OS=Homo sapiens OX=9606 GN=ICAM1 PE=1 SV=2 | 4.75 | 12.42 | 2.61 | 1.39 |
| sp\|Q96HR9-2\|REEP6_HUMAN Isoform 2 of Receptor expression-enhancing protein 6 OS=Homo sapiens OX=9606 GN=REEP6 | 4.74 | 10.51 | 2.22 | 1.15 |
| tr\|J3KQ48\|J3KQ48_HUMAN Peptidyl-tRNA hydrolase 2, mitochondrial OS=Homo sapiens OX=9606 GN=PTRH2 PE=1 SV=1 | 4.74 | 6.21 | 1.31 | 0.39 |
| sp\|Q4VC31\|CCD58_HUMAN Coiled-coil domain-containing protein 58 OS=Homo sapiens OX=9606 GN=CCDC58 PE=1 SV=1 | 4.73 | 6.54 | 1.38 | 0.47 |
| sp\|P43304-2\|GPDM_HUMAN Isoform 2 of Glycerol-3-phosphate dehydrogenase, mitochondrial OS=Homo sapiens OX=9606 GN=GPD2 | 4.71 | 9.99 | 2.12 | 1.08 |
| sp\|P28482\|MK01_HUMAN Mitogen-activated protein kinase 1 OS=Homo sapiens OX=9606 GN=MAPK1 PE=1 SV=3 | 4.69 | 11.82 | 2.52 | 1.33 |
| sp\|Q92600-3\|CNOT9_HUMAN Isoform 3 of CCR4-NOT transcription complex subunit 9 OS=Homo sapiens OX=9606 GN=CNOT9 | 4.65 | 7.90 | 1.70 | 0.76 |
| sp\|P51398-2\|RT29_HUMAN Isoform 2 of 28S ribosomal protein S29, mitochondrial OS=Homo sapiens OX=9606 GN=DAP3 | 4.59 | 4.88 | 1.06 | 0.09 |
| tr\|A0A0A0MRF4\|A0A0A0MRF4_HUMAN 39S ribosomal protein L19, mitochondrial OS=Homo sapiens OX=9606 GN=MRPL19 PE=1 SV=1 | 4.59 | 12.20 | 2.66 | 1.41 |
| tr\|U3KQB5\|U3KQB5_HUMAN Homeobox protein TGIF2 (Fragment) OS=Homo sapiens OX=9606 GN=TGIF2 PE=4 SV=1 | 4.59 | 17.76 | 3.87 | 1.95 |
| sp\|Q96Q11-2\|TRNT1_HUMAN Isoform 2 of CCA tRNA nucleotidyltransferase 1, mitochondrial OS=Homo sapiens OX=9606 GN=TRNT1 | 4.57 | 7.98 | 1.75 | 0.81 |
| sp\|Q9UDY4\|DNJB4_HUMAN DnaJ homolog subfamily B member 4 OS=Homo sapiens OX=9606 GN=DNAJB4 PE=1 SV=1 | 4.56 | 31.88 | 6.99 | 2.80 |
| tr\|E9PLY5\|E9PLY5_HUMAN Microtubule-actin cross-linking factor 1, isoforms 1/2/3/5 (Fragment) OS=Homo sapiens OX=9606 GN=MACF1 PE=1 SV=1 | 4.55 | 19.70 | 4.33 | 2.11 |
| sp\|O76094\|SRP72_HUMAN Signal recognition particle subunit SRP72 OS=Homo sapiens OX=9606 GN=SRP72 PE=1 SV=3 | 4.55 | 22.35 | 4.91 | 2.30 |
| sp\|O75131\|CPNE3_HUMAN Copine-3 OS=Homo sapiens OX=9606 GN=CPNE3 PE=1 SV=1 | 4.55 | 8.00 | 1.76 | 0.81 |
| sp\|Q9H5Q4\|TFB2M_HUMAN Dimethyladenosine transferase 2, mitochondrial OS=Homo sapiens OX=9606 GN=TFB2M PE=1 SV=1 | 4.53 | 8.07 | 1.78 | 0.83 |

| | | | | |
|---|---|---|---|---|
| tr\|H0YDP7\|H0YDP7_HUMAN 39S ribosomal protein L49, mitochondrial (Fragment) OS=Homo sapiens OX=9606 GN=MRPL49 PE=1 SV=1 | 4.52 | 18.92 | 4.19 | 2.07 |
| sp\|Q8NCA5-2\|FA98A_HUMAN Isoform 2 of Protein FAM98A OS=Homo sapiens OX=9606 GN=FAM98A | 4.52 | 8.49 | 1.88 | 0.91 |
| sp\|Q8WU90\|ZC3HF_HUMAN Zinc finger CCCH domain-containing protein 15 OS=Homo sapiens OX=9606 GN=ZC3H15 PE=1 SV=1 | 4.52 | 0.78 | 0.17 | -2.53 |
| tr\|Q5T8A0\|Q5T8A0_HUMAN 28S ribosomal protein S2, mitochondrial (Fragment) OS=Homo sapiens OX=9606 GN=MRPS2 PE=1 SV=1 | 4.51 | 10.67 | 2.36 | 1.24 |
| sp\|Q9HAV7\|GRPE1_HUMAN GrpE protein homolog 1, mitochondrial OS=Homo sapiens OX=9606 GN=GRPEL1 PE=1 SV=2 | 4.51 | 22.20 | 4.93 | 2.30 |
| sp\|Q5JTJ3-2\|COA6_HUMAN Isoform 2 of Cytochrome c oxidase assembly factor 6 homolog OS=Homo sapiens OX=9606 GN=COA6 | 4.51 | 0.65 | 0.14 | -2.80 |
| sp\|Q13561-2\|DCTN2_HUMAN Isoform 2 of Dynactin subunit 2 OS=Homo sapiens OX=9606 GN=DCTN2 | 4.50 | 3.57 | 0.79 | -0.33 |
| sp\|Q12874\|SF3A3_HUMAN Splicing factor 3A subunit 3 OS=Homo sapiens OX=9606 GN=SF3A3 PE=1 SV=1 | 4.50 | 12.98 | 2.89 | 1.53 |
| sp\|Q06203\|PUR1_HUMAN Amidophosphoribosyltransferase OS=Homo sapiens OX=9606 GN=PPAT PE=1 SV=1 | 4.49 | 4.29 | 0.96 | -0.06 |
| tr\|A0A087X2D5\|A0A087X2D5_HUMAN 39S ribosomal protein L45, mitochondrial OS=Homo sapiens OX=9606 GN=MRPL45 PE=1 SV=1 | 4.48 | 16.23 | 3.62 | 1.86 |
| tr\|A0A0A0MT59\|A0A0A0MT59_HUMAN Mortality factor 4-like protein 2 OS=Homo sapiens OX=9606 GN=MORF4L2 PE=1 SV=1 | 4.47 | 6.45 | 1.44 | 0.53 |
| sp\|O00273-2\|DFFA_HUMAN Isoform DFF35 of DNA fragmentation factor subunit alpha OS=Homo sapiens OX=9606 GN=DFFA | 4.43 | 12.55 | 2.83 | 1.50 |
| tr\|A0A087X279\|A0A087X279_HUMAN Interferon-induced protein with tetratricopeptide repeats 2 OS=Homo sapiens OX=9606 GN=IFIT2 PE=1 SV=1 | 4.42 | 6.75 | 1.53 | 0.61 |
| tr\|E9PFR3\|E9PFR3_HUMAN Serine/threonine-protein phosphatase 2A 56 kDa regulatory subunit OS=Homo sapiens OX=9606 GN=PPP2R5D PE=1 SV=1 | 4.40 | 2.34 | 0.53 | -0.91 |
| tr\|A0A1B0GVN9\|A0A1B0GVN9_HUMAN Uroporphyrinogen decarboxylase OS=Homo sapiens OX=9606 GN=UROD PE=1 SV=1 | 4.39 | 11.84 | 2.70 | 1.43 |
| sp\|Q8N183\|NDUF2_HUMAN NADH dehydrogenase [ubiquinone] 1 alpha subcomplex assembly factor 2 OS=Homo sapiens OX=9606 GN=NDUFAF2 PE=1 SV=1 | 4.39 | 15.22 | 3.47 | 1.79 |

| | | | | |
|---|---|---|---|---|
| sp\|P55957-2\|BID_HUMAN Isoform 2 of BH3-interacting domain death agonist OS=Homo sapiens OX=9606 GN=BID | 4.38 | 18.38 | 4.19 | 2.07 |
| tr\|F5GWV0\|F5GWV0_HUMAN Armadillo repeat-containing protein 6 (Fragment) OS=Homo sapiens OX=9606 GN=ARMC6 PE=1 SV=8 | 4.38 | 8.09 | 1.85 | 0.89 |
| sp\|P36405\|ARL3_HUMAN ADP-ribosylation factor-like protein 3 OS=Homo sapiens OX=9606 GN=ARL3 PE=1 SV=2 | 4.37 | 2.31 | 0.53 | -0.92 |
| tr\|C9JJ19\|C9JJ19_HUMAN 28S ribosomal protein S34, mitochondrial OS=Homo sapiens OX=9606 GN=MRPS34 PE=1 SV=2 | 4.36 | 9.82 | 2.25 | 1.17 |
| sp\|Q96IR7\|HPDL_HUMAN 4-hydroxyphenylpyruvate dioxygenase-like protein OS=Homo sapiens OX=9606 GN=HPDL PE=1 SV=1 | 4.36 | 26.58 | 6.10 | 2.61 |
| sp\|Q9Y295\|DRG1_HUMAN Developmentally-regulated GTP-binding protein 1 OS=Homo sapiens OX=9606 GN=DRG1 PE=1 SV=1 | 4.36 | 12.58 | 2.89 | 1.53 |
| sp\|P23258\|TBG1_HUMAN Tubulin gamma-1 chain OS=Homo sapiens OX=9606 GN=TUBG1 PE=1 SV=2 | 4.36 | 10.49 | 2.41 | 1.27 |
| sp\|P19525\|E2AK2_HUMAN Interferon-induced, double-stranded RNA-activated protein kinase OS=Homo sapiens OX=9606 GN=EIF2AK2 PE=1 SV=2 | 4.35 | 4.34 | 1.00 | 0.00 |
| sp\|O95801\|TTC4_HUMAN Tetratricopeptide repeat protein 4 OS=Homo sapiens OX=9606 GN=TTC4 PE=1 SV=3 | 4.34 | 7.35 | 1.69 | 0.76 |
| sp\|Q96KP4\|CNDP2_HUMAN Cytosolic non-specific dipeptidase OS=Homo sapiens OX=9606 GN=CNDP2 PE=1 SV=2 | 4.34 | 25.42 | 5.86 | 2.55 |
| tr\|A0A2R8Y6Y7\|A0A2R8Y6Y7_HUMAN Succinate--CoA ligase [ADP-forming] subunit beta, mitochondrial OS=Homo sapiens OX=9606 GN=SUCLA2 PE=1 SV=1 | 4.33 | 7.54 | 1.74 | 0.80 |
| sp\|Q99598\|TSNAX_HUMAN Translin-associated protein X OS=Homo sapiens OX=9606 GN=TSNAX PE=1 SV=1 | 4.29 | 16.48 | 3.84 | 1.94 |
| sp\|P11802\|CDK4_HUMAN Cyclin-dependent kinase 4 OS=Homo sapiens OX=9606 GN=CDK4 PE=1 SV=2 | 4.29 | 23.35 | 5.44 | 2.44 |
| tr\|B4DJ81\|B4DJ81_HUMAN NADH-ubiquinone oxidoreductase 75 kDa subunit, mitochondrial OS=Homo sapiens OX=9606 GN=NDUFS1 PE=1 SV=1 | 4.29 | 12.86 | 3.00 | 1.58 |
| tr\|B4DXZ6\|B4DXZ6_HUMAN cDNA FLJ58644, highly similar to Fragile X mental retardation syndrome-related protein 1 OS=Homo sapiens OX=9606 GN=FXR1 PE=1 SV=1 | 4.29 | 12.63 | 2.95 | 1.56 |
| sp\|Q9H6T3-2\|RPAP3_HUMAN Isoform 2 of RNA polymerase II-associated protein 3 OS=Homo sapiens OX=9606 GN=RPAP3 | 4.27 | 8.73 | 2.05 | 1.03 |

| | | | | |
|---|---|---|---|---|
| tr\|Q5VXN0\|Q5VXN0_HUMAN Ribosome production factor 2 homolog (Fragment) OS=Homo sapiens OX=9606 GN=RPF2 PE=1 SV=1 | 4.26 | 6.06 | 1.42 | 0.51 |
| sp\|Q8WW12-2\|PCNP_HUMAN Isoform 2 of PEST proteolytic signal-containing nuclear protein OS=Homo sapiens OX=9606 GN=PCNP | 4.26 | 6.93 | 1.63 | 0.70 |
| sp\|Q8N0X7\|SPART_HUMAN Spartin OS=Homo sapiens OX=9606 GN=SPART PE=1 SV=1 | 4.22 | 8.20 | 1.94 | 0.96 |
| tr\|A0A0C4DGS0\|A0A0C4DGS0_HUMAN NADH dehydrogenase [ubiquinone] 1 alpha subcomplex subunit 6 OS=Homo sapiens OX=9606 GN=NDUFA6 PE=1 SV=1 | 4.21 | 16.02 | 3.80 | 1.93 |
| sp\|Q9H4A6\|GOLP3_HUMAN Golgi phosphoprotein 3 OS=Homo sapiens OX=9606 GN=GOLPH3 PE=1 SV=1 | 4.20 | 11.65 | 2.77 | 1.47 |
| sp\|Q8IVT2\|MISP_HUMAN Mitotic interactor and substrate of PLK1 OS=Homo sapiens OX=9606 GN=MISP PE=1 SV=1 | 4.20 | 3.29 | 0.78 | -0.35 |
| sp\|O95232\|LC7L3_HUMAN Luc7-like protein 3 OS=Homo sapiens OX=9606 GN=LUC7L3 PE=1 SV=2 | 4.20 | 19.33 | 4.60 | 2.20 |
| tr\|A0A2R8YFH5\|A0A2R8YFH5_HUMAN Protein transport protein SEC23 OS=Homo sapiens OX=9606 GN=SEC23B PE=1 SV=1 | 4.18 | 6.00 | 1.44 | 0.52 |
| tr\|I3L1L9\|I3L1L9_HUMAN Ethanolamine-phosphate cytidylyltransferase (Fragment) OS=Homo sapiens OX=9606 GN=PCYT2 PE=1 SV=1 | 4.16 | 5.63 | 1.35 | 0.44 |
| sp\|Q14247-2\|SRC8_HUMAN Isoform 2 of Src substrate cortactin OS=Homo sapiens OX=9606 GN=CTTN | 4.15 | 5.41 | 1.31 | 0.38 |
| sp\|O94826\|TOM70_HUMAN Mitochondrial import receptor subunit TOM70 OS=Homo sapiens OX=9606 GN=TOMM70 PE=1 SV=1 | 4.14 | 7.92 | 1.91 | 0.94 |
| sp\|Q9NPD3\|EXOS4_HUMAN Exosome complex component RRP41 OS=Homo sapiens OX=9606 GN=EXOSC4 PE=1 SV=3 | 4.13 | 17.52 | 4.24 | 2.08 |
| tr\|A0A0A0MSK5\|A0A0A0MSK5_HUMAN Torsin-1A-interacting protein 1 OS=Homo sapiens OX=9606 GN=TOR1AIP1 PE=1 SV=1 | 4.12 | 6.27 | 1.52 | 0.61 |
| sp\|Q00325-2\|MPCP_HUMAN Isoform B of Phosphate carrier protein, mitochondrial OS=Homo sapiens OX=9606 GN=SLC25A3 | 4.10 | 10.06 | 2.45 | 1.29 |
| tr\|A0A075B6F9\|A0A075B6F9_HUMAN Nitric oxide synthase-interacting protein OS=Homo sapiens OX=9606 GN=NOSIP PE=1 SV=1 | 4.09 | 5.71 | 1.39 | 0.48 |
| sp\|Q8N752\|KC1AL_HUMAN Casein kinase I isoform alpha-like OS=Homo sapiens OX=9606 GN=CSNK1A1L PE=2 SV=2 | 4.09 | 10.34 | 2.52 | 1.34 |
| sp\|Q96KA5-2\|CLP1L_HUMAN Isoform 2 of Cleft lip and palate transmembrane protein 1- | 4.09 | 4.70 | 1.15 | 0.20 |

| | | | | |
|---|---|---|---|---|
| like protein OS=Homo sapiens OX=9606 GN=CLPTM1L | | | | |
| tr\|C9JLU1\|C9JLU1_HUMAN DNA-directed RNA polymerases I, II, and III subunit RPABC3 (Fragment) OS=Homo sapiens OX=9606 GN=POLR2H PE=1 SV=8 | 4.07 | 14.37 | 3.53 | 1.82 |
| sp\|Q9NRN7\|ADPPT_HUMAN L-aminoadipate-semialdehyde dehydrogenase-phosphopantetheinyl transferase OS=Homo sapiens OX=9606 GN=AASDHPPT PE=1 SV=2 | 4.06 | 4.74 | 1.17 | 0.22 |
| sp\|Q9H845\|ACAD9_HUMAN Acyl-CoA dehydrogenase family member 9, mitochondrial OS=Homo sapiens OX=9606 GN=ACAD9 PE=1 SV=1 | 4.02 | 1.66 | 0.41 | -1.28 |
| sp\|P60981\|DEST_HUMAN Destrin OS=Homo sapiens OX=9606 GN=DSTN PE=1 SV=3 | 4.02 | 9.25 | 2.30 | 1.20 |
| sp\|Q8N1G4\|LRC47_HUMAN Leucine-rich repeat-containing protein 47 OS=Homo sapiens OX=9606 GN=LRRC47 PE=1 SV=1 | 4.00 | 2.96 | 0.74 | -0.44 |
| sp\|Q10713\|MPPA_HUMAN Mitochondrial-processing peptidase subunit alpha OS=Homo sapiens OX=9606 GN=PMPCA PE=1 SV=2 | 4.00 | 9.79 | 2.45 | 1.29 |
| tr\|A0A087WTB8\|A0A087WTB8_HUMAN Ubiquitin carboxyl-terminal hydrolase OS=Homo sapiens OX=9606 GN=UCHL3 PE=1 SV=1 | 3.94 | 3.87 | 0.98 | -0.03 |
| sp\|Q8WXX5\|DNJC9_HUMAN DnaJ homolog subfamily C member 9 OS=Homo sapiens OX=9606 GN=DNAJC9 PE=1 SV=1 | 3.94 | 12.81 | 3.25 | 1.70 |
| sp\|P40937-2\|RFC5_HUMAN Isoform 2 of Replication factor C subunit 5 OS=Homo sapiens OX=9606 GN=RFC5 | 3.93 | 2.16 | 0.55 | -0.86 |
| sp\|P51116\|FXR2_HUMAN Fragile X mental retardation syndrome-related protein 2 OS=Homo sapiens OX=9606 GN=FXR2 PE=1 SV=2 | 3.90 | 7.54 | 1.93 | 0.95 |
| sp\|P52306-2\|GDS1_HUMAN Isoform 2 of Rap1 GTPase-GDP dissociation stimulator 1 OS=Homo sapiens OX=9606 GN=RAP1GDS1 | 3.89 | 9.68 | 2.48 | 1.31 |
| sp\|O15460-2\|P4HA2_HUMAN Isoform IIa of Prolyl 4-hydroxylase subunit alpha-2 OS=Homo sapiens OX=9606 GN=P4HA2 | 3.89 | 2.65 | 0.68 | -0.56 |
| tr\|D6REA1\|D6REA1_HUMAN Nucleotide exchange factor SIL1 OS=Homo sapiens OX=9606 GN=SIL1 PE=1 SV=1 | 3.89 | 4.63 | 1.19 | 0.25 |
| tr\|H0YHU4\|H0YHU4_HUMAN tRNA (guanine-N(7)-)-methyltransferase (Fragment) OS=Homo sapiens OX=9606 GN=METTL1 PE=1 SV=1 | 3.88 | 5.33 | 1.37 | 0.46 |
| sp\|P17858\|PFKAL_HUMAN ATP-dependent 6-phosphofructokinase, liver type OS=Homo sapiens OX=9606 GN=PFKL PE=1 SV=6 | 3.88 | 10.52 | 2.72 | 1.44 |

| | | | | |
|---|---|---|---|---|
| sp\|Q96C86\|DCPS_HUMAN m7GpppX diphosphatase OS=Homo sapiens OX=9606 GN=DCPS PE=1 SV=2 | 3.86 | 10.47 | 2.71 | 1.44 |
| tr\|A0A087WUB9\|A0A087WUB9_HUMAN Beta-catenin-like protein 1 OS=Homo sapiens OX=9606 GN=CTNNBL1 PE=1 SV=1 | 3.86 | 3.95 | 1.02 | 0.03 |
| tr\|F5GYF7\|F5GYF7_HUMAN COP9 signalosome complex subunit 7a (Fragment) OS=Homo sapiens OX=9606 GN=COPS7A PE=1 SV=8 | 3.85 | 20.68 | 5.37 | 2.43 |
| tr\|Q5SWC8\|Q5SWC8_HUMAN Heterochromatin protein 1-binding protein 3 (Fragment) OS=Homo sapiens OX=9606 GN=HP1BP3 PE=1 SV=1 | 3.83 | 11.70 | 3.05 | 1.61 |
| tr\|J3KNF5\|J3KNF5_HUMAN Centrosomal protein of 290 kDa OS=Homo sapiens OX=9606 GN=CEP290 PE=1 SV=1 | 3.78 | 15.12 | 4.00 | 2.00 |
| sp\|Q9BQ75-2\|CMS1_HUMAN Isoform 2 of Protein CMSS1 OS=Homo sapiens OX=9606 GN=CMSS1 | 3.78 | 15.12 | 4.00 | 2.00 |
| sp\|Q9Y3C8\|UFC1_HUMAN Ubiquitin-fold modifier-conjugating enzyme 1 OS=Homo sapiens OX=9606 GN=UFC1 PE=1 SV=3 | 3.76 | 15.32 | 4.07 | 2.03 |
| sp\|Q3LXA3\|TKFC_HUMAN Triokinase/FMN cyclase OS=Homo sapiens OX=9606 GN=TKFC PE=1 SV=2 | 3.74 | 5.59 | 1.49 | 0.58 |
| sp\|Q9Y697-2\|NFS1_HUMAN Isoform Cytoplasmic of Cysteine desulfurase, mitochondrial OS=Homo sapiens OX=9606 GN=NFS1 | 3.74 | 2.50 | 0.67 | -0.58 |
| sp\|Q9P0I2-2\|EMC3_HUMAN Isoform 2 of ER membrane protein complex subunit 3 OS=Homo sapiens OX=9606 GN=EMC3 | 3.73 | 8.21 | 2.20 | 1.14 |
| sp\|O60749-2\|SNX2_HUMAN Isoform 2 of Sorting nexin-2 OS=Homo sapiens OX=9606 GN=SNX2 | 3.71 | 25.62 | 6.91 | 2.79 |
| tr\|C9JFK9\|C9JFK9_HUMAN BAG family molecular chaperone regulator 3 (Fragment) OS=Homo sapiens OX=9606 GN=BAG3 PE=1 SV=1 | 3.70 | 1.30 | 0.35 | -1.51 |
| tr\|H3BNB9\|H3BNB9_HUMAN RBR-type E3 ubiquitin transferase (Fragment) OS=Homo sapiens OX=9606 GN=ARIH1 PE=1 SV=1 | 3.69 | 8.36 | 2.27 | 1.18 |
| sp\|P21281\|VATB2_HUMAN V-type proton ATPase subunit B, brain isoform OS=Homo sapiens OX=9606 GN=ATP6V1B2 PE=1 SV=3 | 3.66 | 5.63 | 1.54 | 0.62 |
| sp\|Q9NVJ2\|ARL8B_HUMAN ADP-ribosylation factor-like protein 8B OS=Homo sapiens OX=9606 GN=ARL8B PE=1 SV=1 | 3.66 | 18.93 | 5.18 | 2.37 |
| tr\|F5GZY7\|F5GZY7_HUMAN Gamma-aminobutyric acid receptor-associated protein-like 1 (Fragment) OS=Homo sapiens OX=9606 GN=GABARAPL1 PE=1 SV=1 | 3.65 | 19.68 | 5.39 | 2.43 |

| | | | | |
|---|---|---|---|---|
| sp\|Q9UJX3-2\|APC7_HUMAN Isoform 2 of Anaphase-promoting complex subunit 7 OS=Homo sapiens OX=9606 GN=ANAPC7 | 3.63 | 9.20 | 2.53 | 1.34 |
| sp\|P41223-2\|BUD31_HUMAN Isoform 2 of Protein BUD31 homolog OS=Homo sapiens OX=9606 GN=BUD31 | 3.63 | 2.70 | 0.75 | -0.42 |
| sp\|Q8IZ83-3\|A16A1_HUMAN Isoform 3 of Aldehyde dehydrogenase family 16 member A1 OS=Homo sapiens OX=9606 GN=ALDH16A1 | 3.62 | 7.04 | 1.95 | 0.96 |
| sp\|O00567\|NOP56_HUMAN Nucleolar protein 56 OS=Homo sapiens OX=9606 GN=NOP56 PE=1 SV=4 | 3.62 | 6.37 | 1.76 | 0.82 |
| sp\|Q8IYB8\|SUV3_HUMAN ATP-dependent RNA helicase SUPV3L1, mitochondrial OS=Homo sapiens OX=9606 GN=SUPV3L1 PE=1 SV=1 | 3.61 | 6.61 | 1.83 | 0.87 |
| sp\|Q5TFE4\|NT5D1_HUMAN 5'-nucleotidase domain-containing protein 1 OS=Homo sapiens OX=9606 GN=NT5DC1 PE=1 SV=1 | 3.60 | 5.73 | 1.59 | 0.67 |
| sp\|Q9Y6N5\|SQOR_HUMAN Sulfide:quinone oxidoreductase, mitochondrial OS=Homo sapiens OX=9606 GN=SQOR PE=1 SV=1 | 3.60 | 6.89 | 1.92 | 0.94 |
| tr\|A0A0C4DGX4\|A0A0C4DGX4_HUMAN Cullin-1 OS=Homo sapiens OX=9606 GN=CUL1 PE=1 SV=1 | 3.59 | 10.13 | 2.82 | 1.49 |
| sp\|P09543-2\|CN37_HUMAN Isoform CNPI of 2',3'-cyclic-nucleotide 3'-phosphodiesterase OS=Homo sapiens OX=9606 GN=CNP | 3.59 | 6.89 | 1.92 | 0.94 |
| tr\|G3V0I5\|G3V0I5_HUMAN NADH dehydrogenase [ubiquinone] flavoprotein 1, mitochondrial OS=Homo sapiens OX=9606 GN=NDUFV1 PE=1 SV=1 | 3.57 | 5.96 | 1.67 | 0.74 |
| sp\|Q9UHR4\|BI2L1_HUMAN Brain-specific angiogenesis inhibitor 1-associated protein 2-like protein 1 OS=Homo sapiens OX=9606 GN=BAIAP2L1 PE=1 SV=2 | 3.56 | 3.53 | 0.99 | -0.01 |
| sp\|Q8TBQ9\|KISHA_HUMAN Protein kish-A OS=Homo sapiens OX=9606 GN=TMEM167A PE=1 SV=1 | 3.56 | 8.29 | 2.33 | 1.22 |
| sp\|Q13177\|PAK2_HUMAN Serine/threonine-protein kinase PAK 2 OS=Homo sapiens OX=9606 GN=PAK2 PE=1 SV=3 | 3.55 | 6.16 | 1.73 | 0.79 |
| tr\|A2A2Q9\|A2A2Q9_HUMAN Protein AAR2 homolog OS=Homo sapiens OX=9606 GN=AAR2 PE=1 SV=1 | 3.53 | 6.88 | 1.95 | 0.96 |
| sp\|Q9UI42-2\|CBPA4_HUMAN Isoform 2 of Carboxypeptidase A4 OS=Homo sapiens OX=9606 GN=CPA4 | 3.50 | 5.65 | 1.61 | 0.69 |
| tr\|A0A2R8Y5Q8\|A0A2R8Y5Q8_HUMAN Tubulin-specific chaperone E OS=Homo sapiens OX=9606 GN=TBCE PE=1 SV=1 | 3.50 | 6.68 | 1.91 | 0.93 |
| sp\|Q7L5D6-2\|GET4_HUMAN Isoform 2 of Golgi to ER traffic protein 4 homolog OS=Homo sapiens OX=9606 GN=GET4 | 3.50 | 7.17 | 2.05 | 1.04 |

| | | | | |
|---|---|---|---|---|
| sp\|Q9NZN4\|EHD2_HUMAN EH domain-containing protein 2 OS=Homo sapiens OX=9606 GN=EHD2 PE=1 SV=2 | 3.48 | 15.04 | 4.32 | 2.11 |
| sp\|Q6UXH1-2\|CREL2_HUMAN Isoform 2 of Cysteine-rich with EGF-like domain protein 2 OS=Homo sapiens OX=9606 GN=CRELD2 | 3.48 | 5.74 | 1.65 | 0.72 |
| sp\|Q9ULX3\|NOB1_HUMAN RNA-binding protein NOB1 OS=Homo sapiens OX=9606 GN=NOB1 PE=1 SV=1 | 3.48 | 5.32 | 1.53 | 0.61 |
| tr\|H0YK99\|H0YK99_HUMAN DnaJ homolog subfamily C member 17 OS=Homo sapiens OX=9606 GN=DNAJC17 PE=1 SV=1 | 3.46 | 10.83 | 3.13 | 1.65 |
| sp\|Q13505-3\|MTX1_HUMAN Isoform 3 of Metaxin-1 OS=Homo sapiens OX=9606 GN=MTX1 | 3.45 | 5.77 | 1.67 | 0.74 |
| tr\|H0Y2Y8\|H0Y2Y8_HUMAN Zyxin (Fragment) OS=Homo sapiens OX=9606 GN=ZYX PE=1 SV=1 | 3.45 | 7.02 | 2.03 | 1.02 |
| tr\|X6R5C5\|X6R5C5_HUMAN Carboxypeptidase OS=Homo sapiens OX=9606 GN=CTSA PE=1 SV=1 | 3.44 | 15.19 | 4.41 | 2.14 |
| sp\|Q86UE4\|LYRIC_HUMAN Protein LYRIC OS=Homo sapiens OX=9606 GN=MTDH PE=1 SV=2 | 3.44 | 6.40 | 1.86 | 0.90 |
| sp\|P78346-2\|RPP30_HUMAN Isoform 2 of Ribonuclease P protein subunit p30 OS=Homo sapiens OX=9606 GN=RPP30 | 3.39 | 6.08 | 1.79 | 0.84 |
| sp\|P10155-3\|RO60_HUMAN Isoform 3 of 60 kDa SS-A/Ro ribonucleoprotein OS=Homo sapiens OX=9606 GN=TROVE2 | 3.39 | 10.08 | 2.97 | 1.57 |
| sp\|Q15654\|TRIP6_HUMAN Thyroid receptor-interacting protein 6 OS=Homo sapiens OX=9606 GN=TRIP6 PE=1 SV=3 | 3.39 | 12.47 | 3.68 | 1.88 |
| tr\|H0YDX4\|H0YDX4_HUMAN Mitochondrial amidoxime-reducing component 1 (Fragment) OS=Homo sapiens OX=9606 GN=MARC1 PE=1 SV=1 | 3.39 | 9.64 | 2.85 | 1.51 |
| sp\|Q9UG63-2\|ABCF2_HUMAN Isoform 2 of ATP-binding cassette sub-family F member 2 OS=Homo sapiens OX=9606 GN=ABCF2 | 3.38 | 10.78 | 3.19 | 1.67 |
| tr\|E9PBF6\|E9PBF6_HUMAN Lamin-B1 OS=Homo sapiens OX=9606 GN=LMNB1 PE=1 SV=1 | 3.37 | 10.60 | 3.14 | 1.65 |
| sp\|Q92905\|CSN5_HUMAN COP9 signalosome complex subunit 5 OS=Homo sapiens OX=9606 GN=COPS5 PE=1 SV=4 | 3.37 | 9.18 | 2.73 | 1.45 |
| sp\|Q8TAT6-2\|NPL4_HUMAN Isoform 2 of Nuclear protein localization protein 4 homolog OS=Homo sapiens OX=9606 GN=NPLOC4 | 3.36 | 9.47 | 2.82 | 1.50 |
| sp\|Q86TU7\|SETD3_HUMAN Histone-lysine N-methyltransferase setd3 OS=Homo sapiens OX=9606 GN=SETD3 PE=1 SV=1 | 3.35 | 0.61 | 0.18 | -2.46 |

| | | | | |
|---|---|---|---|---|
| tr\|H0YK61\|H0YK61_HUMAN ER membrane protein complex subunit 4 OS=Homo sapiens OX=9606 GN=EMC4 PE=1 SV=1 | 3.32 | 15.74 | 4.74 | 2.24 |
| sp\|P08240-2\|SRPRA_HUMAN Isoform 2 of Signal recognition particle receptor subunit alpha OS=Homo sapiens OX=9606 GN=SRPRA | 3.32 | 7.95 | 2.39 | 1.26 |
| sp\|Q01581\|HMCS1_HUMAN Hydroxymethylglutaryl-CoA synthase, cytoplasmic OS=Homo sapiens OX=9606 GN=HMGCS1 PE=1 SV=2 | 3.31 | 6.14 | 1.86 | 0.89 |
| tr\|B1AK81\|B1AK81_HUMAN GPI-anchor transamidase OS=Homo sapiens OX=9606 GN=PIGK PE=1 SV=1 | 3.30 | 5.65 | 1.71 | 0.78 |
| sp\|Q9UKD2\|MRT4_HUMAN mRNA turnover protein 4 homolog OS=Homo sapiens OX=9606 GN=MRTO4 PE=1 SV=2 | 3.30 | 5.22 | 1.58 | 0.66 |
| tr\|H0Y8C2\|H0Y8C2_HUMAN 60S ribosomal protein L22-like 1 (Fragment) OS=Homo sapiens OX=9606 GN=RPL22L1 PE=1 SV=1 | 3.28 | 13.36 | 4.08 | 2.03 |
| tr\|A0A1W2PNP0\|A0A1W2PNP0_HUMAN GPI transamidase component PIG-T (Fragment) OS=Homo sapiens OX=9606 GN=PIGT PE=1 SV=1 | 3.26 | 9.28 | 2.84 | 1.51 |
| tr\|F5H1S9\|F5H1S9_HUMAN tRNA pseudouridine synthase OS=Homo sapiens OX=9606 GN=PUS1 PE=1 SV=1 | 3.25 | 4.71 | 1.45 | 0.53 |
| sp\|Q96GM8\|TOE1_HUMAN Target of EGR1 protein 1 OS=Homo sapiens OX=9606 GN=TOE1 PE=1 SV=1 | 3.25 | 8.39 | 2.58 | 1.37 |
| sp\|Q15126\|PMVK_HUMAN Phosphomevalonate kinase OS=Homo sapiens OX=9606 GN=PMVK PE=1 SV=3 | 3.25 | 2.67 | 0.82 | -0.28 |
| sp\|P36639-2\|8ODP_HUMAN Isoform p22 of 7,8-dihydro-8-oxoguanine triphosphatase OS=Homo sapiens OX=9606 GN=NUDT1 | 3.24 | 3.85 | 1.19 | 0.25 |
| tr\|A0A087WVC4\|A0A087WVC4_HUMAN cAMP-dependent protein kinase catalytic subunit beta OS=Homo sapiens OX=9606 GN=PRKACB PE=1 SV=1 | 3.24 | 7.59 | 2.34 | 1.23 |
| tr\|K7EP06\|K7EP06_HUMAN mRNA cap guanine-N7 methyltransferase OS=Homo sapiens OX=9606 GN=RNMT PE=1 SV=1 | 3.22 | 9.24 | 2.87 | 1.52 |
| tr\|I3LCI2\|I3LCI2_PIG Pyruvate dehydrogenase E1 component subunit alpha OS=Sus scrofa GN=PDHA1 PE=1 SV=1 | 3.20 | 4.47 | 1.40 | 0.48 |
| sp\|O96007\|MOC2B_HUMAN Molybdopterin synthase catalytic subunit OS=Homo sapiens OX=9606 GN=MOCS2 PE=1 SV=1 | 3.20 | 7.43 | 2.32 | 1.22 |
| sp\|Q9BYG3\|MK67I_HUMAN MKI67 FHA domain-interacting nucleolar phosphoprotein OS=Homo sapiens OX=9606 GN=NIFK PE=1 SV=1 | 3.19 | 6.32 | 1.98 | 0.98 |

| | | | | |
|---|---|---|---|---|
| sp\|Q15428\|SF3A2_HUMAN Splicing factor 3A subunit 2 OS=Homo sapiens OX=9606 GN=SF3A2 PE=1 SV=2 | 3.19 | 0.25 | 0.08 | -3.67 |
| tr\|G3V2E7\|G3V2E7_HUMAN Kinesin light chain 1 OS=Homo sapiens OX=9606 GN=KLC1 PE=1 SV=1 | 3.12 | 1.98 | 0.64 | -0.65 |
| sp\|P56182\|RRP1_HUMAN Ribosomal RNA processing protein 1 homolog A OS=Homo sapiens OX=9606 GN=RRP1 PE=1 SV=1 | 3.12 | 0.93 | 0.30 | -1.74 |
| sp\|Q9Y305-2\|ACOT9_HUMAN Isoform 2 of Acyl-coenzyme A thioesterase 9, mitochondrial OS=Homo sapiens OX=9606 GN=ACOT9 | 3.08 | 5.85 | 1.90 | 0.93 |
| sp\|Q9NX62\|IMPA3_HUMAN Inositol monophosphatase 3 OS=Homo sapiens OX=9606 GN=IMPAD1 PE=1 SV=1 | 3.07 | 7.17 | 2.33 | 1.22 |
| sp\|Q9Y3T9\|NOC2L_HUMAN Nucleolar complex protein 2 homolog OS=Homo sapiens OX=9606 GN=NOC2L PE=1 SV=4 | 3.05 | 3.73 | 1.22 | 0.29 |
| sp\|Q9H6Y2\|WDR55_HUMAN WD repeat-containing protein 55 OS=Homo sapiens OX=9606 GN=WDR55 PE=1 SV=2 | 3.02 | 3.02 | 1.00 | 0.00 |
| sp\|Q14181\|DPOA2_HUMAN DNA polymerase alpha subunit B OS=Homo sapiens OX=9606 GN=POLA2 PE=1 SV=2 | 3.01 | 4.56 | 1.51 | 0.60 |
| tr\|B1AXG1\|B1AXG1_HUMAN Non-specific serine/threonine protein kinase OS=Homo sapiens OX=9606 GN=RPS6KA3 PE=1 SV=2 | 3.01 | 4.99 | 1.66 | 0.73 |
| tr\|E5RIJ3\|E5RIJ3_HUMAN Tumor necrosis factor alpha-induced protein 8 (Fragment) OS=Homo sapiens OX=9606 GN=TNFAIP8 PE=1 SV=1 | 3.00 | 12.57 | 4.19 | 2.07 |
| tr\|A0A087X0W7\|A0A087X0W7_HUMAN Acyl-coenzyme A thioesterase 2, mitochondrial OS=Homo sapiens OX=9606 GN=ACOT2 PE=1 SV=1 | 2.98 | 4.91 | 1.65 | 0.72 |
| sp\|Q6VUC0\|AP2E_HUMAN Transcription factor AP-2-epsilon OS=Homo sapiens OX=9606 GN=TFAP2E PE=2 SV=1 | 2.97 | 3.69 | 1.24 | 0.31 |
| tr\|C9JZR2\|C9JZR2_HUMAN Catenin delta-1 OS=Homo sapiens OX=9606 GN=CTNND1 PE=1 SV=2 | 2.97 | 1.39 | 0.47 | -1.10 |
| sp\|Q96BN8\|OTUL_HUMAN Ubiquitin thioesterase otulin OS=Homo sapiens OX=9606 GN=OTULIN PE=1 SV=3 | 2.96 | 7.93 | 2.67 | 1.42 |
| sp\|Q08J23-2\|NSUN2_HUMAN Isoform 2 of tRNA (cytosine(34)-C(5))-methyltransferase OS=Homo sapiens OX=9606 GN=NSUN2 | 2.96 | 12.49 | 4.22 | 2.08 |
| sp\|P49770\|EI2BB_HUMAN Translation initiation factor eIF-2B subunit beta OS=Homo sapiens OX=9606 GN=EIF2B2 PE=1 SV=3 | 2.95 | 12.03 | 4.07 | 2.03 |
| tr\|A0A0X1KG71\|A0A0X1KG71_HUMAN Negative elongation factor B OS=Homo sapiens OX=9606 GN=NELFB PE=1 SV=1 | 2.94 | 7.96 | 2.71 | 1.44 |

| | | | | |
|---|---|---|---|---|
| sp\|P49761-1\|CLK3_HUMAN Isoform 1 of Dual specificity protein kinase CLK3 OS=Homo sapiens OX=9606 GN=CLK3 | 2.93 | 4.51 | 1.54 | 0.62 |
| sp\|Q9H814\|PHAX_HUMAN Phosphorylated adapter RNA export protein OS=Homo sapiens OX=9606 GN=PHAX PE=1 SV=1 | 2.92 | 5.72 | 1.96 | 0.97 |
| sp\|Q6ZRP7\|QSOX2_HUMAN Sulfhydryl oxidase 2 OS=Homo sapiens OX=9606 GN=QSOX2 PE=1 SV=3 | 2.92 | 1.40 | 0.48 | -1.06 |
| tr\|F8W7U8\|F8W7U8_HUMAN Double-strand break repair protein OS=Homo sapiens OX=9606 GN=MRE11 PE=1 SV=1 | 2.91 | 10.65 | 3.66 | 1.87 |
| tr\|J3KTC1\|J3KTC1_HUMAN RNA-binding protein Musashi homolog 2 (Fragment) OS=Homo sapiens OX=9606 GN=MSI2 PE=1 SV=8 | 2.90 | 13.24 | 4.56 | 2.19 |
| tr\|A0A087X0K1\|A0A087X0K1_HUMAN Calcium-binding protein 39 OS=Homo sapiens OX=9606 GN=CAB39 PE=1 SV=1 | 2.90 | 6.12 | 2.11 | 1.08 |
| tr\|A0A1W2PNX8\|A0A1W2PNX8_HUMAN Protein unc-45 homolog A OS=Homo sapiens OX=9606 GN=UNC45A PE=1 SV=1 | 2.88 | 3.29 | 1.14 | 0.19 |
| tr\|E7ENJ6\|E7ENJ6_HUMAN AP-1 complex subunit mu-1 OS=Homo sapiens OX=9606 GN=AP1M1 PE=1 SV=1 | 2.86 | 5.83 | 2.04 | 1.03 |
| sp\|Q9NZL4-3\|HPBP1_HUMAN Isoform 3 of Hsp70-binding protein 1 OS=Homo sapiens OX=9606 GN=HSPBP1 | 2.86 | 0.16 | 0.05 | -4.20 |
| sp\|Q13084\|RM28_HUMAN 39S ribosomal protein L28, mitochondrial OS=Homo sapiens OX=9606 GN=MRPL28 PE=1 SV=4 | 2.85 | 5.81 | 2.04 | 1.03 |
| tr\|E5RGS4\|E5RGS4_HUMAN Prefoldin subunit 1 OS=Homo sapiens OX=9606 GN=PFDN1 PE=1 SV=1 | 2.85 | 7.03 | 2.47 | 1.30 |
| sp\|Q8NF37\|PCAT1_HUMAN Lysophosphatidylcholine acyltransferase 1 OS=Homo sapiens OX=9606 GN=LPCAT1 PE=1 SV=2 | 2.81 | 3.29 | 1.17 | 0.23 |
| sp\|Q05048\|CSTF1_HUMAN Cleavage stimulation factor subunit 1 OS=Homo sapiens OX=9606 GN=CSTF1 PE=1 SV=1 | 2.80 | 5.79 | 2.07 | 1.05 |
| tr\|I3L0N3\|I3L0N3_HUMAN Vesicle-fusing ATPase OS=Homo sapiens OX=9606 GN=NSF PE=1 SV=1 | 2.79 | 7.74 | 2.78 | 1.47 |
| sp\|O00629\|IMA3_HUMAN Importin subunit alpha-3 OS=Homo sapiens OX=9606 GN=KPNA4 PE=1 SV=1 | 2.77 | 4.37 | 1.58 | 0.66 |
| tr\|X6RAY8\|X6RAY8_HUMAN 39S ribosomal protein L4, mitochondrial OS=Homo sapiens OX=9606 GN=MRPL4 PE=1 SV=1 | 2.73 | 10.87 | 3.98 | 1.99 |
| tr\|G5E9V5\|G5E9V5_HUMAN 28S ribosomal protein S22, mitochondrial OS=Homo sapiens OX=9606 GN=MRPS22 PE=1 SV=1 | 2.71 | 18.38 | 6.77 | 2.76 |

| | | | | |
|---|---|---|---|---|
| sp\|Q8NI60-3\|COQ8A_HUMAN Isoform 3 of Atypical kinase COQ8A, mitochondrial OS=Homo sapiens OX=9606 GN=COQ8A | 2.70 | 4.70 | 1.74 | 0.80 |
| sp\|O14879\|IFIT3_HUMAN Interferon-induced protein with tetratricopeptide repeats 3 OS=Homo sapiens OX=9606 GN=IFIT3 PE=1 SV=1 | 2.68 | 14.16 | 5.28 | 2.40 |
| sp\|Q8N7H5-2\|PAF1_HUMAN Isoform 2 of RNA polymerase II-associated factor 1 homolog OS=Homo sapiens OX=9606 GN=PAF1 | 2.68 | 4.01 | 1.50 | 0.58 |
| sp\|P36404\|ARL2_HUMAN ADP-ribosylation factor-like protein 2 OS=Homo sapiens OX=9606 GN=ARL2 PE=1 SV=4 | 2.68 | 9.29 | 3.47 | 1.79 |
| sp\|P50583\|AP4A_HUMAN Bis(5'-nucleosyl)-tetraphosphatase [asymmetrical] OS=Homo sapiens OX=9606 GN=NUDT2 PE=1 SV=3 | 2.66 | 3.91 | 1.47 | 0.56 |
| tr\|J3QKW2\|J3QKW2_HUMAN 28S ribosomal protein S7, mitochondrial OS=Homo sapiens OX=9606 GN=MRPS7 PE=1 SV=1 | 2.66 | 9.50 | 3.57 | 1.84 |
| tr\|A0A0B4J287\|A0A0B4J287_HUMAN COMM domain-containing protein 4 OS=Homo sapiens OX=9606 GN=COMMD4 PE=1 SV=1 | 2.65 | 12.28 | 4.63 | 2.21 |
| sp\|Q8TBC4-2\|UBA3_HUMAN Isoform 2 of NEDD8-activating enzyme E1 catalytic subunit OS=Homo sapiens OX=9606 GN=UBA3 | 2.65 | 2.90 | 1.09 | 0.13 |
| sp\|O75251\|NDUS7_HUMAN NADH dehydrogenase [ubiquinone] iron-sulfur protein 7, mitochondrial OS=Homo sapiens OX=9606 GN=NDUFS7 PE=1 SV=3 | 2.63 | 7.51 | 2.85 | 1.51 |
| tr\|A0A0C4DGG1\|A0A0C4DGG1_HUMAN Protein kinase C and casein kinase substrate in neurons protein 3 (Fragment) OS=Homo sapiens OX=9606 GN=PACSIN3 PE=1 SV=1 | 2.62 | 16.93 | 6.46 | 2.69 |
| tr\|A0A0A0MS51\|A0A0A0MS51_HUMAN Gelsolin OS=Homo sapiens OX=9606 GN=GSN PE=1 SV=1 | 2.62 | 3.25 | 1.24 | 0.31 |
| tr\|A0A1B0GUA3\|A0A1B0GUA3_HUMAN KIF1-binding protein OS=Homo sapiens OX=9606 GN=KIF1BP PE=1 SV=1 | 2.61 | 17.87 | 6.83 | 2.77 |
| tr\|A0A0A0MQR2\|A0A0A0MQR2_HUMAN Replication termination factor 2 OS=Homo sapiens OX=9606 GN=RTF2 PE=1 SV=1 | 2.57 | 4.66 | 1.81 | 0.86 |
| sp\|P33947\|ERD22_HUMAN ER lumen protein-retaining receptor 2 OS=Homo sapiens OX=9606 GN=KDELR2 PE=1 SV=1 | 2.57 | 2.11 | 0.82 | -0.29 |
| tr\|B5ME97\|B5ME97_HUMAN Septin 10, isoform CRA_c OS=Homo sapiens OX=9606 GN=SEPT10 PE=1 SV=2 | 2.56 | 7.54 | 2.94 | 1.56 |
| sp\|P31937\|3HIDH_HUMAN 3-hydroxyisobutyrate dehydrogenase, mitochondrial OS=Homo sapiens OX=9606 GN=HIBADH PE=1 SV=2 | 2.56 | 5.28 | 2.07 | 1.05 |
| tr\|A0A2U3TZY2\|A0A2U3TZY2_HUMAN Caseinolytic peptidase B protein homolog | 2.55 | 3.67 | 1.44 | 0.53 |

| | | | | |
|---|---|---|---|---|
| OS=Homo sapiens OX=9606 GN=CLPB PE=1 SV=1 | | | | |
| sp\|O96006\|ZBED1_HUMAN Zinc finger BED domain-containing protein 1 OS=Homo sapiens OX=9606 GN=ZBED1 PE=1 SV=1 | 2.55 | 2.07 | 0.81 | -0.30 |
| tr\|M0R0P9\|M0R0P9_HUMAN Non-specific serine/threonine protein kinase OS=Homo sapiens OX=9606 GN=AKT2 PE=1 SV=1 | 2.54 | 3.30 | 1.30 | 0.38 |
| sp\|Q04446\|GLGB_HUMAN 1,4-alpha-glucan-branching enzyme OS=Homo sapiens OX=9606 GN=GBE1 PE=1 SV=3 | 2.54 | 19.12 | 7.54 | 2.91 |
| sp\|Q13724\|MOGS_HUMAN Mannosyl-oligosaccharide glucosidase OS=Homo sapiens OX=9606 GN=MOGS PE=1 SV=5 | 2.53 | 14.30 | 5.65 | 2.50 |
| tr\|F8VX04\|F8VX04_HUMAN Sodium-coupled neutral amino acid transporter 1 OS=Homo sapiens OX=9606 GN=SLC38A1 PE=1 SV=1 | 2.53 | 0.12 | 0.05 | -4.41 |
| tr\|H0Y3C5\|H0Y3C5_HUMAN Tyrosine-protein kinase OS=Homo sapiens OX=9606 GN=HCK PE=1 SV=1 | 2.52 | 3.50 | 1.39 | 0.47 |
| sp\|O76031\|CLPX_HUMAN ATP-dependent Clp protease ATP-binding subunit clpX-like, mitochondrial OS=Homo sapiens OX=9606 GN=CLPX PE=1 SV=2 | 2.52 | 12.22 | 4.86 | 2.28 |
| sp\|P83111\|LACTB_HUMAN Serine beta-lactamase-like protein LACTB, mitochondrial OS=Homo sapiens OX=9606 GN=LACTB PE=1 SV=2 | 2.51 | 3.36 | 1.34 | 0.42 |
| sp\|Q9NSI2-2\|F207A_HUMAN Isoform B of Protein FAM207A OS=Homo sapiens OX=9606 GN=FAM207A | 2.49 | 8.08 | 3.24 | 1.70 |
| tr\|H0YF61\|H0YF61_HUMAN Non-specific lipid-transfer protein (Fragment) OS=Homo sapiens OX=9606 GN=SCP2 PE=1 SV=1 | 2.49 | 11.95 | 4.80 | 2.26 |
| sp\|Q6PI48\|SYDM_HUMAN Aspartate--tRNA ligase, mitochondrial OS=Homo sapiens OX=9606 GN=DARS2 PE=1 SV=1 | 2.47 | 6.33 | 2.57 | 1.36 |
| sp\|Q13564-2\|ULA1_HUMAN Isoform 2 of NEDD8-activating enzyme E1 regulatory subunit OS=Homo sapiens OX=9606 GN=NAE1 | 2.46 | 5.32 | 2.16 | 1.11 |
| sp\|P07384\|CAN1_HUMAN Calpain-1 catalytic subunit OS=Homo sapiens OX=9606 GN=CAPN1 PE=1 SV=1 | 2.44 | 8.95 | 3.67 | 1.88 |
| tr\|A0A087WZ13\|A0A087WZ13_HUMAN Ribonucleoprotein PTB-binding 1 OS=Homo sapiens OX=9606 GN=RAVER1 PE=1 SV=1 | 2.42 | 12.75 | 5.27 | 2.40 |
| tr\|A8MYV2\|A8MYV2_HUMAN LUC7-like (S. cerevisiae), isoform CRA_f OS=Homo sapiens OX=9606 GN=LUC7L PE=1 SV=1 | 2.40 | 6.58 | 2.74 | 1.45 |
| tr\|C9IY70\|C9IY70_HUMAN 60S ribosomal export protein NMD3 (Fragment) OS=Homo sapiens OX=9606 GN=NMD3 PE=1 SV=1 | 2.40 | 1.65 | 0.69 | -0.54 |

| | | | | |
|---|---|---|---|---|
| sp\|Q9UMX0-2\|UBQL1_HUMAN Isoform 2 of Ubiquilin-1 OS=Homo sapiens OX=9606 GN=UBQLN1 | 2.38 | 5.87 | 2.47 | 1.30 |
| tr\|B1AP22\|B1AP22_HUMAN Presenilin OS=Homo sapiens OX=9606 GN=PSEN2 PE=1 SV=1 | 2.35 | 6.47 | 2.75 | 1.46 |
| tr\|E7EMB1\|E7EMB1_HUMAN Switch-associated protein 70 OS=Homo sapiens OX=9606 GN=SWAP70 PE=1 SV=1 | 2.35 | 2.29 | 0.98 | -0.04 |
| sp\|Q9H307\|PININ_HUMAN Pinin OS=Homo sapiens OX=9606 GN=PNN PE=1 SV=5 | 2.35 | 1.81 | 0.77 | -0.37 |
| sp\|Q6L8Q7-2\|PDE12_HUMAN Isoform 2 of 2',5'-phosphodiesterase 12 OS=Homo sapiens OX=9606 GN=PDE12 | 2.32 | 4.49 | 1.93 | 0.95 |
| tr\|A0A087WT20\|A0A087WT20_HUMAN DDB1- and CUL4-associated factor 13 OS=Homo sapiens OX=9606 GN=DCAF13 PE=1 SV=1 | 2.32 | 1.34 | 0.58 | -0.79 |
| sp\|Q9H6S3\|ES8L2_HUMAN Epidermal growth factor receptor kinase substrate 8-like protein 2 OS=Homo sapiens OX=9606 GN=EPS8L2 PE=1 SV=2 | 2.31 | 2.30 | 0.99 | -0.01 |
| tr\|K7ER96\|K7ER96_HUMAN Thioredoxin-like protein 1 (Fragment) OS=Homo sapiens OX=9606 GN=TXNL1 PE=1 SV=1 | 2.28 | 6.88 | 3.02 | 1.60 |
| tr\|E7EQU1\|E7EQU1_HUMAN High mobility group protein B3 (Fragment) OS=Homo sapiens OX=9606 GN=HMGB3 PE=1 SV=1 | 2.26 | 9.39 | 4.16 | 2.06 |
| sp\|Q96S52-2\|PIGS_HUMAN Isoform 2 of GPI transamidase component PIG-S OS=Homo sapiens OX=9606 GN=PIGS | 2.23 | 3.66 | 1.64 | 0.71 |
| sp\|Q9Y2T2\|AP3M1_HUMAN AP-3 complex subunit mu-1 OS=Homo sapiens OX=9606 GN=AP3M1 PE=1 SV=1 | 2.21 | 3.67 | 1.66 | 0.73 |
| sp\|Q96P11-2\|NSUN5_HUMAN Isoform 2 of Probable 28S rRNA (cytosine-C(5))-methyltransferase OS=Homo sapiens OX=9606 GN=NSUN5 | 2.20 | 3.55 | 1.62 | 0.69 |
| tr\|H0YA56\|H0YA56_HUMAN Apoptosis regulator BAX (Fragment) OS=Homo sapiens OX=9606 GN=BAX PE=1 SV=1 | 2.19 | 5.28 | 2.41 | 1.27 |
| tr\|H0YCY8\|H0YCY8_HUMAN Dipeptidyl peptidase 1 (Fragment) OS=Homo sapiens OX=9606 GN=CTSC PE=1 SV=8 | 2.19 | 7.35 | 3.35 | 1.75 |
| tr\|E9PDQ8\|E9PDQ8_HUMAN Succinate--CoA ligase [GDP-forming] subunit beta, mitochondrial OS=Homo sapiens OX=9606 GN=SUCLG2 PE=1 SV=1 | 2.17 | 6.53 | 3.00 | 1.59 |
| sp\|Q9UKN8\|TF3C4_HUMAN General transcription factor 3C polypeptide 4 OS=Homo sapiens OX=9606 GN=GTF3C4 PE=1 SV=2 | 2.17 | 1.65 | 0.76 | -0.39 |
| sp\|Q14232\|EI2BA_HUMAN Translation initiation factor eIF-2B subunit alpha OS=Homo sapiens OX=9606 GN=EIF2B1 PE=1 SV=1 | 2.15 | 6.19 | 2.88 | 1.53 |

| | | | | |
|---|---|---|---|---|
| sp\|Q9Y2X3\|NOP58_HUMAN Nucleolar protein 58 OS=Homo sapiens OX=9606 GN=NOP58 PE=1 SV=1 | 2.14 | 7.73 | 3.61 | 1.85 |
| sp\|Q9BYN8\|RT26_HUMAN 28S ribosomal protein S26, mitochondrial OS=Homo sapiens OX=9606 GN=MRPS26 PE=1 SV=1 | 2.12 | 3.29 | 1.55 | 0.63 |
| tr\|Q5T760\|Q5T760_HUMAN Serine/arginine-rich-splicing factor 11 (Fragment) OS=Homo sapiens OX=9606 GN=SRSF11 PE=1 SV=1 | 2.12 | 6.45 | 3.04 | 1.60 |
| tr\|A6NG10\|A6NG10_HUMAN cDNA FLJ59459, highly similar to WW domain-binding protein 2 OS=Homo sapiens OX=9606 GN=WBP2 PE=1 SV=2 | 2.12 | 10.40 | 4.91 | 2.30 |
| sp\|Q92820\|GGH_HUMAN Gamma-glutamyl hydrolase OS=Homo sapiens OX=9606 GN=GGH PE=1 SV=2 | 2.10 | 4.04 | 1.92 | 0.94 |
| tr\|J3KMX2\|J3KMX2_HUMAN SWI/SNF-related matrix-associated actin-dependent regulator of chromatin subfamily D member 2 OS=Homo sapiens OX=9606 GN=SMARCD2 PE=1 SV=1 | 2.10 | 2.72 | 1.30 | 0.38 |
| sp\|P20340-2\|RAB6A_HUMAN Isoform 2 of Ras-related protein Rab-6A OS=Homo sapiens OX=9606 GN=RAB6A | 2.08 | 6.11 | 2.94 | 1.56 |
| sp\|Q9UBV8\|PEF1_HUMAN Peflin OS=Homo sapiens OX=9606 GN=PEF1 PE=1 SV=1 | 2.07 | 3.88 | 1.87 | 0.90 |
| sp\|Q8WVY7\|UBCP1_HUMAN Ubiquitin-like domain-containing CTD phosphatase 1 OS=Homo sapiens OX=9606 GN=UBLCP1 PE=1 SV=2 | 2.07 | 3.21 | 1.55 | 0.63 |
| sp\|Q96IJ6-2\|GMPPA_HUMAN Isoform 2 of Mannose-1-phosphate guanyltransferase alpha OS=Homo sapiens OX=9606 GN=GMPPA | 2.07 | 1.92 | 0.93 | -0.11 |
| sp\|Q13895\|BYST_HUMAN Bystin OS=Homo sapiens OX=9606 GN=BYSL PE=1 SV=3 | 2.06 | 6.86 | 3.33 | 1.73 |
| tr\|J3QQV6\|J3QQV6_HUMAN Pyridoxine-5'-phosphate oxidase (Fragment) OS=Homo sapiens OX=9606 GN=PNPO PE=1 SV=2 | 2.03 | 6.36 | 3.13 | 1.65 |
| sp\|Q9Y4W6\|AFG32_HUMAN AFG3-like protein 2 OS=Homo sapiens OX=9606 GN=AFG3L2 PE=1 SV=2 | 2.02 | 3.94 | 1.95 | 0.97 |
| sp\|P32929-2\|CGL_HUMAN Isoform 2 of Cystathionine gamma-lyase OS=Homo sapiens OX=9606 GN=CTH | 2.01 | 5.06 | 2.51 | 1.33 |
| sp\|O14530-2\|TXND9_HUMAN Isoform 2 of Thioredoxin domain-containing protein 9 OS=Homo sapiens OX=9606 GN=TXNDC9 | 2.01 | 7.18 | 3.58 | 1.84 |
| tr\|G3V155\|G3V155_HUMAN Thioredoxin domain containing 14, isoform CRA_a OS=Homo sapiens OX=9606 GN=TMX2 PE=4 SV=1 | 2.00 | 3.92 | 1.96 | 0.97 |
| tr\|A0A2R8Y3X5\|A0A2R8Y3X5_HUMAN Dynamin-like 120 kDa protein, mitochondrial OS=Homo sapiens OX=9606 GN=OPA1 PE=1 SV=1 | 2.00 | 0.51 | 0.26 | -1.97 |

| | | | | |
|---|---|---|---|---|
| sp\|Q9C005\|DPY30_HUMAN Protein dpy-30 homolog OS=Homo sapiens OX=9606 GN=DPY30 PE=1 SV=1 | 2.00 | 10.24 | 5.13 | 2.36 |
| sp\|O43823\|AKAP8_HUMAN A-kinase anchor protein 8 OS=Homo sapiens OX=9606 GN=AKAP8 PE=1 SV=1 | 2.00 | 1.70 | 0.85 | -0.23 |
| tr\|H3BMU1\|H3BMU1_HUMAN IST1 homolog (Fragment) OS=Homo sapiens OX=9606 GN=IST1 PE=1 SV=1 | 1.97 | 3.63 | 1.84 | 0.88 |
| tr\|A0A087WX97\|A0A087WX97_HUMAN Bcl-2-like protein 13 OS=Homo sapiens OX=9606 GN=BCL2L13 PE=1 SV=1 | 1.96 | 0.15 | 0.08 | -3.71 |
| sp\|Q96H20-2\|SNF8_HUMAN Isoform 2 of Vacuolar-sorting protein SNF8 OS=Homo sapiens OX=9606 GN=SNF8 | 1.94 | 2.39 | 1.23 | 0.30 |
| sp\|Q2TAY7\|SMU1_HUMAN WD40 repeat-containing protein SMU1 OS=Homo sapiens OX=9606 GN=SMU1 PE=1 SV=2 | 1.93 | 1.36 | 0.70 | -0.51 |
| sp\|Q8IXB1-2\|DJC10_HUMAN Isoform 2 of DnaJ homolog subfamily C member 10 OS=Homo sapiens OX=9606 GN=DNAJC10 | 1.91 | 1.53 | 0.80 | -0.32 |
| sp\|Q8TC07-2\|TBC15_HUMAN Isoform 2 of TBC1 domain family member 15 OS=Homo sapiens OX=9606 GN=TBC1D15 | 1.90 | 4.77 | 2.50 | 1.32 |
| sp\|Q14790-2\|CASP8_HUMAN Isoform 2 of Caspase-8 OS=Homo sapiens OX=9606 GN=CASP8 | 1.90 | 2.08 | 1.09 | 0.13 |
| sp\|Q8WXF1-2\|PSPC1_HUMAN Isoform 2 of Paraspeckle component 1 OS=Homo sapiens OX=9606 GN=PSPC1 | 1.88 | 6.42 | 3.42 | 1.77 |
| sp\|Q13363-2\|CTBP1_HUMAN Isoform 2 of C-terminal-binding protein 1 OS=Homo sapiens OX=9606 GN=CTBP1 | 1.87 | 3.47 | 1.85 | 0.89 |
| sp\|P35813-2\|PPM1A_HUMAN Isoform Alpha-2 of Protein phosphatase 1A OS=Homo sapiens OX=9606 GN=PPM1A | 1.87 | 2.24 | 1.20 | 0.26 |
| sp\|Q12996\|CSTF3_HUMAN Cleavage stimulation factor subunit 3 OS=Homo sapiens OX=9606 GN=CSTF3 PE=1 SV=1 | 1.87 | 8.93 | 4.78 | 2.26 |
| sp\|Q9BVC6\|TM109_HUMAN Transmembrane protein 109 OS=Homo sapiens OX=9606 GN=TMEM109 PE=1 SV=1 | 1.85 | 14.14 | 7.64 | 2.93 |
| sp\|Q9BZE4-2\|NOG1_HUMAN Isoform 2 of Nucleolar GTP-binding protein 1 OS=Homo sapiens OX=9606 GN=GTPBP4 | 1.82 | 1.92 | 1.06 | 0.08 |
| sp\|O43447\|PPIH_HUMAN Peptidyl-prolyl cis-trans isomerase H OS=Homo sapiens OX=9606 GN=PPIH PE=1 SV=1 | 1.82 | 4.38 | 2.41 | 1.27 |
| sp\|O60825-2\|F262_HUMAN Isoform 2 of 6-phosphofructo-2-kinase/fructose-2,6-bisphosphatase 2 OS=Homo sapiens OX=9606 GN=PFKFB2 | 1.82 | 1.44 | 0.79 | -0.34 |
| tr\|H0YAT2\|H0YAT2_HUMAN 28S ribosomal protein S28, mitochondrial (Fragment) | 1.82 | 5.47 | 3.01 | 1.59 |

| | | | | |
|---|---|---|---|---|
| OS=Homo sapiens OX=9606 GN=MRPS28 PE=1 SV=1 | | | | |
| sp\|Q9H223\|EHD4_HUMAN EH domain-containing protein 4 OS=Homo sapiens OX=9606 GN=EHD4 PE=1 SV=1 | 1.80 | 9.83 | 5.45 | 2.45 |
| tr\|K7ENE3\|K7ENE3_HUMAN Vacuolar protein-sorting-associated protein 25 (Fragment) OS=Homo sapiens OX=9606 GN=VPS25 PE=1 SV=1 | 1.80 | 6.03 | 3.35 | 1.74 |
| tr\|C9JCC6\|C9JCC6_HUMAN Dr1-associated corepressor OS=Homo sapiens OX=9606 GN=DRAP1 PE=1 SV=1 | 1.79 | 7.23 | 4.03 | 2.01 |
| sp\|P56545-2\|CTBP2_HUMAN Isoform 2 of C-terminal-binding protein 2 OS=Homo sapiens OX=9606 GN=CTBP2 | 1.77 | 8.29 | 4.69 | 2.23 |
| tr\|A0A0C4DFN3\|A0A0C4DFN3_HUMAN Monoglyceride lipase OS=Homo sapiens OX=9606 GN=MGLL PE=1 SV=1 | 1.76 | 5.60 | 3.18 | 1.67 |
| sp\|O00186\|STXB3_HUMAN Syntaxin-binding protein 3 OS=Homo sapiens OX=9606 GN=STXBP3 PE=1 SV=2 | 1.74 | 9.45 | 5.44 | 2.44 |
| sp\|O00505\|IMA4_HUMAN Importin subunit alpha-4 OS=Homo sapiens OX=9606 GN=KPNA3 PE=1 SV=2 | 1.73 | 4.12 | 2.38 | 1.25 |
| sp\|Q9H2J4\|PDCL3_HUMAN Phosducin-like protein 3 OS=Homo sapiens OX=9606 GN=PDCL3 PE=1 SV=1 | 1.73 | 5.36 | 3.10 | 1.63 |
| sp\|P48163-2\|MAOX_HUMAN Isoform 2 of NADP-dependent malic enzyme OS=Homo sapiens OX=9606 GN=ME1 | 1.71 | 3.27 | 1.91 | 0.93 |
| sp\|P40692-3\|MLH1_HUMAN Isoform 3 of DNA mismatch repair protein Mlh1 OS=Homo sapiens OX=9606 GN=MLH1 | 1.69 | 7.32 | 4.33 | 2.11 |
| sp\|Q96LD4-2\|TRI47_HUMAN Isoform 2 of E3 ubiquitin-protein ligase TRIM47 OS=Homo sapiens OX=9606 GN=TRIM47 | 1.69 | 3.63 | 2.14 | 1.10 |
| sp\|P80217-2\|IN35_HUMAN Isoform 2 of Interferon-induced 35 kDa protein OS=Homo sapiens OX=9606 GN=IFI35 | 1.67 | 9.38 | 5.62 | 2.49 |
| sp\|Q9NRF9\|DPOE3_HUMAN DNA polymerase epsilon subunit 3 OS=Homo sapiens OX=9606 GN=POLE3 PE=1 SV=1 | 1.66 | 8.59 | 5.18 | 2.37 |
| sp\|Q969U7-2\|PSMG2_HUMAN Isoform 2 of Proteasome assembly chaperone 2 OS=Homo sapiens OX=9606 GN=PSMG2 | 1.62 | 12.36 | 7.65 | 2.93 |
| tr\|J3QLD9\|J3QLD9_HUMAN Flotillin-2 OS=Homo sapiens OX=9606 GN=FLOT2 PE=1 SV=1 | 1.61 | 4.43 | 2.75 | 1.46 |
| tr\|A0A2R8YEM2\|A0A2R8YEM2_HUMAN Cyclin-H OS=Homo sapiens OX=9606 GN=CCNH PE=1 SV=1 | 1.60 | 7.65 | 4.79 | 2.26 |
| tr\|H3BU51\|H3BU51_HUMAN Neuroplastin OS=Homo sapiens OX=9606 GN=NPTN PE=1 SV=1 | 1.59 | 3.99 | 2.51 | 1.33 |

| | | | | |
|---|---|---|---|---|
| sp\|Q9HCC0-2\|MCCB_HUMAN Isoform 2 of Methylcrotonoyl-CoA carboxylase beta chain, mitochondrial OS=Homo sapiens OX=9606 GN=MCCC2 | 1.59 | 3.21 | 2.02 | 1.01 |
| sp\|P61225\|RAP2B_HUMAN Ras-related protein Rap-2b OS=Homo sapiens OX=9606 GN=RAP2B PE=1 SV=1 | 1.58 | 3.93 | 2.49 | 1.31 |
| sp\|Q7L2J0\|MEPCE_HUMAN 7SK snRNA methylphosphate capping enzyme OS=Homo sapiens OX=9606 GN=MEPCE PE=1 SV=1 | 1.57 | 2.71 | 1.72 | 0.78 |
| tr\|A0A1B0GTB8\|A0A1B0GTB8_HUMAN Carnitine O-palmitoyltransferase 2, mitochondrial OS=Homo sapiens OX=9606 GN=CPT2 PE=1 SV=1 | 1.57 | 1.40 | 0.89 | -0.16 |
| sp\|Q9H6F5\|CCD86_HUMAN Coiled-coil domain-containing protein 86 OS=Homo sapiens OX=9606 GN=CCDC86 PE=1 SV=1 | 1.55 | 1.28 | 0.83 | -0.27 |
| sp\|Q6PI78\|TMM65_HUMAN Transmembrane protein 65 OS=Homo sapiens OX=9606 GN=TMEM65 PE=1 SV=2 | 1.55 | 4.00 | 2.59 | 1.37 |
| tr\|G5E9W3\|G5E9W3_HUMAN Cleavage and polyadenylation specific factor 3, 73kDa, isoform CRA_b OS=Homo sapiens OX=9606 GN=CPSF3 PE=1 SV=1 | 1.55 | 4.29 | 2.78 | 1.47 |
| tr\|F5GWE5\|F5GWE5_HUMAN Phosphatidylinositol transfer protein alpha isoform OS=Homo sapiens OX=9606 GN=PITPNA PE=1 SV=1 | 1.53 | 11.43 | 7.48 | 2.90 |
| sp\|Q9NQT5\|EXOS3_HUMAN Exosome complex component RRP40 OS=Homo sapiens OX=9606 GN=EXOSC3 PE=1 SV=3 | 1.51 | 2.56 | 1.70 | 0.77 |
| tr\|A0A2R8Y8A7\|A0A2R8Y8A7_HUMAN Sorting nexin-27 OS=Homo sapiens OX=9606 GN=SNX27 PE=1 SV=1 | 1.50 | 5.36 | 3.58 | 1.84 |
| tr\|A0A0A0MQX8\|A0A0A0MQX8_HUMAN Muscleblind-like protein 1 OS=Homo sapiens OX=9606 GN=MBNL1 PE=1 SV=1 | 1.49 | 8.22 | 5.51 | 2.46 |
| sp\|Q86YP4-2\|P66A_HUMAN Isoform 2 of Transcriptional repressor p66-alpha OS=Homo sapiens OX=9606 GN=GATAD2A | 1.49 | 6.94 | 4.66 | 2.22 |
| sp\|Q9P032\|NDUF4_HUMAN NADH dehydrogenase [ubiquinone] 1 alpha subcomplex assembly factor 4 OS=Homo sapiens OX=9606 GN=NDUFAF4 PE=1 SV=1 | 1.48 | 6.67 | 4.50 | 2.17 |
| sp\|P08243-2\|ASNS_HUMAN Isoform 2 of Asparagine synthetase [glutamine-hydrolyzing] OS=Homo sapiens OX=9606 GN=ASNS | 1.46 | 0.94 | 0.64 | -0.64 |
| sp\|P37235\|HPCL1_HUMAN Hippocalcin-like protein 1 OS=Homo sapiens OX=9606 GN=HPCAL1 PE=1 SV=3 | 1.45 | 5.58 | 3.86 | 1.95 |
| tr\|D6RC74\|D6RC74_HUMAN Protein SDA1 OS=Homo sapiens OX=9606 GN=SDAD1 PE=1 SV=1 | 1.41 | 1.63 | 1.16 | 0.21 |

| | | | | |
|---|---|---|---|---|
| sp\|O15031\|PLXB2_HUMAN Plexin-B2 OS=Homo sapiens OX=9606 GN=PLXNB2 PE=1 SV=3 | 1.40 | 2.20 | 1.57 | 0.65 |
| sp\|Q9NYK5-2\|RM39_HUMAN Isoform 2 of 39S ribosomal protein L39, mitochondrial OS=Homo sapiens OX=9606 GN=MRPL39 | 1.37 | 5.17 | 3.77 | 1.91 |
| sp\|Q06124-2\|PTN11_HUMAN Isoform 2 of Tyrosine-protein phosphatase non-receptor type 11 OS=Homo sapiens OX=9606 GN=PTPN11 | 1.37 | 7.04 | 5.16 | 2.37 |
| sp\|O60678-2\|ANM3_HUMAN Isoform 2 of Protein arginine N-methyltransferase 3 OS=Homo sapiens OX=9606 GN=PRMT3 | 1.36 | 4.72 | 3.48 | 1.80 |
| sp\|O43747-2\|AP1G1_HUMAN Isoform 2 of AP-1 complex subunit gamma-1 OS=Homo sapiens OX=9606 GN=AP1G1 | 1.35 | 2.83 | 2.09 | 1.06 |
| sp\|P49902-2\|5NTC_HUMAN Isoform 2 of Cytosolic purine 5'-nucleotidase OS=Homo sapiens OX=9606 GN=NT5C2 | 1.35 | 8.11 | 6.01 | 2.59 |
| sp\|Q9BV20\|MTNA_HUMAN Methylthioribose-1-phosphate isomerase OS=Homo sapiens OX=9606 GN=MRI1 PE=1 SV=1 | 1.33 | 7.16 | 5.38 | 2.43 |
| sp\|Q9NUQ8-2\|ABCF3_HUMAN Isoform 2 of ATP-binding cassette sub-family F member 3 OS=Homo sapiens OX=9606 GN=ABCF3 | 1.33 | 4.96 | 3.74 | 1.90 |
| sp\|Q9BVI4\|NOC4L_HUMAN Nucleolar complex protein 4 homolog OS=Homo sapiens OX=9606 GN=NOC4L PE=1 SV=1 | 1.31 | 4.02 | 3.07 | 1.62 |
| sp\|Q13451\|FKBP5_HUMAN Peptidyl-prolyl cis-trans isomerase FKBP5 OS=Homo sapiens OX=9606 GN=FKBP5 PE=1 SV=2 | 1.31 | 1.42 | 1.09 | 0.12 |
| tr\|H3BRY6\|H3BRY6_HUMAN Integrator complex subunit 14 OS=Homo sapiens OX=9606 GN=INTS14 PE=1 SV=1 | 1.29 | 1.29 | 1.00 | -0.01 |
| tr\|B0YIW6\|B0YIW6_HUMAN Archain 1, isoform CRA_a OS=Homo sapiens OX=9606 GN=ARCN1 PE=1 SV=1 | 1.29 | 4.60 | 3.56 | 1.83 |
| tr\|H3BND4\|H3BND4_HUMAN Pyridoxal-dependent decarboxylase domain-containing protein 1 OS=Homo sapiens OX=9606 GN=PDXDC1 PE=1 SV=1 | 1.28 | 2.66 | 2.08 | 1.05 |
| tr\|E9PGT3\|E9PGT3_HUMAN Ribosomal protein S6 kinase OS=Homo sapiens OX=9606 GN=RPS6KA1 PE=1 SV=1 | 1.28 | 2.92 | 2.28 | 1.19 |
| sp\|Q14677-3\|EPN4_HUMAN Isoform 3 of Clathrin interactor 1 OS=Homo sapiens OX=9606 GN=CLINT1 | 1.27 | 3.05 | 2.40 | 1.26 |
| sp\|Q8NGA1\|OR1M1_HUMAN Olfactory receptor 1M1 OS=Homo sapiens OX=9606 GN=OR1M1 PE=2 SV=1 | 1.25 | 2.77 | 2.21 | 1.14 |
| tr\|A0A0R4J2F6\|A0A0R4J2F6_HUMAN Interferon-related developmental regulator 2 OS=Homo sapiens OX=9606 GN=IFRD2 PE=1 SV=1 | 1.25 | 3.02 | 2.42 | 1.28 |

| | | | | |
|---|---|---|---|---|
| sp\|Q8TCD5\|NT5C_HUMAN 5'(3')-deoxyribonucleotidase, cytosolic type OS=Homo sapiens OX=9606 GN=NT5C PE=1 SV=2 | 1.24 | 0.17 | 0.14 | -2.84 |
| tr\|A0A1B0GTC3\|A0A1B0GTC3_HUMAN Alpha-ketoglutarate-dependent dioxygenase FTO OS=Homo sapiens OX=9606 GN=FTO PE=1 SV=1 | 1.22 | 5.16 | 4.23 | 2.08 |
| sp\|Q92917\|GPKOW_HUMAN G-patch domain and KOW motifs-containing protein OS=Homo sapiens OX=9606 GN=GPKOW PE=1 SV=2 | 1.22 | 3.41 | 2.80 | 1.49 |
| tr\|G3V158\|G3V158_HUMAN 2-deoxyribose-5-phosphate aldolase homolog (C. elegans), isoform CRA_a OS=Homo sapiens OX=9606 GN=DERA PE=1 SV=1 | 1.21 | 8.76 | 7.26 | 2.86 |
| sp\|Q15291-2\|RBBP5_HUMAN Isoform 2 of Retinoblastoma-binding protein 5 OS=Homo sapiens OX=9606 GN=RBBP5 | 1.19 | 2.33 | 1.96 | 0.97 |
| tr\|H3BM38\|H3BM38_HUMAN Synaptosomal-associated protein (Fragment) OS=Homo sapiens OX=9606 GN=SNAP23 PE=1 SV=1 | 1.19 | 5.46 | 4.60 | 2.20 |
| tr\|A0A1W2PQ47\|A0A1W2PQ47_HUMAN Squalene synthase OS=Homo sapiens OX=9606 GN=FDFT1 PE=1 SV=1 | 1.18 | 7.74 | 6.57 | 2.72 |
| sp\|P53701\|CCHL_HUMAN Cytochrome c-type heme lyase OS=Homo sapiens OX=9606 GN=HCCS PE=1 SV=1 | 1.16 | 4.41 | 3.81 | 1.93 |
| sp\|Q52LJ0-2\|FA98B_HUMAN Isoform 2 of Protein FAM98B OS=Homo sapiens OX=9606 GN=FAM98B | 1.16 | 1.48 | 1.28 | 0.35 |
| tr\|A0A140TA73\|A0A140TA73_HUMAN Beta-2-syntrophin (Fragment) OS=Homo sapiens OX=9606 GN=SNTB2 PE=1 SV=1 | 1.15 | 1.06 | 0.92 | -0.12 |
| sp\|Q969S3\|ZN622_HUMAN Zinc finger protein 622 OS=Homo sapiens OX=9606 GN=ZNF622 PE=1 SV=1 | 1.13 | 4.04 | 3.58 | 1.84 |
| tr\|A0A087WWY3\|A0A087WWY3_HUMAN Filamin-A OS=Homo sapiens OX=9606 GN=FLNA PE=1 SV=1 | 1.13 | 4.08 | 3.62 | 1.86 |
| sp\|P09914-2\|IFIT1_HUMAN Isoform 2 of Interferon-induced protein with tetratricopeptide repeats 1 OS=Homo sapiens OX=9606 GN=IFIT1 | 1.12 | 2.32 | 2.07 | 1.05 |
| sp\|P38432\|COIL_HUMAN Coilin OS=Homo sapiens OX=9606 GN=COIL PE=1 SV=1 | 1.11 | 1.67 | 1.50 | 0.59 |
| sp\|Q9BV44\|THUM3_HUMAN THUMP domain-containing protein 3 OS=Homo sapiens OX=9606 GN=THUMPD3 PE=1 SV=1 | 1.11 | 2.15 | 1.94 | 0.95 |
| sp\|Q9Y6Y0\|NS1BP_HUMAN Influenza virus NS1A-binding protein OS=Homo sapiens OX=9606 GN=IVNS1ABP PE=1 SV=3 | 1.07 | 1.59 | 1.50 | 0.58 |
| sp\|Q8IYS1\|P20D2_HUMAN Peptidase M20 domain-containing protein 2 OS=Homo sapiens OX=9606 GN=PM20D2 PE=1 SV=2 | 1.06 | 2.23 | 2.10 | 1.07 |

| | | | | |
|---|---|---|---|---|
| sp\|Q96D15\|RCN3_HUMAN Reticulocalbin-3 OS=Homo sapiens OX=9606 GN=RCN3 PE=1 SV=1 | 1.04 | 0.20 | 0.20 | -2.35 |
| tr\|A0A1W2PPX5\|A0A1W2PPX5_HUMAN Lysosome membrane protein 2 OS=Homo sapiens OX=9606 GN=SCARB2 PE=1 SV=1 | 1.04 | 3.83 | 3.69 | 1.88 |
| sp\|Q7Z4Q2\|HEAT3_HUMAN HEAT repeat-containing protein 3 OS=Homo sapiens OX=9606 GN=HEATR3 PE=1 SV=2 | 1.04 | 2.34 | 2.26 | 1.18 |
| tr\|D6W592\|D6W592_HUMAN Heterogeneous nuclear ribonucleoprotein L-like, isoform CRA_d OS=Homo sapiens OX=9606 GN=HNRNPLL PE=1 SV=1 | 1.03 | 0.68 | 0.66 | -0.60 |
| sp\|Q9NPQ8-2\|RIC8A_HUMAN Isoform 2 of Synembryn-A OS=Homo sapiens OX=9606 GN=RIC8A | 1.01 | 3.17 | 3.15 | 1.66 |
| sp\|Q08AM6\|VAC14_HUMAN Protein VAC14 homolog OS=Homo sapiens OX=9606 GN=VAC14 PE=1 SV=1 | 1.00 | 2.07 | 2.06 | 1.04 |
| sp\|P53396-2\|ACLY_HUMAN Isoform 2 of ATP-citrate synthase OS=Homo sapiens OX=9606 GN=ACLY | 0.98 | 6.43 | 6.57 | 2.72 |
| tr\|A0A087WW40\|A0A087WW40_HUMAN Endophilin-B1 OS=Homo sapiens OX=9606 GN=SH3GLB1 PE=1 SV=1 | 0.98 | 2.90 | 2.97 | 1.57 |
| sp\|Q9BTE3-2\|MCMBP_HUMAN Isoform 2 of Mini-chromosome maintenance complex-binding protein OS=Homo sapiens OX=9606 GN=MCMBP | 0.97 | 4.76 | 4.92 | 2.30 |
| sp\|Q9Y5A9-2\|YTHD2_HUMAN Isoform 2 of YTH domain-containing family protein 2 OS=Homo sapiens OX=9606 GN=YTHDF2 | 0.97 | 4.22 | 4.37 | 2.13 |
| sp\|Q15061\|WDR43_HUMAN WD repeat-containing protein 43 OS=Homo sapiens OX=9606 GN=WDR43 PE=1 SV=3 | 0.95 | 0.16 | 0.17 | -2.54 |
| tr\|H7BZI9\|H7BZI9_HUMAN ATP-binding cassette sub-family A member 13 (Fragment) OS=Homo sapiens OX=9606 GN=ABCA13 PE=4 SV=1 | 0.95 | 6.81 | 7.16 | 2.84 |
| sp\|P61964\|WDR5_HUMAN WD repeat-containing protein 5 OS=Homo sapiens OX=9606 GN=WDR5 PE=1 SV=1 | 0.95 | 5.12 | 5.39 | 2.43 |
| tr\|E9PKP7\|E9PKP7_HUMAN Nucleolar transcription factor 1 OS=Homo sapiens OX=9606 GN=UBTF PE=1 SV=1 | 0.95 | 3.72 | 3.92 | 1.97 |
| sp\|P50570-2\|DYN2_HUMAN Isoform 2 of Dynamin-2 OS=Homo sapiens OX=9606 GN=DNM2 | 0.95 | 1.56 | 1.65 | 0.73 |
| sp\|Q8IU81\|I2BP1_HUMAN Interferon regulatory factor 2-binding protein 1 OS=Homo sapiens OX=9606 GN=IRF2BP1 PE=1 SV=1 | 0.94 | 3.76 | 4.01 | 2.00 |
| tr\|Q5TDF0\|Q5TDF0_HUMAN Cancer-related nucleoside-triphosphatase OS=Homo sapiens OX=9606 GN=NTPCR PE=1 SV=1 | 0.94 | 3.92 | 4.18 | 2.06 |

| | | | | |
|---|---|---|---|---|
| tr\|M0QZX5\|M0QZX5_HUMAN Acetolactate synthase-like protein (Fragment) OS=Homo sapiens OX=9606 GN=ILVBL PE=1 SV=8 | 0.94 | 2.45 | 2.62 | 1.39 |
| sp\|B5ME19\|EIFCL_HUMAN Eukaryotic translation initiation factor 3 subunit C-like protein OS=Homo sapiens OX=9606 GN=EIF3CL PE=3 SV=1 | 0.90 | 5.97 | 6.61 | 2.72 |
| sp\|Q9UBD5-2\|ORC3_HUMAN Isoform 2 of Origin recognition complex subunit 3 OS=Homo sapiens OX=9606 GN=ORC3 | 0.88 | 2.19 | 2.49 | 1.32 |
| sp\|P50416-2\|CPT1A_HUMAN Isoform 2 of Carnitine O-palmitoyltransferase 1, liver isoform OS=Homo sapiens OX=9606 GN=CPT1A | 0.86 | 0.12 | 0.14 | -2.88 |
| tr\|F5GZF0\|F5GZF0_HUMAN Cyclin-dependent kinase 2-associated protein 1 (Fragment) OS=Homo sapiens OX=9606 GN=CDK2AP1 PE=1 SV=1 | 0.86 | 4.41 | 5.12 | 2.36 |
| tr\|E5RHX8\|E5RHX8_HUMAN Transcription and mRNA export factor ENY2 OS=Homo sapiens OX=9606 GN=ENY2 PE=1 SV=1 | 0.86 | 4.14 | 4.82 | 2.27 |
| tr\|Q5H907\|Q5H907_HUMAN Melanoma antigen family D, 2, isoform CRA_d OS=Homo sapiens OX=9606 GN=MAGED2 PE=1 SV=1 | 0.85 | 5.54 | 6.54 | 2.71 |
| tr\|A0A0D9SG72\|A0A0D9SG72_HUMAN Syntaxin-binding protein 1 OS=Homo sapiens OX=9606 GN=STXBP1 PE=1 SV=2 | 0.84 | 1.13 | 1.34 | 0.42 |
| sp\|Q96TA2-2\|YMEL1_HUMAN Isoform 2 of ATP-dependent zinc metalloprotease YME1L1 OS=Homo sapiens OX=9606 GN=YME1L1 | 0.83 | 6.06 | 7.34 | 2.88 |
| tr\|H0YCA5\|H0YCA5_HUMAN Spermatogenesis-associated protein 5-like protein 1 (Fragment) OS=Homo sapiens OX=9606 GN=SPATA5L1 PE=1 SV=1 | 0.81 | 0.97 | 1.20 | 0.26 |
| sp\|P30837\|AL1B1_HUMAN Aldehyde dehydrogenase X, mitochondrial OS=Homo sapiens OX=9606 GN=ALDH1B1 PE=1 SV=3 | 0.81 | 3.18 | 3.94 | 1.98 |
| sp\|Q96EY7\|PTCD3_HUMAN Pentatricopeptide repeat domain-containing protein 3, mitochondrial OS=Homo sapiens OX=9606 GN=PTCD3 PE=1 SV=3 | 0.81 | 2.66 | 3.30 | 1.72 |
| sp\|Q15172-2\|2A5A_HUMAN Isoform 2 of Serine/threonine-protein phosphatase 2A 56 kDa regulatory subunit alpha isoform OS=Homo sapiens OX=9606 GN=PPP2R5A | 0.80 | 1.96 | 2.46 | 1.30 |
| tr\|H0YJV7\|H0YJV7_HUMAN Transcriptional repressor protein YY1 (Fragment) OS=Homo sapiens OX=9606 GN=YY1 PE=1 SV=1 | 0.79 | 3.35 | 4.25 | 2.09 |
| tr\|H3BPK7\|H3BPK7_HUMAN Alanine--tRNA ligase, cytoplasmic (Fragment) OS=Homo sapiens OX=9606 GN=AARS PE=1 SV=3 | 0.77 | 1.75 | 2.26 | 1.18 |
| tr\|R4GN94\|R4GN94_HUMAN Ankyrin repeat and SOCS box protein 9 (Fragment) OS=Homo sapiens OX=9606 GN=ASB9 PE=1 SV=1 | 0.76 | 2.02 | 2.65 | 1.41 |

| | | | | |
|---|---|---|---|---|
| tr\|I3L387\|I3L387_HUMAN Serine/threonine-protein kinase PLK1 OS=Homo sapiens OX=9606 GN=PLK1 PE=1 SV=1 | 0.75 | 1.62 | 2.16 | 1.11 |
| tr\|A0A024RCR6\|A0A024RCR6_HUMAN HLA-B associated transcript 3, isoform CRA_a OS=Homo sapiens OX=9606 GN=BAG6 PE=1 SV=1 | 0.75 | 1.40 | 1.87 | 0.91 |
| sp\|Q9UKX7-2\|NUP50_HUMAN Isoform 2 of Nuclear pore complex protein Nup50 OS=Homo sapiens OX=9606 GN=NUP50 | 0.72 | 5.18 | 7.19 | 2.85 |
| sp\|Q9NXF1-2\|TEX10_HUMAN Isoform 2 of Testis-expressed protein 10 OS=Homo sapiens OX=9606 GN=TEX10 | 0.71 | 2.85 | 4.02 | 2.01 |
| tr\|A0A0A0MSI8\|A0A0A0MSI8_HUMAN Exocyst complex component 5 OS=Homo sapiens OX=9606 GN=EXOC5 PE=1 SV=1 | 0.70 | 1.99 | 2.84 | 1.51 |
| sp\|Q9UKR5\|ERG28_HUMAN Probable ergosterol biosynthetic protein 28 OS=Homo sapiens OX=9606 GN=ERG28 PE=1 SV=1 | 0.70 | 1.78 | 2.54 | 1.34 |
| tr\|A0A0A0MRE1\|A0A0A0MRE1_HUMAN Exocyst complex component 7 (Fragment) OS=Homo sapiens OX=9606 GN=EXOC7 PE=1 SV=1 | 0.70 | 4.70 | 6.71 | 2.75 |
| tr\|F5H7C2\|F5H7C2_HUMAN Death domain-containing protein CRADD OS=Homo sapiens OX=9606 GN=CRADD PE=1 SV=1 | 0.68 | 2.00 | 2.93 | 1.55 |
| sp\|Q9BT67\|NFIP1_HUMAN NEDD4 family-interacting protein 1 OS=Homo sapiens OX=9606 GN=NDFIP1 PE=1 SV=1 | 0.68 | 0.04 | 0.06 | -4.14 |
| tr\|K7EP32\|K7EP32_HUMAN UBX domain-containing protein 6 (Fragment) OS=Homo sapiens OX=9606 GN=UBXN6 PE=1 SV=1 | 0.67 | 2.17 | 3.23 | 1.69 |
| tr\|H3BP20\|H3BP20_HUMAN Beta-hexosaminidase OS=Homo sapiens OX=9606 GN=HEXA PE=1 SV=1 | 0.65 | 2.67 | 4.12 | 2.04 |
| sp\|Q9UH99-2\|SUN2_HUMAN Isoform 2 of SUN domain-containing protein 2 OS=Homo sapiens OX=9606 GN=SUN2 | 0.62 | 1.35 | 2.18 | 1.12 |
| sp\|P41214-2\|EIF2D_HUMAN Isoform 2 of Eukaryotic translation initiation factor 2D OS=Homo sapiens OX=9606 GN=EIF2D | 0.61 | 0.34 | 0.55 | -0.85 |
| sp\|Q96G46-3\|DUS3L_HUMAN Isoform 3 of tRNA-dihydrouridine(47) synthase [NAD(P)(+)]-like OS=Homo sapiens OX=9606 GN=DUS3L | 0.61 | 2.50 | 4.10 | 2.03 |
| sp\|Q9H1E5\|TMX4_HUMAN Thioredoxin-related transmembrane protein 4 OS=Homo sapiens OX=9606 GN=TMX4 PE=1 SV=1 | 0.61 | 1.62 | 2.66 | 1.41 |
| tr\|A0A087WUL4\|A0A087WUL4_HUMAN Retrotransposon-derived protein PEG10 OS=Homo sapiens OX=9606 GN=PEG10 PE=1 SV=1 | 0.61 | 3.69 | 6.09 | 2.61 |
| tr\|H0UI80\|H0UI80_HUMAN Negative elongation factor C/D OS=Homo sapiens OX=9606 GN=NELFCD PE=1 SV=1 | 0.60 | 1.55 | 2.60 | 1.38 |

| | | | | |
|---|---|---|---|---|
| sp\|Q9H900-2\|ZWILC_HUMAN Isoform 2 of Protein zwilch homolog OS=Homo sapiens OX=9606 GN=ZWILCH | 0.57 | 1.46 | 2.56 | 1.35 |
| sp\|Q15053\|K0040_HUMAN Uncharacterized protein KIAA0040 OS=Homo sapiens OX=9606 GN=KIAA0040 PE=1 SV=2 | 0.56 | 0.60 | 1.07 | 0.10 |
| sp\|Q8N584-2\|TT39C_HUMAN Isoform 2 of Tetratricopeptide repeat protein 39C OS=Homo sapiens OX=9606 GN=TTC39C | 0.54 | 2.68 | 4.93 | 2.30 |
| tr\|K7EP90\|K7EP90_HUMAN RNA-binding protein 42 OS=Homo sapiens OX=9606 GN=RBM42 PE=1 SV=1 | 0.53 | 1.27 | 2.40 | 1.26 |
| sp\|Q9UID3-2\|VPS51_HUMAN Isoform 2 of Vacuolar protein sorting-associated protein 51 homolog OS=Homo sapiens OX=9606 GN=VPS51 | 0.52 | 1.60 | 3.10 | 1.63 |
| sp\|Q99808-2\|S29A1_HUMAN Isoform 2 of Equilibrative nucleoside transporter 1 OS=Homo sapiens OX=9606 GN=SLC29A1 | 0.50 | 1.47 | 2.94 | 1.56 |
| sp\|O75394\|RM33_HUMAN 39S ribosomal protein L33, mitochondrial OS=Homo sapiens OX=9606 GN=MRPL33 PE=1 SV=1 | 0.50 | 2.70 | 5.44 | 2.44 |
| sp\|Q9H773\|DCTP1_HUMAN dCTP pyrophosphatase 1 OS=Homo sapiens OX=9606 GN=DCTPP1 PE=1 SV=1 | 0.50 | 2.20 | 4.43 | 2.15 |
| sp\|Q8IXI1\|MIRO2_HUMAN Mitochondrial Rho GTPase 2 OS=Homo sapiens OX=9606 GN=RHOT2 PE=1 SV=2 | 0.43 | 1.87 | 4.35 | 2.12 |
| sp\|Q27J81-2\|INF2_HUMAN Isoform 2 of Inverted formin-2 OS=Homo sapiens OX=9606 GN=INF2 | 0.43 | 0.84 | 1.96 | 0.97 |
| sp\|P18084\|ITB5_HUMAN Integrin beta-5 OS=Homo sapiens OX=9606 GN=ITGB5 PE=1 SV=1 | 0.41 | 0.86 | 2.08 | 1.06 |
| sp\|Q5T6V5\|QSPP_HUMAN Queuosine salvage protein OS=Homo sapiens OX=9606 GN=C9orf64 PE=1 SV=1 | 0.30 | 2.23 | 7.31 | 2.87 |
| sp\|Q9NUY8-2\|TBC23_HUMAN Isoform 2 of TBC1 domain family member 23 OS=Homo sapiens OX=9606 GN=TBC1D23 | 0.24 | 0.71 | 2.90 | 1.54 |
| tr\|H3BMX9\|H3BMX9_HUMAN PSME3-interacting protein (Fragment) OS=Homo sapiens OX=9606 GN=FAM192A PE=1 SV=1 | 0.22 | 0.73 | 3.37 | 1.75 |
| tr\|A0A2R8YD95\|A0A2R8YD95_HUMAN Vacuolar protein sorting-associated protein 45 OS=Homo sapiens OX=9606 GN=VPS45 PE=1 SV=1 | 0.18 | 1.30 | 7.03 | 2.81 |
| sp\|Q5F1R6-2\|DJC21_HUMAN Isoform 2 of DnaJ homolog subfamily C member 21 OS=Homo sapiens OX=9606 GN=DNAJC21 | 0.13 | 0.27 | 2.01 | 1.01 |
| tr\|A0A0A0MTL5\|A0A0A0MTL5_HUMAN S-phase kinase-associated protein 2 OS=Homo sapiens OX=9606 GN=SKP2 PE=1 SV=1 | 0.12 | 0.73 | 6.10 | 2.61 |

| | | | | |
|---|---|---|---|---|
| sp\|O43819\|SCO2_HUMAN Protein SCO2 homolog, mitochondrial OS=Homo sapiens OX=9606 GN=SCO2 PE=1 SV=3 | 0.10 | 0.79 | 7.52 | 2.91 |
| tr\|Q5TH61\|Q5TH61_HUMAN DnaJ homolog subfamily C member 11 (Fragment) OS=Homo sapiens OX=9606 GN=DNAJC11 PE=1 SV=1 | 0.08 | 0.16 | 1.90 | 0.93 |

*TABLE S 9. MS DATA ACQUISITION PARAMETERS*

| Parameter | Value |
|---|---|
| **Full scan** | |
| Scan range | 350-1700 m/z |
| Micro-scans | 1 |
| Resolution | 60,000 |
| Lock mass | 445.120025 m/z (polysiloxane) |
| Data mode | Profile |
| AGC target | 3.00E+06 |
| Maximum IT | 50ms |
| **dd-MS2** | |
| Micro-scans | 1 |
| Resolution | 15,000 |
| Data mode | Centroid |
| AGC target | 1.00E+05 |
| Maximum IT | 30ms |
| Top N | 12 |
| MSX | no |
| Isolation window | 1.6 m/z |
| Fixed first mass | 140 m/z |
| NCE (no stepped NCE) | 25 |
| Apex trigger | off |
| Charge exclusion | Unassigned, 1 and >8 |
| Peptide match | Off |
| Exclude isotopes | On |
| Dynamic exclusion | 30s |
